# Eph-Ephrin Tetramerization Inhibitors Target Bidirectional Signaling to Combat Pain and Addiction

**DOI:** 10.64898/2025.12.08.692997

**Authors:** Hanghang Wang, Amav Khambete, Frances Sprouse, Miah Henkemeyer, Melody Karsi, Karina Cortez, Eric Bellinghausen, Tamdan Khuu, Natalie Block, Asghar Talebian, Mahmoud Salama Ahmed, Chinthaka Mahesh Udamulle Gedara, Michael Ortiz, Uttam K. Tambar, Patrice N. Mimche, Mark Henkemeyer

## Abstract

Ephs and Ephrins are a large group of interacting membrane-anchored receptor-ligand molecules that form dimers, tetramers, and tetramer superclusters to become activated and signal upon cell-cell contact. While most noted for bidirectional phosphotyrosine signals in development, Ephs/Ephrins also become overexpressed and participate in chronic pathologies, including EphB1 in pain and EphB2 in fibrosis. We searched for small molecules that disrupt EphB-EphrinB receptor-ligand interactions and discovered a large set of 8-hydroxyquinoline compounds that target tetramer-driven EphB1-EphrinB2 and EphB2-EphrinB2 interactions, with less action toward the more dimer-driven EphB4-EphrinB2 interaction. The compounds are orally available, reduce EphB forward signaling and EphrinB reverse signaling, and act to blunt inflammatory pain and opioid withdrawal. Our data indicates 8-hydroxyquinolines, which are known metal-binding compounds, effectively target the high-affinity circular tetramer structure needed for bidirectional signaling.

**SIGNIFICANCE STATEMENT:** We report here and in the accompanying Khambete et al manuscript the discovery of small molecules that disrupt formation of the very high-affinity Eph-Ephrin circular tetramer necessary for activation of bidirectional signaling. We show their potential therapeutic uses here for pain and opioid addiction and in additional manuscripts on diet-induced liver fibrosis and pancreatic cancer cell migration. Altogether, our work indicates metal-binding compounds can exert a powerful effect on Eph-Ephrin interactions and bidirectional cell-cell signaling. This may, in part, help explain why chelation therapy could improve various chronic conditions.

## INTRODUCTION

Studies of Eph receptor tyrosine kinases and their Ephrin ligands have posed many challenges (1). There are 14 receptors and 8 ligands, all membrane-attached, that promiscuously interact upon cell-cell contact to form dimers, tetramers, and tetramer superclusters (2), which become activated to transduce bidirectional phosphotyrosine-mediated signals into both the Eph-expressing cell (forward signaling) and Ephrin-expressing cell (reverse signaling) (Figure 1A). Beyond roles in axon pathfinding, cell migration, and vascular organization, Ephs and Ephrins regulate adult tissue homeostasis, notably in stem cells and neural plasticity (3, 4).

**Figure 1.**
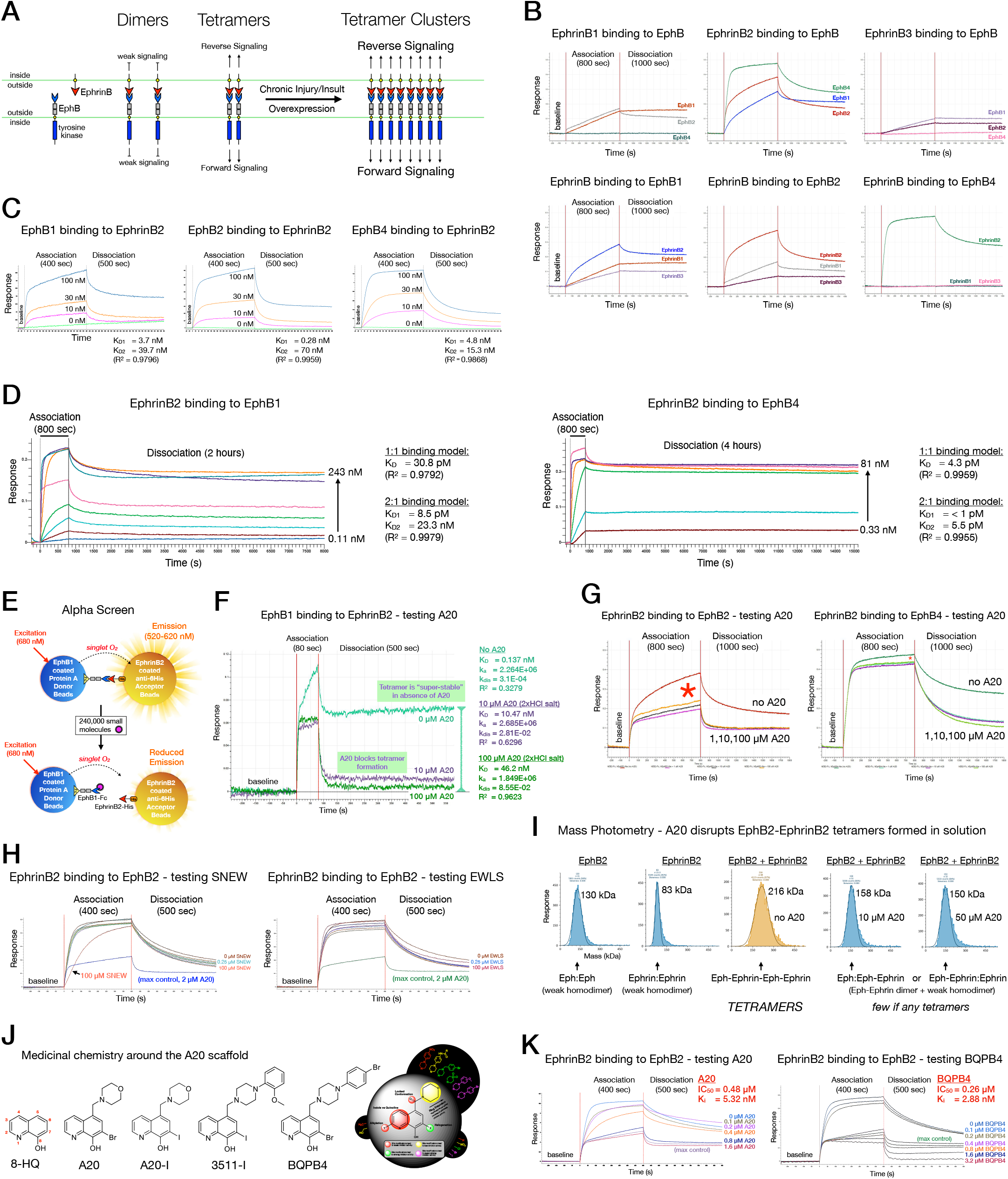
Discovery of small molecules that disrupt the high-affinity Eph-Ephrin tetramer. (A) Transmembrane EphB receptors and EphrinB ligands form dimers, tetramers, and tetramer superclusters at cell-cell contact to activate bidirectional signaling. (B) BLI sensorgram traces of AHC sensorchip immobilized human EphrinB1-Fc, EphrinB2-Fc, and EphrinB3-Fc binding to 50 nM soluble human EphB1-His, EphB2-His, and EphB4-His. (C) BLI of SSA-immobilized biotinylated rat EphB1-Fc, mouse EphB2-Fc, and mouse EphB4-Fc binding to mouse EphrinB2-His; global fit with the 2:1 model gave K_D2_ (dimer) and K_D1_ (tetramer). (D) BLI of AHC2-immobilized human EphrinB2-Fc binding to human EphB1-His or EphB4-His for extended dissociation global fit analysis, with K_D_ values from 1:1 and 2:1 models. (E) Alpha assay with rat EphB1-Fc on Protein A donor beads and mouse EphrinB2-His on anti-6His acceptor beads used for HTS. (F) AHC sensorchip immobilized rat EphB1-Fc binding to 100 nM soluble mouse EphrinB2-His showed A20 (2xHCl salt) reduced tetramer formation. (G) Human EphrinB2-Fc binding to 50 nM human EphB2-His or EphB4-His; A20 (2xHCl salt) reduced tetramer formation, much more for EphB2-EphrinB2 (red asterisks). (H) Human EphrinB2-Fc binding to 100 nM human EphB2-His with peptide SNEW (targets EphB2) or EWLS (targets EphB1) and 2 μM A20 as max control; SNEW reduced the dimer slope at 100 μM (arrow). (I) Mass photometry of human EphB2-His binding to mouse EphrinB2-His with A20 (2xHCl salt). Alone, EphB2-His and EphrinB2-His formed homodimers of ∼130 and 83 kDa, respectively, and when combined assembled into a tetramer complex centered around 216 KDa that shifted to 158 or 150 kDa with A20. (J) Medicinal chemistry identified more potent 8-HQ analogs A20-I, 3511-I, and BQPB4. The schematic highlights changes that show improve (green and purple), poor (yellow), or weak/no (red) activity. (K) Human EphrinB2-Fc binding to 25 nM human EphB2-His with A20 (2xHCl salt) and BQPB4 (free base); area under the curve (AUC) analysis determined relative IC_50_ and K_i_.

Dysregulated Eph/Ephrin signaling is implicated in cancer (5), and in chronic inflammatory/fibrotic/autoimmune/wound healing diseases and metabolic disorders (6). EphB2 and EphrinB2 knockout/knockdown protects against fibrosis of the skin (7, 8), lung (7), kidney (9), liver (10), and heart (11).

Eph/Ephrin dysregulation is also associated with neurological disorders (12). EphB1 and EphB2 are overexpressed in chronic pain (13). EphB2 signals in primary sensory nociceptors by interacting with Ephrin ligands in peripheral target tissues (14), while EphB1 becomes overexpressed and signals in the dorsal horn (DH) neurons within the spinal cord by interacting with EphrinB2 on pre-synaptic nociceptor fibers from the dorsal root ganglion (DRG) (15–20). In both cases, the activated post-synaptic EphB receptor associates with and tyrosine phosphorylates the N-methyl-D-aspartate receptor (NMDAR) (21, 22), allowing calcium influx, strengthening of DH synapses (long-term potentiation, LTP), and central sensitization. People with rheumatoid arthritis show increased synovial fibroblast *EPHRINA3* (*EFNA3*) (23), *EPHRINA5* (*EFNA5*) (24), and *EPHRINB1* (*EFNB1*) (25), those with diabetic neuropathic pain *EPHRINB3* (*EFNB3*) in the DRG (26), and those with cervical neuropathic pain *EPHRINB2* (*EFNB2*) in DRG (27). Nociceptor-specific conditional knockout (KO) studies in mice showed the key role for EphrinB2 in development of inflammatory and neuropathic pain (18).

*EphB1*-/- homozygous germline KO mice, even *EphB1*-/-;*EphB2*-/-;*EphB3*-/- triple KO mice, are viable animals with normal acute pain (28). Like nociceptor EphrinB2 conditional knockouts, *EphB1* KO mice subjected to nerve injury models did not develop enhanced pain sensitivity like wild-type (WT) mice (15, 19, 20). The *EphB1* KO mice further did not generate spinal LTP following C-fiber stimulation of the sciatic nerve in live animals (17). Even *EphB1* +/- heterozygous (HET) mice are refractory to developing pain (15). This indicates a pharmacological strategy to target this receptor would only need to reduce expression/signaling 50%. We therefore searched for small molecules that interfere with EphB1 binding to EphrinB2 and report our discovery of compounds that disrupt formation of the very high-affinity Eph-Ephrin circular tetramer. We show the potential therapeutic uses of Eph-Ephrin tetramerization inhibitors here for pain and addiction, in accompanying Kamdem et al. (29) for diet-induced liver inflammation and fibrosis, and in Bolton et al. (30) to counter pancreatic cancer cell migration. We further present in accompanying Khambete et al. (31) a mechanistic study on how the Eph-Ephrin circular tetramer forms, and how metal-binding compounds can interfere with assembly of this high-affinity macromolecular complex necessary for activation of bidirectional signaling.

## RESULTS

### EphB-EphrinB binding kinetics

Soluble epitope-tagged Eph and Ephrin ectodomains were tested by biolayer interferometry (BLI) on an Octet RED384 or R8e instrument with Studio Analysis 13.1 software (Sartorius). Figure S1A illustrates how BLI immobilizes one protein as “bait” on a biosensor and exposes it to a soluble “prey” in timed association-dissociation steps to assess formation of dimers, tetramers, and tetramer superclusters. Figure 1B shows typical BLI sensorgram data in which human EphrinB1-Fc, EphrinB2-Fc, or EphrinB3-Fc ectodomain proteins were sparsely immobilized onto Anti-Human Fc Capture (AHC) biosensors and, after baseline measurements, exposed to 50 nM soluble human EphB1-His, EphB2-His, or EphB4-His ectodomains for an 800 sec association step followed by a 1,000 sec dissociation step. EphB1 and EphB2 promiscuously interacted with all three EphrinB proteins, whereas cardiovascular-specific EphB4, bound only to EphrinB2 as expected. All three EphB ectodomains exhibited strongest binding to EphrinB2, with EphB4 showing the fastest/strongest binding with a very steep response slope and binding nearly complete within the first 200 sec of the association step, at which point the response quickly flattened. In contrast, EphB1 and EphB2 binding to EphrinB2 occurred more slowly with no sign of a plateau, even at the end of the 800 sec association step. Once placed into dissociation, all three interactions exhibited similar profiles, with some EphB protein coming off the EphrinB2-Fc-bound sensorchips, though ∼50% or more of the receptor remained bound at the end of dissociation, representing very stable complexes (tetramers). To confirm the EphB-EphrinB2 interactions, we biotinylated and immobilized onto Super Streptavidin (SSA) biosensor chips rat EphB1-Fc, mouse EphB2-Fc, and mouse EphB4-Fc ectodomains and assessed binding to 10, 30, and 100 nM soluble mouse EphrinB2-His ectodomain (Figure 1C) and also SSA immobilized biotinylated mouse EphrinB2-His and assessed binding to 30 and 100 nM soluble rat EphB1-Fc ectodomain (Figure S1B). All runs produced similar binding curves regardless of which protein was immobilized or whether human or rodent proteins were used.

Consistent with dimer and tetramer formation, Studio Analysis 13.1 models indicated complex 2:1 interactions rather than simple 1:1 dimer on-off interactions (Figure S1B-1D). With sensorgram data that better fits a 2:1 model, two equilibrium dissociation constants (K_D_) are calculated from two rates of association (k_a_) and two rates of dissociation (k_dis_), K_D2_ for the dimer and K_D1_ for the tetramer. Analysis of the single concentration BLI experiments in Figure 1B confirmed high-affinity binding of all EphrinB-EphB pairs, with K_D_ values in the low nanomolar to picomolar range (Figure S1E), similar to earlier surface plasmon resonance reports (32). SSA biosensor global fit analysis testing three concentrations of prey using the 2:1 model indicated tetramer affinities for EphrinB2-EphB interactions of 0.28-5 nM, with dimer affinities of 10-70 nM (Figures 1C and S1E).

High affinity Eph-Ephrin binding was attributed to a very strong k_a2_ (dimer) and slower k_a1_ (tetramer) rates of association and very slow or unmeasurable off rates of the tetramer (k_dis1_). To gain more accurate kinetic information, especially for the tetramer, the dissociation step was extended to 2 or 4 hr (7,200 or 14,400 sec) and 4-8 prey concentrations were used for global fit analysis. This revealed extremely high affinities for all Eph-Ephrin pairs, including EphrinB2 bound to either EphB1 with a K_D_ = 30.8 pM (1:1 model) and K_D1/D2_ = 8.5 pM/23.3 nM (2:1 model), or EphB4 with a K_D_ = 4.3 pM (1:1 binding model) and K_D1/D2_ = <1 pM/5.5 pM (2:1 binding model) (Figure 1D). BLI analysis of 23 extended global fit experiments using a 2 or 4 hr dissociation step provided rigorous sensorgram data for all three EphrinB ligands binding EphB1, EphB2, EphB4, and EphA4 (Figure S1F) which was analyzed using Studio Analysis 13.1 (Table S1). This revealed extremely stable EphrinB-EphB interactions in the low to mid picomolar K_D_ range, whether using the 1:1 or 2:1 model (Figure S1G). EphA4 bound with higher affinity to EphrinB3 than to EphrinB1 or EphrinB2.

In the above extended dissociation BLI studies, we typically observed an initial fast loss of dimer signal followed by an extremely slow decline that often flatlined. Studio Analysis 13.1 therefore often reported an undetectable tetramer off rate (k_dis1_ < 1 X 10^-7^ s^-1^) and default KD1 values of < 1 pM. We developed our own “dimer:tetramer” mathematical model (see Supplementary Information), and using the same sensorgram data from the 23 extended global fit experiments, found it better captures dimer/ tetramer kinetics (Figure S1H and Table S2). The new model indicates Eph-Ephrin tetramer affinities between 28 pM and 3 nM, and dimer affinities of 6-56 nM, with a tetramer half-life of 6 hours to 7 days, and dimer half-life much shorter from 0.8 to 3.5 minutes. Adding 0.05% BSA blocking agent enhanced tetramer affinity of EphrinB2-EphB2 and EphrinB3-EphB2 interactions >5-fold to 70 pM and 382 pM, respectively (Figure S1I and Table S2). Tetramer half-life correspondingly rose from about 1 day to over 4 days.

### EphB1-EphrinB2 high-throughput screen

We developed sensitive chemiluminescent bead-based Alpha proximity assays in 384 well plates to measure the protein-protein interactions formed between the EphB1 ectodomain and EphrinB1, EphrinB2, and EphrinB3 ectodomains (Figure S2A), selecting the EphB1-EphrinB2 interaction to run a high-throughput screen (HTS) (Figure 1E). The assay was validated with competition experiments using 12-mer peptides EWLS and SNEW that bind within the dimerization pocket of EphB1 or EphB2, respectively, and antagonize EphrinB2 binding (33). EWLS reduced the EphB1-EphrinB2 Alpha signal in a concentration-dependent manner with observed half-maximal inhibitory concentration (IC_50_) of ∼4 µM (Figure S2B), similar to the ∼10 µM reported for the peptide using a different binding assay (33). Soluble Reelin, which can also bind the EphB1 ectodomain (34), was then tested to see if it interfered with the EphB1-EphrinB2 Alpha assay. This revealed a strong concentration-dependent reduction in Alpha signal with an IC_50_ of ∼50 nM Reelin and almost complete inhibition at 200 nM (Figure S2C).

We executed a full HTS at the UT Southwestern HTS Core, assessing at 5 µM a library of ∼240,000 small molecular weight chemicals. The average Z’ was 0.92, indicating a high-quality screen. A total of 1,246 hits that reduced the Alpha signal >10% were identified, of which 950 were retested in the Alpha assay, with positives advancing to additional Alpha, ELISA, and pulldown assays. This identified a single compound (SW056428), called A20 for simplicity, that strongly reduced both EphB1-EphrinB2 and EphB2-EphrinB2 interactions, with much less activity towards the EphB4-EphrinB2 interaction (Figures S2D-S2G).

### Lead compound A20 antagonizes the Eph-Ephrin tetramer

We used BLI to test how A20 affects immobilized EphB1 interacting with 100 nM soluble EphrinB2 for an 80 sec association step (Figure 1F). In the absence of A20, EphB1 bound EphrinB2 in the first 10 sec with an extremely high 87° slope that quickly shifted to a lower 60o slope for the remainder of the association step. Once placed in the 500 sec dissociation step, the slope abruptly reversed for the first 20 sec, indicating a very fast-off species, and then the response quickly flatlined for the remaining time at a level significantly higher than baseline, indicating a very stable complex that did not melt (tetramers). Remarkably, A20 had no effect on the initial 10 sec 87° slope at the start of the association step, though additional binding was lost in a concentration-dependent manner, blunted by 10 µM A20 and flatlined by 100 µM A20. In dissociation, the responses with A20 quickly fell to near baseline levels. A20 thus converts a complex interaction producing stable tetramers into a simple 1:1 dimer-type fast-on/fast-off interaction. The K_D_ calculated for the EphB1-EphrinB2 interaction using 1:1 binding model without A20 indicates a high affinity of 0.137 nM, while with 100 µM A20 the KD was 46.2 nM, a 337-fold weaker affinity, similar to the K_D2_ value for the dimer of 39.7 nM obtained in global fit studies (Figure 1C). A20 did not significantly change the rate of association (k_a_), which is mainly driven by the fast-on dimer species. However, 100 µM A20 caused a striking 275-fold increase in k_dis_, reflecting rapid loss of most EphrinB2 binding to EphB1 in dissociation, consistent with an absence of stable tetramers.

In the absence of A20, shortened association times showed that >15 sec of association is needed before any EphB1-EphrinB2 tetramers can be detected (Figure S3A). This suggests two fast-on/fast-off dimers need 15-20 sec to form before associating into a stable tetramer. A20 also affected immobilized human EphrinB2 binding to soluble human EphB2 and EphB4 testing 0, 1, 10, and 100 µM A20 in longer (800 sec) association times. As with EphB1, A20 reduced tetramer formation during the association step, with a much stronger effect on the EphB2-EphrinB2 interaction than on the EphB4-EphrinB2 interaction (Figures 1G and S3B). Interestingly, A20 inhibition of either interaction maxed out at 1 µM, as 10-fold (10 µM) and 100-fold (100 µM) more compound provided no enhanced inhibition. This indicates the half-maximal inhibitory concentration (IC_50_) for A20 is less than 1 µM. Similar to EphB1, A20 had no effect on the initial very high dimerization slopes observed at the beginning of either the EphB2-EphrinB2 or EphB4-EphrinB2 association step. The reduced effect of A20 on EphB4-EphrinB2 suggests this interaction is mainly dimer-driven.

The SNEW peptide that binds the EphB2 dimerization pocket reduced the early, extreme EphB2-EphrinB2 response slope, confirming an effect on dimer binding (Figures 1H and S3C). Additional studies showed A20 acts as a reversible competitive antagonist of EphB-EphrinB tetramer binding (Figures S3D-S3F), and that A20 can melt pre-existing tetramers (Figure S3G). Mass photometry showed EphB2 and EphrinB2 form tetramers in solution that A20 disrupts (Figure 1I). We synthesized a closely related analog, A20-I (see below), and found it also strongly reduced tetramer formation between EphB1, EphB2, and cross-class EphA4 binding to EphrinB1, EphrinB2, or EphrinB3, again with little effect on the more dimer-driven EphB4-EphrinB2 interaction (Figure S3H and S3I). A global fit of the dimer-driven EphB4-EphrinB2 interaction with A20-I gave a dimer K_D2_ of 19 nM (Figure S3J), similar to the above studies.

A20 further targeted EphB2-EphrinA5 cross-subclass tetramers as well as strict A-subclass tetramers formed between EphA3 and EphrinA1 or EphrinA5 (Figure S3K and S3L). A20 thus targets tetramers formed by most if not all Eph-Ephrin family members, consistent with conserved tetramer interfaces (2, 31).

### Tetramer inhibitors can be improved

In addition to A20, 66 of the top 1,246 primary HTS hits were related to A20 in that they all contained a metal binding 8-hydroxyquinoline (8-HQ) ring and could disrupt Eph-Ephrin tetramers (Figure S4A and Table S3). Medicinal chemistry generated soluble salts and 25 additional novel 8-HQ chemicals (Figure 1J) (Figures S4A-S4G, and Table S4).

We used two BLI methods to characterize tetramer inhibitor activities. The first involved sensorgram area under the curve (AUC) analysis to calculate the percentage of interaction lost upon addition of 3.2 µM of a selected compound during an 80 sec association time. At this max concentration a potent compound will inhibit the majority of tetramers (surmountable), though a low level of insurmountable tetramers can still accumulate, especially in runs with extended association times. For example, 3.2 µM novel compound A20-I shows typical strong tetramer inhibitor activity with greatest ability to reduce the EphB2-EphrinB2 interaction (61.8% inhibited), followed by EphB1-EphrinB2 (36.6%), and least toward the dimer-driven EphB4-EphrinB2 interaction (13.9% inhibited) (Figure S4H).

The second method was to determine the concentration of a compound needed to reduce the protein-protein interaction 50% (IC50), which combined with a known K_D_ in the Cheng-Prusoff equation allowed calculation of the inhibition constant (K_i_), estimating how strongly a competitive inhibitor binds its target. IC_50_ calculations also use AUC information, though here a concentration-response series of sensorgrams for a compound of interest, typically 0.0, 0.1, 0.2, 0.4, 0.8, 1.6, and 3.2 μM, are obtained and compared to a max control (3.2 µM of A20 or A20-I). The AUC of each biosensor response is then used to determine both an absolute IC_50_ and a relative IC_50_. The relative IC_50_ is determined by first subtracting the AUC of the max control run from the AUCs of the concentration-response runs of the compound. This eliminates the contribution of dimer binding and insurmountable tetramer binding (Figure S3B) from the IC_50_ calculation, providing a more accurate determination of a compound’s intrinsic tetramer inhibitor activity. Examples of concentration-response BLI runs and AUC comparisons of novel compounds are provided (Figures S4I and S4J), as are details of a new program, Find50.py, we developed to collect and compare AUCs and calculate IC_50_ and K_i_ values (Figure S4K), presented in Tables S3 and S4. The K_i_ values center on the EphrinB2-EphB2 interaction where the K_D1_ was fixed at 0.28 nM as obtained in early global fit studies in Figure 1C. A particularly potent novel compound, BQPB4, strongly blocked the tetramer with improved IC_50_ and K_i_ values compared to A20 (Figure 1K).

### Tetramer inhibitor pharmacology

Salts of interesting compounds were assessed *in vitro* for solubility, plasma stability, and metabolic stability at the UT Southwestern Preclinicial Core facility, and were generally soluble and stable, with BQPB4 showing superior stability (Figures S5A and S5B). A20 was evaluated for absorption, distribution, metabolism, excretion, and toxicity (ADMET) following a single intravenous (i.v.), intraperitoneal (i.p.), or oral (p.o.) dose, and by all three routes quickly distributed to the plasma, liver, and skin and crossed the blood brain barrier (BBB) to reach the brain and spinal cord (Figures S5C-S5D). Following a single oral dose of the A20 salt, the maximum observed concentration (C_max_) in plasma, brain, and liver was 626 ng/ml, 302 ng/ml, and 19,014 ng/ml, respectively, and terminal half-life (t_1/2_) was 153, 314, and 175 min, respectively. Likewise, single doses of salts of novel compounds A20-I (p.o. dosed; C_max_ in plasma, brain, and liver was 297 ng/ml, 369 ng/ml, and 34,933 ng/ml, and t_1/2_ was 288, 273, and 150 min, respectively) and QPB4-Bn (i.p. dosed; C_max_ in plasma, brain, liver, skin, and spinal cord was 203 ng/ml, 1,566 ng/ml, 6,380, 958 ng/ml, and 660 ng/ml ng/ml, and t_1/2_ was 128, 178, 251, 201, and 385 min, respectively) also indicated favorable ADMET properties, indicating compounds are orally available and maintained concentrations close to their IC_50_ values up to 24 hr post dosing (Figures S5E and S5F). Multiple consecutive dosings of A20, A20-I, 3511-I, and BQPB4 were well tolerated (Figure S5G).

### Tetramer inhibitors target forward signaling

We next investigated the effect of tetramer inhibitors on Eph-Ephrin bidirectional signaling. Focusing on forward signaling mediated by the EphB intracellular tyrosine kinase catalytic domain, rigorous methods were first developed to quantitatively measure a receptor’s activation state. To do this, commercially available antibodies advertised against EphB1 or EphB2 were tested on total cell protein lysates from wild-type (WT) animals to immunoprecipitate (IP) the receptor, which was then visualized by adding ATP in a classic *in vitro* tyrosine kinase autophosphorylation assay (35). Gene-targeted *EphB1* and *EphB2* mutant mice provided control lysates. No antibody detected EphB1, but EphB2 reagents precipitated high levels of active receptor from mid-gestation mouse embryo lysates (Figure 2A) and adult brains (Figure S6A). Following ATP addition, the EphB2 protein IPed from lysates of WT and F620D point mutant (which expresses a dysregulated overactive kinase) animals showed an intense phosphotyrosine signal detected with multiple anti-phosphotyrosine antibodies. Activated, tyrosine phosphorylated pEphB2 migrated slightly slower, just above the 130 kDa standard, distinct from the unphosphorylated, total tEphB2 species, which migrated just below the 130 kDa marker. Key mutant control lysates prove the signals were EphB2 protein: the knockout (KO) animals showed no EphB2 protein, the lacZ truncated mutant expressed a larger ∼250 kDa EphB2-βgal fusion protein lacking the tyrosine kinase domain and showed no phosphorylation, and the K661R kinase-dead point mutation that produces full-length EphB2 also did not become phosphorylated.

**Figure 2.**
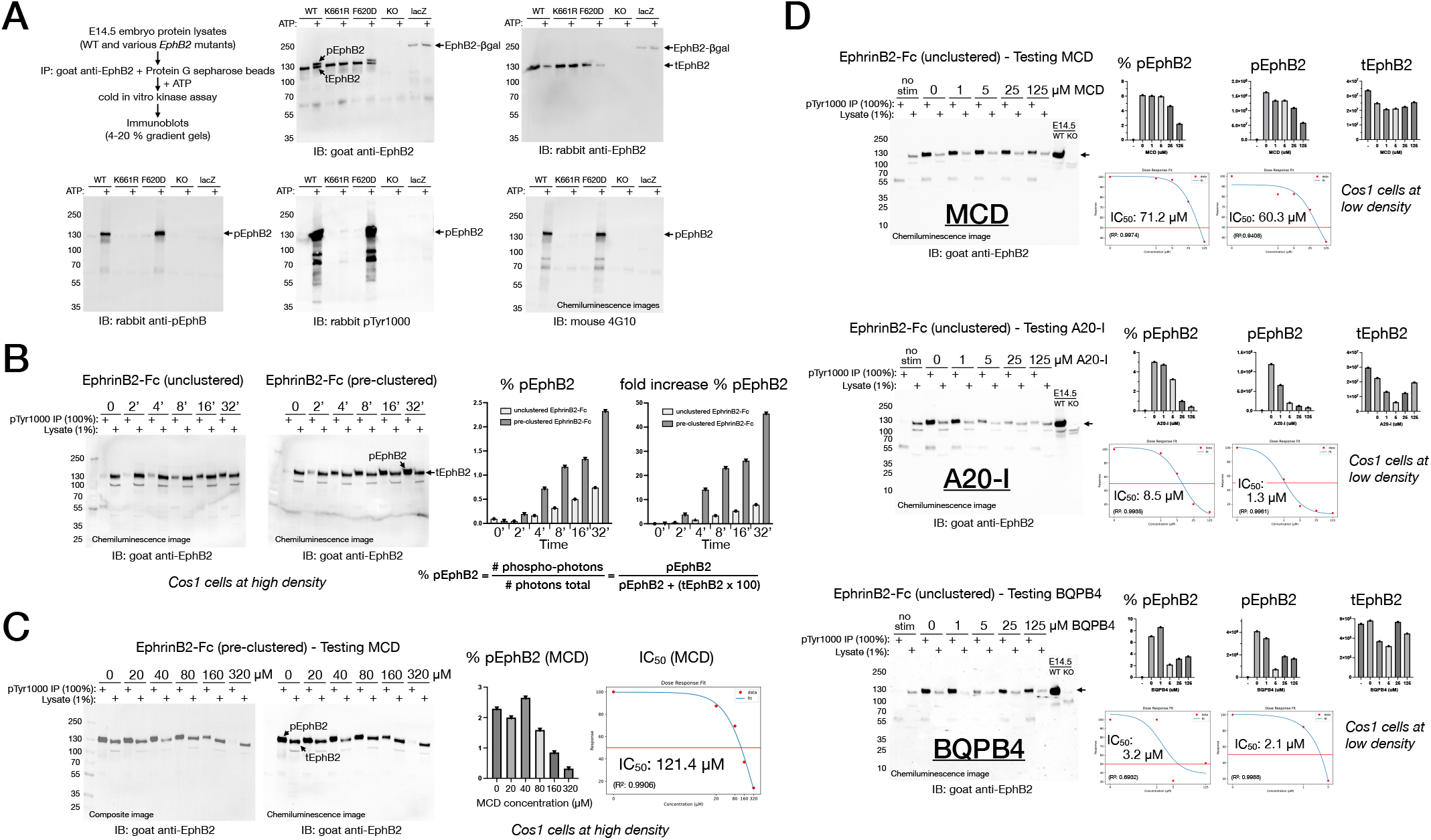
Tetramer inhibitors disrupt EphB2 tyrosine kinase forward signaling *in vitro*. (A) EphB2 immunoprecipitation (IP) from lysates of WT and mutant E14.5 mouse embryos followed by *in vitro* tyrosine kinase assay with ATP. Immunoblots (IB) show a pEphB2 band at or above 130 kDa only in WT and F620D catalytic overactive point mutant lysates. K661R kinase-dead, knockout (KO), and lacZ intracellular truncated EphB2 mutants showed no phosphorylation. Goat anti-EphB2 recognized both pEphB2 and tEphB2, whereas rabbit anti-EphB2 recognized only tEphB2. (B) Anti-pTyr1000 antibody IP from Cos1 cells stimulated with soluble EphrinB2-Fc for indicated times. Goat anti-EphB2 IB shows pEphB2 (IP lanes) migrating slightly slower than tEphB2 (lysate lanes). (C) MCD added to Cos1 cells stimulated with EphrinB2-Fc resulted in a concentration-dependent reduction in % pEphB2. (D) MCD, A20-I, or BQPB4 added to Cos1 cells stimulated with EphrinB2-Fc resulted in concentration-dependent reductions in pEphB2 levels and % pEphB2.

We next developed methods to measure how much EphB2 protein in a lysate is in the active, tyrosine phosphorylated pEphB2 state. Cos1 cells that endogenously express EphB2 were stimulated with EphrinB2-Fc, testing unclustered and pre-clustered soluble ectodomain, and cell lysates were mixed with pTyr1000-conjugated beads and 100% of the precipitated phosphotyrosine-containing proteins were run on a gel next to a lane loaded with 1% of the whole cell lysate taken prior to IP. After immunoblotting for EphB2, band signals corresponding to pEphB2 (pTyr1000 IP lane) and tEphB2 (lysate lane) were used in a simple equation to calculate the percentage of EphB2 in a lysate that is in the tyrosine phosphorylated pEphB2 state (% pEphB2) (Figures 2B and S6B). Unstimulated cells showed <0.1% pEphB2, whereas EphrinB2-Fc caused a time-dependent increase reaching 0.72% (unclustered) and 2.27% (pre-clustered) at 32 min, which was 8-fold and 45-fold higher than the unstimulated cells, respectively. Cos1 cells were then stimulated with pre-clustered EphrinB2-Fc for 32 min in the presence of increasing amounts of a combination of three tetracyclines previously reported to act as a pan EphB tyrosine kinase inhibitor (Minocycline/ Chlortetracycline/Demeclocycline, MCD) (36). This decreased pEphB2 in a concentration-dependent manner with an IC_50_ of 121.4 μM (Figures 2C and S6C). Cos1 cells growing at lower density were then used to test novel compounds A20-I and BQPB4, and compare to MCD. More sparsely plated cells responded much better to EphrinB2-Fc, achieving 5-6% pEphB2 after 32 min of stimulation with the unclustered ectodomain. Addition of increasing MCD again resulted in a concentration-dependent decrease in pEphB2 levels and calculated % pEphB2, here with another relatively poor IC_50_ of 71.2 μM, while addition of A20-I reduced pEphB2 levels and % pEphB2 at much lower concentrations with an IC_50_ of 8.5 μM, and BQPB4 performed even better with an IC_50_ of 3.2 μM (Figures 2D and S6D). A20-I and BQPB4 were thus >8-fold and >22-fold more potent than MCD at reducing % pEphB2.

A20 also affected EphrinB2-Fc binding, clustering, and activation of EphB2 on the cell surface. Cos1 cells on cover slips were stimulated with pre-clustered EphrinB2-Fc for 30 min without or with 40 μM A20 (Figure S6E). A20 significantly reduced the number of spots and the fraction that were phosphotyrosine positive.

We next assessed pEphB2 *in vivo* to examine the effect of tetramer inhibitors in living animals using similar protocols as above, this time with protein lysates of *EphB2* WT and KO mouse embryos collected at E14.5. This revealed a low % pEphB2 (0.6-1.0%) in WT specimens at mid-gestation (Figures 3A and S6F). We then attempted to IP pEphB2 from the adult mouse brain, where the receptor also shows very high expression, but detected no activated protein (Figure 3B). Realizing pEphB2 *in vivo* might be under extreme tight control, brain lysates at different developmental stages revealed the E16.5 embryonic brain contained 1.39% active pEphB2, which gradually decreased through postnatal stages to an almost undetectable 0.09% by 84 days (Figures 3C and S6G).

**Figure 3.**
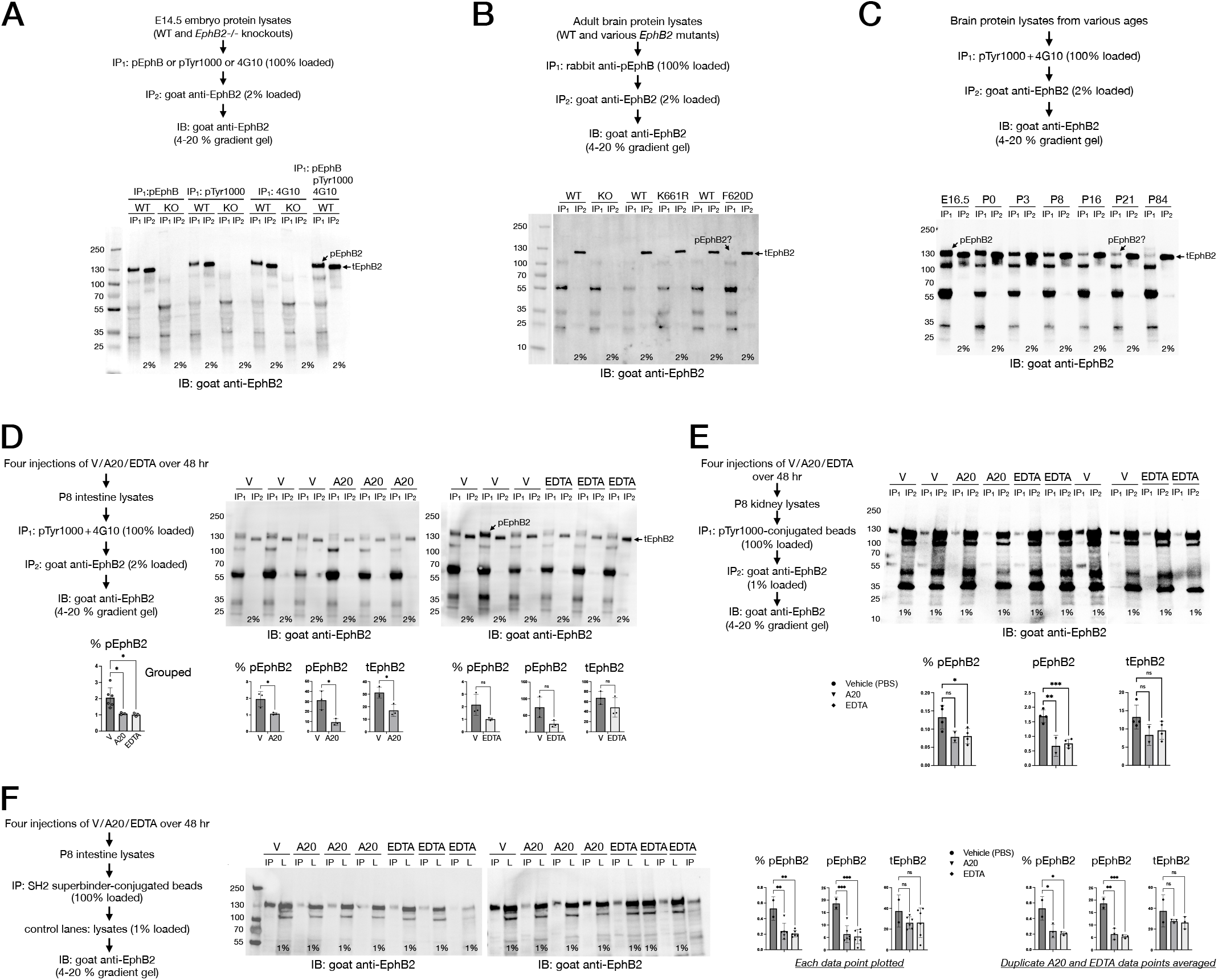
Tetramer inhibitors disrupt EphB2 tyrosine kinase forward signaling when dosed into animals *in vivo.* (A) E14.5 embryo lysates show WT pEphB2 (IP1 lanes) migrating slightly slower than unphosphorylated EphB2 (IP2 lanes). % pEphB2 is ≤1%, and no EphB2 is detected in knockout (KO) animals. (B) No pEphB2 was detected in IP_1_ lanes in adult mouse brain lysates. (C) Whole brain lysates from indicated embryonic (E) and postnatal (P) stages showed pEphB2 gradually diminished with age from 1.39% to almost undetectable. (D) P8 intestines showed reduced pEphB2 levels and % pEphB2 after dosing pups with A20 or EDTA compared to vehicle controls. Unpaired t-tests showed A20, but not EDTA, significantly reduced % pEphB2, pEphB2, and tEphB2; combined one-way analysis of variance (ANOVA) with Dunnett’s post-hoc tests showed significant % pEphB2 reductions for both. Error bars in all graphs are ± standard error of the mean (± SEM) with significance indicated as: ns non-significant; *p < 0.05, **p < 0.01, ***p < 0.001, ****p < 0.0001. Table S5 provides full statistical information for all immunoblot analysis. (E) P8 kidneys showed reduced pEphB2 levels and % pEphB2 in tetramer inhibitor treated animals using one-way ANOVA with Dunnett’s post-hoc test. (F) Tetramer inhibitor treated P8 intestines reduced pEphB2 levels and % pEphB2 using one-way ANOVA with Dunnett’s post-hoc test analyzing individual data points (left) or averages (right).

The barely detectable pEphB2 in adult tissues made it difficult to assess tetramer inhibitors *in vivo*. We therefore turned to postnatal stages and found sufficient pEphB2 in the intestines of vehicle dosed pups collected at P8, and that this was significantly reduced in animals i.p. injected with A20 (Figures 3D and S6H). This experiment also tested ethylenediaminetetraacetic acid (EDTA, tetrasodium salt), finding it also reduced pEphB2 in animals. As described in the accompanying manuscript (31), EDTA is another metal binding, potent Eph-Ephrin tetramer inhibitor. A20 and EDTA were dosed into additional postnatal pups, with kidney and intestine lysates similarly prepared for IP analysis using pTyr1000-conjugated or SH2 superbinder-conjugated beads to precipitate phosphotyrosine-containing proteins. The immunoblots showed both A20 and EDTA led to significant reductions in % pEphB2 as well as highly significant reductions in pEphB2 levels (Figures 3E, 3F, S6I, and S6J).

### Tetramer inhibitors target reverse signaling

For reverse signaling, we used the same strategy to measure phosphorylated pEphrinB2 in mid-gestation embryo lysates, with gene-targeted *EphrinB2* lacZ and 6YFdV mutants as controls (37, 38). *EphrinB2* lacZ expresses a properly localized membrane-anchored EphrinB2-βgal fusion protein lacking its intracellular tail, and 6YFdV is a multiple point mutant in which the six intracellular tyrosine residues are changed to phenylalanine to prevent phosphorylation and coupling to SH2 domain proteins. A small amount of pEphrinB2 was detected in WT but not 6YFdV lysates (Figures 4A and S7A). Quantification of the pTyr1000 bead IP indicated 0.435% of EphrinB2 was in the active tyrosine phosphorylated pEphrinB2 state in mid-gestation embryos. SH2 superbinder beads could also IP activated pEphrinB2 protein from embryo lysates (Figures 4B and S7B). Here, SH2 superbinder beads gave 0.14% pEphrinB2, comparable to 0.13% from pTyr1000 beads.

**Figure 4.**
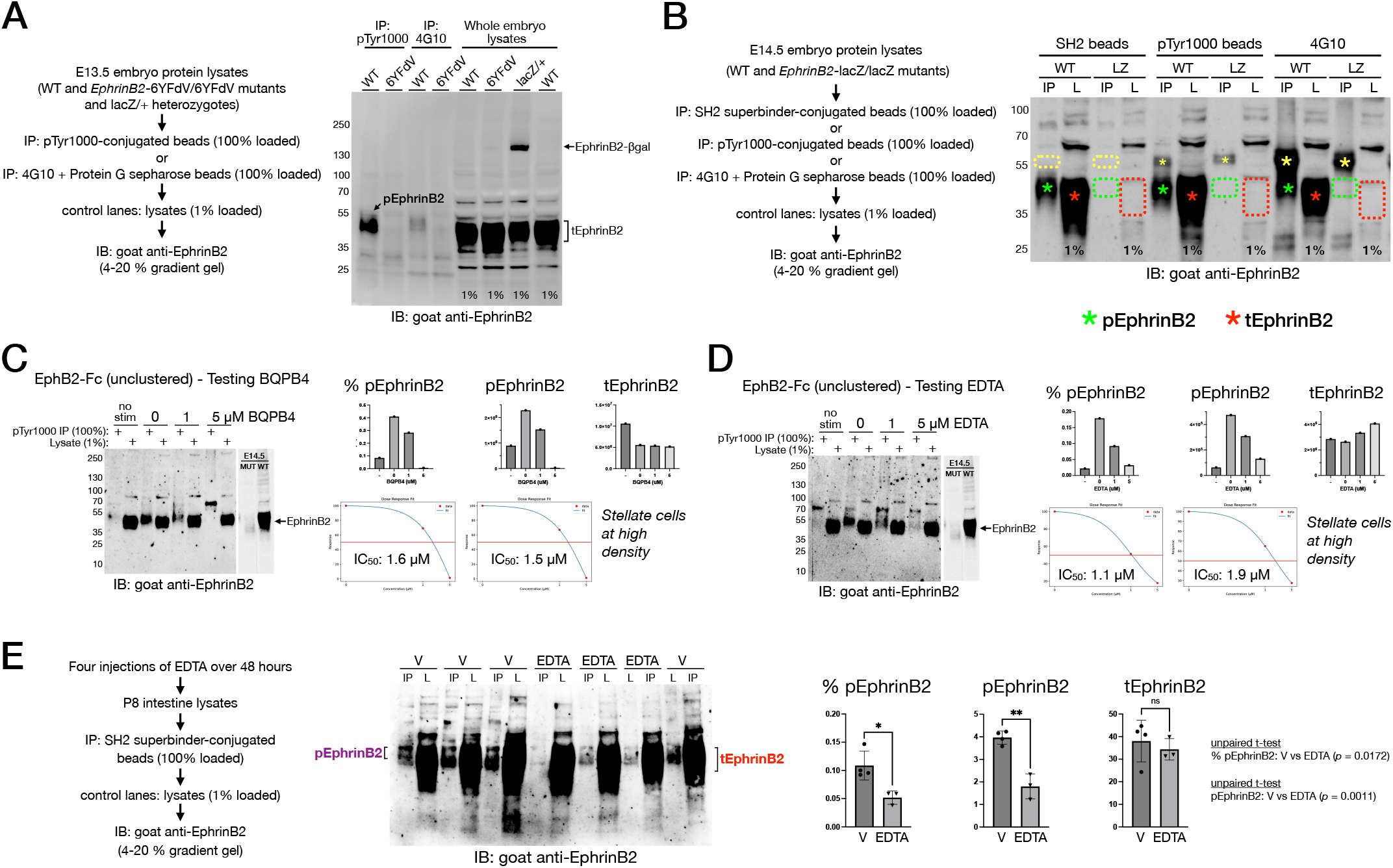
Tetramer inhibitors disrupt EphrinB2 reverse signaling *in vitro* and *in vivo*. (A) pEphrinB2 was precipitated as a 40-50 kDa protein from EphrinB2 +/+ (WT) but not 6YFdV/6YFdV (6YFdV) E13.5 embryo lysates; the lacZ/+ lysate controls for antibody specificity. (B) Activated pEphrinB2 (green asterisks, 40-50 kDa) was detected in *EphrinB2* +/+ (WT), but not in *EphrinB2* lacZ/lacZ (LZ) mutants from E14.5 embryo lysates. Background bands were strong, diminished, or absent with 4G10 Sepharose (large yellow asterisks), pTyr1000-conjugated beads (small yellow asterisks), and SH2 domain-conjugated beads (dotted yellow boxes), respectively. (C) BQPB4 added to pancreatic stellate cells stimulated with EphB2-Fc show a concentration-dependent reduction pEphrinB2. (D) EDTA added to pancreatic stellate cells stimulated with EphB2-Fc resulted in a concentration-dependent reduction in pEphrinB2. (E) EDTA treatment of P8 intestines significantly reduced pEphrinB2 levels and % pEphrinB2 without affecting tEphrinB2 levels using unpaired t-test.

Pancreatic stellate cells that endogenously express EphrinB2 (30) were stimulated with unclustered EphB2-Fc with or without tetramer inhibitor. BQPB4 reduced reverse signaling in a concentration-dependent fashion, shown by reduced % pEphrinB2 (IC_50_ = 1.6 μM) and an overall reduction in pEphrinB2 levels (IC_50_ = 1.5 μM) (Figures 4C and S7C). EDTA exhibited a similar concentration-dependent ability to reduce the % pEphrinB2 (IC_50_ = 1.1 μM) and pEphrinB2 levels (IC_50_ = 1.9 μM) (Figures 4D and S7D). *In vivo* reduction of EphrinB2 phosphorylation by tetramer inhibitor was assessed using lysates from P8 intestines collected from a group of animals dosed with vehicle or EDTA. Compound significantly reduced % pEphrinB2 and pEphrinB2 levels without affecting total tEphrinB2 protein levels (Figures 4E and S7E).

### Tetramer inhibitors blunt inflammatory pain

We next investigated if A20 compounds can counter chronic pain. Intraplantar injection of complete freund’s adjuvant (CFA) into the mouse hind paw is a robust model producing ∼2 weeks of painful swelling with heat (thermal) and touch (mechanical) sensitivity (Figure 5A).

**Figure 5.**
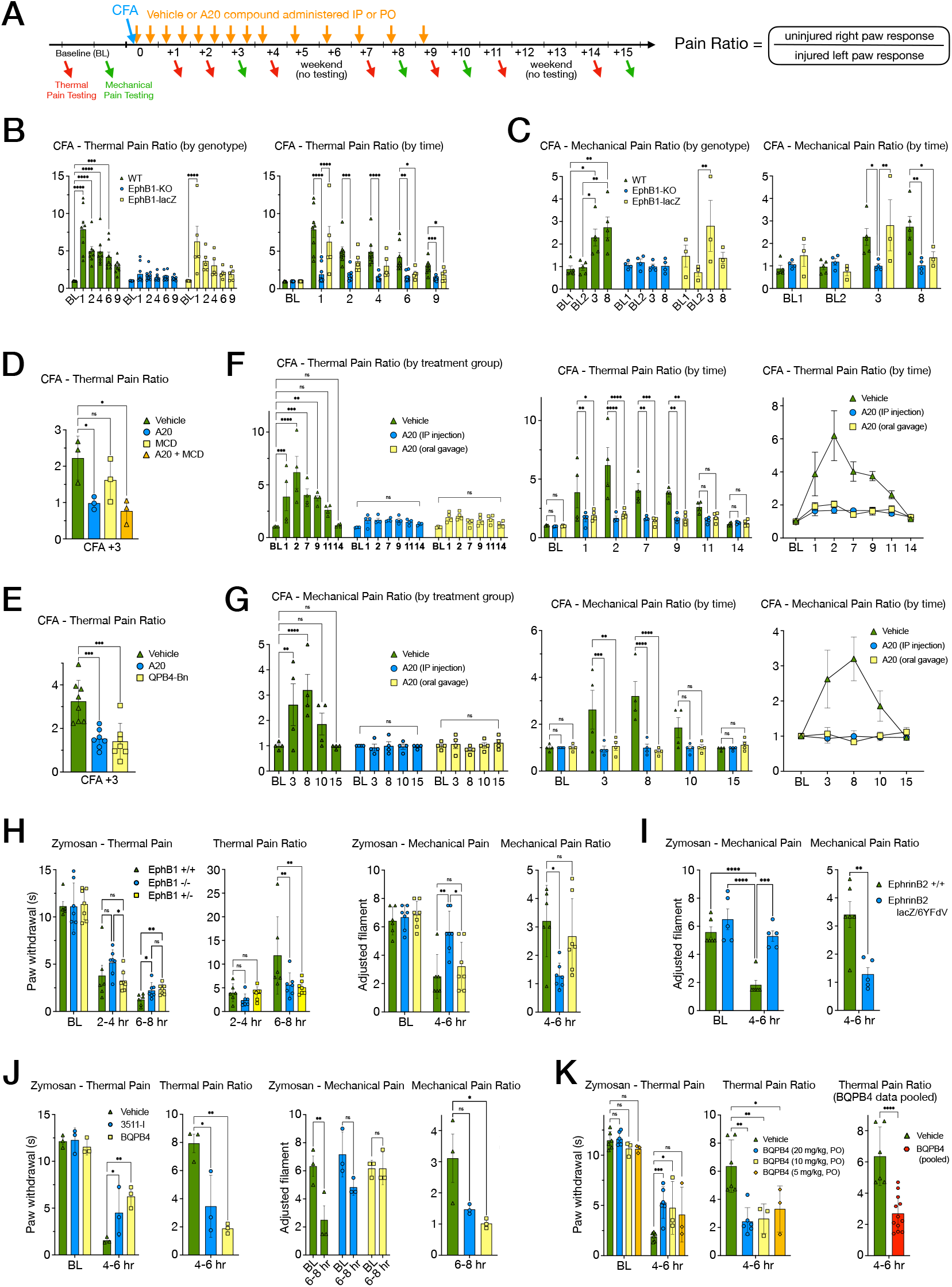
Tetramer inhibitors blunt injury-induced inflammatory pain. (A) General flow of CFA studies. Pain ratios were analyzed by t-tests or ANOVA with post-hoc tests. Error bars in graphs are ± SEM with significance indicated as: ns non-significant; *p < 0.05, **p < 0.01, ***p < 0.001, ****p < 0.0001 (most ns comparisons are omitted from the graphs). Figure S8 and Table S6 provide full statistical information for all pain studies. (B and C) Thermal (B) and mechanical (C) pain ratios obtained on multiple days from CFA-injured littermate-matched *EphB1* +/+ wild-type (WT), -/-knockout (KO), and lacZ/ lacZ intracellular truncated mice. (D-E) Thermal pain ratios +3 days post-CFA in WT mice i.p. injected with: (D) vehicle only, A20 free base (20 mg/kg), MCD kinase inhibitor (10 mg/kg), or both A20+MCD, or (E) vehicle only, A20 salt (20 mg/kg), or QPB4-Bn salt (20 mg/kg). (F and G) Thermal (F) and mechanical (G) pain ratios in WT mice gavaged with vehicle, i.p. injected with A20 salt (20 mg/kg), or gavaged with A20 salt (20 mg/kg) per (A). (H) Thermal and mechanical pain measurements and pain ratios from Zymosan-injured littermate-matched *EphB1* +/+, -/-, and +/-mice. (I) Mechanical pain measurements and pain ratios from Zymosan-injured littermate-matched *EphrinB2* +/+ and lacZ/6YFdV reverse signaling deficient mice. (J) Thermal and mechanical pain measurements and pain ratios from Zymosan-injured WT mice orally dosed with vehicle only, or 3511-I salt (20 mg/kg) or BQPB4 salt (20 mg/kg). (K) Thermal pain measurements and pain ratios from Zymosan-injured WT mice orally dosed with vehicle only or BQPB4 salt.

CFA pain was first assessed in *EphB1* -/-KO (20) and *EphB1* lacZ/lacZ (LZ) mutant mice (39) that have not previously been studied for pain. As the *EphB1* LZ mice express an intracellular truncated EphB1-βgal fusion protein that lacks its tyrosine kinase domain, it is unable to transduce classic forward signals. The EphB1-βgal fusion protein does, however, retain its extracellular, transmembrane, and juxtamembrane segments so it is still membrane localized and able to bind EphrinB proteins to activate reverse signaling, like the EphB2-βgal fusion protein (35). WT mice exhibited long-lasting increased thermal pain in their CFA-injured hind paw (Figures 5B and S8A). The *EphB1* KO mice showed no increased thermal pain over baseline, also as expected and consistent with previous work demonstrating that deletion of this receptor leads to mice refractory to chronic pain (15, 17, 19, 20). Curiously, the *EphB1* LZ mice did not behave like the KOs; they showed a transient, highly significant increase in thermal pain like WT mice, but within a couple of days looked more like KO mice with low thermal pain. At day +1 post-CFA, both WT and LZ mice exhibited strong increases in pain ratios that were significantly different from KO mice, who remained pain free.

However, by days +6 and +9, only WT mice exhibited significant pain, while LZ mice mimicked the KO mice and appeared relatively pain free. Mechanical pain gave similar results, with both WT and LZ mice showing significantly increased touch sensitivity early at +3 days post-CFA, though by day +8 only WT mice continued to experience elevated pain (Figures 5C and S8A). As LZ mice exhibit increased pain early after CFA injury and the truncated EphB1-βgal fusion protein cannot transduce forward signals but can still bind Ephrins and activate reverse signaling, the data suggest reverse signaling may be more important for early stages of chronic pain, while EphB1 forward signaling becomes more important for later stages (see below).

We next tested thermal pain in WT mice i.p. injected with A20, MCD, or A20+MCD, finding animals that received 20 mg/kg of tetramer inhibitor exhibited significantly reduced thermal pain at +3 days post-CFA, whereas the MCD kinase inhibitor had no significant effect (Figure 5D). Other mice i.p. injected with 20 mg/kg of A20 or QPB4-Bn showed significant pain reductions compared to vehicle controls (Figures 5E and S8B). A two week CFA study where A20 was i.p. injected or p.o. oral dosed at 20 mg/kg and mice tested for thermal pain (Figures 5F and S8C-S8D) and mechanical pain (Figures 5G and S8E-S8F) showed compound strongly blunted pain whether injected or orally dosed.

Another inflammatory agent, Zymosan, was used to induce an intense short-lasting increase in thermal and mechanical pain, first testing WT and *EphB1* KO mice (Figures 5H and S8G), revealing that EphB1 loss significantly blunted both elevated heat and touch pain responses compared to WT controls. Reverse signaling mutant mice (*EphrinB2* lacZ/6YFdV), were also highly refractory to zymosan-induced pain (Figures 5I and S8H), indicating reverse signaling through the EphrinB2 intracellular domain is key for inflammatory pain.

Oral dosing of novel A20 tetramer inhibitors 3511-I and BQPB4 at 20 mg/kg significantly blunted pain in the Zymosan model (Figures 5J and S8I). Oral doses of 5, 10, and 20 mg/kg of A20 (Figure S8J) and, to a greater extent, BQPB4 (Figures 5K and S8K) significantly diminished pain, even at the lowest dose.

### Tetramer inhibitors reduce dorsal horn neuron activation

Chronic pain generating nerve injuries/insults overstimulate nociceptor C-fibers whose central projections form synapses with DH neurons in the superficial spinal cord. The excessive firing of C-fibers increases synaptic activity and post-synaptic activation of the NMDAR, leading to calcium influx and downstream signaling that activates DH neurons and triggers plasticity (LTP). Consistent with the post-synaptic localization of EphB1 (28), its role in pain (15, 17, 19, 20), and its ability to associate with the NMDAR via GluN1/NR1 (21) and phosphorylate GluN2/NR2 (22), KO mice exhibit significantly reduced numbers of c-Fos expressing DH neurons after an inflammatory insult (20). We visualized EphB1 stimulation of pain signaling and DH neuron activation using a genetic tool, *Trap2*, that expresses CreERT2 only in activated neurons (40). When combined with the *Ai9 Rosa26-stop-tdTomato* Cre indicator (41) and i.p. injection of 4-hydroxytamoxifen (4-OHT), *Trap2* permanently labels with red fluorescence only neurons activated during a short window (∼8 hr) until the 4-OHT is degraded. Mice carrying *Trap2*/*Ai9* elements and the *EphB1* KO mutation were injected with CFA, and after 4 hr to allow DH neuron activation, were injected with 4-OHT. Two weeks later, lumbar spinal cord sections revealed red neurons activated ∼4-12 hr post-CFA, with ∼3 times more red labeled neurons on the injured versus uninjured side in WT mice, whereas +/-(HET) and -/-(KO) spinal cords showed reduced ratios, with the KO exhibiting a near equal number of neurons on both sides, indicating little, if any, activation of DH neurons in the mutant (Figure 6A). A20 and MCD were then assessed, revealing tetramer inhibitor, but not kinase inhibitor, significantly reduced DH neuron activation (Figure 6B). We then developed methods to rigorously document DH neuron activation in many sections (Figure S9A), including a new computer program, EasyCellCounting.py, that can count and quantify labeled cells (Figure S9B-S9C).

**Figure 6.**
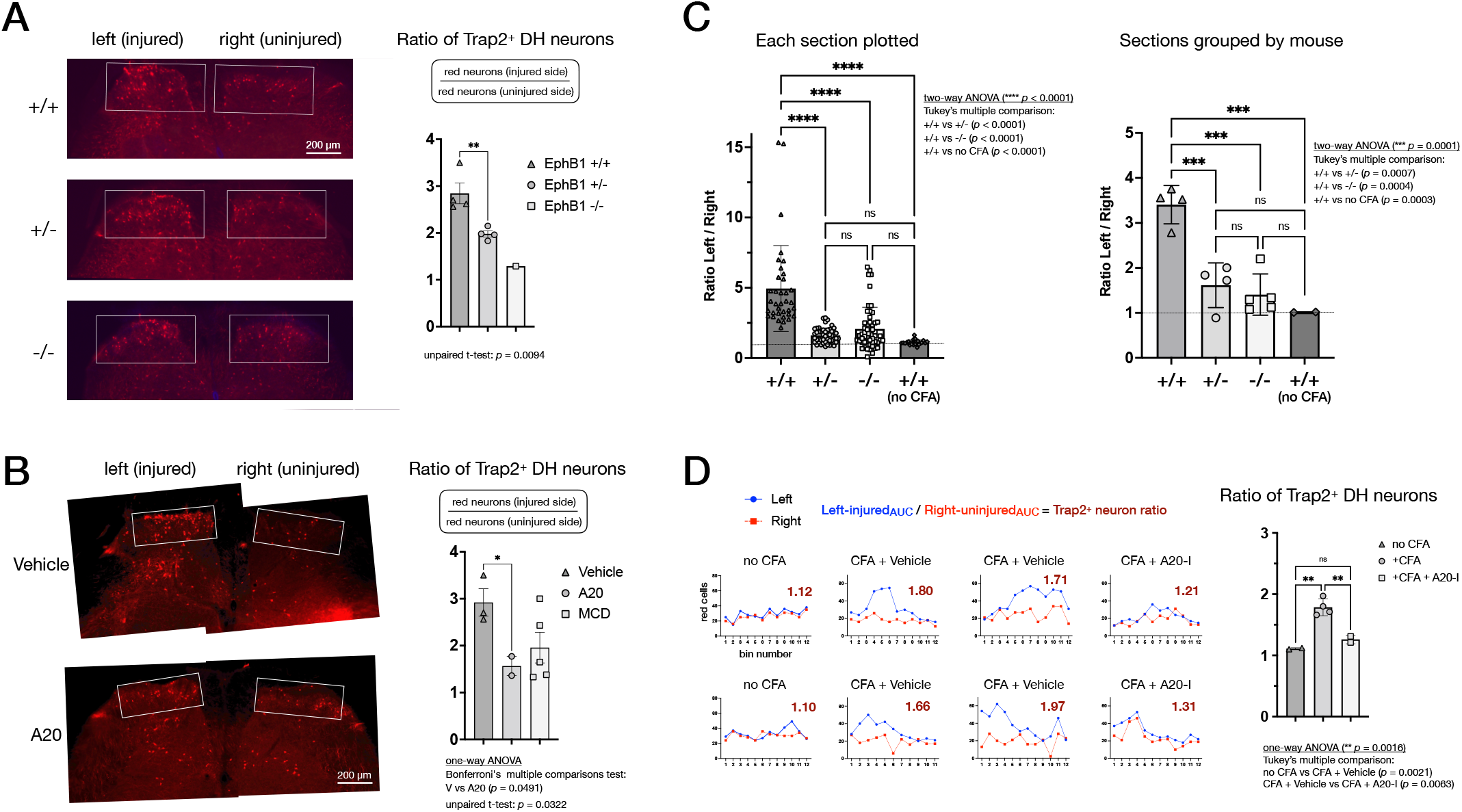
A20 blunts dorsal horn (DH) neuron activation in the superficial spinal cord after injury-induced inflammatory pain. (A) Red fluorescent DH neurons in lumbar spinal cord sections from CFA-injured Trap2/ Ai9-tdTomato *EphB1*+/+, +/-, and -/-littermates. Unpaired t-test shows reduced activation in +/- (HET) versus +/+ (WT). (B) Red fluorescent DH neurons in lumbar spinal cord from CFA-injured WT mice. A20 significantly reduced labeled neurons in the injured side by one-way ANOVA. (C) EasyCellCounting.py counted activated red DH neurons in 160 lumbar spinal cord sections from CFA-injured male and female *EphB1* +/+, +/-, and -/- mice. Plots show the injured/uninjured red neuron ratio per section (left) and per spinal cord (right). Two-way ANOVA shows injured +/+ (WT) animals had a significant >3-fold increase in ratio versus uninjured +/+ controls, whereas injured +/-(HET) and -/-(MUT) animals had highly significant reductions versus injured WT, similar to uninjured controls. (D) CFA-injured WT female mice were studied. Graphs show red DH neurons on the left (blue) and right (red) of one 50 µm section from each of 12 bins spanning ∼6 mm of lumbar spinal cord (10 sections x 50 µm = 0.5 mm per bin). The DH neuron ratio (dark red) is the left (injured) over right (uninjured) AUC. One-way ANOVA showed A20-I significantly reduced DH neuron activation, giving a left/right ratio closer to uninjured controls.

These methods were tested on CFA-injured *EphB1* WT, HET, and KO littermates, demonstrating highly significant differences in neuronal activation can be detected (Figure 6C). We then used these methods to assess A20-I (Figure 6D) and 3511-I and BQPB4 (Figure S9D), showing all three compounds reduced DH neuron activation in the lumbar spinal cord.

### Tetramer inhibitors blunt opioid withdrawal

Like in chronic pain, EphB1 is reportedly upregulated in the dorsal horn of the spinal cord in mice given escalating doses of morphine (42), and similarly dosed *EphB1* -/-KO mice exhibit greatly reduced withdrawal behaviors following administration of naloxone (15). A role for EphB1 in mediating opioid dependence is consistent with its ability to control the NMDAR (21, 22, 28), which is also implicated in mediating the adverse effects of opioid withdrawal (43). We first confirmed the role of EphB1 in withdrawal by subjecting *EphB1* KO and WT littermates to increasing morphine every 12 hr for 5 days. Withdrawal was then precipitated with a subcutaneous injection of naloxone, and animals monitored for 30 min. Major withdrawal events in mice include easily scored jumping, diarrhea, tremor, wet dog shakes, and ptosis. Total jumps per mouse showed that WT mice exhibited significantly more jumps than KO mice (Figure 7A). We then assessed whether i.p. dosing A20 affected withdrawal behaviors in WT mice, using CD1 mice in one experiment and C57BL/6 mice in another. Combined, A20 treated animals exhibited a highly significant reduction in jumps versus vehicle (Figure 7B). The C57BL/6 mice were additionally scored at 5 min intervals for diarrhea, tremor, and ptosis. The tetramer inhibitor treated mice showed reduced incidence of each parameter (Figure 7C).

**Figure 7.**
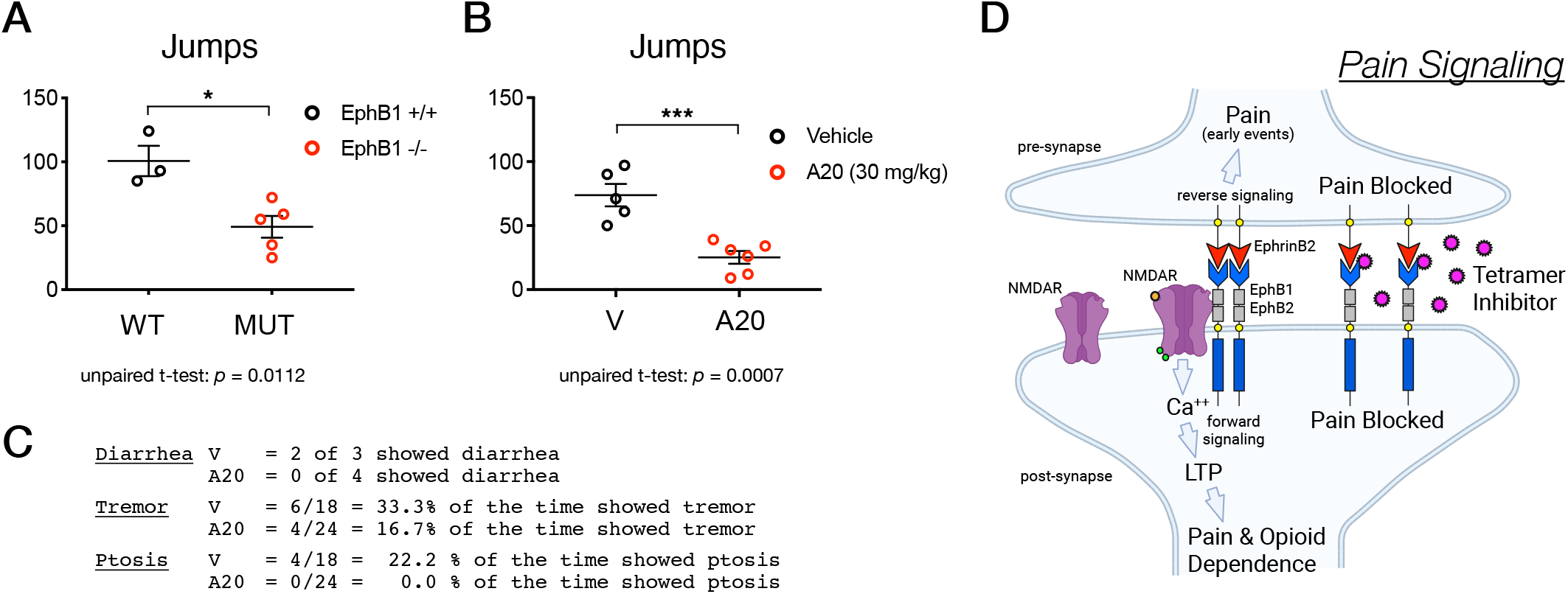
A20 blunts opioid withdrawal behaviors. (A) Morphine withdrawal study of eight month old male *EphB1* +/+ and -/- littermates. The number of jumps is plotted as an indicator of withdrawal intensity. (B) Pooled number of jumps from two opioid withdrawal experiments comparing vehicle to A20 dosed mice, one using CD1 strain mice (2 V and 3 A20) and the other using C57BL/6 strain mice (3 V and 4 A20). (C) Incidence of diarrhea, tremor, and ptosis scored during the withdrawal study using C57BL/6 strain mice in (B). (D) Schematic summarizing how tetramer inhibitors may block synaptic EphB-EphrinB2 forward and reverse signaling to blunt pain and addiction.

## DISCUSSION

We identified and characterized small molecules that disrupt the Eph-Ephrin circular tetramer, providing a novel method to counter chronic diseases associated with excessive Eph-Ephrin signaling, as in pain (Figure 7D). Our drug discovery efforts targeted the interaction of EphB1 with its ligand EphrinB2. The focus was to disrupt the receptor-ligand interaction, as we predicted this would interfere with both EphB forward and EphrinB reverse signaling. This contrasts with the traditional search for a tyrosine kinase inhibitor (like MCD) that would only target Eph catalytic function (36).

Work presented here on pain and on pancreatic cancer cell migration by our London collaborators (30) indicate reverse signaling may be involved in both pathologies and can be countered by A20-based tetramer inhibitors. Regarding the apparent early role we uncovered for nociceptor-expressed pre-synaptic EphrinB2 reverse signaling in chronic pain (Figure 5B, 5C, and 5I), we liken it to the previously described pre-synaptic retrograde signaling by EphrinB proteins in the induction of LTP of hippocampal mossy fiber–CA3 synapses (44). Other attempts to target Eph-Ephrin receptor-ligand interactions have focused on the dimerization interface, like the SNEW and EWLS peptides (33) incorporated in our studies, as well as small molecules (45). However, none of these dimerization inhibitors approach the activities of tetramerization inhibitors, as A20 compounds have submicromolar IC_50_ and low nanomolar K_i_ values, block both forward and reverse signaling, and blunt chronic pain and opioid withdrawal *in vivo*. Our approach started with an HTS of almost 240,000 compounds that identified 67 A20 type 8-HQ chemicals among the top 1,246 hits (Table S3), and an extensive medicinal chemistry campaign that yielded more potent tetramer inhibitors, like BQPB4 (Figure 1K and Table S4).

While the BLI binding kinetics described above for the various EphB-EphrinB interactions are complicated and show clear differences, they are all extremely high affinity, with K_D_ values in the low nanomolar to picomolar range. One striking feature is that only some Eph-Ephrin interactions are tetramer-driven and strongly disrupted by tetramer inhibitors, like EphB1-EphrinB2 and EphB2-EphrinB2, whereas other interactions are more dimer-driven and only slightly disrupted by the tetramer inhibitors, like EphB4-EphrinB2. While the co-crystal of EphB2-EphrinB2 showed these two proteins assemble into a circular tetramer (2), co-crystals and other biophysical studies reported for EphB4-EphrinB2 do not indicate tetramer formation (46, 47).

EphB4 is the only essential EphB receptor, knockout mice are embryonic lethal with cardiovascular malformations (48). EphrinB2 is also a key player in cardiovascular development, with EphrinB2 expressed on arterial cells and EphB4 on venous cells (48, 49). Thus, EphB4-EphrinB2 cell-cell contact in endothelial cells is essential to regulate blood vessel remodeling and integrity and may need to be preserved. As tetramer inhibitors exhibit only mild effects on the EphB4-EphrinB2 interaction, we anticipate they may robustly counter pathological expression and signaling of the tetramer-driven EphB1 and EphB2 receptors while leaving EphB4 interactions intact and minimizing unwanted vascular effects. Indeed, we noticed no negative effects of compound administration in animals, with no signs of endothelial issues at the doses tested. We also uncovered a similar tetramer/dimer difference with A-class Eph/Ephrin molecules, as EphA3 interacts mainly as a dimer with EphrinA5 and as a tetramer with EphrinA1 (Figure S3L).

Our study also uncovered another important concept regarding whether the binding affinity behind assembly of Eph-Ephrin macromolecular complexes is due to dimer or tetramer interactions. While the evidence is not strong, the literature states that Eph-Ephrin dimer binding is the high-affinity interaction that drives assembly. Our studies indicate this notion needs to be reconsidered. The BLI experiment shown in Figure 1F of EphB1-EphrinB2 binding particularly brings this to light, demonstrating A20 effectively disrupts all tetramer formation at 100 μM without any effect on dimer binding kinetics. As such, 100 μM A20 isolates dimer formation, which exhibits simple 1:1 binding kinetics with very fast-on/fast-off dynamics. The K_D_ calculated with 100 μM A20 was 46.2 nM and reflects simple dimer binding, whereas the K_D_ for the full interaction without A20, reflecting both dimer and tetramer binding, was 0.137 nM. These results indicate dimer binding contributes marginally to the overall binding affinity of the interaction, and tetramer formation drives the strength of the macromolecular assembly.

Target specificity is another consideration: a small molecule could be exquisitely specific for a single interaction or hit multiple targets. While our work focuses on EphB1/EphB2 in chronic pain, other Eph receptors may be involved, including EphB3, EphA4, EphA7, and EphA8 (13). An effective anti-Eph/Ephrin therapeutic to counter excessive signaling in chronic pain might need to target not just EphB1 and EphB2 tetramers, but tetramers formed by these other Eph receptors. Likewise, in addition to EphB2, EphA2 and EphA3 interacting with EphrinA1 may also contribute to chronic fibrotic pathologies (50, 51). We show tetramerization inhibitors disrupt tetramers formed by both B- and A-class molecules, as well as cross-class A/B tetramers. Thus, an innovative therapeutic might be a small molecule with broad capabilities to target both A- and B-class tetramers. This may be particularly important because many people suffer from multiple comorbid chronic conditions and may overexpress many of the 14 Eph receptors in different affected organs. In contrast, if tetramer inhibitors with greater selectively are warranted, our ongoing medicinal chemistry campaign indicate compounds with more specific abilities to disrupt either EphB1 or EphB2 tetramers are possible and could be further explored.

## MATERIALS & METHODS SUMMARY

Eph-Ephrin binding was measured by biolayer interferometry (BLI) using an Octet RED384 or R8e with Fc, His, or biotin tagged ectodomains captured on AHC, AHC2, or SSA biosensors in PBS containing 0.05% Tween-20 at 25°C. Kinetic information was alayyzed using Octet Studio Analysis 13.1 for 1:1 and 2:1 binding models, a custom Python dimer:tetramer binding model, and custom Find50.py for AUC-based IC_50_ and Cheng-Prusoff K_i_ values. EphB1-EphrinB2 tetramer inhibitors were identified through Alpha-based screening followed by ELISA and pull-down assays. Cell signaling was assessed by immunoprecipitation and immunoblotting of EphrinB2-Fc stimulated Cos1 cells and EphB2-Fc stimulated pancreatic stellate cells, while levels of activated pEphB2 and pEphrinB2 were obtained using mouse embryo and brain lysates, with tetramer inhibitor effects assessed in postnatal intestines and kidneys. Inflammatory pain was assessed in mice using CFA and Zymosan models with thermal and mechanical sensitivity measurements, while pain-activated spinal cord neurons were quantified with Trap2/Ai9 labeling and custom EasyCellCounting.py. Opioid withdrawal was evaluated following escalating morphine and naloxone. Statistical analyses were performed using GraphPad Prism with t-tests and ANOVA, with sample sizes, controls, and statistical details provided in the Figures, Figure Legends, and Supplementary Information Appendix.

## ACKNOWLEDGEMENTS

We thank our colleagues at UT Southwestern for use of core facilities and technical assistance: Bruce Posner (HTS), Chad Brautigam (Biophysics), Noelle Williams (Pharmacology), and Sherry Birnbaum (Behavior). We also thank Marc Diamond for use of the Octet RED384 instrument and Joachim Herz for Reelin protein. Dean Sherry, Ted Price, Hesham Sadek, Nick Carruthers, Tracy Saxton, Chad Cowan, and Derrick Rossi all provided valuable discussion. The HTS core was supported by NIH awards 2P30CA142543-16 and 1S10OD026758-01, and the mass photometer in Biophysics core by S10OD030312-01. A.K. was supported in part by a Herchel Smith summer fellowship. This research was made possible by the U.S. Department of Defense (DOD) / U.S. Army Medical Research and Development Command (USAMRDC) / Congressionally Directed Medical Research Program (CDMRP) awards W81XWH-14-1-0220 and W81XWH-21-1-0949 (to M.H), U.S. Department of Defense (DOD) / Defense Health Agency Contracting Activity (DHACA) award HT9425-26-1-E064 (to M.H.), NIH award R01DK128819 (to M.H. and P.M.), and an award from the Lyda Hill Philanthropies (to M.H.).

## AUTHOR CONTRIBUTIONS

H.W. performed chemical synthesis, DH neuron analysis, and assisted with other aspects of the study; A.K. performed chemical synthesis, biophysical studies, DH neuron analysis, and wrote computer programs; F.S. was lab manager, preformed biochemistry, DH neuron analysis, and helped with animals; M.H. conducted DH neuron analysis, biophysical studies, and genotyping; M.K. did the HTS screen and early work on A20 chemicals; K.C. conducted the pain behavioral testing; E.B. performed biophysical studies; T.K. performed biochemistry; A.T. conducted the cell clustering experiment; M.S.A. performed chemical synthesis, C.M.U.G. performed chemical synthesis, Alpha assays, and mass photometry; M.O. performed chemical synthesis; U.T. helped guide chemistry; P.M. helped guide biological studies; M.H. raised funds, planned and conducted experiments, supervised and guided the work, and wrote the manuscript with input from the other authors.

