## Supplementary Information Appendix for "Eph-Ephrin Tetramerization Inhibitors Target Bidirectional Signaling to Combat Pain and Addiction"

**This PDF file includes:**

Supplementary Materials & Methods (pages 2 - 13)
Supplementary Figures S1 to S9 (pages 14 - 125)
Supplementary Chemistry Information (pages 126 - 141)
Supplementary Computer Program Information (pages 142 - 143)
Supplementary Tables S1 to S8 (pages 144)

### Materials & Methods

**Biolayer Interferometry**

Eph-Ephrin binding kinetics were analyzed using BLI and either an Octet RED384 or R8e instrument in 384 tilted well plates (Sartorius) and various commercially available recombinant Eph and Ephrin ectodomain proteins that contained C-terminal Fc or His tagged epitopes (sources indicated below). Unconjugated human Fc only protein was also utilize in BLI as negative control and showed no binding in the experiments. Reconstituted ectodomain proteins were routinely passed through Zeba desalting columns before use (Thermo Scientific, 89890). The buffer conditions used were PBS pH 7.4 containing 0.05% Tween 20 (PBST), and plates incubated at 25$^{o}$C with shaking at 1,000 RPM. In most experiments, Fc-tagged Eph or Ephrin ectodomains were first sparsely loaded onto Octet Anti-Human IgG Fc Capture biosensor chips (AHC and AHC2, Sartorius, 18-5060 and 18-5143) using 0.25 - 2.0 µg/ml of the Fc protein, and then after washing and obtaining baseline measurements, the biosensors were placed into wells that contained the desired concentration of soluble interacting analyte partner for the association step for the desired period of time, after which the biosensors were returned to the baseline well for dissociation step. In other experiments ectodomain proteins were sparsely biotinylated using EZ-Link NHS-PEG4-Biotin (Thermo Scientific, A39259) and then passed through Zeba desalting columns before loading onto Octet Super Streptavidin (SSA) Biosensor Chips (Sartorius, 18-5057) and subjecting to baseline, association, and dissociation steps. In experiments that assessed a tetramer inhibitor compound, the baseline and association wells also contained the diluted compound in 1% DMSO final as compounds were dissolved and diluted to 100X final concentrations in DMSO prior to being diluted 1:100 in PBST. For global fit kinetic studies using multiple concentrations of analyte and a long, extended dissociation step, 96 well flat bottom plates were used with an Octet AE Evaporation Cover (Sartorius, 18-5152). Data acquisition was performed using the bundled Octet BLI Discovery 13.1 software to automate the process and generate output files that were then analyzed using Octet Studio Analysis 13.1 software to visualize the sensorgram data and calculate the kinetic parameters of the molecular interaction under study.

**Dimer:Tetramer kinetic binding model computer program**

Due to the very slow dissociation of the circular tetramer and limitations of the Octet Studio Analysis 13.1 software, a new mathematical model was developed in attempts to better analyze dimer:tetramer binding dynamics from global fit BLI sensorgram traces. A multi-step model was developed to account for the binding situation in which a biosensor immobilized Fc-dimerized Ephrin-Fc protein complex when placed into the association step (1) first binds one soluble Eph interacting molecule to form a single fast-on/fast-off heterodimer to produce an increase in response signal, and then (2) subsequently if a second Eph molecule binds the other tethered immobilized Ephrin-Fc molecule to make a second fast-on/fast-off heterodimer, that also leads to an equal sized increase in response signal, and then (3) if the two closely juxtaposed Eph–Ephrin heterodimers close into a 2:2 circular heterotetramer; closure conserves bound mass and produces no change in response. Because the circular tetramer is extremely stable and dissociates far more slowly than heterodimers, the vast majority of the response contributed by tetramer formation persists through to the end of the dissociation.

$$\mathrm{L} + \mathrm{E} \underset{k_{\mathrm{dis2}}}{\overset{k_{\mathrm{a2}}}{\rightleftharpoons}} \mathrm{LE} \tag{1}$$

$$\mathrm{LE} + \mathrm{E} \underset{k_{\mathrm{dis2}}}{\overset{k_{\mathrm{a2}}}{\rightleftharpoons}} \mathrm{LE}_2 \tag{2}$$

$$\mathrm{LE}_2 \underset{k_{\mathrm{dis1}}}{\overset{k_{\mathrm{a1}}}{\rightleftharpoons}} \mathrm{T} \tag{3}$$

where L is the immobilized Fc-dimerized Ephrin-Fc, E is the soluble Eph, LE and $LE_2$ are the Ephrin-Fc with one and two Eph bound, and T is the closed 2:2 heterotetramer.

Accordingly, the binding events were treated as biphasic, with the faster process always assigned to the heterodimer and the slower to the tetramer (assignment by rate, not amplitude). Reference-subtracted sensorgram data was exported from Octet Data Analysis; with individual sensor data excluded if the response signal was low (peaked at ≤ 0.02 nm) or drifted upward during dissociation. Dissociation phases from all retained sensors were fitted globally to a bi-exponential decay (Eq. 4) with shared rates and per-sensor amplitudes, giving $k_{dis2}$, dimer (fast) and $k_{dis1}$, tetramer (slow).

$$R_i(t) = A_{\mathrm{f},i}\mathrm{e}^{-k_{\mathrm{dis2}}t} + A_{\mathrm{s},i}\mathrm{e}^{-k_{\mathrm{dis1}}t} \tag{4}$$

Association phases were fitted globally to a two-phase model (Eq. 5) with observed rates linear in analyte concentration (Eq. 6) and amplitudes free per sensor; $k_{a2}$, dimer (fast) was taken as the slope of the fast observed rate against analyte concentration.

$$R_i(t) = A_{\mathrm{f},i}\left(1 - \mathrm{e}^{-k_{\mathrm{f,i}}t}\right) + A_{\mathrm{s},i}\left(1 - \mathrm{e}^{-k_{\mathrm{s,i}}t}\right) \tag{5}$$

$$k_{\mathrm{f,i}} = k_{\mathrm{a2}}C_i + k_{\mathrm{f0}}, \qquad k_{\mathrm{s,i}} = k_{\mathrm{a1}}C_i + k_{\mathrm{s0}} \tag{6}$$

where $C_i$ is the analyte concentration of sensor i (in M), $k_{f0}$ and $k_{s0}$ are the rate intercepts, and $k_a$ and $k_{dis}$ are in $M^{-1}$ $s^{-1}$ and $s^{-1}$, respectively.

This slope was reported only if it was bounded on both sides by a 5% sum-of-squares profile and at least three sensors carried a fast phase making up ≥ 10% of their signal. Otherwise $k_{a2}$, dimer (fast) was estimated from the top concentration (Eq. 7) and reported as approximate.

$$k_{\mathrm{a2}} = \frac{k_{\mathrm{fast}} - k_{\mathrm{dis2}}}{C_{\mathrm{top}}} \tag{7}$$

Because the slow observed rate depended only weakly on concentration, $k_{a1}$, tetramer was obtained from a two-exponential fit to the highest-concentration trace (Eq. 8), and was reported only when that fit had $R^2 \geq 0.95$.

$$k_{a1} = \frac{k_{slow} - k_{dis1}}{C_{top}} \tag{8}$$

Equilibrium constants were calculated as in Eq. 9 (both in nM), and half-lives were calculated from the dissociation rate constants as in Eq. 10 (dimer in minutes, tetramer in days).

$$K_{D2} = \frac{k_{dis2}}{k_{a2}} \text{ (dimer)}, \quad K_{D1} = \frac{k_{dis1}}{k_{a1}} \text{ (tetramer)} \tag{9}$$

$$t_{½,dimer} = \frac{\ln 2}{k_{dis2}}, \quad t_{½,tetramer} = \frac{\ln 2}{k_{dis1}} \tag{10}$$

All fitting used bounded trust-region least squares in custom Python code (Python 3.13, NumPy 2.3, SciPy 1.16).

**Alpha assays and HTS**

Recombinant rat EphB1-Fc, mouse EphB2-Fc-His, human EphB4-Fc ectodomain proteins, and human Fc only protein as negative control and various EphrinB-His ectodomain proteins were combined with Alpha Protein A Donor Beads (PerkinElmer/Revvity, AS102D) and Alpha Anti-6xHis Acceptor Beads (PerkinElmer/Revvity, AL178C) in 1x AlphaLISA Hiblock Buffer (PerkinElmer/Revvity, AL004C)(25 mM HEPES, pH 7.4, 0.1% Casein, 1 mg/mL Dextran-500, 0.5% Triton X-100, 0.5% Blocking reagent, 0.5%BSA and 0.05% Proclin-300) in 384 well untreated AlphaPlates (PerkinElmer/Revvity, 6057350). The EphB-Fc ectodomain decorated Protein A donor beads and EphrinB-His ectodomain decorated anti-6His acceptor beads were allowed to bind each other and reach equilibrium (>3 hr), after which plates were placed into an EnVision Multilabel Plate Reader (PerkinElmer) to generate singlet oxygen molecules by high energy irradiation of the donor beads which will travel over a constrained distance to excite the acceptor beads and produce a chemiluminescent signal. The basic idea is that Eph-Ephrin binding will bring donor and acceptor beads into close proximity to transfer energy and give a strong signal, whereas a compound from the HTS library that interferes with Eph-Ephrin binding would lead to diminished chemiluminescence. The 240,000 compound chemical library file that was screened in the UT Southwestern HTS Core facility was composed of compounds obtained from various suppliers, including ChemBridge, Chemical Diversity, ComGenex, TimTek, Prestwick, and the NIH phase 1 collection. The chemical library is stored as five identical master sets ([compound] ~10 mM) in uniquely barcoded, 384-well microtiter plates. Three cherry-picking plate sets (384-well plates, [compound] ~5 mM) are derived from one master plate set. Five to six screening masters (384-well plates, [compound]~0.5 mM) are prepared from one of the cherry-picking plate sets. Each screening master set supplies 30 to 50 high throughput screens. All of the plates sets derived from a master plate set are barcoded and tracked using an inventory management system. Copies of the chemical library are stored frozen in DMSO in plates at -80 or -20 °C. The quality of compounds in the library is monitored by tracking the hit rates in experiments and freeze thaw cycles are counted. Copies of the library are archived usually due to deterioration of the plates as a result of repeated resealing rather than any decrease in biological activity of the compounds. Small amounts of lead hits were cherry-picked from master plates and used for confirmation Alpha, ELISA, and pull-down assays. Commercial sources were used for resupply of interesting compounds and to obtain additional available analogs.

**ELISA assays**

Recombinant rat EphB1-Fc, mouse EphB2-Fc-His, human EphB4-Fc ectodomain proteins, and human Fc only protein as negative control were added for 1 hr to 96 well Pierce Protein A Coated Plates (Thermo Scientific, 15154) that had been pre-blocked in 1x AlphaLISA Hiblock Buffer. Wells were then washed in PBST and incubated for 1 hr with soluble mouse EphrinB2-His ectodomain protein in 1x AlphaLISA Hiblock Buffer in the absence or presence of selected concentration of compound diluted in DMSO (1% DMSO final), washed again, reacted with HRP-conjugated 6xHis Tag (C-term) Monoclonal Antibody (3D5) (Thermo Scientific, R931-25), and then visualized using ELISA Pico Chemiluminescent Substrate (Thermo Scientific, 37069) and a luminometer.

**Pull-down assays**

Recombinant rat EphB1-Fc, mouse EphB2-Fc-His, human EphB4-Fc ectodomain proteins, and human Fc only protein as negative control were mixed for 1 hr with Protein G Sepharose beads (Thermo Scientific, 101242) that had been pre-blocked in 1x HiBlock Buffer. Beads were then washed and incubated for 1 hr with soluble mouse EphrinB2-His ectodomain protein in 1x AlphaLISA Hiblock Buffer in the absence or presence of selected concentration of compound diluted in DMSO (1% DMSO final), washed again, and then proteins eluted from the beads in 6X SDS sample buffer, resolved on SDS-PAGE gels, transferred to membrane, and immunoblotted with anti-His or anti-Fc antibodies.

**Chemical synthesis**

Detailed information concerning chemical synthesis is provided in the Supplementary Chemistry Information below.

**Cell stimulations and protein lysates**

Cos1 cells were grown in 6 well culture plates in DMEM medium containing 10% fetal bovine serum, 2 mM L-Glutamine and antibiotics (1% penicillin and streptomycin). Pancreatic Stellate cells were grown in 6 well culture plates in DMEM/F12 medium containing 10% fetal bovine serum and antibiotics (1% penicillin and streptomycin). Serum-starved cells were stimulated with 1.5 ug/ml (30 nM) unclustered or pre-clustered mouse EphrinB2-Fc (to stimulate forward signaling) or mouse EphB2-Fc (to stimulate reverse signaling) for 32 min or other indicated times in protein free DMEM or DMEM/F12 supplemented with 10 mM HEPES. For pre-clustering, selected Fc protein was combined with goat anti-human IgG at a 2:1 ratio for at least 1 hr prior to cell stimulations. Following stimulations, plates were placed on ice, cells washed with cold PBS, and then lysed using 1 ml/well ice cold PLC Lysis Buffer (50mM HEPES (pH 7.5), 150mM NaCl, 1.5mM $MgCl_2$, 10% Glycerol, 1% Triton-X100, 1 mM EGTA, 10 mM

Na4P2O7 .10H2O, 10 mM NaF, 1 mM Na3VO4, 1X Halt Protease Inhibitor Cocktail, 1 mM PMSF). After Nutating the lysate for 30 min in a cold room, the insoluble material was pelleted, and cleared supernatants frozen on dry ice and placed in -80 freezer for long-term storage.

Protein lysates from ground frozen powders of mouse tissues (embryos, brains, intestines, kidneys) were made by dounce homogenization in ice cold PLC Lysis Buffer (3 ml PLC lysis buffer for every 0.1 g of ground tissue powder). After Nutating the lysate for 60 min in a cold room, the insoluble material was pelleted, and 1 ml aliquots of cleared supernatant frozen on dry ice and placed in -80 freezer for long-term storage.

Forward signaling by EphB receptors is driven by activation of their intracellular tyrosine kinase catalytic domain, autophosphorylation of key juxtamembrane tyrosine residues, and coupling to intracellular SH2 and PDZ domain proteins, while reverse signaling by EphrinB ligands involves phosphorylation of conserved tyrosine residues on their short intracellular tail and coupling to SH2 and PDZ domain proteins.

The intracellular juxtamembrane tyrosine residue Y596 of EphB2 receptor becomes autophosphorylated when the catalytic domain is activated by addition of ATP (+), resulting in reduced mobility of ~50% of the protein in SDS-PAGE gels from just under the 130 kDa marker for the unphosphorylated EphB2 protein to a phosphorylated pEphB2 band just at or above the 130 kDa protein marker.

##### *In vitro* kinase assay

Frozen protein lysates (1 ml) prepared in advance from mouse embryo or brain tissues were thawed on ice and 1 ug of goat anti-EphB2 antibody and 20 ul of washed G sepharose bead slurry added to immunoprecipitate the receptor protein. Beads were washed 3X in ice cold HNTGV (20 mM HEPES pH 7.5, 150 mM NaCl, 1% Triton-X100, 10% Glycerol, 1 mM Na3VO4), washed briefly in 1X Kinase Buffer (20 mM HEPES pH 7.5, 10 mM MgCl2, 4 mM MnCl2, 0.1 mM Na3VO4), and then incubated at room temperature RT for 15 minutes in 45 ul of 1X Kinase Buffer containing 200 µM ATP. Kinase reaction was terminated by adding 15 µl 6X SDS Sample Buffer containing DTT and heating at 65°C for 5 minutes, proteins resolved on SDS-PAGE gels, transferred to membranes, and immunoblotted with indicated antibodies.

##### Phosphotyrosine quantification

Frozen protein lysates (1 ml) prepared in advance from desired source (mouse embryo, brain, intestine, kidney, Cos1/stellate cells) were thawed on ice and after removing a small sample (1% - 2% of the whole protein lysates) for the immunoblot lysate lanes either 1 ug of desired anti-phosphotyrosine antibody and 20 ul of washed Protein G Sepharose bead slurry or 20 ul of bead-conjugated pTyr1000 antibody or bead-conjugated SH2 superbinder domain were added and allowed to Nutate in a cold room for 2 hr to immunoprecipitate phosphotyrosine-containing proteins. Beads were washed 3X in ice cold low salt HNTGV (20 mM HEPES, pH 7.5, 50 mM NaCl, 0.02% Triton-X100, 10% Glycerol, 1 mM Na3VO4), and proteins eluted by adding 15 µl 6X SDS Sample Buffer containing DTT and heating at 65°C for 5 minutes, then resolved on SDS-PAGE gels, transferred to membranes, and immunoblotted with indicated antibodies.

The percent level of phosphorylated pEphB2 (% pEphB2) for each lysate was quantified by dividing the number of phospho-specific pEphB2 photons in the IP lane by the total number of tEphB2 photons using the provided equation:

% pEphB2 = # phospho-photons / # photons total

= pEphB2 / (pEphB2 + (tEphB2 x 100)

To assess the effect of tetramer inhibitors, ~2 ul of a 50 mg/ml (equal to a dose of 25 mg/kg) solution of A20, EDTA, or vehicle only (100% PBS) were administered where needed.

**Animal studies**
All animal procedures were approved by the Institutional Animal Care and Use Committee at UT Southwestern Medical Center (IACUC Protocol #2016-101651).

**CFA pain.** In the week prior to beginning a study, mice were first acclimated to the chambers used in pain assessments and their left and right hind paws were subjected to thermal (heat) and mechanical (touch) pain measurements to obtain baseline data. Thermal pain was assessed by placing mice on a glass surface and subjecting their hind paws to an infrared heat source (Hargreaves device, Ugo Basile) that is connected to a timer to measure how long the animal takes to notice the heat and lift the paw. Mechanical pain was assessed by placing mice on a wire grid and repeatably testing hind paws for their sensitivity to a set of 20 von Frey (VF) monofilaments of different stiffnesses (Bioseb Instruments) with the mechanical withdrawal threshold for each mouse hind paw defined as the minimum gauge filament needed to elicit a reflex. In these tests, both hind paws of mice were subjected to multiple measurements over a 2-3 hr period to (1) obtain 6-8 good thermal response measurements which are averaged to provide the data points for heat pain or (2) narrow down the VF filament stiffness threshold to plot the touch pain. Chronic pain was initiated by subcutaneously injecting 30 µl of CFA (Sigma-Aldrich, F5881) into the plantar surface of the left hind paws to induce inflammatory pain. In subsequent days mice were then tested for either thermal or mechanical pain responses as indicated up to 15 days. In all of the experiments, the person doing the actual pain testing of mice was kept blinded to the mouse genotypes (e.g. wild-type or *EphB1* knockout) and/or treatment conditions (e.g. vehicle only or dosed with an A20 chemical). Up to 12 mice were assessed in any one experiment. Thermal pain ratios were calculated by dividing the average response of uninjured right paw by the average response obtained from the injured left paw. Mechanical pain ratios were calculated by dividing the VF response of uninjured right paw by the VF response obtained from the injured left paw.

$$\text{Thermo Pain Ratio} = \frac{\text{the average response of uninjured right paw}}{\text{the averaged response of injured left paw}}$$

$$\text{Mechanical Pain Ratio} = \frac{\text{the VF response of uninjured right paw}}{\text{the VF response of unjured left paw}}$$

To assess tetramer inhibitors for their ability to blunt CFA-induced pain, mice were injected intraperitoneally (i.p.) or administered treatments by oral gavage either vehicle only, or with tetramer inhibitor, or with MCD kinase inhibitor, or with dual compounds.

**Zymosan pain.** Mice were first acclimated to the chambers used in pain assessments and their left hind paws were subjected to thermal (heat) and mechanical (touch) pain measurements to obtain baseline data. On a subsequent day, the plantar surface of the left hind paws were injected subcutaneously with 30 µl of 5 mg/ml Zymosan (Sigma-Aldrich, Z4250) dissolved in PBS and then assessed for thermal pain using Hargreaves device (2-4 hr), for mechanical pain using VF filaments (4-6 hr), and then again for thermal pain (6-8 hr). In some experiments, mechanical pain was assessed first (2-4 hr), followed by tests for thermal pain (4-6 hr), and then again for mechanical pain (6-8 hr). Pain ratios were calculated by dividing the response of left paw obtained during baseline (uninjured) measurements by the response obtained from the same hind paw post-Zymosan (injured).

$$\text{Thermo Pain Ratio} = \frac{\text{the response of left hind paw obtained during baseline (uninjured)}}{\text{the response of the left hind paw post-Zymosan (injured)}}$$

To assess tetramer inhibitors for their ability to bunt Zymosan-induced pain, mice were orally dosed 15 minutes post-Zymosan with either vehicle only, or with tetramer inhibitor. Two hours after Zymosan injection, the injured left hind paws were assessed for mechanical pain (2-4 hr), then mice were orally dosed again with vehicle only or tetramer inhibitors, and then hind paws tested for thermal pain (4-6 hr), and again for mechanical pain (6-8 hr).

**DH neurons.** Mice bred to contain the *Trap2-CreERT2* driver and the *Ai9-tdTomato* reporter were injected with CFA to induce inflammatory pain. Four hours later, mice were then injected IP with 40 mg/kg 4-hydroxytamoxifen (4-OHT) (Hello Bio, HB6040) dissolved in sunflower seed oil to activate Cre recombinase and permanently label with red fluorescence neurons that were activated during the short window of time (~8 hr) until the 4-OHT is degraded. Fourteen days later, animals were perfused with 4% paraformaldehyde dissolved in PBS and then 50 µm thick vibratome serial sections throughout the lumbar spinal cord were obtained, slices mounted, and then viewed under fluorescence microscopy to identify and image red labeled neurons in the superficial dorsal horn of the spinal cords that were activated between ~4-12 hr after the CFA induction of inflammatory pain. Superficial red neurons were counted on the left (injured) and right (uninjured) side dorsal horns from 8-12 sections along the span of lumbar cord of each animal. The ratio of Trap2+ activated neurons for each animal was then determined by dividing the average number of red neurons from the injured side by the number obtained from the uninjured side.

$$\text{Ratio of Trap2+ Activated Neurons for each animal} = \frac{\text{the average number of red neurons from the injured side}}{\text{the average number of red neurons from the uninjured side}}$$

**Morphine withdrawal.** Mice were injected intraperitoneally (IP) with increasing morphine doses in normal saline every 12 hr for 4 consecutive days (day 1: 20 mg/kg, day 2: 40 mg/kg, day 3: 60 mg/kg, day 4: 80 mg/kg) and on the fifth day in the morning they were injected with a final 100 mg/kg morphine dose which was staggered by one mouse every 30 minutes. Exactly 3 hr after the last morphine injection, each mouse was weighed, and withdrawal precipitated using naloxone hydrochloride that was subcutaneously injected at 1 mg/kg. Withdrawal signs were then monitored by an examiner who was blinded to animal genotypes or treatment groups for 30 min after naloxone administration to identify the number of jumps, wet dog shakes, and diarrhea events observed in the 30 min period, and % change in weight from before naloxone administration to 30 min post withdrawal. For tremor and ptosis, the presence of tremor and/or a drooping upper eyelid is monitored at the beginning of each 5 min interval during the 30 min monitoring period.

**Statistics and reproducibility**

Statistical analysis (t-test, one-way ANOVA, and two-way ANOVA) was done using GraphPad software v.10.6.1. The tests used are noted in the figure legends. All tests are two sided and unpaired. A significance level of 0.05 was used to reject the null hypothesis. The sample size, age and sex of experimental animals, as well as the summary of each statistical test (including degrees of freedom, confidence intervals and P values) can be found in the Figures, associated legends, Supplementary Figures, and Tables S5-S8. All animal experiments were done using biologically independent samples (individual animals) with each data point plotted on scatter graphs unless noted. Representative fluorescent images shown are from one animal per group, selected from multiple animals that consistently showed similar results.

**$IC_{50}$ computer program**

**Find50.py** Was developed to analyze biophysical kinetic data generated by the Octet RED384 instrument and calculate (1) the % of a protein-protein interaction that is inhibited at a given concentration of an inhibitor and (2) the $IC_{50}$ value of an inhibitor using area under the curve (AUC) analysis. Relative $IC_{50}$ values were calculated by subtracting the AUC of the max control sensorgram from the AUCs of the concentration-response runs, eliminating the contribution of dimer binding and insurmountable tetramer binding for a more accurate determination of a compound's tetramer inhibitor activity. Find50.py can also be utilized with other types of dose/concentration-dependent data that generates curves needing AUC analysis and $IC_{50}$ calculations. If an accurate $K_D$ is known, FIND50.py will further use the calculated $IC_{50}$ value and concentration of the soluble protein used to determine $IC_{50}$ [L], to calculate the $K_i$ (inhibition constant) using the Cheng-Prussof equation: $K_i = IC_{50}/(1+ [L]/K_D)$. The $K_i$ is a dissociation constant specific for the protein-inhibitor complex; a lower value indicates a stronger binding affinity as it indicates a lower concentration of the competitive inhibitor is needed to occupy 50% of its target protein. This equation defines the theoretical relationship between the $IC_{50}$ and $K_i$ of an inhibitory compound based on the $K_D$ of the interaction it is inhibiting and provides a useful constant to describe a specific inhibitor's activity towards its target.

**Cell counting computer program**

**EasyCellCounting.py** Was developed to count and quantify labeled cells from microscope images within a selected area of interest to the end user. EasyCellCounting.py can be used with precision for any type of cell labeling system, as long as signal-to-noise ratios are discernible over background levels, and it is particularly useful for counting images of fluorescent labeled cells. EasyCellCounting.py was developed to replace the labor-intensive manual process of counting Tom+ red fluorescent neurons in the spinal cord with a highly optimized and efficient program, though the program should be of broad use to anyone needing to count and quantify cell numbers no matter what was used to label them. Further, EasyCellCounting.py eliminates bias and increases the overall precision of counting cells compared to manual counting.

**Proteins**

**Recombinant Eph/Ephrin ectodomain proteins:**

Human Fc protein (R&D Systems, 110-HG-100). Is a 26.6 kDa protein (with human Fc Pro100-Lys330) that migrates as a 30-35 kDa protein in SDS-PAGE under reducing conditions.

Rat EphB1-Fc protein (R&D Systems, 1596-B1-200). Is an 85 kDa Fc fusion protein (with rat EphB1: Met18-Gln538) that migrates as a 102 kDa protein in SDS-PAGE under reducing conditions.

Mouse EphB2-Fc-His protein (R&D Systems, 467-B2-200). Is an 85 kDa Fc fusion protein (with mouse EphB2: Val27-Lys548) that migrates as a 100-110 kDa protein in SDS-PAGE under reducing conditions.

Human EphB4-Fc (R&D Systems, 11307-B4). Is an 85.2 kDa fusion Fc protein (with human EphB4: Leu16-Arg539) that migrates as a 95-108 kDa protein in SDS-PAGE under reducing conditions.

Human EphA3-Fc (R&D Systems, 6444-A3-050). Is an 85.4 kDa Fc fusion protein (with human EphA3: Met1-Gln541) that migrates as a 100-110 kDa protein in SDS-PAGE under reducing conditions.

Human EphrinB1-Fc (R&D Systems, 7654-EB-050). Is a 49.5 kDa Fc fusion protein (with human EphrinB1: Leu28-Ser236) that migrates as a 62-67 kDa protein in SDS-PAGE under reducing conditions.

Mouse EphrinB2-Fc-His protein (R&D Systems, 496-EB-200). Is a 49.6 kDa Fc-His fusion protein (with mouse EphrinB2: Arg27-Ala227) that migrates as a 60-65 kDa protein in SDS-PAGE under reducing conditions.

Human EphrinB2-Fc (R&D Systems, 7397-EB-050). Is a 48.8 kDa Fc fusion protein (with human EphrinB2: Met1-Ala229) that migrates as a 58-66 kDa protein in SDS-PAGE under reducing conditions.

Human EphrinB3-Fc (R&D Systems, 7924-EB-050). Is a 48.3 kDa Fc fusion protein (with human EphrinB3: Leu28-Ser224) that migrates as a 57-61 kDa protein in SDS-PAGE under reducing conditions.

Human EphB1-His (Sino Biologicals, 11963-H08H). Is a 60 kDa His fusion protein (human EphB1: Met1-Pro540) that migrates as a 66 kDa protein in SDS-PAGE under reducing conditions.

Human EphB2-His (Sino Biologicals, 10762-H08H). Is a 59.7 kDa His fusion protein (human EphB2: Met1-Leu543) that migrates as a 70 kDa protein in SDS-PAGE under reducing conditions.

Human EphB4-His (Sino Biologicals, 10235-H08H). Is a 58.5 kDa His fusion protein (human EphB4: Met1-Ala539) that migrates as a 72 kDa protein in SDS-PAGE under reducing conditions.

Mouse EphrinB1-His protein (Sino Biologicals, 50580-M08H). Is a 23.3 kDa His fusion protein (with mouse EphrinB1: Met1-Ser229) that migrates as a 33-38 kDa protein in SDS-PAGE under reducing conditions.

Mouse EphrinB2-His protein (Sino Biologicals, 50598-M08H). Is a 23.5 kDa His fusion protein (with mouse EphrinB2: Met1-Ala232) that migrates as a 30-40 kDa protein in SDS-PAGE under reducing conditions.

Rat EphrinB2-His protein (Sino Biologicals, 80107-R08H). Is a 23.5 kDa His fusion protein (with rat EphrinB2: Met1-Ala229) that migrates as a 33-39 kDa protein in SDS-PAGE under reducing conditions.

Human EphrinB2-His protein (Sino Biologicals, 10881-H08H). Is a 23.5 kDa His fusion protein (with human EphrinB2: Met1-Ala229) that migrates as a 35-40 kDa protein in SDS-PAGE under reducing conditions.

Human EphrinA1-His protein (Sino Biologicals, 10882-H08H). Is a 20.8 kDa His fusion protein (with mouse EphrinA1: Met1-Ser182) that migrates as a 26 kDa protein in SDS-PAGE under reducing conditions.

Human EphrinA5-His protein (Sino Biologicals, 10192-H08H). Is a 23 kDa His fusion protein (with mouse EphrinA1: Met1-Asn203) that migrates as a 27 kDa protein in SDS-PAGE under reducing conditions.

**Primary Antibodies and phosphotyrosine detection reagents:**
Goat anti-EphB2 antibody (R&D Systems, AF467). Polyclonal antibody raised against mouse EphB2 ectodomain Val27-Lys548. Recognizes both unphosphorylated EphB2 band (<130 kDa) and tyrosine phosphorylated pEphB2 (>130 kDa).

Rabbit anti-EphB2 antibody [D2X2I] (Cell Signaling, 83029). Monoclonal antibody raised against an unphosphorylated intracellular peptide corresponding to juxtamembrane residues surrounding Ala573 of human EphB2 protein, just after the TM segment. Only recognizes the unphosphorylated EphB2 protein (<130 kDa).

Rabbit anti-phospho-EphB1/2 (Abcam, 61791). Polyclonal antibody raised against the conserved tyrosine phosphorylated intracellular juxtamembrane peptide at residue pY594 in EphB1 and pY596 in EphB2 protein (pYIDP). Only recognizes the tyrosine phosphorylated EphB2 protein (>130 kDa).

Goat anti-EphrinB2 (R&D Systems, AF496). Polyclonal antibody raised against mouse EphrinB2 ectodomain Arg27-Ala227. Recognizes both unphosphorylated EphrinB2 in a broad band (35-50 kDa) and tyrosine phosphorylated pEphrinB2 (40-50 kDa)..

Rabbit anti-phosphotyrosine MultiMab™ Rabbit mAb mix (pTyr1000) (Cell Signaling, 8954). A mixture of rabbit monoclonal antibodies that recognizes a broad range of tyrosine-phosphorylated proteins and peptides, and does not cross-react with proteins or peptides containing phospho-Ser or phospho-Thr residues.

4G10® Platinum anti-phosphotyrosine Antibody (mouse monoclonal cocktail IgG2b) (Millipore Sigma, 05-1050). A mixture of mouse 4G10 and PY20 monoclonal antibodies that recognizes a broad range of tyrosine-phosphorylated proteins.

Rabbit anti-phosphotyrosine MultiMab™ Rabbit mAb mix (pTyr1000 / Sepharose Bead Conjugate) (Cell Signaling, 14005). A mixture of rabbit monoclonal antibodies that recognizes a broad range of tyrosine-phosphorylated proteins and peptides conjugated to sepharose beads for use in immunoprecipitations.

SH2 Superbinder (SH2S) Agarose Conjugate (Precision Proteomics, PPI-001). The SH2 Superbinder (SH2S) is an engineered SH2 domain protein conjugated to agarose beads that binds to phosphotyrosine containing proteins with antibody-like affinity. The SH2 superbinder beads may be used like an anti-phosphotyrosine antibody in numerous applications, including immunoprecipitation and affinity purification to measure global or target protein tyrosine phosphorylation.

**Secondary Antibodies and detection reagents:**

Bovine anti-Goat HRP secondary:(Jackson ImmunoResearch, 805-035-180). Peroxidase AffiniPure Bovine Anti-Goat IgG (H+L) (min X Bov, Ck, GP, Sy Hms, Hrs, Hu, Ms, Rb, Rat Sr Prot).

Goat anti-Rabbit HRP secondary (Jackson ImmunoResearch, 111-035-144). Peroxidase AffiniPure Goat Anti-Rabbit IgG (H+L) (min X Hu, Ms, Rat Sr Prot).

Donkey anti-Mouse HRP secondary (Jackson ImmunoResearch, 715-035-151). Peroxidase AffiniPure Donkey Anti-Mouse IgG (H+L) (min X Bov, Ck, Gt, GP, Sy Hms, Hrs, Hu, Rb, Rat, Shp Sr Prot).

6xHis Tag (C-term) Monoclonal Antibody (3D5), HRP conjugated (Thermo Scientific, R931-25). Used to detect His-tagged recombinant proteins.

Mouse Penta·His Antibody (Qiagen, 34660). A combination of mouse monoclonal anti·His antibodies used to detect His-tagged recombinant proteins.

Peroxidase AffiniPure Goat Anti-Human IgG, Fcγ fragment specific (Jackson ImmunoResearch, 109-035-098). A polyclonal anti-Fc antibody used to detect Fc-tagged recombinant proteins.

AffiniPure® Goat Anti-Human IgG, Fcγ fragment specific (Jackson ImmunoResearch, 109-005-008). A polyclonal anti-Fc antibody used to pre-cluster EphrinB2-Fc and EphB2-Fc ectodomain proteins for cell stimulations.

ELISA Pico Chemiluminescent Substrate (Thermo Scientific, 37069). ELISA Pico Chemiluminescent Substrate is optimized to generate an intense light signal and provide exceptional performance in luminometer-based assays.

SuperSignal West Dura Extended Duration Substrate (Thermo Scientific, 34076). A luminol-based enhanced chemiluminescence (ECL) horseradish peroxidase (HRP) substrate ideal for quantitative western blotting with stable light output for mid-femtogram level detection for western blot analysis.

### Supplementary Figures

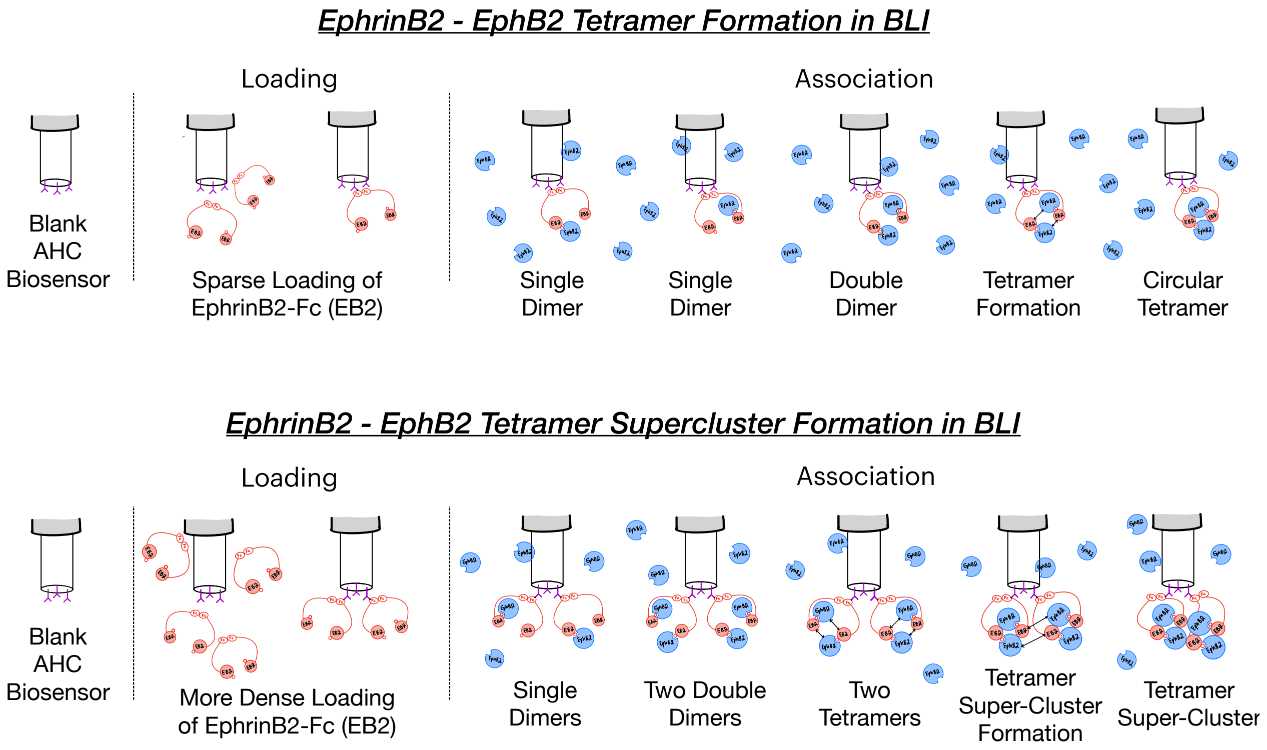

**Figure S1A.** Cartoons visualizing how biolayer interferometry (BLI) can be used to study the complex Eph-Ephrin receptor-ligand protein-protein binding interactions that lead to formation of dimers, tetramers, and tetramer superclusters. Here, on the **top drawing** as example, low concentrations of soluble EphrinB2 ectodomain available as a fusion protein with the Fc region of human IgG1 attached to the C-terminus (EphrinB2-Fc or simply EB2 in the cartoon, red) is first sparsely loaded in buffer (PBST: PBS containing 0.05% Tween-20) onto AHC or AHC2 Anti-Human Fc Capture sensorchips to immobilize the "bait" protein using an Octet RED384 (RH16) or R8e instrument (Sartorius). As the Fc domain forms a strong Fc-Fc homodimer in solution, sparse loading leads to an isolated island of two closely positioned EphrinB2-Fc ectodomains. After washing the EphrinB2-Fc loaded biosensors and obtaining baseline measurements in buffer, each biosensor is then placed in a well containing the desired soluble "prey" target protein (e.g. the EphB2 ectodomain protein fused to a poly-Histidine tag at the C-terminus, EphB2-His, blue) which is diluted to a specific concentration in buffer for a timed association step. As illustrated, each of the two available EphrinB2 ectodomain "baits" can independently bind to an EphB2 ectodomain "prey" using the dimerization interface, which leads to an increase in sensorgram response signal. At some point, both of the immobilized EphrinB2 proteins may each form a dimer with available EphB2 molecules, forming two adjacent dimers, and when that happens it now becomes possible to form the circular tetramer through the bringing together of the two distinct juxtaposed Eph-Ephrin tetramerization interfaces. After the association step, biosensors are then placed back into buffer only for a timed dissociation step. Here, the dimers, which exhibit fast-on/fast-off binding kinetics, rapidly dissociate/melt with EphB2 being released from the immobilized EphrinB2, reducing biosensor response signal and leaving behind the stable slow-on/slow-off circular tetramer macromolecular structure. As the circular tetramer is stable and can persist through to the end of the dissociation step, the sensorgram response signal remains well above that of baseline at the end of the experiment. On the **bottom drawing** is an illustration of what may occur when EphrinB2-Fc "bait" protein is loaded at higher density on the biosensor surface where two Fc-Fc homodimers become immobilized to occupy nearby adjacent regions of the chip. Here, dimer and circular tetramer formation proceed as above, but now with two adjacent circular tetramers nearby, the protein complexes can interact and assemble through other Eph-Ephrin contact interfaces to form tetramer superclusters, which are extremely stable macromolecular complexes less prone to be targeted by tetramer inhibitors (insurmountable). The cartoons shown are simplified and do not take into account that soluble EphB2 and EphrinB2 ectodomains can also each assemble into homodimers (see **Figure 1I**). Reference biosensors loaded with the human Fc only protein or blank loaded with no protein were included in the BLI studies to control for any non-specific binding and typically showed no response signal (or only slight signals) when exposed to an Eph-His protein during the association step, compared to sensors loaded with an EphrinB-Fc protein which exhibited obvious strong response signals. Biosensors loaded with EphrinB-Fc protein but only exposed to buffer only during the association step were used to control for and subtract baseline drift. These control EphrinB-Fc loaded sensors typically exhibited little if any response drift, indicating protein remained stably immobilized onto the AHC/AHC2 chips throughout the baseline, association, and dissociation steps.

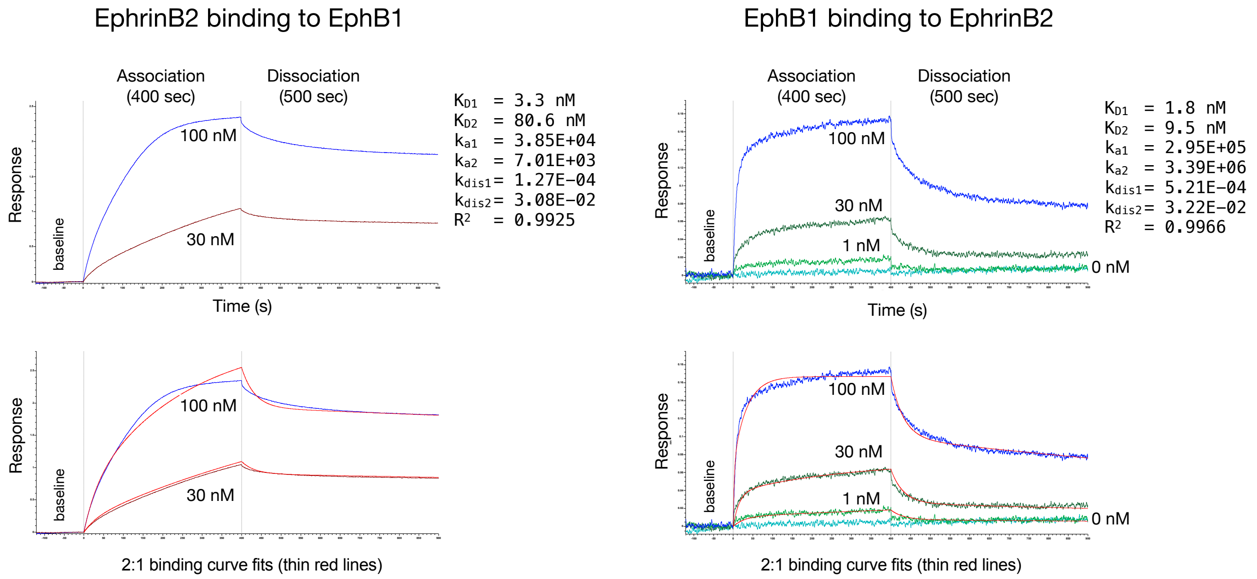

**Figure S1B (related to Figures 1B and 1C).** Shown are sensorgram traces of BLI kinetic binding studies using 2 or 3 concentrations of the soluble component to calculate using global fit analysis important information regarding the protein-protein interaction, including the association rate constant ($k_a/k_{on}$), dissociation rate constant ($k_d/k_{dis}/k_{off}$), and equilibrium dissociation constant ($K_D$) used to describe the biophysical properties and strength/affinity of an interaction under study, in this case EphB1-EphrinB2 binding. Here, Super Streptavidin (SSA) biosensors were used to either sparsely immobilize biotinylated mouse EphrinB2-His ectodomain and assess binding to 30 and 100 nM soluble rat EphB1-Fc ectodomain (left) or sparsely immobilize biotinylated rat EphB1-Fc ectodomain and assess binding to 1, 30, and 100 nM soluble mouse EphrinB2-His ectodomain (right). In both cases the association time was 400 sec and dissociation time was 500 sec, buffer was PBST. Mathematical models are used to assess the resulting protein binding sensorgram response data and attempt to "curve fit" (thin red lines) the data using either (1) a simple 1:1 type model that assumes a basic dimer binding kinetics prevail where one soluble analyte molecule binds to a single, specific site on an immobilized partner ligand molecule, or (2) a more complex 2:1 heterologous model which assumes as a basic premise that analyte can bind to two independent sites on the ligand with different affinities. Consistent with their ability to form dimers and tetramers, the curve fits calculated from the above EphB1-EphrinB2 sensorgram traces and the 2:1 binding model indicated these proteins form complex heterologous interactions. With 2:1 type interactions, the model attempts to best fit the curve by calculating two different values for each parameter to explain the complex interactions, including two different $k_a$, $k_{dis}$, and $K_D$ values, with $k_{a2}/k_{dis2}/K_{D2}$ used here to describe dimer formation and $k_{a1}/k_{dis1}/K_{D1}$ used to describe tetramer formation. In these calculations, the R-squared ($R^2$) informs how well the actual sensorgram trace data fits the mathematical curve, with values close to 1.0 indicating a perfect fit.

BLI studies of immobilized human EphrinB ectodomains binding to soluble human EphB ectodomains

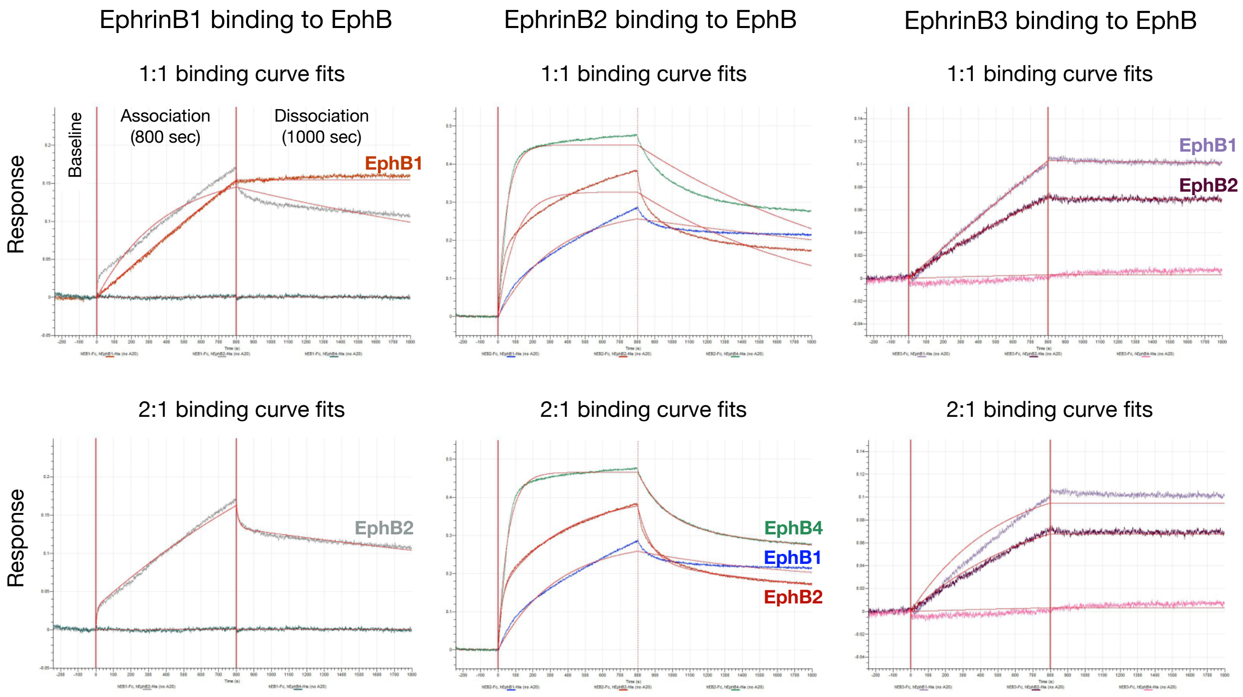

**Figure S1C (related to Figure 1B).** Curve fits to show both 1:1 and 2:1 binding models (thin red lines) superimposed on the sensorgram traces for the BLI kinetic studies shown in **Figure 1B** using AHC biosensor immobilized human EphrinB1-Fc, EphrinB2-Fc, and EphrinB3-Fc ectodomains binding to 50 nM soluble human EphB1-His, EphB2-His, and EphB4-His ectodomains. The $K_D$ and $R^2$ values calculated from these single 50 nM concentration BLI sensorgram runs are provided in **Figure S1E**.

As expected, the immobilized EphrinB1 ectodomain specifically bound EphB1 and EphB2 ectodomains, but not EphB4. EphrinB1 bound very slowly to EphB1 presented at 50 nM in the association step and with a straight slope of 45$^o$, without any sign of response plateauing after 800 sec, and showed no loss of receptor binding during the 1,000 sec dissociation step. This indicates formation of an extremely high affinity/super-stable EphrinB1-EphB1 protein complex. The slow, straight slope of response from the very beginning of the association step indicates EphrinB1-EphB1 proteins may assemble directly into tetramers in a 1:1 binding model and the curve fit (thin red lines) supports this, apparently bypassing the need for dimer formation. EphrinB1 binding to 50 nM EphB2 was slightly different to that of EphB1. Here, EphB2 exhibited a very brief extremely high slope of response at the very beginning of the association step that quickly shifted within 10-20 sec to a slow straight association of 45$^o$ with no sign of response plateauing, quite similar to that of EphB1. Once placed into dissociation, the sensorgram response showed a rapid loss of bound EphB2 for the initial 20-40 sec and then it flatlined to lose hardly any more of the bound protein. The cure fit (thin red lines) here precisely matches a 2:1 heterologous binding model. Particularly, the immediate high slope of response from 0-20 sec at the start of the association step and initial high reverse slope of response for the first 40 sec once placed into dissociation observed for EphrinB1-EphB2 binding is similar to the kinetics observed for the three different EphB receptors binding to EphrinB2 (below), and visualizes a rapid on/off rate for the lower affinity EphrinB-EphB dimer, which are fast-on/fast-off type interactions. The rapid shift in response from a high-slope fast-on dimer formation to a less steep 45$^{o}$ slope visualizes the slow, gradual accumulation of circular tetramers during the association step. Likewise, the initial rapid loss of response signal once placed into the dissociation step visualizes melting of the fast-off dimers, with response then quickly flatlining to lose very little signal as the tetramers are super-stable and essentially do not dissociate.

As expected, the immobilized EphrinB2 ectodomain exhibited binding to all three EphB ectodomains when presented at 50 nM, with the EphB4 protein showing the fastest/strongest binding with a very steep and high initial response slope and with binding nearly complete within the first 200 sec of association as the response slope quickly flattened. In contrast, the binding of EphB1 and EphB2 to immobilized EphrinB2 occurred more slowly and with no sign of the response slope reaching a plateau, even at the end of the 800 sec association step. Once placed into dissociation, all three EphrinB2-EphB interactions exhibited similar profiles with a fair amount of EphB protein coming off the EphrinB2-bound sensorchips, though ~50% or more of the respective EphB protein remained bound at the end of the 1,000 sec dissociation step. The curve fits clearly indicate the EphrinB2 ectodomain forms complex 2:1 type heterologous interactions with all three EphB1, EphB2, and EphB4 ectodomains.

The EphrinB3 ectodomain also specifically bound to only EphB1 and EphB2, again as expected. In both cases, EphrinB3 bound very slowly to 50 nM EphB1 and EphB2 in the association step and with straight slopes, without any sign of response plateauing after 800 sec, and they showed no loss of response signal once placed in the dissociation step. This indicates extremely high affinity/stability of the EphrinB3-EphB1 and EphrinB3-EphB2 protein complexes. The slow, straight slopes of response from the very beginning of the association step indicates EphrinB3-EphB1 and EphrinB3-EphB2 proteins, like EphrinB1-EphB1, at 50 nM in these early BLI studies assemble directly into tetramers, apparently bypassing the need for dimer formation.

Curve fits for global binding affinities using BLI to test immobilized EphB binding to soluble EphrinB2

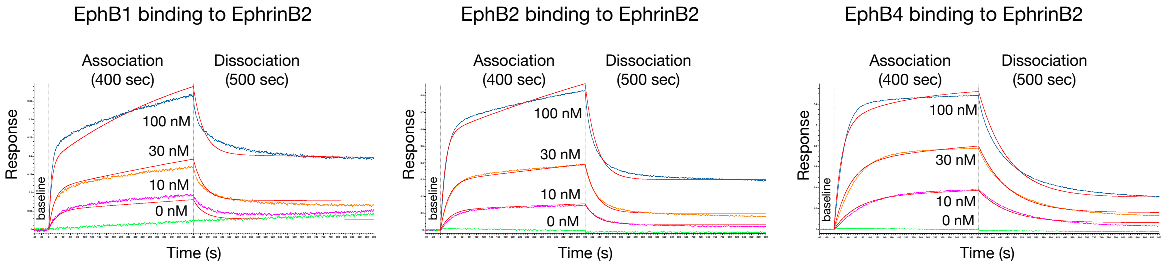

| EphB1 | EphB2 | EphB4 |
|---|---|---|
| $K_{D1}$ = 3.7 nM | $K_{D1}$ = 0.28 nM | $K_{D1}$ = 4.8 nM |
| $K_{D2}$ = 39.7 nM | $K_{D2}$ = 70 nM | $K_{D2}$ = 15.3 nM |
| $k_{a1}$ = 2.15E+04 | $k_{a1}$ = 8.25E+03 | $K_{a1}$ = 3.20E+05 |
| $k_{a2}$ = 8.87E+05 | $k_{a2}$ = 4.58E+05 | $K_{a2}$ = 1.01E+06 |
| $kdis_1$ = 7.98E-05 | $kdis_1$ = 2.28E-06 | $Kdis_1$ = 1.52E-03 |
| $kdis_2$ = 3.52E-02 | $kdis_2$ = 3.20E-02 | $Kdis_2$ = 1.55E-02 |
| $R^2$ = 0.9796 | $R^2$ = 0.9959 | $R^2$ = 0.9868 |

**Figure S1D (related to Figure 1C).** Curve fits for 2:1 heterologous binding model (thin red lines) superimposed on the sensorgrams for the BLI kinetic studies shown in **Figure 1C** using SSA immobilized biotinylated rat EphB1-Fc, mouse EphB2-Fc, and mouse EphB4-Fc ectodomains binding to 0, 10, 30, and 100 nM soluble mouse EphrinB2-His ectodomain in early global fit studies. The $K_D$, $k_a$, $k_{dis}$, and $R^2$ values calculated from the sensorgram traces provide accurate descriptions of the complex high-affinity 2:1 heterologous type protein-protein interactions consistent with the ability of these molecules to form dimers and tetramers.

$K_D$ calculations of various EphB-EphrinB binding interactions determined using BLI

| | | Single Run Kinetics (binding at 50 nM) | | | | | Global Fit Kinetics (binding at 10, 30, 100 nM) | | |
|---|---|---|---|---|---|---|---|---|---|
| | | 1:1 interaction model | | 2:1 heterologous interaction model | | | 2:1 heterologous interaction model | | |
| EphrinB | EphB | $K_D$ | $R^2$ | $K_{D1}$ | $K_{D2}$ | $R^2$ | $K_{D1}$ | $K_{D2}$ | $R^2$ |
| EphrinB1 | EphB1 | 11.4 pM | 0.9950 | nd | nd | | | | |
| EphrinB1 | EphB2 | 8.57 nM | 0.9122 | 41 nM | 40 nM | 0.9870 | | | |
| EphrinB1 | EphB4 | no binding | | | | | | | |
| EphrinB2 | EphB1 | 4.3 nM | 0.9661 | < 1 pM | 27 nM | 0.9723 | 3.7 nM | 39.7 nM | 0.9796 |
| EphrinB2 | EphB2 | 4.84 nM | 0.8504 | 5.9 nM | 16.8 nM | 0.9965 | 0.28 nM | 70 nM | 0.9959 |
| EphrinB2 | EphB4 | 1.8 nM | 0.9249 | 0.35 nM | 42 nM | 0.9948 | 4.8 nM | 15.3 nM | 0.9868 |
| EphrinB3 | EphB1 | 8.67 nM | 0.9969 | 1.4 pM | 1.2 pM | 0.9296 | | | |
| EphrinB3 | EphB2 | 1.58 nM | 0.9947 | 1.8 pM | 1.8 pM | 0.9845 | | | |
| EphrinB3 | EphB4 | no binding | | | | | | | |

**Figure S1E.** The equilibrium dissociation constants ($K_D$) for the different EphrinB-EphB interaction pairs were calculated using the BLI kinetic sensorgram data shown in **Figure 1B** and **Figure 1C** and bundled Octet Studio Analysis software. The single run values from **Figure 1B** (and curve fit in **Figure S1C**) were acquired using AHC immobilized human EphrinB-Fc binding to 50 nM soluble human EphB-His. The global fit values from **Figure 1C** (and curve fit in **Figure S1D**) were acquired using SSA immobilized biotinylated rat EphB1-Fc, mouse EphB2-Fc, and mouse EphB4-Fc binding to 10, 30, and 100 nM soluble mouse EphrinB2-His.

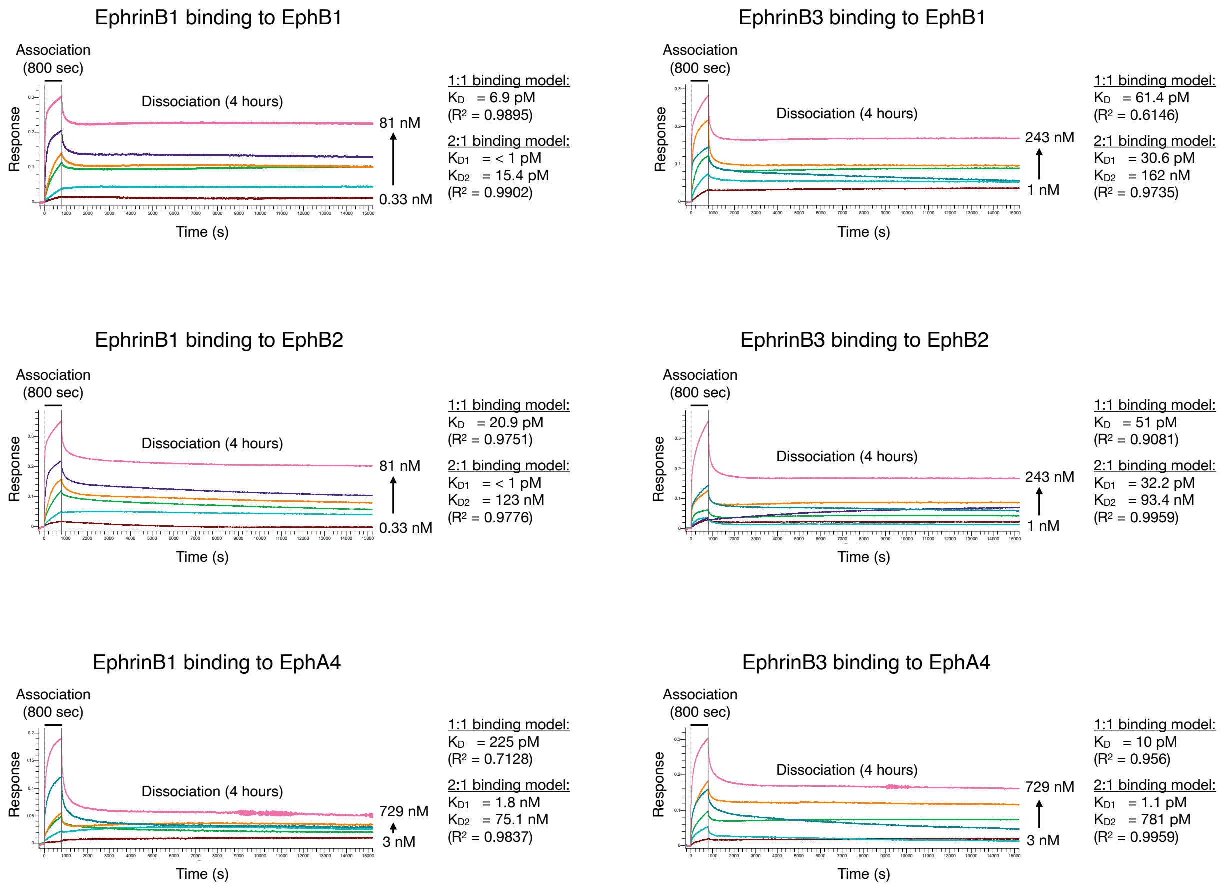

**Figure S1F (related to Figure 1D).** Shown are example sensorgrams and calculated equilibrium dissociation constants ($K_D$) using Octet Studio Analysis software from expanded global fit kinetic BLI studies using AHC2 biosensors sparsely loaded with the indicated human EphrinB-Fc bait protein and exposed to 6-7 different concentrations of soluble analyte prey protein (0.33-243 nM for human EphB1-His and EphB2-His, 3-729 nM for human EphA4-His) for an 800 sec association step followed by an extended 4 hr dissociation step. A reference biosensor loaded with the human EphrinB-Fc protein and exposed to buffer only was used to measure for possible biosensor response drift, which was generally non-existent or minimal, and for preprocessing signal subtraction/ correction. Additional control biosensors were loaded with human Fc only protein and exposed to high concentrations of tested analyte to identify any non-specific binding when exposed to an Eph-His protein, which typically, as expected, showed no or only a slight increase response signals during the association step.

Focusing on the key 800 sec association step, all tested Eph receptors quickly bound the immobilized Ephrin ligands as indicated by a concentration-dependent increase in response signal that begins immediately upon placement of biosensor tip into its association well, and with high immediate response slopes in wells testing high concentrations of the analyte Eph protein that gradually bends as tetramers accumulate.

Once placed into the dissociation step, biosensors exposed to high concentrations of the Eph molecule showed a rapid loss of a portion of the signal (the dimers), but then the response plateaued to maintain a high signal significantly above baseline that showed little, if any, further dissociation for the remaining 2 hr, indicating formation of an extremely stable complex (the tetramers). Likewise, the responses observed in biosensors exposed to low concentrations of analyte in the association step similarly showed some initial loss of signal once placed into the dissociation well, but that loss quickly plateaued well above baseline for the remaining 4 hr. Combined, the response traces for all tested analyte concentrations of an Eph molecule under study provide extensive global fit data for determination of important kinetic information concerning the protein-protein interaction (see Tables S1 and S2).

Average $K_D$ values calculated from multiple global fit runs using immobilized EphrinB-Fc ectodomains binding wide concentration ranges of indicated Eph analyte ectodomain and with an extended 2-4 hour long dissociation step, analyzed using Octet Studio Analysis software

| | | 1:1 interaction model | | 2:1 heterologous interaction model | | | |
|---|---|---|---|---|---|---|---|
| Ephrin | Eph | $K_D$ | $R^2$ | $K_{D1}$ | $K_{D2}$ | $R^2$ | n |
| EphrinB1 | EphB1 | 5 pM | 0.9735 | < 1 pM | 17 pM | 0.9737 | 2 |
| EphrinB1 | EphB2 | 21 pM | 0.9751 | < 1 pM | 123 nM | 0.9776 | 1 |
| EphrinB1 | EphB4 | no binding | | | | | |
| EphrinB1 | EphA4 | 225 pM | 0.7128 | 1820 pM | 75 nM | 0.9837 | 1 |
| EphrinB2 | EphB1 | 11 pM | 0.9856 | 3.5 pM | 132 nM | 0.9792 | 3 |
| EphrinB2 | EphB2 | 193 pM | 0.8760 | 3.3 pM | 158 nM | 0.9057 | 4 |
| EphrinB2 | EphB4 | 16 pM | 0.9799 | < 1 pM | 0.33 nM | 0.9692 | 3 |
| EphrinB2 | EphA4 | 529 pM | 0.9648 | 634 pM | 30 nM | 0.9795 | 3 |
| EphrinB3 | EphB1 | 36 pM | 0.7695 | 16 pM | 81 nM | 0.9424 | 2 |
| EphrinB3 | EphB2 | 26 pM | 0.9407 | 11.4 pM | 31 nM | 0.9705 | 3 |
| EphrinB3 | EphB4 | no binding | | | | | |
| EphrinB3 | EphB4 | 10 pM | 0.9560 | < 1 pM | 0.78 nM | 0.9612 | 1 |

**Figure S1G (related to Figure 1D).** Ephrin-Eph equilibrium dissociation constants ($K_D$) were calculated by averaging the results of multiple independent BLI global fit kinetic experiments that were analyzed using Octet Studio Analysis software. AHC/AHC2 biosensors sparsely loaded with the indicated human EphrinB-Fc ectodomain bait protein and exposed to increasing concentrations of indicated soluble analyte Eph-His ectodomain prey protein for an 800 sec association step followed by an extended 2-4 hr dissociation step in PBST. The concentration of soluble Eph molecules in the association step ranged from 110 pM up to 729 nM, depending on the protein, with EphA4-His in some cases requiring higher concentrations due to its lower affinity binding to EphrinB1/EphrinB2 ectodomains. A reference biosensor loaded with the human EphrinB-Fc protein and exposed to buffer only was used to measure for possible biosensor response drift and for signal subtraction/correction. Additional control biosensors were loaded with human Fc only protein and exposed to high concentrations of tested analyte to identify any non-specific binding when exposed to an Eph-His protein, which typically, as expected, showed no or only a slight increase response signals during the association step. The number of independent experiments for each Ephrin-Eph interaction assessed is indicated, with the full kinetic information from all runs presented in Table S1.

Average $K_D$ values and tetramer half-life ($t_{1/2}$) from multiple global fit runs using immobilized EphrinB-Fc ectodomains binding wide concentration ranges of indicated Eph analyte ectodomain and with an extended 2-4 hour long dissociation step, analyzed using the newly developed dimer:tetramer model

| | | dimer:tetramer interaction model | | | | | | |
|---|---|---|---|---|---|---|---|---|
| Ephrin | Eph | $K_{D1}$ (tetramer) | $K_{D2}$ (dimer) | $t_{1/2}$ (tetramer) | $t_{1/2}$ (dimer) | $R^2$ (association) | $R^2$ (dissociation) | n |
| EphrinB1 | EphB1 | 28 pM | 16.4 nM | 7 days | 1.6 minutes | 0.9994 | 0.9941 | 2 |
| EphrinB1 | EphB2 | 815 pM | 8 nM | 0.47 days | 1.6 minutes | 0.9992 | 0.9938 | 1 |
| EphrinB1 | EphB4 | no binding | | | | | | |
| EphrinB1 | EphA4 | 2815 pM | 56 nM | 0.42 days | 1.4 minutes | 0.9993 | 0.9824 | 1 |
| EphrinB2 | EphB1 | 1.3 nM | 9.2 nM | 0.81 days | 2.3 minutes | 0.9982 | 0.9940 | 3 |
| EphrinB2 | EphB2 | 0.38 nM | 8.6 nM | 1.12 days | 2.2 minutes | 0.9986 | 0.9947 | 4 |
| EphrinB2 | EphB4 | 1.77 nM | 6.3 nM | 0.53 days | 3.5 minutes | 0.9963 | 0.9847 | 3 |
| EphrinB2 | EphA4 | 1.68 nM | 47.9 nM | 0.35 days | 1.3 minutes | 0.9991 | 0.9751 | 3 |
| EphrinB3 | EphB1 | 0.593 nM | 54.8 nM | 1.92 days | 0.8 minutes | 0.9968 | 0.9888 | 2 |
| EphrinB3 | EphB2 | 3.05 nM | 27.4 nM | 0.28 days | 1.5 minutes | 0.9998 | 0.9955 | 3 |
| EphrinB3 | EphB4 | no binding | | | | | | |
| EphrinB3 | EphB4 | 1.6 nM | 43.9 nM | 0.94 days | 2 minutes | 0.9995 | 0.9913 | 1 |

**Figure S1H (related to Figure 1D and S1G).** A new "dimer:tetramer" mathematical model was developed to better analyze Ephrin-Eph equilibrium dissociation constants ($K_D$) that are associated with dimer and tetramer binding kinetics (see Methods for full description of the model). The dimer:tetramer model was tested on the same sensorgram data analyzed above in Supplementary Figure S1G, with data presented including overall equilibrium dissociation binding affinities ($K_D$) for both tetramer and dimer ($K_{D1}$), as well as dissociation half-life ($t_{1/2}$) for both dimer and tetramer. The R-squared shows the new dimer:tetramer model fits the data extremely well. The number of independent experiments for each Ephrin-Eph interaction assessed is indicated, with the full kinetic information from all runs analyzed using dimer:tetramer model presented in Table S2.

Average $K_D$ values and tetramer half-life ($t_{1/2}$) from multiple global fit runs using immobilized EphrinB-Fc ectodomains binding wide concentration ranges of indicated Eph analyte ectodomain and with an extended 15 hour long dissociation step, testing addition of BSA as blocking agent to PBST binding buffer, analyzed using the newly developed dimer:tetramer model

| | | dimer:tetramer interaction model | | | | | | |
|---|---|---|---|---|---|---|---|---|
| Ephrin | Eph | $K_{D1}$ (tetramer) | $K_{D2}$ (dimer) | $t_{1/2}$ (tetramer) | $t_{1/2}$ (dimer) | $R^2$ (association) | $R^2$ (dissociation) | n |
| EphrinB2 | EphB2 | 70 pM | 7.1 nM | 4.3 days | 7.6 minutes | 0.9996 | 0.9985 | 2 |
| EphrinB3 | EphB2 | 382 pM | 2.0 nM | 7.5 days | 6.8 minutes | 1.0 | 0.9995 | 2 |

**Figure S1I (related to Figure 1D, S1G, and S1H).** BSA was added to the PBST buffer to determine if use of a blocking agent in global fit studies of EphrinB2-EphB2 and EphrinB3-EphB2 binding would alter dimer/tetramer association-dissociation kinetics, here using a very long 15 hour extended dissociation step. Addition of BSA was found to improve the binding kinetics, providing higher overall binding affinities for both the dimer and tetramer, as especially increasing the half-life of the tetramer. The number of independent experiments for each Ephrin-Eph interaction assessed is indicated, with the full kinetic information from all runs analyzed using dimer:tetramer model presented in Table S2.

Alpha Assay Development - Testing EphB1 binding to EphrinB1, EphrinB2, and EphrinB3 in 384-well plates

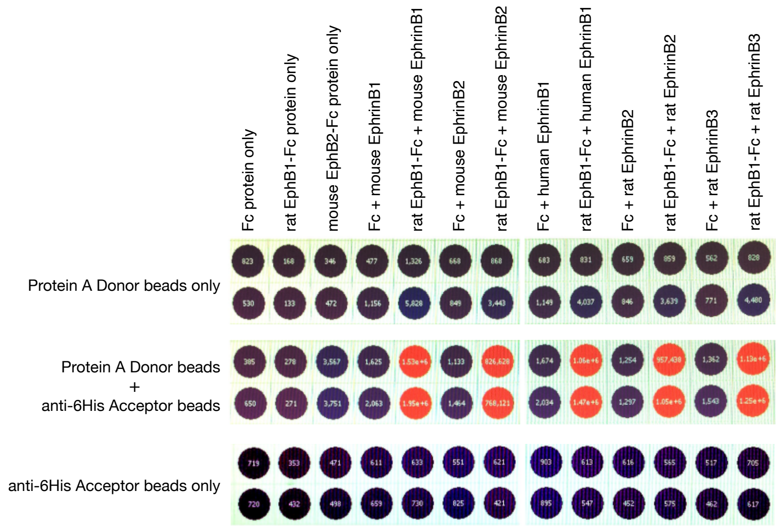

<u>Conditions:</u> 384-well plates
AlphaLISA 1X HiBlock Buffer
40 nM EphB1-Fc
40 nM EphrinB-His
125 ng Protein A donor beads per well
125 ng anti-6His acceptor beads per well

<u>Scan device:</u> PerkinElmer EnVision Multilabel Plate Reader

<u>AlphaLISA 1X HiBlock Buffer:</u> 25 mM HEPES, pH 7.4, 0.1% Casein, 1 mg/ml Dextran-500, 0.5% Triton X-100, 0.5% blocking reagent, 0.5% BSA

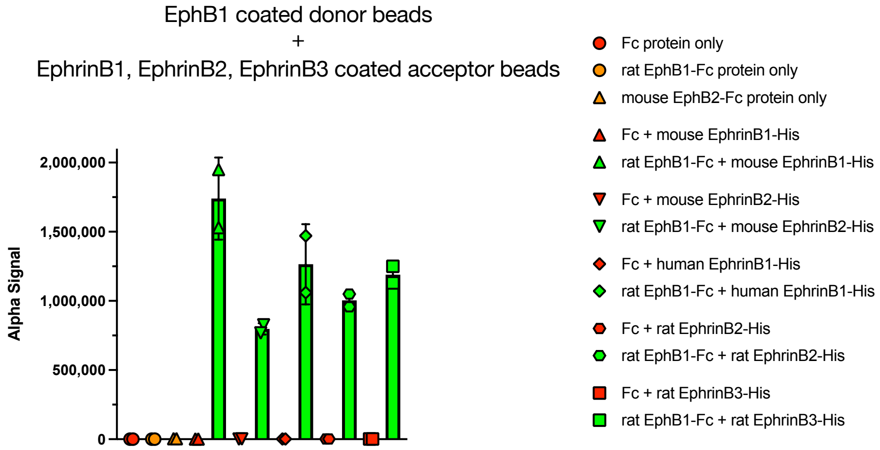

**Figure S2A.** Development of EphB1-EphrinB1, EphB1-EphrinB2, and EphB1-EphrinB3 proximity-based Alpha assays for high-throughput screening (HTS). Testing 40 nM rat EphB1-Fc ectodomain or Fc only coated Protein A donor beads with 40 nM of mouse/ human EphrinB1-His, mouse/rat EphrinB2-His, and rat EphrinB3-His ectodomain coated anti-6His acceptor beads in 384-well plates. After overnight incubation at RT, plates were placed in an EnVision plate reader for excitation and chemiluminescent detection. Extremely strong Alpha signals were detected for the wells with EphB1-Fc and any of the EphrinB-His proteins, whereas no signal was detected if Fc control protein replaced the EphB1-Fc protein. Likewise, no Alpha signals were detected in wells contained only the EphB1-Fc protein or only the EphrinB-His protein.

Alpha Assay Development - With 0.8 nM EphB1 and EphrinB2, testing EWLS peptide

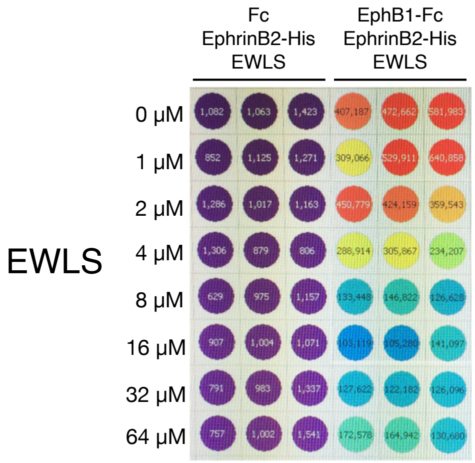

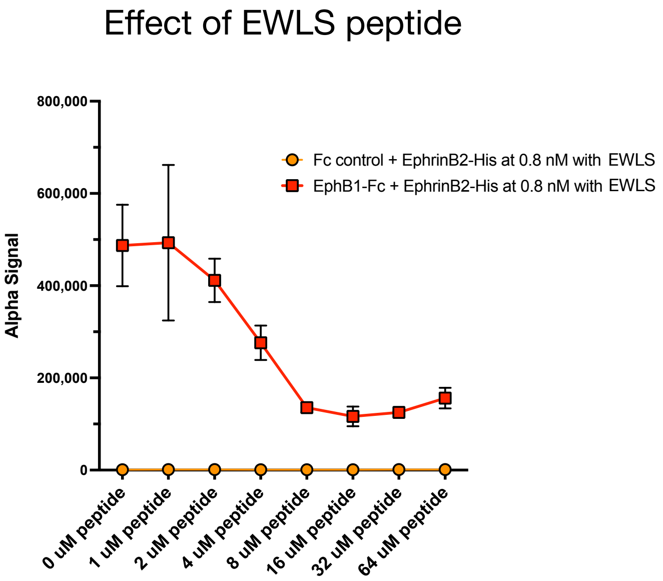

Conditions: 384-well plates
AlphaLISA HiBlock Buffer
0.8 nM rat EphB1-Fc
0.8 nM mouse EphrinB2-His
125 ng each bead per well

**Figure S2B.** Assessment of EphB1-EphrinB2 Alpha assay for HTS using 0.8 nM rat EphB1-Fc ectodomain coated Protein A donor beads and 0.8 nM mouse EphrinB2-His ectodomain coated anti-6His acceptor beads, testing 1-64 μM of soluble EWLS peptide in 384-well plates.

#### Alpha Assay Development - With 0.8 nM EphB1 and EphrinB2, testing Reelin

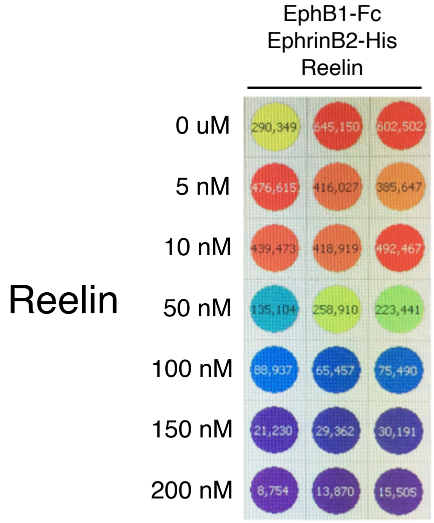

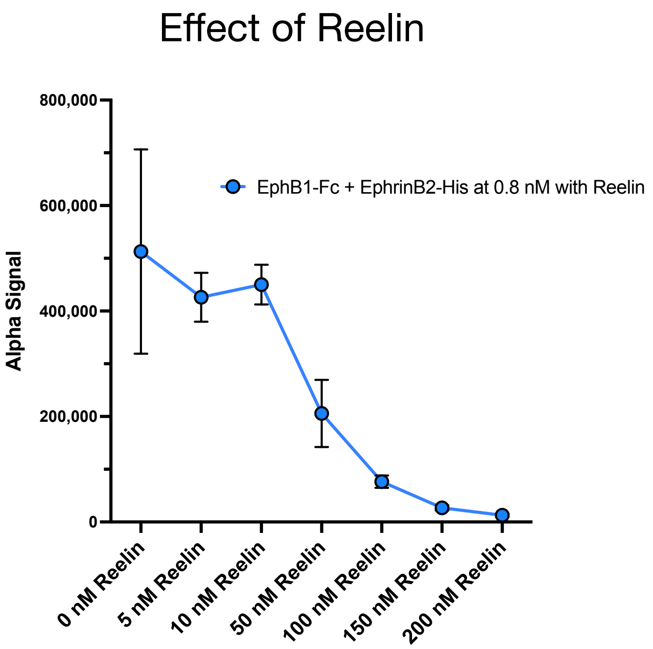

Conditions: 384-well plates
AlphaLISA HiBlock Buffer
0.8 nM rat EphB1-Fc
0.8 nM mouse EphrinB2-His
125 ng each bead per well

**Figure S2C.** Assessment of EphB1-EphrinB2 Alpha assay for HTS using 0.8 nM rat EphB1-Fc ectodomain coated Protein A donor beads and 0.8 nM mouse EphrinB2-His ectodomain coated anti-6His acceptor beads, testing 5-200 nM soluble Reelin in 384-well plates.

Sample of an HTS plate with a possible hit

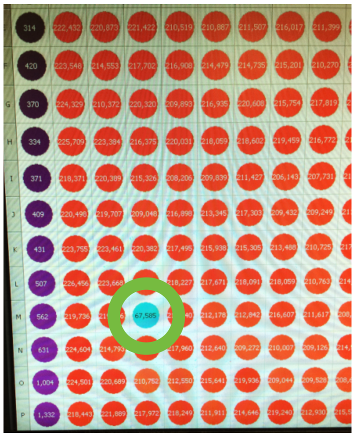

HTS Conditions: 384-well plates
AlphaLISA HiBlock Buffer
0.8 nM rat EphB1-Fc
0.8 nM mouse EphrinB2-His
50 ng each bead per well
5 μM compound, 2% DMSO

Results of HTS campaign

240,000 compounds screened at 5 μM in EphB1-EphrinB2 AlphaScreen

↓ Average Z' = 0.92

1,055 compounds reduced Alpha signal >10%

↓
- •243 compounds reduced signal ≥25%
- •383 compounds reduced signal 20-24.99%
- •429 compounds reduced signal 10-19.99%

950 hits cherry-picked and retested in EphB1-EphrinB2 AlphaScreen

↓

Hits confirmed in above retest subjected to single-protein Alpha counter screen to eliminate false positives

↓

Remaining positives tested in EphB1-EphrinB2, EphB2-EphrinB2, and EphB4-EphrinB2 ELISA assays

↓

Remaining positives tested in EphB1-EphrinB2, EphB2-EphrinB2, and EphB4-EphrinB2 pulldown assays

↓

Lead compound A20 (SW056428) identified

**Figure S2D.** Execution of EphB1-EphrinB2 AlphaScreen HTS of a ~240,000 small molecule chemical library at 5 μM using 0.8 nM rat EphB1-Fc ectodomain coated Protein A donor beads and 0.8 nM mouse EphrinB2-His ectodomain coated anti-6His acceptor beads in 384-well plates. An example of a plate run with a potential hit well circled in green is shown (left) as is a summary of the flow of additional Alpha screens, counter screens, ELISAs, and pulldown assays that resulted in the identification of lead compound A20 (right). A total of 1,055 potential hits that reduced the Alpha signal >10% were identified. 950 of the 1,055 compounds were selected from master plates, avoiding those with "pan-assay interference" (PAINS) properties that tend to nonspecifically react with numerous biological targets and are often false positives, and retested in triplicate in the EphB1-EphrinB2 Alpha assay. Selected compounds were then tested in a counter screen designed to identify non-specific compounds in a single-protein Alpha assay using a dual-tagged EphB2-His-Fc ectodomain protein that can bind both the Protein A donor bead and the anti-6His Acceptor bead to bring both beads into proximity. If a selected compound simply absorbs the chemiluminescent signal or interferes with bead function/binding or protein folding, it was hypothesized that it would also reduce signals in the single protein assay, and would consider it a false positive. Together, the Alpha confirmation and counter screens allowed us to eliminate a large number of the hits, leaving 32 compounds to advance forward that showed repeated ability to reduce the EphB1-EphrinB2 Alpha assay chemiluminescent signal >10% and had no effect on signal in the single-protein assay. The 32 hits were then subjected to ELISA in which EphB1-Fc was first immobilized in 96 well Protein A plates, then soluble EphrinB2-His was added with hit compound in triplicate, again at 5 μM. After 2 hr at RT to allow EphrinB2-His binding to immobilized EphB1-Fc, wells were washed, incubated with nickel-activated horseradish peroxidase (HisProbe-HRP) to detect bound EphrinB2-His protein, washed, and then incubated with ELISA Pico Chemiluminescent Substrate (both from Thermo-Fisher) before being read in a luminometer. A single-protein counter ELISA using the dual-tagged EphB2-His-Fc ectodomain protein was also run. This allowed us to quickly narrow down to one compound from well 268 G15 (SW056428) that reduced signal in the two-protein ELISA but had no effect on signal in the single-protein ELISA. For simplicity, we call this compound A20.

A20 strongly antagonizes EphrinB2 binding to EphB1 and EphB2, with less activity towards EphB4

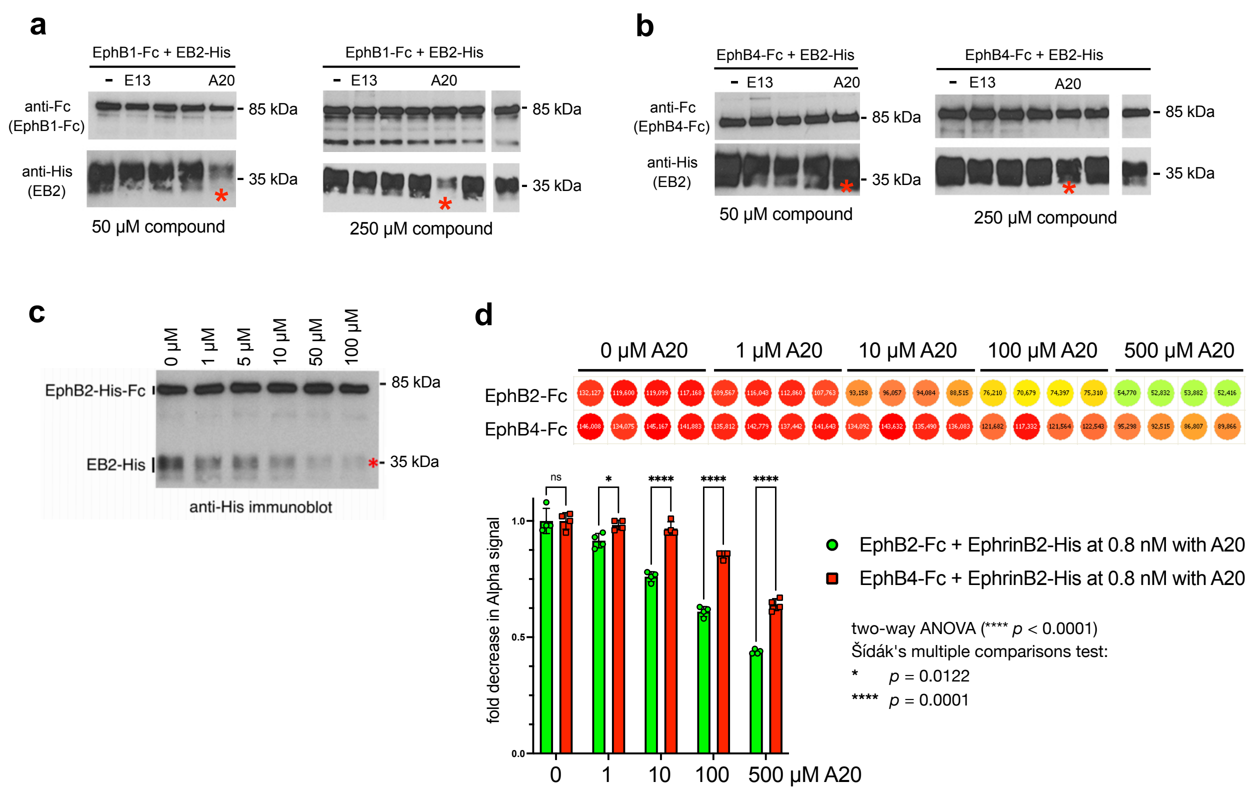

**Figure S2E.** Early biochemical pulldown and additional Alpha studies of lead hit A20 indicate a strong effect of compound on the EphB1-EphrinB2 and EphB2-EphrinB2 protein-protein interactions, with less activity towards the EphB4-EphrinB2 interaction. In **a-c**, the indicated EphB-Fc ectodomain was immobilized on Protein A Sepharose-conjugated beads and mixed with soluble EphrinB2-His ectodomain (EB2-His) in the absence (-) or presence of the indicated concentration of A20 compound and some of the other early potential hit compounds identified in the HTS, including one termed E13. After washing the beads, EphB-bound EphrinB2-His ectodomain that was pulled down in the assay was visualized using SDS-PAGE and anti-His immunoblot analysis, revealing A20 strongly reduce the amount of EphrinB2 that co-precipitated with EphB1 and EphB2, with little effect on the EphB4-EphrinB2 interaction (red asterisks). In **d**, Alpha assays testing EphB2-Fc and EphB4-Fc binding to EphrinB2-His confirmed A20 has a greater ability to antagonize the EphB2-EphrinB2 interaction compared to EphB4-EphrinB2.

### Lead compound A20 and similar early molecules of interest

**SW056428**
**(A20)**

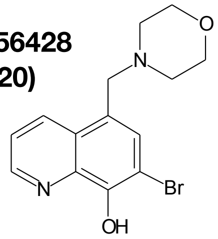

Chemical Formula: $C_{14}H_{15}BrN_2O_2$
Molecular Weight: 323.19

SW022102

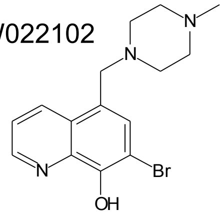

SW022101

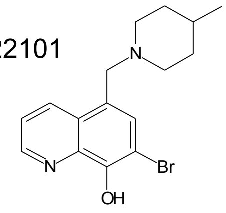

SW134218
(3511-0013)

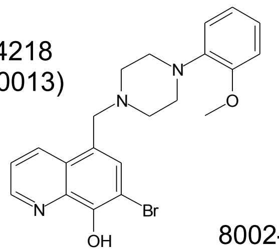

8009-9255

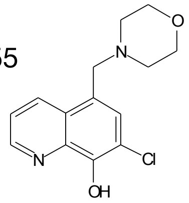

D014-0004

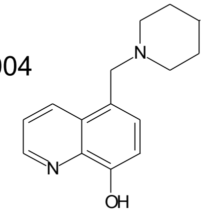

8002-0823

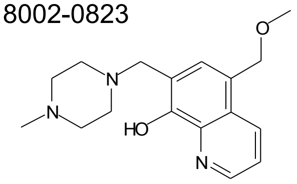

F235-1974

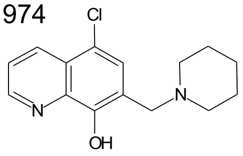

**Figure S2F.** Chemical structure of lead hit A20 and other analogs used in early studies. Three other compounds present in the chemical library screened, SW022101, SW022102, and SW134218 are similar to A20. While SW022102 was not identified in the HTS, SW022101 and SW134218 were both in the initial 950 selected primary hits, though they did not pass through subsequent Alpha confirmation and counter screens. In addition to A20, SW022101, SW022102, and SW134218, four other similar analogs not in the library were identified as being commercially available and were obtained and subjected to Alpha, ELISA, and pulldown experiments. The results from these early studies indicated compounds 3511-0013 and 8009-9255 are as effective as A20 at antagonizing EphB1-EB2 and EphB2-EB2 interactions, again with very little if any effect on the EphB4-EB2 interaction. Compounds D014-0004, F235-1974, and 8002-0823 did not exhibit any inhibitor activity.

### Summary of ELISA and pulldown results with early IC50 values

| | | ELISA | | | Pulldown |
|---|---|---|---|---|---|
| Name/ID | Structures | IC50 of EphB1-EB2 | IC50 of EphB2-EB2 | IC50 of EphB4-EB2 | IC50 of EphB1-EB2 |
| "A20" 8009-7666 | 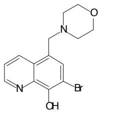 | 18 uM | 32 uM | >1000 uM | 18 uM |
| "3511" 3511-0013 | 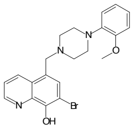 | 14 uM | 30 uM | >1000 uM | 9 uM |
| 8009-9255 | 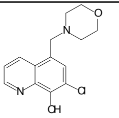 | 7 uM | 14 uM | No inhibition | 12 uM |
| D014-0004 | 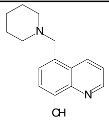 | >1000 uM | >1000 uM | No inhibition | >1000 uM |
| F235-1974 | 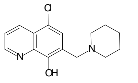 | 425 uM | No inhibition | No inhibition | >1000 uM |
| 8002-0823 | 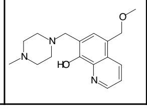 | >1000 uM | No inhibition | No inhibition | >1000 uM |

**Figure S2G.** Summary of ELISA and pulldown results with early $IC_{50}$ values (Inhibitory Concentration 50%, represents the concentration of a compound required to inhibit the protein-protein interaction by 50%) for A20 and other early analogs of interest (all free bases) when tested for their ability to disrupt immobilized rat EphB1-Fc, mouse EphB2-Fc, and mouse EphB4-Fc ectodomains binding to soluble mouse EphrinB2-His (EB2) ectodomain.

EphB1 binding to EphrinB2 for 80-40-20-15-10-5 second association steps

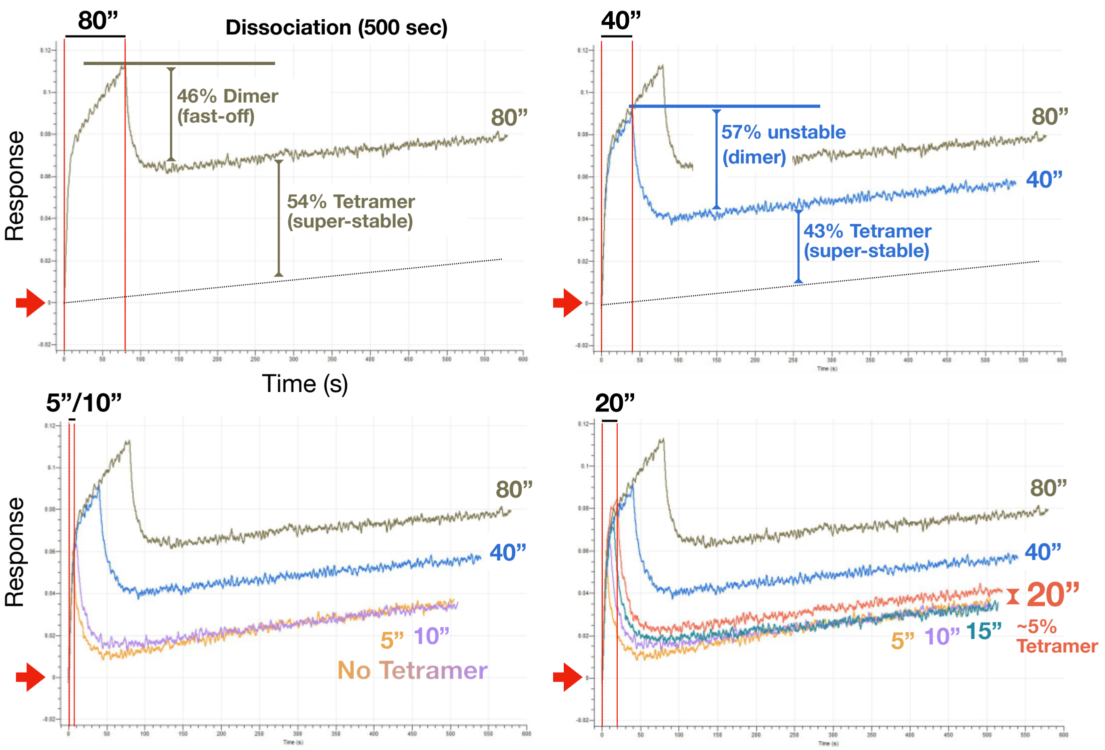

**Figure S3A.** Kinetic studies of early stages of EphB1-EphrinB2 dimer-tetramer assembly were investigated using BLI and very short association steps of 5, 10, 15, 20, 40, and 80 secs with AHC immobilized rat EphB1-Fc ectodomain binding to 100 nM soluble mouse EphrinB2-His ectodomain followed by a 500 sec dissociation step. In the 5, 10, and 15 sec association runs, the EphB1-EphrinB2 dimer is observed to form fast with a very high response slope and without any tetramers being formed as the response signal quickly reverses to baseline (indicated by thick red arrows) after the biosensors were placed into dissociation. If the association step is extended to 20 secs, at the time when the high response slope of the dimer shifts to a lower tetramer accumulation slope, then a very small amount of super-stable tetramers are formed that persists through the end of the dissociation step. Note that longer 40 and 80 sec association steps result in progressively greater formation of super-stable tetramers. Binding assays were conducted in PBST.

Defining levels of Eph-Ephrin dimer and tetramer formation based on A20 tetramer inhibition

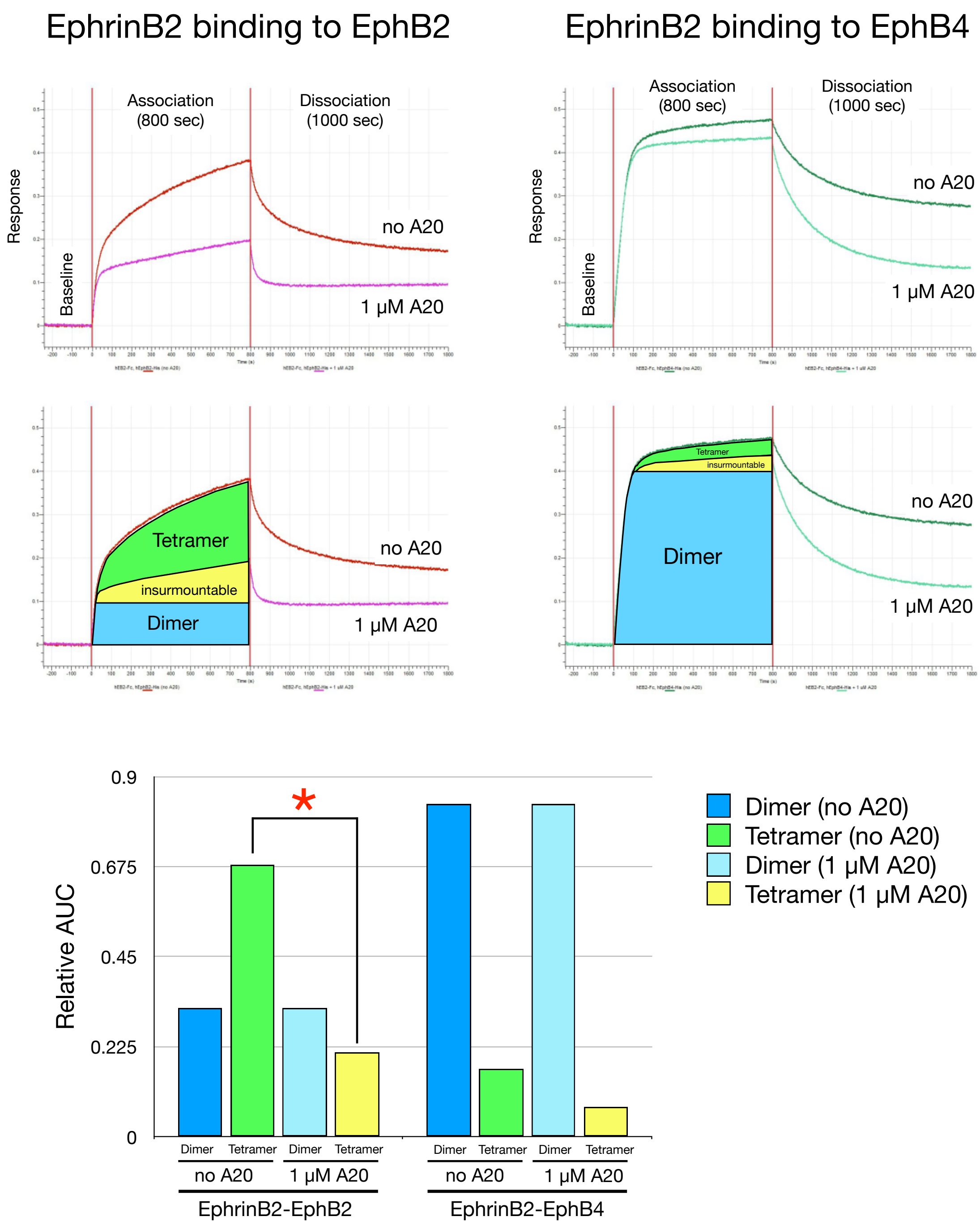

**Figure S3B.** Area under the curve (AUC) analysis of the BLI sensorgram data shown in **Figure 1G** of AHC immobilized human EphrinB2-Fc binding to either soluble human EphB2-His or EphB4-His indicates lead compound A20 exhibits a much greater effect on EphB2-EphrinB2 binding kinetics compared to that of EphB4-EphrinB2. In the presence of 1 µM A20, a large fraction of the EphB2-EphrinB2 interaction is lost during the association step and this can be attributed to reduced formation of the tetramer (green/yellow shading) with no effect on dimer formation (blue/light blue shading). Compound A20 does not prevent all tetramers from forming, a low level of insurmountable tetramers continue to form during the association step in the presence of A20 (yellow shading), and these are very stable tetramers that persist throughout the dissociation step as the response at the end of the Octet run remains significantly above baseline. The insurmountable tetramers likely represent the formation of higher-order tetramer superclusters that are more resistant to A20 inhibition. While A20 has a strong effect on tetramer formation of the EphB2-EphrinB2 interaction, compound shows very little effect on EphB4-EphrinB2 binding kinetics, as this interaction is mainly dimer-driven. Graphing the AUC data indicates the EphB2-EphrinB2 interaction is tetramer-driven, accounting for ~70% of the interaction, and this is reduced to ~20% in the presence of A20 (red asterisk), whereas the EphB4-EphrinB2 interaction is dimer-driven, accounting for ~80% of this interaction. Binding assays were conducted in PBST containing 1% DMSO (PBSTD).

EphrinB2 binding to EphB2 - testing 50, 100, 200, and 400 µM SNEW and EWLS

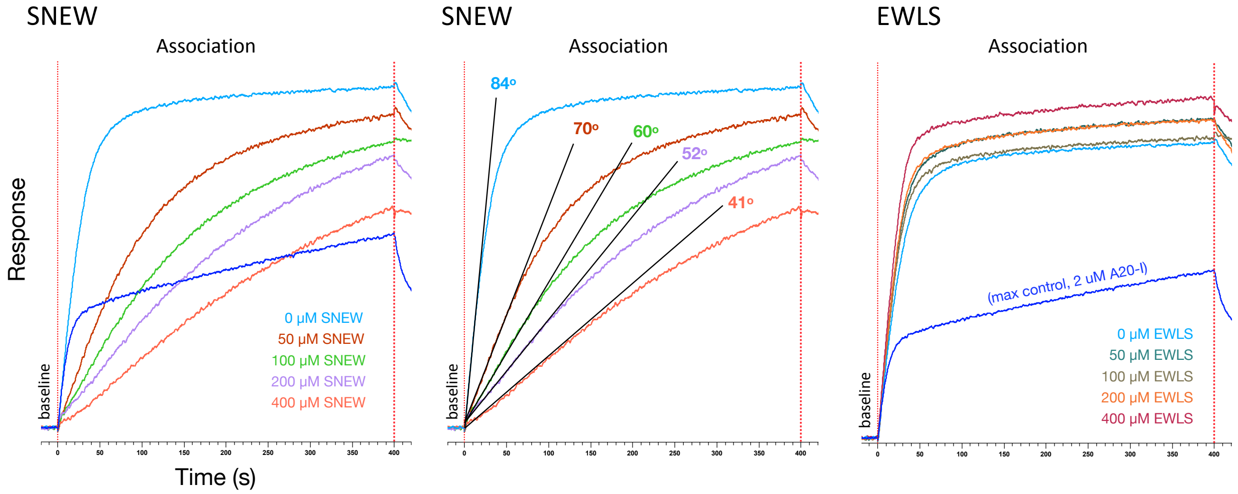

**Figure S3C.** Additional kinetic studies related to **Figure 1H** using higher concentrations of the SNEW and EWLS peptides up to 400 µM show SNEW specifically reduces in a concentration-dependent fashion the dimer slope of AHC immobilized human EphrinB2-Fc ectodomain binding to 100 nM soluble human EphB2-His ectodomain during the 400 sec association step. In contrast, presence of 2 µM of tetramer inhibitor A20-I (dark blue trace) had no effect on dimer binding dynamics during the association step as the initial high slope of response was unaffected, though the compound strongly curtailed the formation and accumulation of tetramers. The EWLS peptide exhibited no effect on the EphrinB2-EphB2 interaction, consistent with this peptide being able to specifically target the EphB1 dimerization interface. Only the baseline and association steps are shown to focus in on the important association dynamics where compound and peptides are present and exert their effects on Eph-Ephrin binding. Binding assays were conducted in PBSTD.

#### A20 is a reversible antagonist of immobilized EphB1 binding to soluble EphrinB2

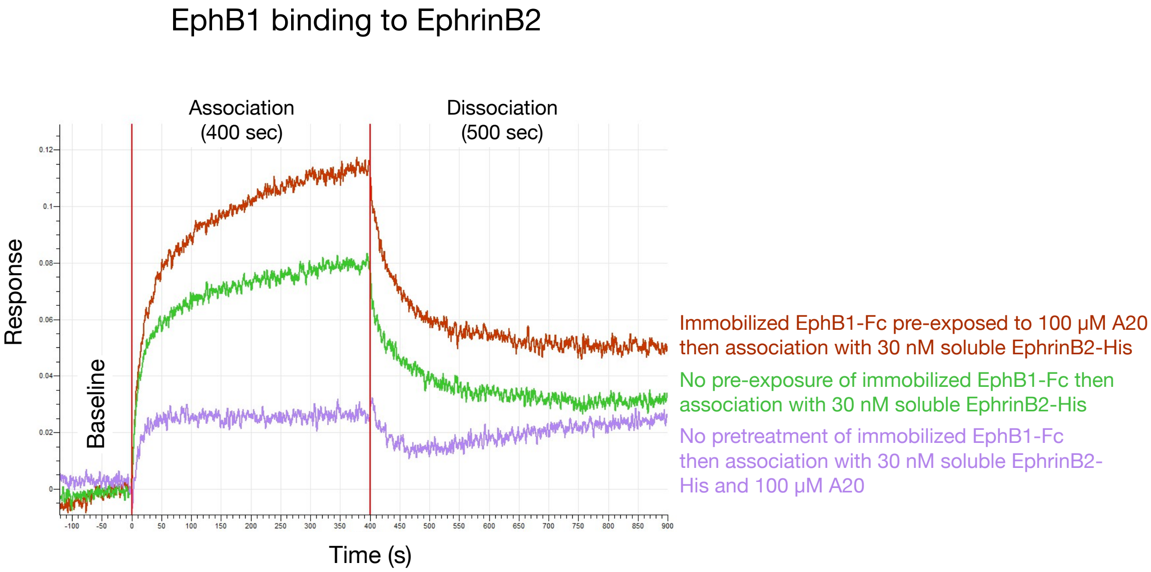

**Figure S3D.** Biophysical studies show A20 is a reversible antagonist of immobilized EphB1 ectodomain binding to soluble EphrinB2 ectodomain. AHC immobilized rat EphB1-Fc was first pre-incubated with either no compound (green sensorgram trace) or with 100 μM of A20 (2xHCl salt) (dark red sensorgram trace). After washing in binding buffer and obtaining baseline measurements, pre-incubated EphB1-Fc was then exposed to 30 nM soluble mouse EphrinB2-His ectodomain in the association step. Pre-exposing EphB1 to A20 did not effect its subsequent ability to bind EphrinB2 (red sensorgram trace), indicating compound acts as a reversible inhibitor of Eph-Ephrin tetramerization. Adding A20 during the association step did, as expected, strongly reduce tetramer formation (purple sensorgram trace). Binding assays were conducted in PBSTD.

#### A20 is a reversible antagonist of immobilized EphrinB2 binding to soluble EphB2

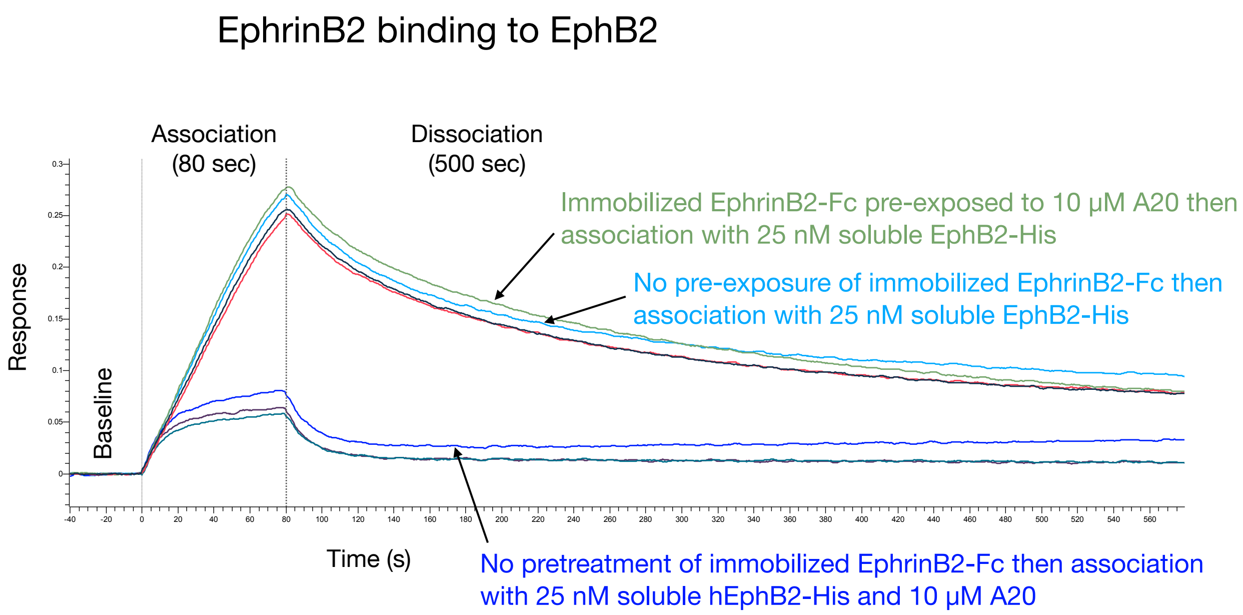

**Figure S3E.** Biophysical studies show A20 is a reversible antagonist of immobilized EphrinB2 ectodomain binding to soluble EphB2 ectodomain. AHC immobilized human EphrinB2-Fc was first pre-incubated with either no compound (light blue sensorgram trace) or with 10 µM of A20 (2xHCl salt) (green sensorgram trace). After washing in buffer and obtaining baseline measurements, pre-incubated EphrinB2-Fc was then exposed to 25 nM soluble human EphB2-His in the association step. Pre-exposing EphrinB2 to A20 had no effect on its subsequent ability to bind EphB2 and form tetramers (green sensorgram trace), indicating compound acts as a reversible inhibitor of Eph-Ephrin tetramerization. Adding A20 during the association step did, as expected, strongly reduce tetramer formation (dark blue sensorgram trace). Binding assays were conducted in PBSTD.

A20 is a competitive antagonist of immobilized EphB1 binding to soluble EphrinB2

**a**

#### EphB1 binding to EphrinB2

### 0 µM A20

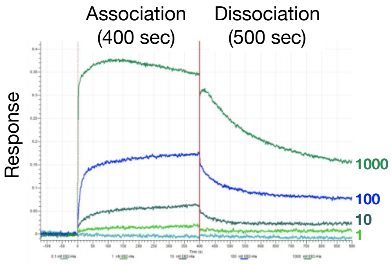

EphrinB2 binding at 0.1-1000 nM with no A20

↓

EphrinB2 binding detected at 1, 10, 100, and 1000 nM

### 10 µM A20

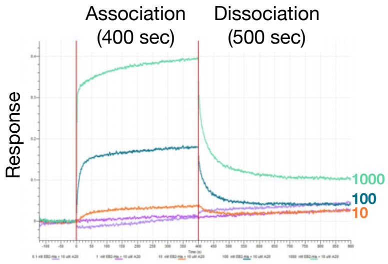

EphrinB2 binding at 0.1-1000 nM with 10 µM A20

↓

EphrinB2 binding only detected at 10, 100, and 1000 nM

### 100 µM A20

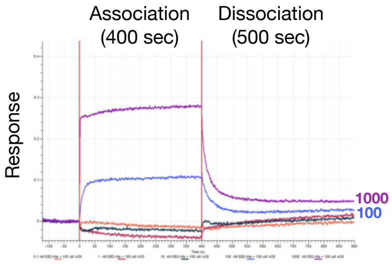

EphrinB2 binding at 0.1-1000 nM with 100 µM A20

↓

EphrinB2 binding only detected at 100 and 1000 nM

**b**

#### EphB1 binding to EphrinB2

##### 0.1 nM EphrinB2

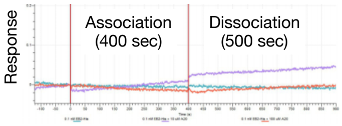

no EphrinB2 binding at 0.1 nM

##### 1 nM EphrinB2

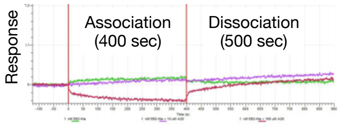

Very low EphrinB2 binding at 1 nM that is competed away with both 10 µM and 100 µM A20

##### 10 nM EphrinB2

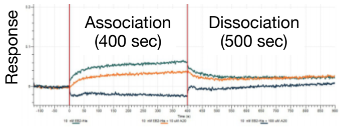

Medium EphrinB2 binding at 10 nM that is competed slightly by 10 µM and completely by 100 µM A20

##### 100 nM EphrinB2

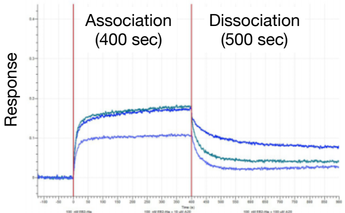

High EphrinB2 binding at 100 nM that is competed away slightly with 10 µM and more so with 100 µM A20

##### 1000 nM EphrinB2

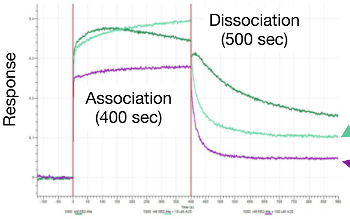

Binding of very high concentrations of EphrinB2 protein can be competed away with as little as 10 µM A20.

And even more so with 100 µM A20.

Very high EphrinB2 binding at 1000 nM that is competed away slightly with 10 µM and more so with 100 µM A20

**Figure S3F.** A20 is a competitive antagonist of EphrinB2 binding to immobilized EphB1. **a**, In the absence of A20, SSA immobilized biotinylated rat EphB1-Fc ectodomain protein can bind soluble mouse EphrinB2-His ectodomain protein when presented at 1, 10, 100, and 1,000 nM concentrations, with no binding detected at 0.1 nM (left), in the presence of 10 µM A20 (2xHCl salt) the binding of EphrinB2 is reduced and detected at only 10, 100, and 1,000 nM (middle), and with 100 µM A20 the binding of EphrinB2 can only be detected with 100 and 1,000 nM of the ligand protein (right). **b**, Grouping the sensorgrams based on the concentration of soluble EphrinB2 ligand used in the association step shows that addition of A20 competes with and reduces EphrinB2 binding to the immobilized EphB1 protein. As increasing concentrations of A20 diminished the ability of EphB1 to bind EphrinB2, the data indicates compound acts as a competitive antagonist. Binding assays were conducted in PBSTD.

#### Pre-existing stable Eph-Ephrin tetramers are also targeted by A20

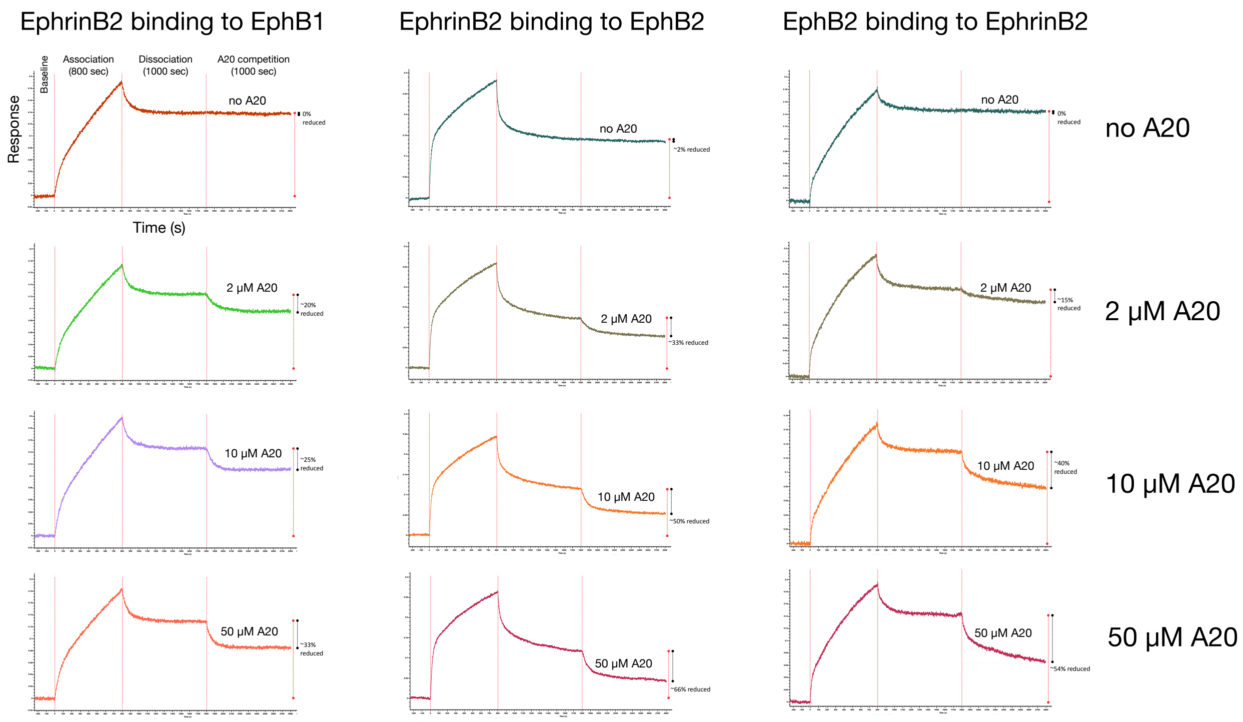

**Figure S3G.** Pre-existing stable Eph-Ephrin tetramers are also targeted by A20. AHC immobilized human EphrinB2-Fc or mouse EphB2-Fc ectodomains were first subjected to a 800 sec association step with 50 nM soluble human EphB1-His and EphB2-His or mouse EphrinB2-His ectodomains followed by a 1,000 sec dissociation step to pre-form the super-stable tetramer species. Biosensors were then placed into a 1,000 sec competition step in the absence or presence of the indicated concentration of A20 (2xHCl salt). Note that tetramers remained super-stable in the competition step without any A20 as the biosensors showed little if any loss of response, whereas addition of A20 produced a concentration-depended loss of tetramers as indicated by the reduced sensorgram response signals. Binding assays were conducted in PBSTD.

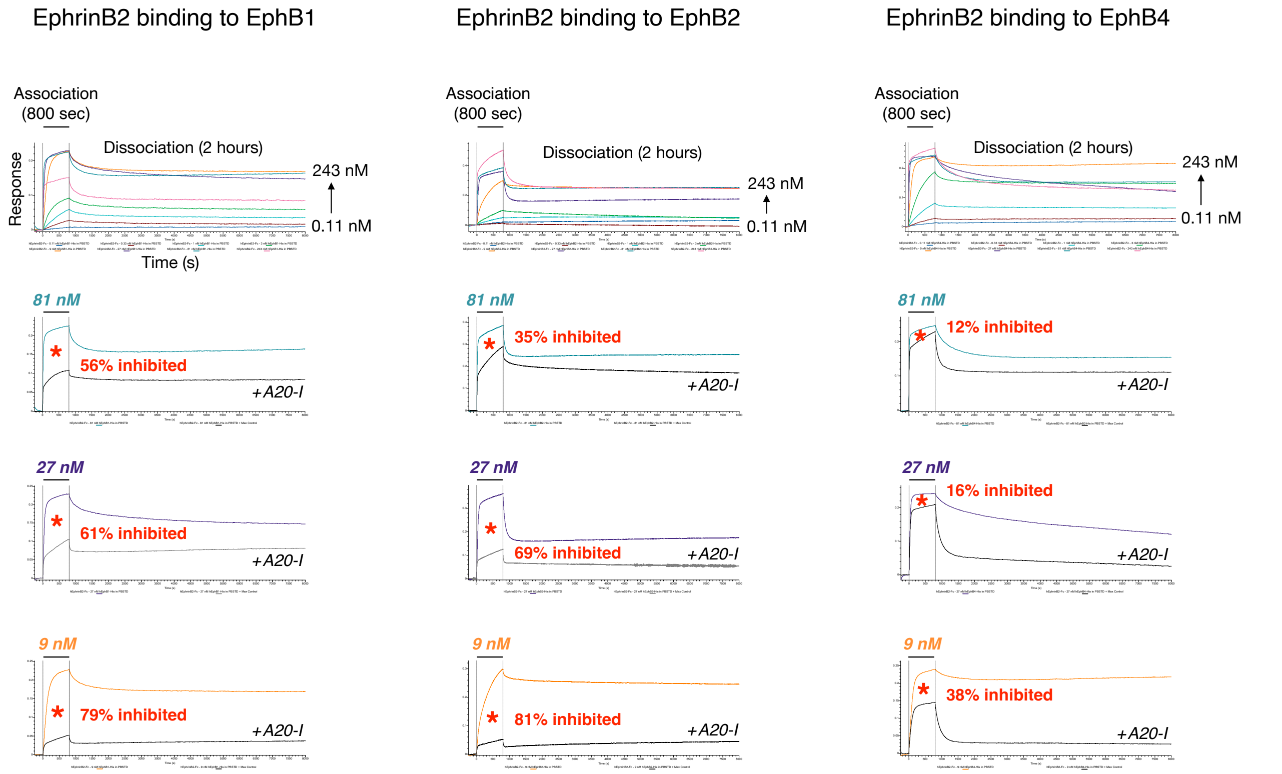

**Figure S3H (related to Figures 1D and S1F).** Many of the global fit studies shown above using multiple concentrations of soluble human EphB1-His, EphB2-His, EphB4-His binding to immobilized human EphrinB2-Fc also tested the addition of tetramer inhibitor A20-I. Shown are example sensorgram traces that included 2.43 μM A20-I with 81 nM Eph protein (tetramer inhibitor:protein molar ratio of 30), 27 nM Eph protein (tetramer inhibitor:protein molar ratio of 90), or 9 nM Eph protein (tetramer inhibitor:protein molar ratio of 270). A20-I strongly inhibited EphB1 and EphB2 binding to EphrinB2 during the association step at all receptor concentrations tested as indicated by the greatly reduced response signals, while compound exhibited less of an effect on the more dimer-driven EphB4-EphrinB2 interaction. For each test, the association step AUC was determined for both the run without compound and the run with compound, and then a % inhibition was calculated (red asterisk).

**Tetramer inhibitor A20-I strongly reduced binding of 27 nM EphB1, EphB2, and EphA4 to immobilized EphrinB2, with significantly less effect on EphB4 binding**

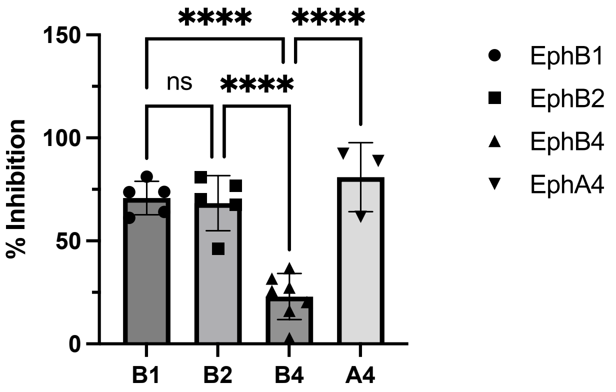

**Tetramer inhibitor A20-I strongly reduced binding of 9 nM EphB1 and EphB2 to immobilized EphrinB2, with significantly less effect on EphB4 binding**

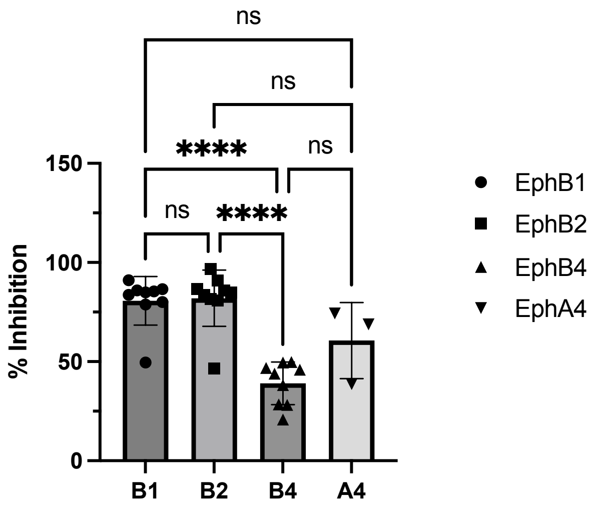

**Figure S3I (related to Figure S3H).** Compound A20-I strongly disrupts the ability of EphrinB2 to bind and associate with EphB1, EphB2, and EphA4, though shows significantly less effect on binding to EphB4. A large number of independent Octet experiments tested immobilized human EphrinB2-Fc binding to 27 or 9 nM soluble human EphB1-His, EphB2-His, EphB4-His, or EphA4-His ectodomains in the absence or presence of 2.43 μM A20-I. For each test, the association step AUC was determined for both the run without compound and the run with compound, and then a % inhibition was calculated and plotted. While A20-I was able to inhibit ~75% of the total interaction of EphrinB2 with EphB1, EphB2, and EphA4, one-way ANOVA shows a highly significant reduced ability of compound to inhibit the EphrinB2-EphB4 interaction.

EphrinB2 binding to EphB4

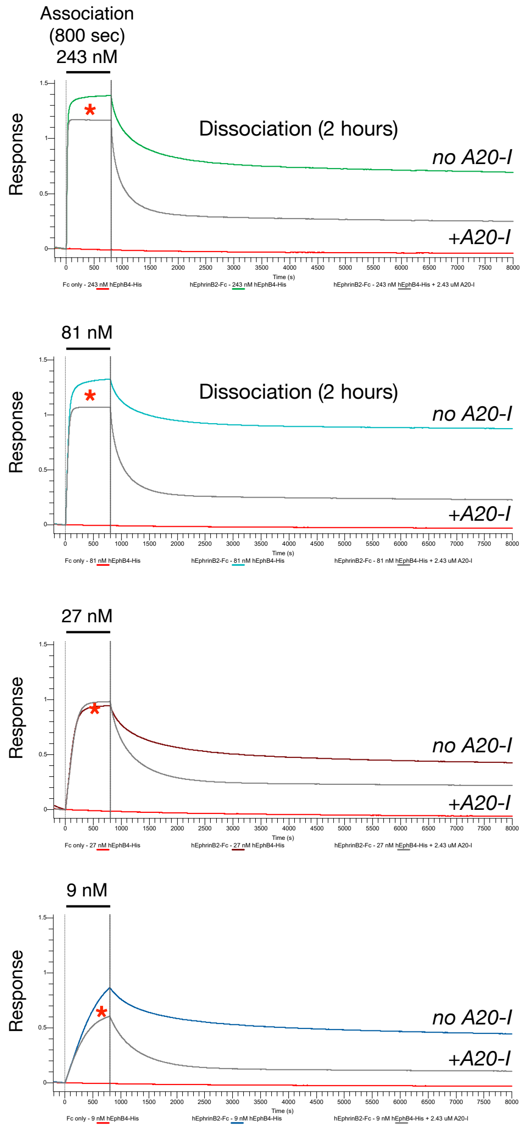

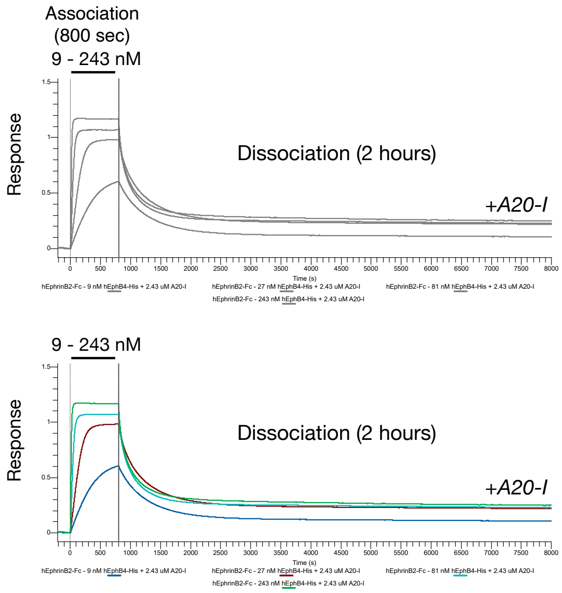

| 1:1 model | 2:1 model |
|---|---|
| $K_D = 0.57$ nM | $K_{D1} = 0.185$ nM |
| $k_a = 5.83E+05$ | **$K_{D2} = 19.22$ nM** |
| $k_{dis} = 3.34E-04$ | $k_{a1} = 2.02E+05$ |
| $R^2 = 0.6281$ | $k_{a2} = 2.02E+05$ |
| | $k_{dis1} = 3.74E-05$ |
| | $k_{dis2} = 3.92E-03$ |
| | $R^2 = 0.9883$ |

**Figure S3J (related to Figures 1F and 1G).** Global fit calculations of the EphrinB2-EphB4 dimer binding affinity. Shown are sensorgrams of AHC2 immobilized human EphrinB2-Fc binding to 243, 81, 27, and 9 nM of soluble human EphB4-His ectodomains in the absence (colored traces) or presence of 2.43 μM A20-I (grey traces) for 800 sec association step and 2 hr dissociation step (left sensorgrams). Fc control protein was also loaded on AHC2 sensors and showed no binding to the EphB4-His ectodomain (red traces). The presence of A20-I led to a greatly flattened slope of response during the association step, especially noted for the runs testing 243 and 81 nM EphB4-His protein, indicating curtailed accumulation of tetramers. Grouping the runs containing A20-I for global fit analysis (right sensorgrams, bottom is color coded) revealed the 2:1 model with $R^2$ of 0.9883 provided the best fit and with a $K_{D2}$ for the dimer of 19.2 nM.

BLI studies show EphrinA ligands also exhibit complex binding interactions with EphA receptors

#### EphA3 and EphB2 binding to EphrinA1 and EphrinA5

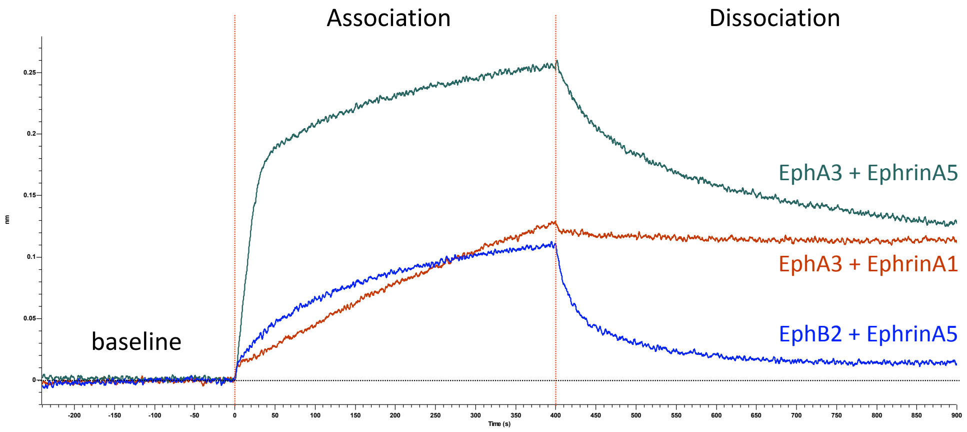

| | | 1:1 binding | 2:1 heterologous binding | |
|---|---|---|---|---|
| **EphrinA** | **Eph** | **KD** | **KD1** | **KD2** |
| **EphrinA1** | EphA3 | 2.79 nM (0.9914) | 2.97 pM | 7.51 nM (0.9915) |
| **EphrinA5** | EphA3 | 2.54 nM (0.9334) | <1 pM | 4.85 nM (0.988) |
| **EphrinA5** | EphB2 | 353 nM (0.9354) | 696 nM | 1.37 uM (0.9794) |

**Figure S3K.** BLI studies of A-subclass and cross A/B-subclass Eph/Ephrin interactions also indicates complex 2:1 heterologous type formation of dimers and tetramers. AHC immobilized human EphA3-Fc ectodomain binding to 50 nM soluble human EphrinA1-His and EphrinA5-His ectodomains, and immobilized mouse EphB2-Fc binding to 50 nM soluble human EphrinA5-His ectodomain. The calculated $K_D$ values for the affinity of both 1:1 and 2:1 type interactions is provided with $R^2$ information.

A20 tetramer inhibitors also target EphrinA ligands binding to EphA3 and EphB2 receptors

EphA3 + EphrinA1

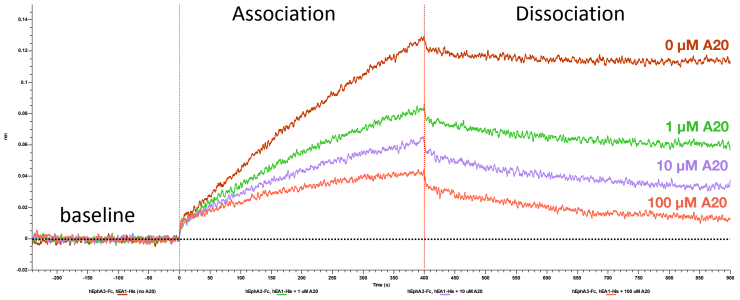

EphA3 + EphrinA5

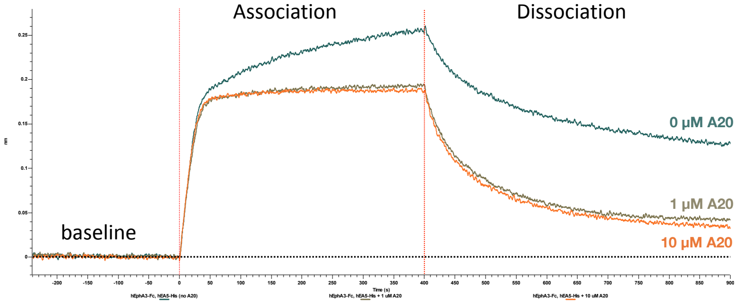

EphB2 + EphrinA5

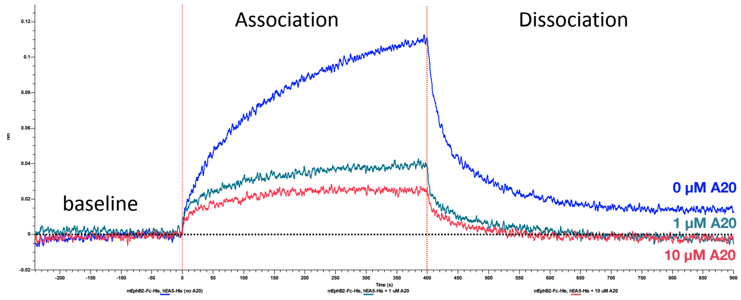

EphB2 + EphrinA5

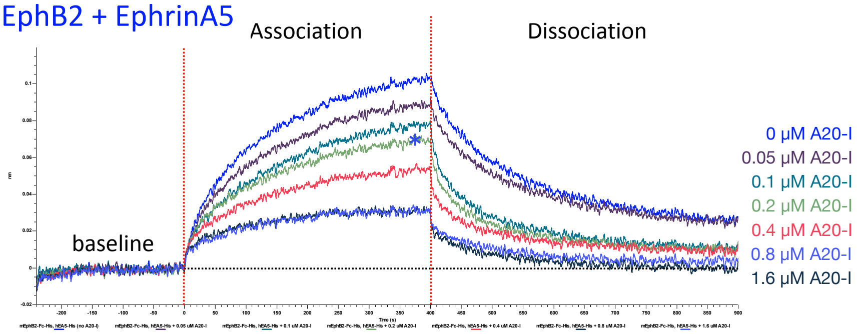

**Figure S3L.** BLI studies of EphA3-EphrinA1 and EphA3-EphrinA5 interactions indicates lead compound A20 can also target tetramers formed by A-subclass molecules, as well

as tetramers formed by the cross A/B-subclass EphB2-EphrinA5 interaction. AHC immobilized human EphA3-Fc ectodomain binding to 50 nM soluble human EphrinA1-His and EphrinA5-His ectodomains, and AHC immobilized mouse EphB2-Fc binding to 50 nM soluble human EphrinA5-His ectodomain. Curiously, the EphA3 receptor exhibits distinct binding kinetics when comparing its interaction with EphrinA1 versus EphrinA5. Here, EphA3-EphrinA1 appears mainly tetramer-driven with a very short/small dimer phase/slope (similar to EphB2-EphrinB1), while the EphA3-EphrinA5 interaction is mainly dimer-driven with a very long/tall dimer phase/slope (similar to EphB4-EphrinB2).

### Summary of HTS and medicinal chemistry campaign

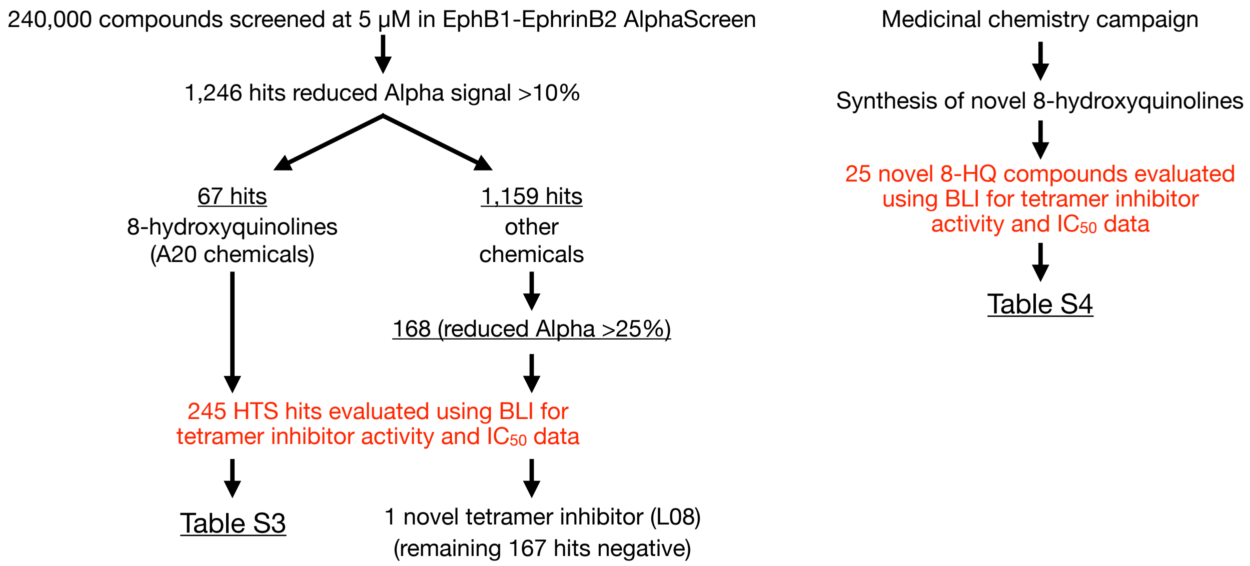

**Figure S4A.** Summary of high-throughput screen (HTS) and medicinal chemistry campaign that led to discovery of A20 and additional related small molecules that antagonize the Eph-Ephrin tetramer. A total of 1,246 primary hits were identified in the HTS that reduced the EphB1-EphrinB2 Alpha signal >10%, including A20 and an additional 66 compounds related to A20 (>5% of the primary hits) that all contained an 8-hydroxyquinoline (8-HQ) bicyclic ring structure known to act as a bidentate metal chelator (***for more details see the accompanying Khambete et al. manuscript***). The top 245 hits from the Alpha-based HTS, including the 67 different 8-HQ compounds, were subsequently assessed using BLI and it was determined that all but one of the 67 different 8-HQ compounds exhibited strong Eph-Ephrin tetramer inhibitor activity (Table S2). Further, of the other 168 non-8-HQ compounds that reduced Alpha signal >25% in the HTS, BLI identified one additional compound, termed L08, that acted as a potent tetramer inhibitor (***L08 is further described in the accompanying Khambete et al. manuscript***). An additional 25 novel 8-HQ compounds were synthesized in a medicinal chemistry campaign and tested using BLI, finding that many of these were also potent tetramer inhibitors (Table S3).

### Generation of the A20 salt

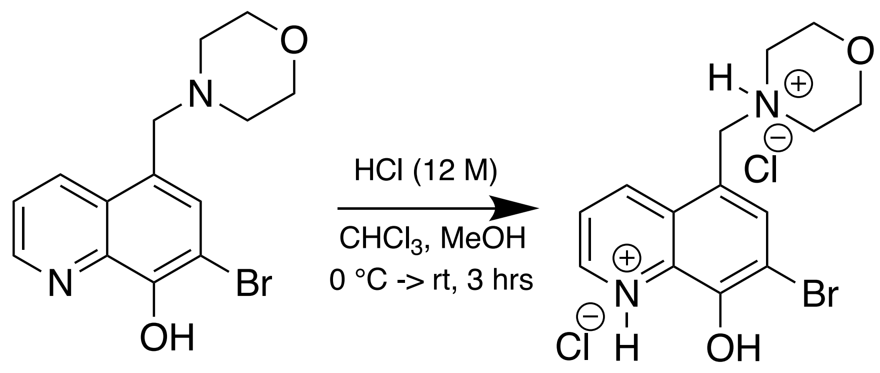

A20 (free-base)
(MW = 323.19)

A20.2xHCl (salt)
(MW = 396.20)

**Figure S4B.** Generation of the A20 salt. To a round bottom flask equipped with a stir bar, free-base A20 (0.200 g, 0.62 mmol) was added and subsequently dissolved in $CHCl_3$ (5 mL). While stirring the solution at 0 °C, concentrated (12.1 M) HCl (106 uL, 1.27 mmol) diluted in methanol (2 mL) was added dropwise over 5 min and then continued to stir over ice for 30 min as yellow precipitate forms, and then allowed to warm to room temperature for 2.5 hr. The resulting precipitate was filtered in ambient conditions over a Buchner funnel, washed with $CHCl_3$, and collected as a pale yellow solid (yield = 215 mg, 88 %).

### Analogs based off A20 and 3511 scaffolds

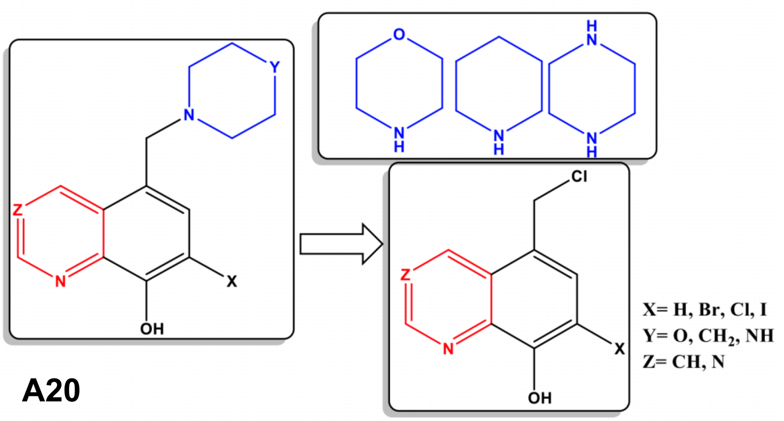

**A20**

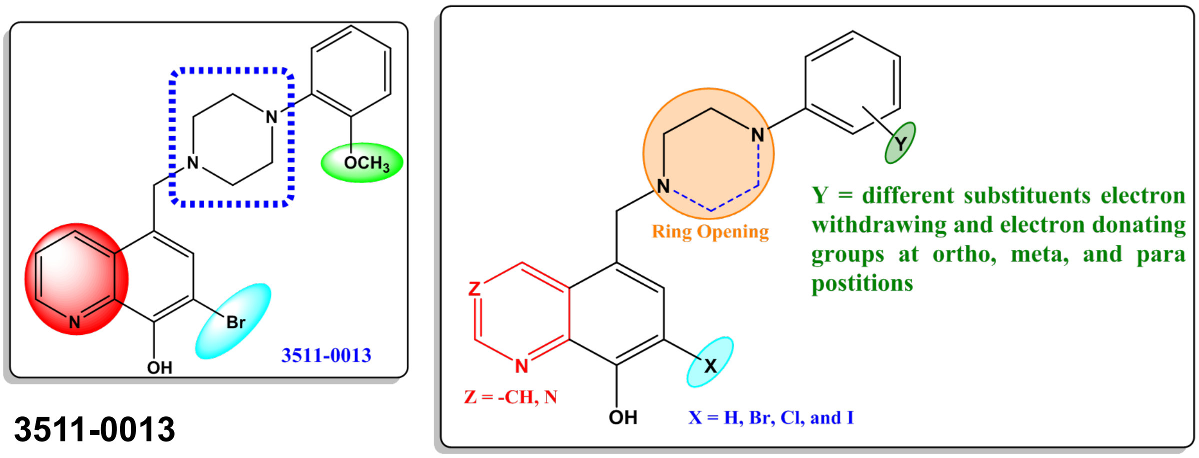

**3511-0013**

**Figure S4C.** Analogs based off A20 and 3511-0013 scaffolds.

Early first-generation medicinal chemistry effort to generate novel A20 analogs

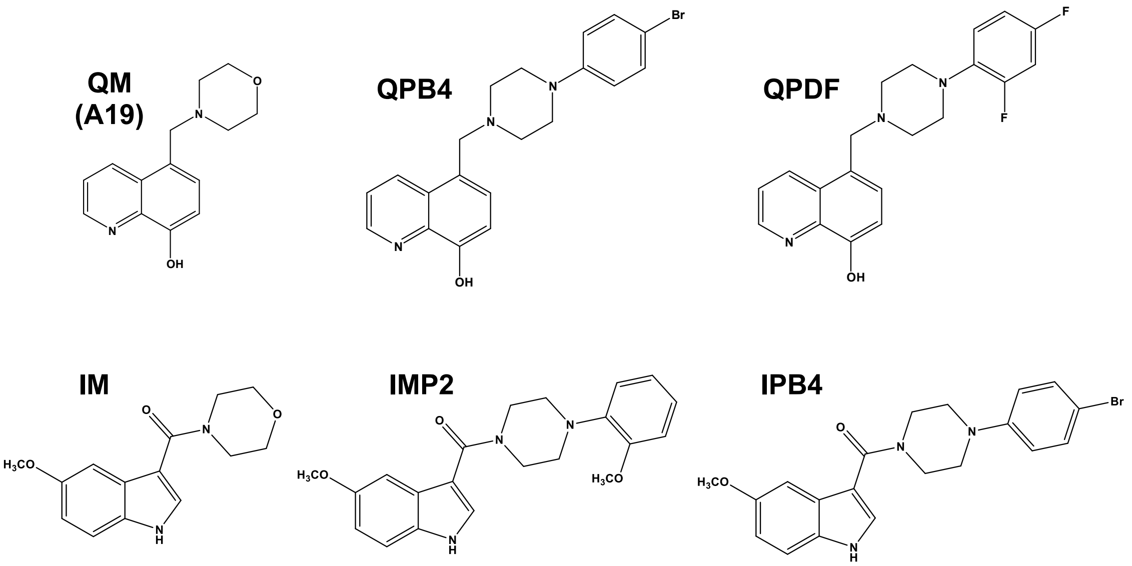

**Figure S4D.** First-generation medicinal chemistry effort to generate novel A20 analogs with extensions off 8-HQ position 5, and synthesis of 5-methoxyindole ring substitutions.

### Alpha testing of first-generation A20 analogs

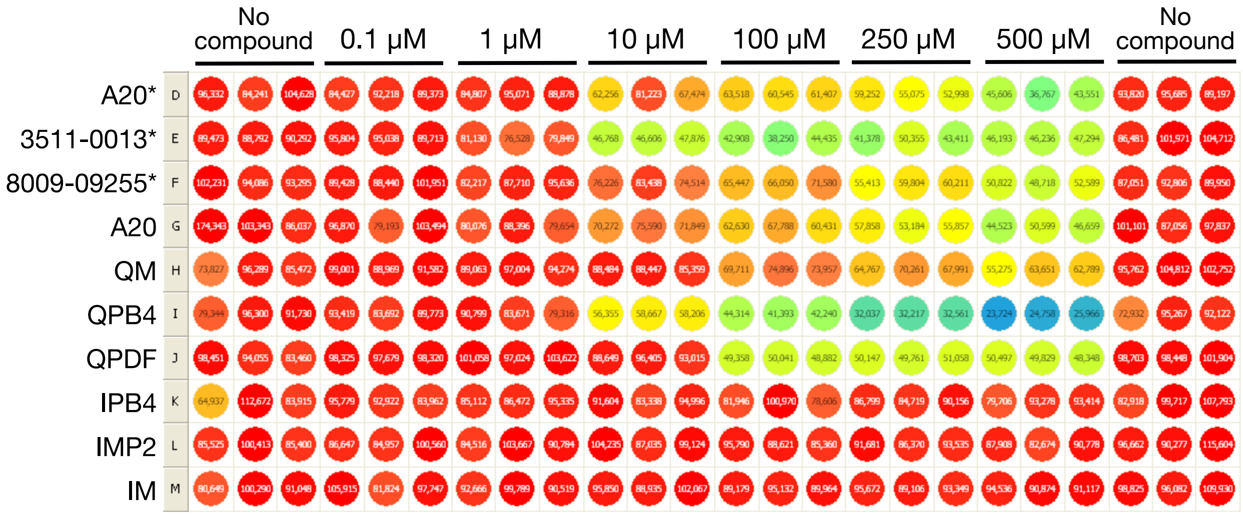

* store-bought compounds

**Figure S4E.** EphB1-EphrinB2 Alpha assay on available A20 related 8-HQ analogs, novel first-generation 8-HQ analogs, and novel indole ring substituted compounds (all free bases). Conditions included 8 nM of EphB1-Fc and EphrinB2-His proteins, 1% DMSO, 50 ng each bead per well, and indicated concentration of compound. QPB4 and 3511-0013 exhibited the strongest ability to reduce the Alpha signal, while the indole-substituted compounds showed no activity.

Compound synthesis schemes for first-generation compounds A20-I, 3511-I, and BQPB4

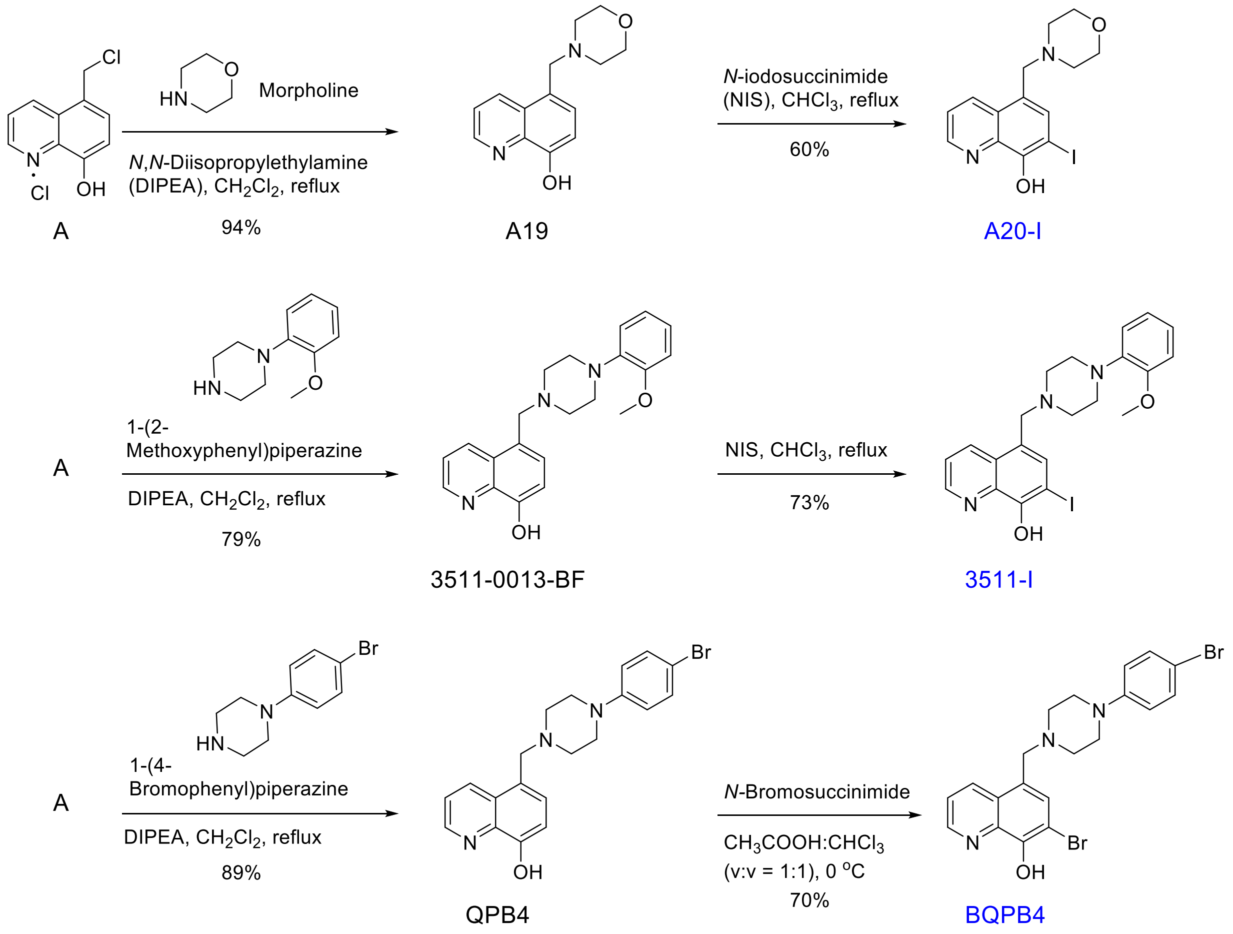

**Figure S4F.** Compound synthesis schemes for first-generation compounds A20-I, 3511-I, and BQPB4. Additional synthetic and purification details are provided in the Supplementary Chemistry Methods.

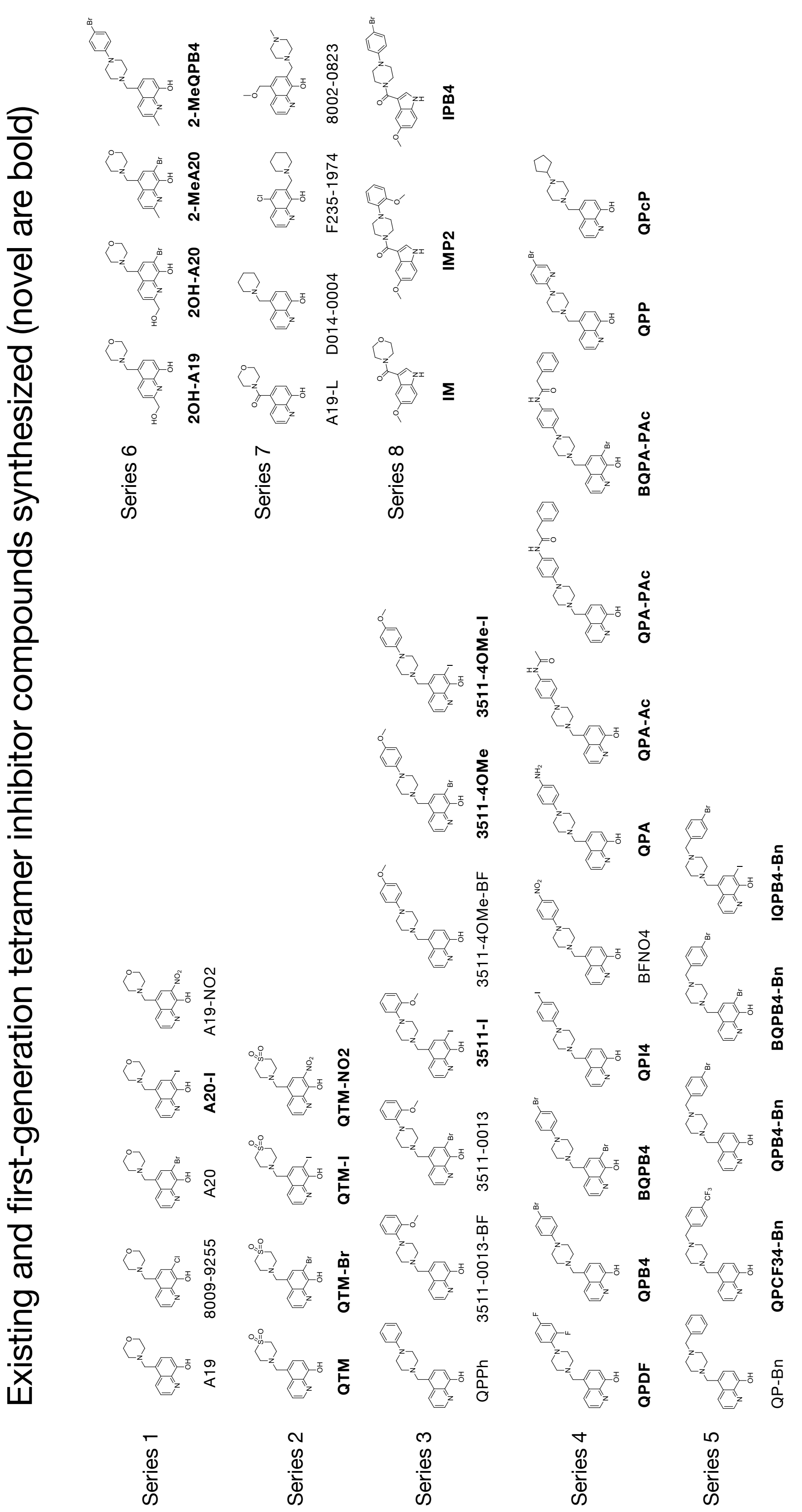

**Figure S4G.** First-generation Eph-Ephrin tetramer inhibitors (novel compounds in bold). A series of novel A20-related analogs were designed, synthesized, and tested using

BLI. Tables S2 and S3 provide tetramer inhibitor activities and additional information.

BLI studies to determine the total percent inhibition of EphrinB2-EphB interactions caused by A20-I

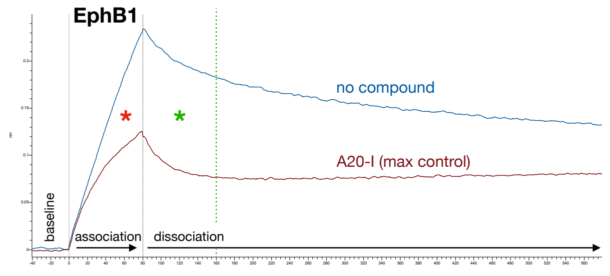

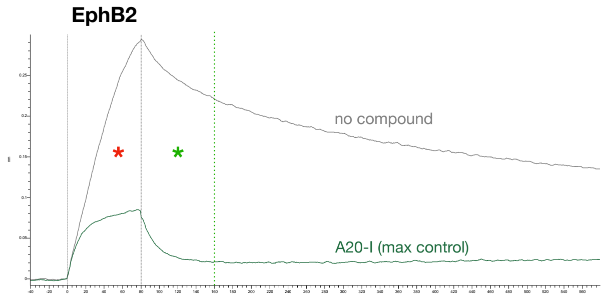

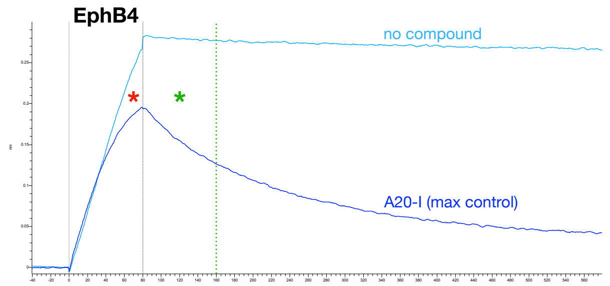

**% Inhibited (AUC)**

| | Association (*) | Association (*) + Dissociation (*) |
|---|---|---|
| EphB1 + A20-I | 36.6 | 48.5 |
| EphB2 + A20-I | 61.8 | 76.5 |
| EphB4 + A20-I | 13.9 | 33.9 |

**Figure S4H.** BLI kinetic studies were used to determine the overall percent inhibition of Ephrin-Eph protein-protein interactions caused by A20 compounds during the association and dissociation steps. Shown are three example sensorgram response traces testing for immobilized human EphrinB2-Fc ectodomain binding to 25 nM soluble human EphB1-His, EphB2-His, and EphB4-His ectodomains for an 80 sec association step and a 500 sec dissociation step in the absence or presence of 3.2 μM novel compound A20-I (2xHCl salt) in PBS with 0.05% Tween-20, 1% DMSO, and +/- indicated chemical. The sensorgram runs with no compound were compared to those with A20-I by determining the area under the curve (AUC) for the 80 sec association step and then calculating the percentage of the interaction that is lost/inhibited when compound is included using the following simple equation: ($AUC_{with\ compound}/AUC_{no\ compound}$). Focusing only on the AUC of the key 80 sec association step when the two proteins are interacting (area identified with a red asterisk), the biophysical data shows that A20-I exhibited greatest ability to reduce the EphrinB2-EphB2 interaction (61.8% of the association interaction was inhibited), followed by the EphrinB2-EphB1 interaction (36.6% of the association interaction was inhibited), and then by the EphrinB2-EphB4 interaction (13.9% of the association interaction was inhibited). In general, during the 80 sec association step, strong tetramer inhibitor chemicals like A20-I exhibit approximately a 60%, 30%, and 15% ability to disrupt binding of the EphrinB2 ligand ectodomain to its cognate EphB2, EphB1, and EphB4 receptor ectodomains, respectively. The AUCs were also calculated to encompass both the 80 sec association step and the first 80 sec of dissociation step (green asterisks). This revealed A20-I inhibited 76.5% of the EphrinB2-EphB2 interaction, 48.5% the EphrinB2-EphB1 interaction, and only 33.9%

the EphrinB2-EphB4 interaction. The % Inhibited AUC calculations determined for A20-I and other tetramer inhibitor compounds are provided in Table S3.

BLI studies to determine the $IC_{50}$ values for tetramer inhibitors A20-I and BQPB4

# A20-I

EphB1

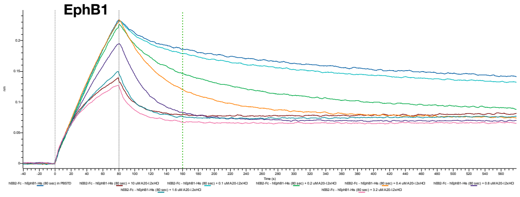

EphB2

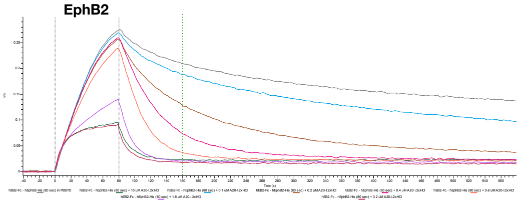

EphB4

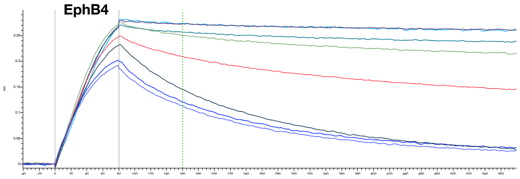

**% Inhibited (AUC)**

| | Association (*) | Association (*) + Dissociation (*) |
|---|---|---|
| EphB1 + A20-I | 32.6 | 49.4 |
| EphB2 + A20-I | 56.4 | 74.7 |
| EphB4 + A20-I | 15.4 | 37.0 |

**IC50 (uM)**

| **EphB1 + A20-I** | Absolute | Relative |
|---|---|---|
| Association (75") | >3.2 | 0.85 |
| Association AUC | >3.2 | 0.98 |
| Dissociation (135") | 0.85 | 0.44 |
| Full AUC | 1.48 | 0.24 |
| **EphB2 + A20-I** | | |
| Association (75") | 1.66 | 1.20 |
| Association AUC | 2.24 | 1.15 |
| Dissociation (135") | 0.41 | 0.34 |
| Full AUC | 0.24 | 0.19 |
| **EphB4 + A20-I** | | |
| Association (75") | >3.2 | 1.70 |
| Association AUC | >3.2 | 1.66 |
| Dissociation (135") | 3.02 | 0.98 |
| Full AUC | 1.26 | 0.89 |

### BQPB4

EphB1

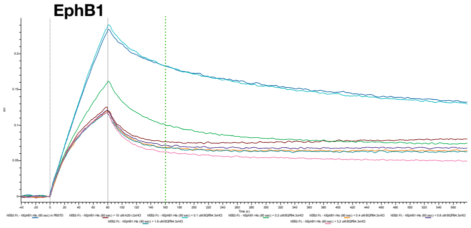

EphB2

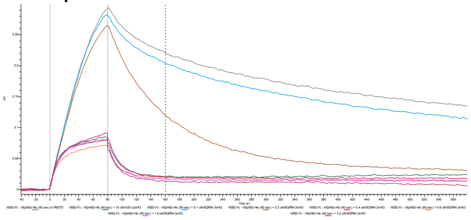

EphB4

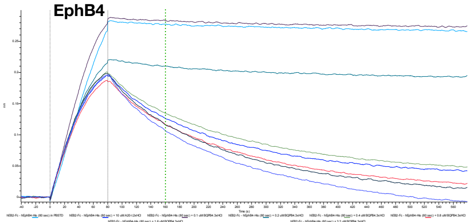

**% Inhibited (AUC)**

| | Association (*) | Association (*) + Dissociation (*) |
|---|---|---|
| EphB1 + BQPB4 | 39.1 | 53.1 |
| EphB2 + BQPB4 | 62.5 | 78.7 |
| EphB4 + BQPB4 | 12.5 | 36.4 |

**IC50 (uM)**

| **EphB1 + BQPB4** | Absolute | Relative |
|---|---|---|
| Association (75") | >3.2 | 0.10 |
| Association AUC | >3.2 | 0.10 |
| Dissociation (135") | 0.36 | 0.08 |
| Full AUC | 0.50 | 0.17 |
| **EphB2 + BQPB4** | | |
| Association (75 sec) | 0.32 | 0.28 |
| Association AUC | 0.36 | 0.28 |
| Dissociation (135") | 0.25 | 0.23 |
| Full AUC | 0.20 | 0.18 |
| **EphB4 + BQPB4** | | |
| Association (75 sec) | >3.2 | 0.08 |
| Association AUC | >3.2 | 0.12 |
| Dissociation (135") | 0.78 | 0.20 |
| Full AUC | 0.32 | 0.26 |

**Figure S4I.** BLI kinetic studies were also used to determine the Ephrin-Eph tetramer inhibitor $IC_{50}$ values for A20 compounds during association and dissociation steps. Shown are example sensorgram response traces testing for immobilized human

EphrinB2-Fc ectodomain binding to 25 nM soluble human EphB1-His, EphB2-His, and EphB4-His ectodomains for an 80 sec association step and a 500 sec dissociation step, in the absence or presence of 0.1, 0.2, 0.4, 0.8, 1.6, and 3.2 μM novel first-generation compounds A20-I (2xHCl salt) or BQPB4 (3xHCl salt), in PBS with 0.05% Tween-20, 1% DMSO, and +/- indicated chemical. In addition to using AUC calculations of the sensorgrams to obtain % inhibition information for the highest dose of A20 chemical tested (typically 3.2 μM) as shown above in **Figure S4H**, the AUCs for the various concentration-response runs were used to calculate $IC_{50}$ values for the association only (0-80”, Association AUC) or for the combined 80 sec association + 500 sec of dissociation (0-580”, Full AUC). Additional $IC_{50}$ values were also determined by analyzing the sensorgram response level at specific timepoints, such as 75 sec into the association step and at 55 sec into the dissociation step (135”). Both absolute and relative $IC_{50}$ values were also determined. Absolute values are based off the full AUC sensorgram traces and take into account both dimer and tetramer formation, whereas relative values are based off of adjusted AUCs in which the AUC of the max control trace is first subtracted before calculating the $IC_{50}$. As this calculation removes the portion of the Ephrin-Eph binding that is due to dimer dynamics and insurmountable tetramer species not affected by the A20 chemicals, the relative $IC_{50}$ values are more useful as they provide a description of a compound’s concentration needed to specifically disrupt 50% of the tetramer.

With an accurate relative $IC_{50}$ value determined for a selected compound and an accurate $K_D$ value for the interaction under analysis, an inhibition constant (the $K_i$) can further be calculated as $K_i = IC_{50}/(1+ [L]/K_D)$ using the Cheng-Prussof equation where [L] is the concentration of the soluble protein used to determine the $IC_{50}$. The $K_i$ provides a dissociation constant to describe an inhibitor’s activity towards a specific target and gives an estimate of how strongly a competitive inhibitor binds to its target protein, a lower value indicates a stronger binding affinity as it indicates a lower concentration of the inhibitor is needed to occupy 50% of the target protein.

The % Inhibited, absolute/relative $IC_{50}$ values, and $K_i$ values determined for these and other tetramer inhibitor compounds are provided in Tables S2 and S3.

### Example BLI studies to determine the $IC_{50}$ values for tetramer inhibitors

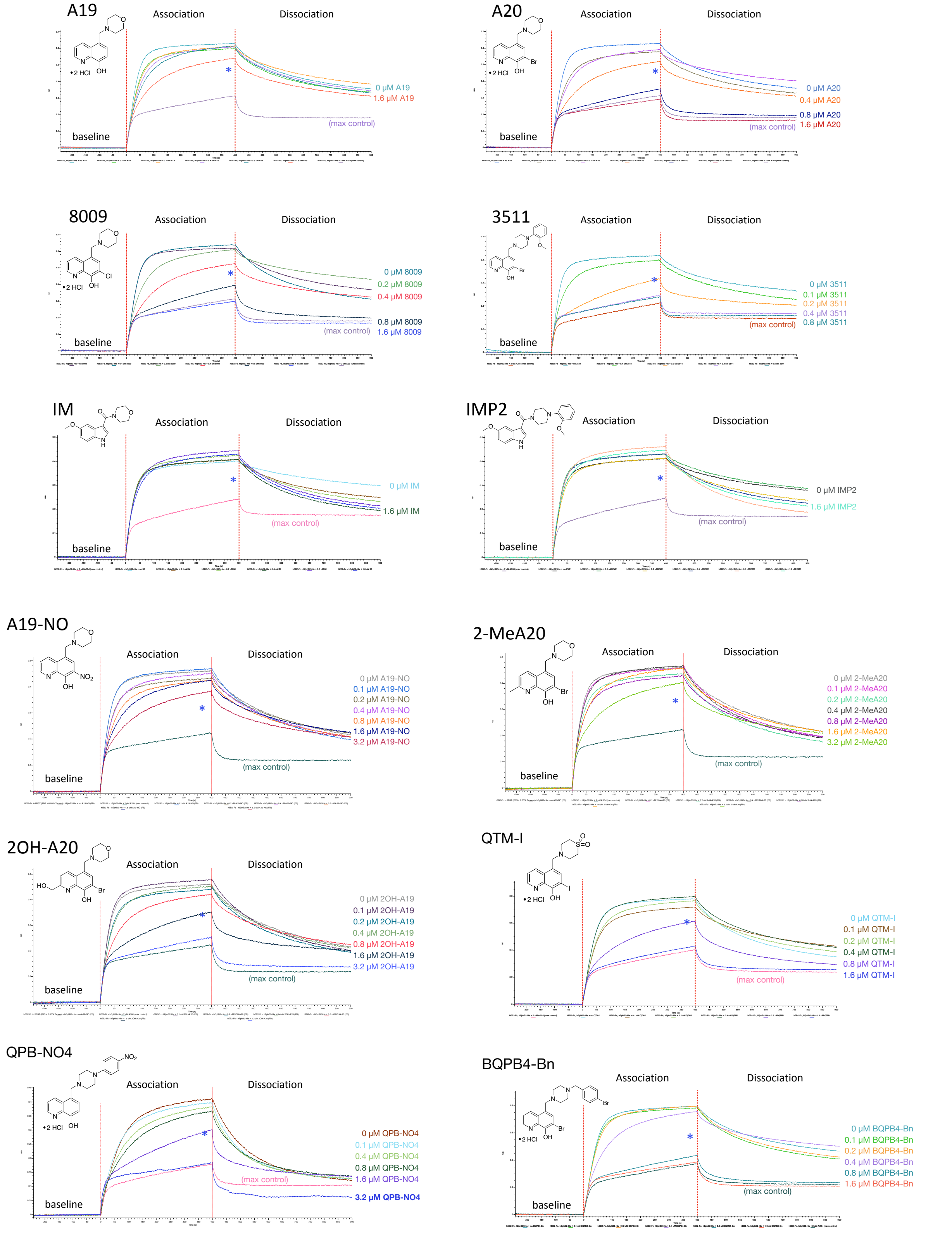

**Figure S4J.** Example BLI kinetic studies used to determine the $IC_{50}$ values for A20

compounds ability to inhibit Ephrin-Eph protein-protein interactions during association and dissociation steps. Shown are sensorgram response traces testing for immobilized human EphrinB2-Fc ectodomain binding to 25 nM soluble human EphB2-His ectodomain for a 400 sec association step and a 500 sec dissociation step, in the absence or presence of 0.1, 0.2, 0.4, 0.8, and 1.6 μM of the indicated compounds in PBS with 0.05% Tween-20, 1% DMSO, and +/- indicated chemical. The blue asterisks denote the 50% point in the tetramer-specific (relative) response measured at the 1/2 point between the no compound sensorgram response and the max control (A20-I) sensorgram response. The absolute/relative $IC_{50}$ values determined by AUC analysis for these and other tetramer inhibitor compounds are presented in Tables S2 and S3.

Find50.py, a new computer program to analyze kinetic data and calculate $IC_{50}$ and $K_i$

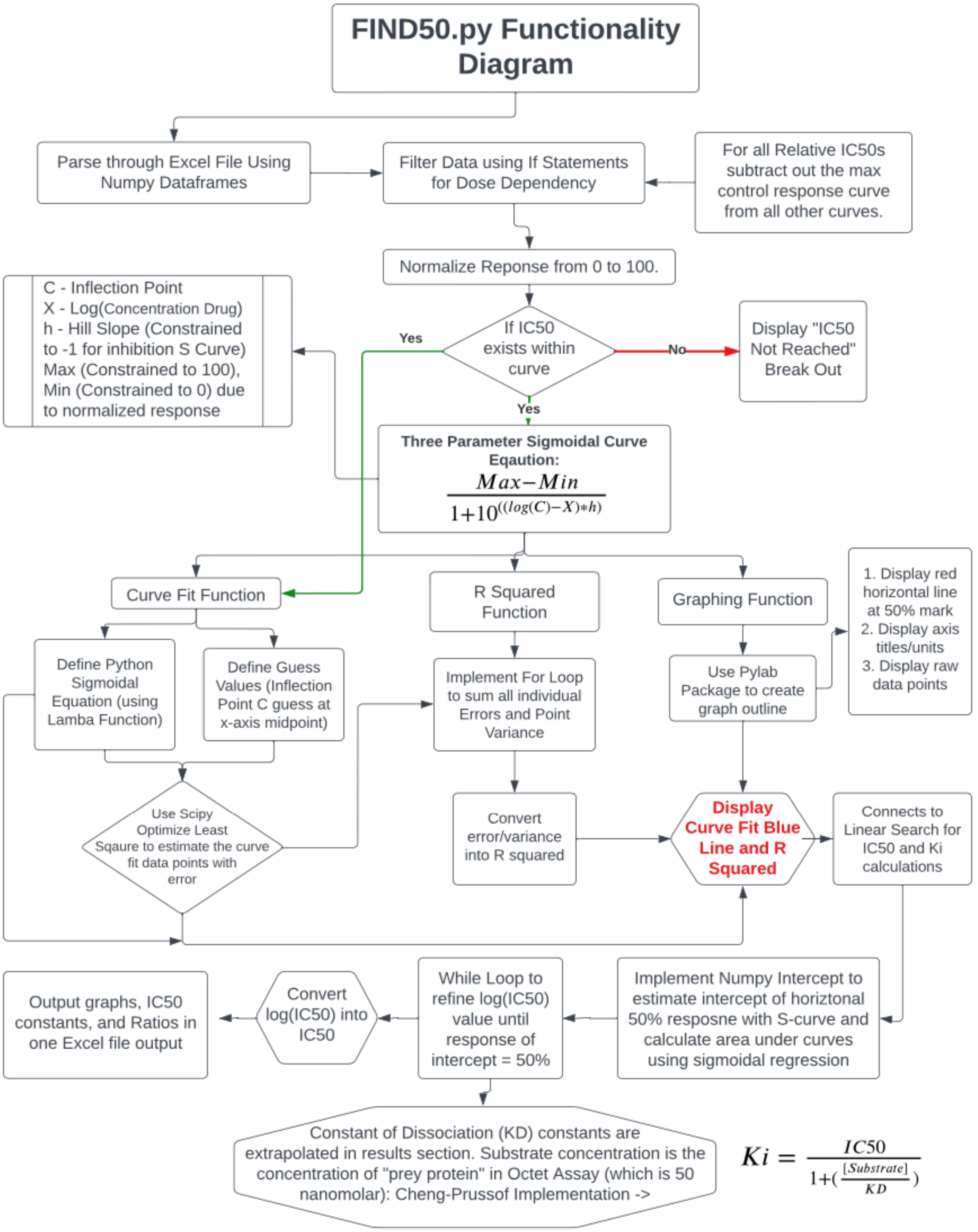

**Figure S4K.** New computer program: Find50.py, used to analyze biophysical kinetic data such as that generated by the Octet RED384 instrument and calculate $IC_{50}$ values using AUC analysis. Find50.py can further be utilized with other types of dose/concentration-dependent data that generates curves needing AUC analysis and $IC_{50}$ calculations. FIND50.py can further calculate the Inhibition Constant ($K_i$), which is an estimate of the binding affinity of an inhibitor for its target protein based off the Cheng-Prussof equation, and represents the concentration of a compound needed to occupy 50% of the binding sites of the target protein or to achieve 50% inhibition of a protein's activity. This equation defines the theoretical relationship between the $IC_{50}$ and $K_i$ of an inhibitory compound based on the $K_D$ of the interaction it is inhibiting and provides a useful constant to describe a specific inhibitor's activity towards its target, though the numerical value can vary with different target proteins (e.g. EphB1 vs EphB2) or by experimental parameters. In summary, FIND50.py allows for full analysis of BLI data or any other similar type data requiring AUC analysis (including 6 different types of $IC_{50}$, $K_i$, and 2 types of ratio calculations, and will display these calculations in a comprehensible, concise manner.

#### Aqueous solubility and in vitro plasma stability of A20 and A20-I

##### aqueous solubility

| | Solubility at pH 7 (mg/mL) |
|---|---|
| A20 (HCl)2 | 1033 |
| A20-I (HCl)2 | 156 |

##### *in vitro* plasma stability

| | Plasma T½ (min) | Saline T½ (min) |
|---|---|---|
| **3511-I.2HCl** | 990.0 | >1440 |
| **A19.2HCl** | 462.0 | >1440 |
| **A19-NO2.2HCl** | >1440 | >1440 |
| **A20.2HCl** | >1440 | >1440 (1732 min) |
| **A20-I.2HCl** | >1440 | >1440 (1732 min) |
| **BQPB4.3HCl** | >1440 | >1440 |
| **QTM-br.2HCl** | 630.0 | 33.5 |
| **QTM-I.2HCl** | 1155.0 | 39.6 |
| **QPB4-bn.2HCl** | 866.3 | 330.0 |
| **QTM-NO2.2HCl** | >1440 | 462.0 |

**Figure S5A.** Aqueous solubility tests show A20 was especially soluble and can be made into a >1% solution. For *in vitro* plasma stability, 10 compounds (2 mM in DMSO) were separately incubated at 37°C in mouse female CD1 pooled plasma (BioIVT, Lot MSE460968) for 0, 10, 30, 60, 120, 240, and 1440 min, quenched with 2:1 ratio of methanol containing formic acid 0.4%, and 100 ng/ml n-benzylbenzamide, and then cleared supernatants analyzed using a 4500 LC-MS/MS.

### *In vitro* metabolic stability of A20 compounds

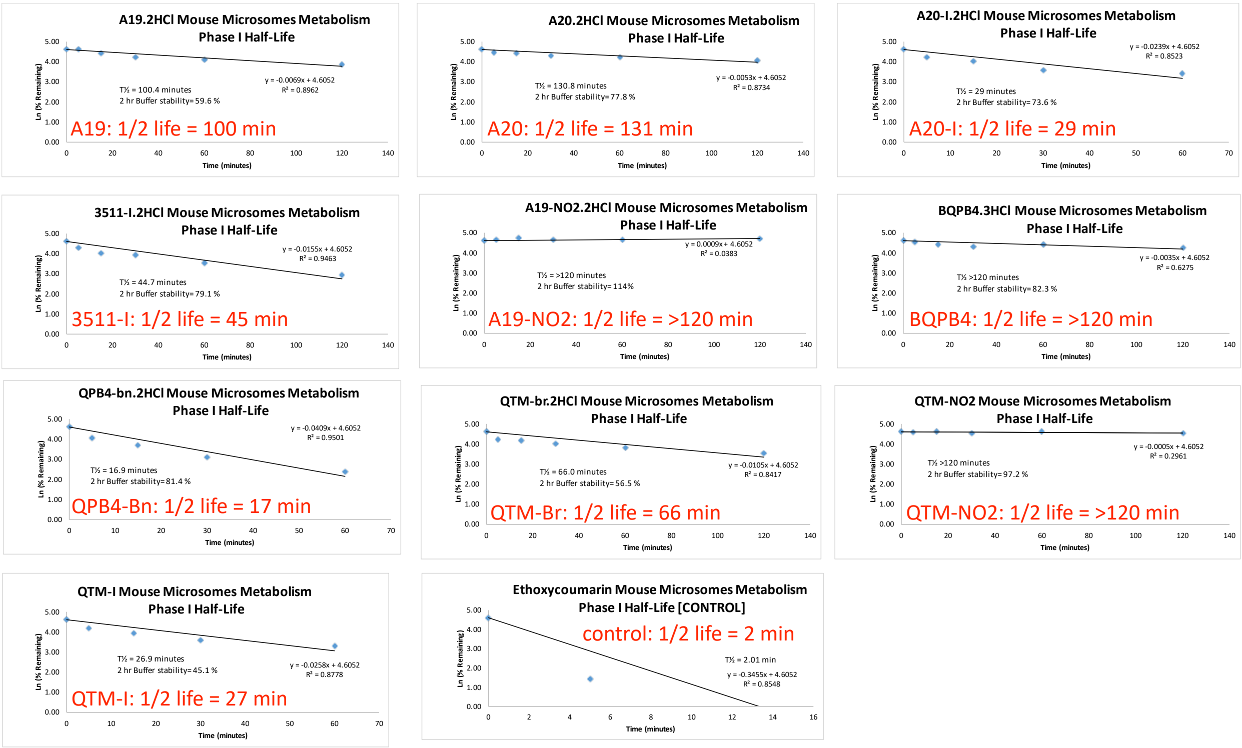

**Figure S5B.** *In vitro* metabolic stability of A20 compounds. 10 compounds (2 mM in DMSO) were separately incubated at 37°C with 0.5 mg/ml mouse microsomes (Xenotech, Male CD1 microsomes M1000 Lot 2210246) and Phase I (NADPH Regenerating System) cofactors for 0-120 min at a final concentration of 2 µM. Reactions were quenched with 0.5 mL (1:1) of methanol containing formic acid 0.4%, and 100 ng/ml n-benzylbenzamide IS (0.2% FA and 50 ng/mL final) and cleared supernatants analyzed using a 4500 LC-MS/MS.

Early pharmacokinetic (PK) studies of A20 free base

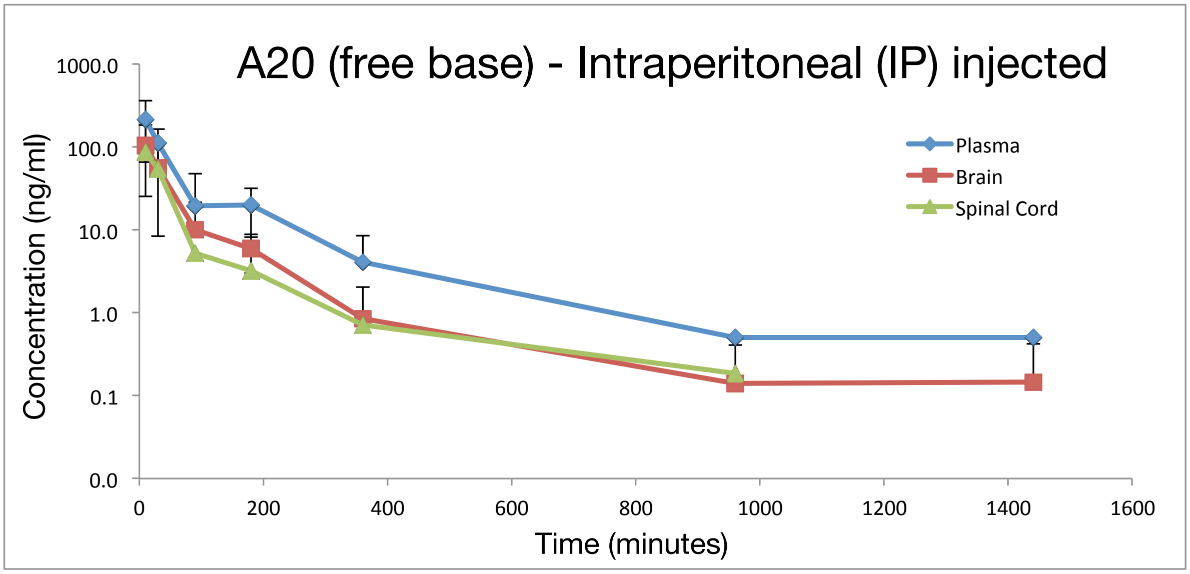

**Figure S5C.** In early pharmacokinetic (PK) studies A20 free base distributed to plasma, brain, and spinal cord following a single intraperitoneal (IP) injection of 20 mg/kg into CD1 mice (vehicle was 100% PBS) and followed for 10, 30, 90, 180, 360, 960, and 1440 min.

### PK studies show the A20 salt is orally bioavailable

**Sample Concentration Data**

**A20 $(HCl)_2$ 3 mg/kg IV & 30 mg/kg PO PKs**

| | IV | | | | | | PO | | | | | | |
|---|---|---|---|---|---|---|---|---|---|---|---|---|---|
| | Plasma | | Brain* | | Liver* | | | Plasma | | Brain* | | Liver* | |
| Time (min) | Conc (ng/mL) | std. dev. | Conc. (ng/g) | Std. Dev | Conc. (ng/g) | Std. Dev | Time (min) | Conc (ng/mL) | std. dev. | Conc. (ng/g) | Std. Dev | Conc. (ng/g) | Std. Dev |
| 5 | 1034 | 92 | 591.5 | 201.6 | 6228 | 623 | 10 | 629 | 84 | 301.8 | 88.2 | 19054 | 6074 |
| 30 | 102 | 59 | 428.3 | 488.2 | 919 | 386 | 30 | 279 | 102 | 103.7 | 27.3 | 6242 | 2328 |
| 90 | 28 | 14 | 0.0 | 0.0 | 622 | 169 | 90 | 125 | 57 | 42.5 | 10.6 | 2675 | 626 |
| 180 | 5.97 | 1.33 | 0.0 | 0.0 | 183 | 11 | 180 | 108 | 29 | 33.1 | 5.5 | 1968 | 290 |
| 360 | 4.21 | 2.41 | 0.0 | 0.0 | 70 | 51 | 360 | 19.27 | 9.35 | 23.2 | 2.8 | 479 | 293 |
| 960 | 1.92 | 1.46 | 3.3 | 5.8 | 9 | 16 | 960 | 2.40 | 0.89 | 0.0 | 0.0 | 73 | 98 |
| 1440 | 2.71 | 0.54 | 9.9 | 0.0 | 7 | 12 | 1440 | 3.06 | 0.90 | 0.0 | 0.0 | 0 | 0 |

**PK Parameters**

| IV | | | | PO | | | |
|---|---|---|---|---|---|---|---|
| 3 mg/kg A20 $(HCl)_2$ IV PK | Plasma | Brain | Liver | 30 mg/kg A20 $(HCl)_2$ PO PK | Plasma | Brain | Liver |
| Terminal T½ (min): | 489 | 29 | 187 | Terminal T½ (min): | 153 | 314 | 175 |
| Tmax (min): | 5 | 5 | 5 | Tmax (min): | 10 | 10 | 10 |
| Cmax (ng/mL or ng/g): | 1034 | 592 | 6228 | Cmax (ng/mL or ng/g): | 629 | 302 | 19054 |
| AUClast (min*ng/mL or min*ng/g): | 30192 | 14617 | 260535 | AUClast (min*ng/mL or min*ng/g): | 54147 | 18026 | 1210206 |
| Vz (mL/kg): | 65896 | 4450 | 3084 | Vz_F (mL/kg): | 120804 | 477807 | 6171 |
| CL (mL/min/kg or g/min/kg): | 93 | 106 | 11 | CL_F (mL/min/kg or g/min/kg): | 547 | 1056 | 24 |
| $MRT_{inf}$ (min): | 223 | 41 | 116 | $MRT_{inf}$ (min): | 183 | 369 | 152 |
| Vss (mL/kg) | 20871 | 4315 | 1321 | | | | |

*A20$(HCl)_2$ average in tissue is calculated by subtracting the amount of compound in the residual blood/plasma within that tissue. The reference for this volume of residual blood for brain is shown below.

| Bioavailability (%) | | |
|---|---|---|
| Plasma | Brain | Liver |
| 17.9 | 12.3 | 46.5 |

**Previous IP Data**

**A20 $(HCl)_2$ 20 mpk IP PKs**

| | Plasma | | Brain* | | Liver* | | Skin* | | Spinal Cord* | |
|---|---|---|---|---|---|---|---|---|---|---|
| Time (min) | Conc (ng/mL) | std. dev. | Conc. (ng/g) | Std. Dev | Conc. (ng/g) | Std. Dev | Conc. (ng/g) | Std. Dev | Conc. (ng/g) | Std. Dev |
| 10 | 2880 | 629 | 2787 | 458 | 22577 | 10392 | 24807 | 4759 | 3310 | 953 |
| 30 | 502 | 133 | 636 | 145 | 9166 | 1444 | 1185 | 318 | 419 | 254 |
| 90 | 43 | 18 | 65 | 17 | 1227 | 639 | 0 | 0 | 58 | 6 |
| 180 | 97 | 75 | 89 | 36 | 2436 | 1428 | 0 | 0 | 67 | 74 |
| 360 | 16 | 7 | 37 | 10 | 502 | 331 | 0 | 0 | 18 | 16 |
| 960 | 0 | 0 | 36 | 8 | 69 | 53 | 0 | 0 | 0 | 0 |
| 1440 | 0 | 0 | 66 | 10 | 18 | 7 | 0 | 0 | 33 | 12 |

**PK Parameters**

| 20 mpk A20 $(HCl)_2$ IP PK | Plasma | Brain | Liver | Skin | Spinal Cord |
|---|---|---|---|---|---|
| Terminal T½ (min): | 69 | 142 | 223 | 5 | 97 |
| Tmax (min): | 10 | 10 | 10 | 10 | 10 |
| Cmax (ng/mL or ng/g): | 2880 | 2787 | 22577 | 24807 | 3310 |
| AUClast (min*ng/mL or min*ng/g): | 81039 | 136933 | 1363490 | 383953 | 86044 |
| Vz_F (mL/kg): | 24226 | 27297 | 4696 | 336 | 31532 |
| CL_F (mL/min/kg or g/min/kg): | 242 | 133 | 15 | 51 | 226 |
| $MRT_{inf}$ (min): | 60 | 483 | 149 | 11 | 57 |

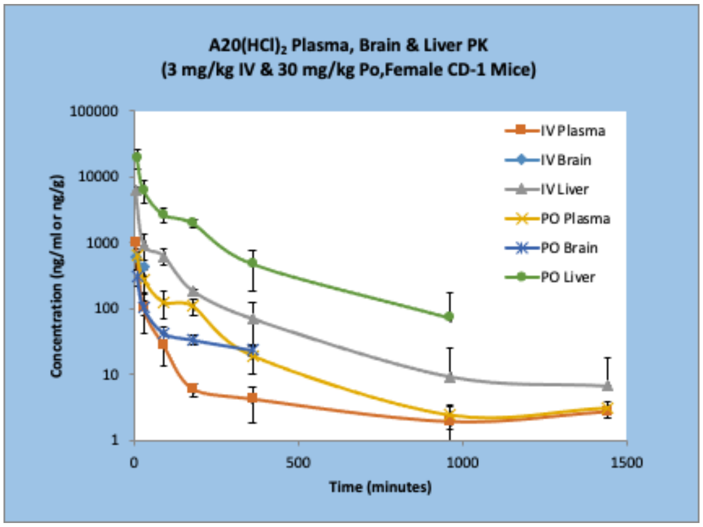

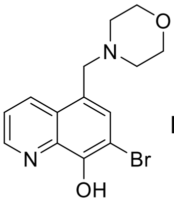

A20
MW = 323.19

| A20 MW = 323.19 g/Mole |
|---|
| 5,000 ng/g = 15.47 µM |
| 1,000 ng/g = 3.09 µM |
| 500 ng/g = 1.55 µM |
| 100 ng/g = 0.31 µM |

**Figure S5D.** A20 is orally bioavailable. Following a single oral dose (PO) or intravenous dose (IV) of 30 mg/kg A20 2xHCl salt dissolved in PBS into CD1 mice and followed for

10, 30, 90, 180, 360, 960, and 1440 min, the compound distributed to plasma, brain, and liver. Relatively high levels of oral-dosed A20 were observe to persist in the liver where it maintained concentrations above the tetramer inhibitor $IC_{50}$ value for this compound (green).

### PK studies show the A20-I salt is orally bioavailable

| A20-I $(HCl)_2$ 30 mg/kg IV & 30 mg/kg PO PKs | | | | | | | | | | | | | |
|---|---|---|---|---|---|---|---|---|---|---|---|---|---|
| IV | | | | | | | PO | | | | | | |
| | Plasma | | Brain* | | Liver* | | | Plasma | | Brain* | | Liver* | |
| Time (min) | Conc (ng/mL) | std. dev. | Conc. (ng/g) | Std. Dev | Conc. (ng/g) | Std. Dev | Time (min) | Conc (ng/mL) | std. dev. | Conc. (ng/g) | Std. Dev | Conc. (ng/g) | Std. Dev |
| 5 | 1073 | 540 | 1426.5 | 740.5 | 7320 | 300 | 10 | 297 | 78 | 368.6 | 147.9 | 34933 | 11097 |
| 30 | 206 | 43 | 143.5 | 22.8 | 744 | 359 | 30 | 73 | 22 | 185.8 | 52.7 | 5696 | 1967 |
| 90 | 14 | 1 | 26.9 | 18.1 | 602 | 187 | 90 | 31 | 21 | 66.9 | 50.8 | 2298 | 1773 |
| 180 | 10.91 | 4.58 | 40.2 | 35.0 | 323 | 145 | 180 | 31 | 15 | 59.0 | 42.0 | 2912 | 1815 |
| 360 | 2.80 | 0.35 | 13.6 | 7.9 | 92.06 | 4.08 | 360 | 19.43 | 5.39 | 59.4 | 24.1 | 878 | 101 |
| 960 | 2.76 | 0.98 | 18.9 | 10.3 | 15.15 | 3.00 | 960 | 4.68 | 3.86 | 9.0 | 3.4 | 70 | 69 |
| 1440 | 2.46 | 0.43 | 54.1 | 1.4 | 26.13 | 9.19 | 1440 | 6.66 | 4.21 | 41.5 | 34.3 | 22 | 9 |

| PK Parameters | | | | | | | |
|---|---|---|---|---|---|---|---|
| IV | | | | PO | | | |
| 3 mg/kg A20-I $(HCl)_2$ IV PK | Plasma | Brain | Liver | 30 mg/kg A20-I $(HCl)_2$ PO PK | Plasma | Brain | Liver |
| Terminal T½ (min): | 113 | 115 | 188 | Terminal T½ (min): | 288 | 273 | 150 |
| Tmax (min): | 5 | 5 | 5 | Tmax (min): | 10 | 10 | 10 |
| Cmax (ng/mL or ng/g): | 1073 | 1426 | 7320 | Cmax (ng/mL or ng/g): | 297 | 369 | 34933 |
| AUClast (min*ng/mL or min*ng/g): | 34286 | 68122 | 309382 | AUClast (min*ng/mL or min*ng/g): | 25572 | 59860 | 1703087 |
| Vz (mL/kg): | 14045 | 6599 | 2569 | Vz_F (mL/kg): | 439486 | 155246 | 3796 |
| CL (mL/min/kg or g/min/kg): | 86 | 40 | 9 | CL_F (mL/min/kg or g/min/kg): | 1059 | 393 | 18 |
| $MRT_{inf}$ (min): | 116 | 529 | 175 | $MRT_{inf}$ (min): | 483 | 767 | 160 |
| Vss (mL/kg) | 10016 | 21052 | 1657 | | | | |

*A20-I$(HCl)_2$ average in tissue is calculated by subtracting the amount of compound in the residual blood/plasma within that tissue. The reference for this volume of residual blood for brain is shown below.

| Bioavailability (%) | | |
|---|---|---|
| Plasma | Brain | Liver |
| 7.5 | 8.8 | 55.0 |

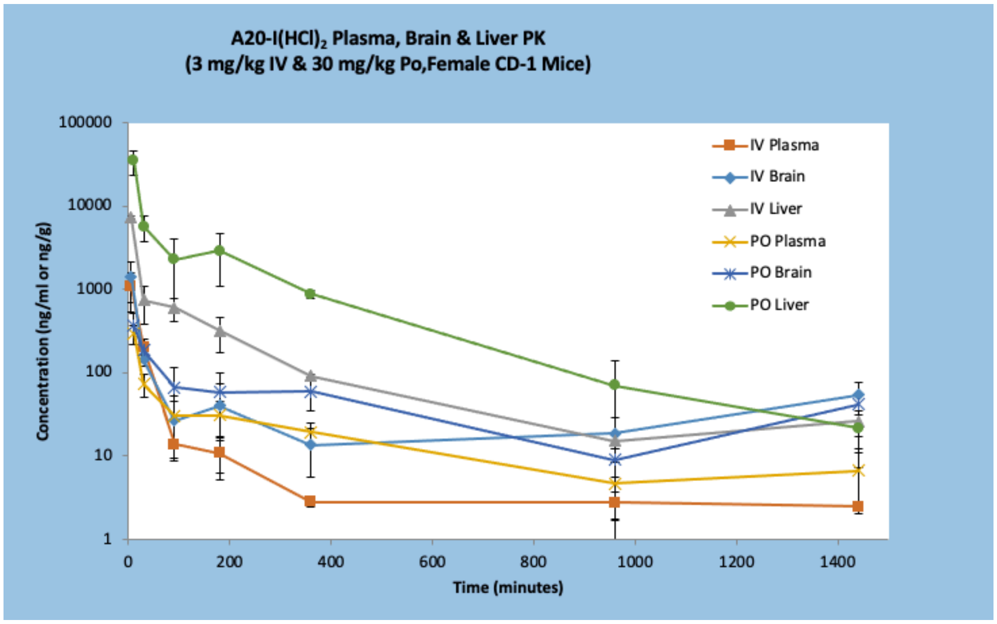

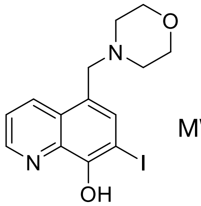

A20-I
MW = 370.19

| A20-I MW = 370.19 g/Mole |
|---|
| 5,000 ng/g = 13.51 µM |
| 1,000 ng/g = 2.70 µM |
| 500 ng/g = 1.35 µM |
| 100 ng/g = 0.27 µM |

**Figure S5E.** A20-I is also orally bioavailable. Following a single oral dose (PO) or intravenous dose (IV) of 30 mg/kg A20-I (2xHCl salt) into CD1 mice and followed for 10,

30, 90, 180, 360, 960, and 1440 min, the compound distributed to plasma, brain, and liver (vehicle was 100% PBS). Relatively high levels of A20-I were observe to persist in the brain and liver where it maintained concentrations close to the tetramer inhibitor $IC_{50}$ values for this compound.

#### PK studies show tetramer inhibitor QPB4-Bn exhibits impressive ADMET properties

| QPB4-Bn(HCl)$_2$ 20 mpk IP PKs | | | | | | | | | | |
|---|---|---|---|---|---|---|---|---|---|---|
| | Plasma | | Brain* | | Liver* | | Skin* | | Spinal cord* | |
| Time (min) | Conc (ng/mL) | std. dev. | Conc. (ng/g) | Std. Dev | Conc. (ng/g) | Std. Dev | Conc. (ng/g) | Std. Dev | Conc. (ng/g) | Std. Dev |
| 10 | 203 | 181 | 1566 | 1636 | 6380 | 1461 | 324 | 219 | 404 | 345 |
| 30 | 99 | 24 | 616 | 161 | 2318 | 1774 | 958 | 234 | 660 | 195 |
| 90 | 43 | 17 | 264 | 92 | 750 | 189 | 709 | 32 | 204 | 115 |
| 180 | 32 | 10 | 206 | 88 | 899 | 191 | 675 | 249 | 214 | 47 |
| 360 | 10 | 4 | 94 | 16 | 363 | 150 | 728 | 311 | 119 | 38 |
| 960 | 5 | 1 | 60 | 2 | 92 | 31 | 56 | 29 | 41 | 17 |
| 1440 | 9 | 1 | 111 | 7 | 108 | 4 | 99 | 32 | 89 | 11 |

| PK Parameters | | | | | |
|---|---|---|---|---|---|
| 20 mpk QPB4-Bn(HCl)$_2$ IP PK | Plasma | Brain | Liver | Skin | Spinal cord |
| Terminal T½ (min): | 128 | 178 | 251 | 201 | 385 |
| Tmax (min): | 10 | 10 | 10 | 30 | 30 |
| Cmax (ng/mL or ng/g): | 203 | 1566 | 6380 | 958 | 660 |
| AUClast (min*ng/mL or min*ng/g): | 23672 | 191753 | 583291 | 539594 | 170635 |
| Vz_F (mL/kg): | 145775 | 23368 | 11657 | 10198 | 50458 |
| CL_F (mL/min/kg or g/min/kg): | 787 | 91 | 32 | 35 | 91 |
| $MRT_{inf}$ (min): | 449 | 632 | 381 | 420 | 814 |
| User's lambda_z bounds (min) | 90-360 | 90-360 | 180-960 | 180-960 | 180-960 |

*QPB4-Bn(HCl)$_2$ average in tissue is calculated by subtracting the amount of compound in the residual blood/plasma within that tissue. The reference for this volume of residual blood for brain is shown below.

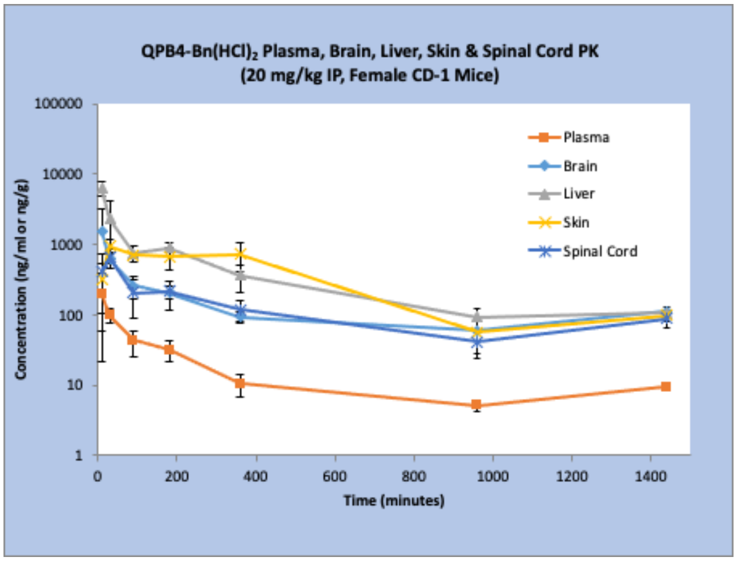

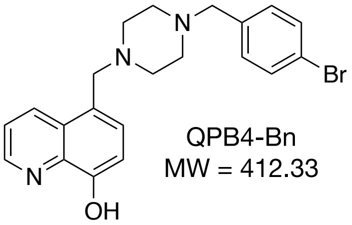

QPB4-Bn
MW = 412.33

| QPB4-Bn MW = 412.33 g/Mole |
|---|
| 5,000 ng/g = 12.13 μM |
| 1,000 ng/g = 2.43 μM |
| 500 ng/g = 1.21 μM |
| 100 ng/g = 0.24 μM |

**Figure S5F.** QPB4-Bn (2xHCl salt form) exhibited strong PK distribution following a single IP injection of 20 mg/kg into mice (vehicle was 2% DMSO, 98% PBS). High levels of QPB4-Bn were observe to persist in the liver, skin, brain, and spinal cord and maintained concentrations close to the tetramer inhibitor $IC_{50}$ values for this compound.

A20 compounds are well tolerated when dosed into mice - Normal nest building behavior

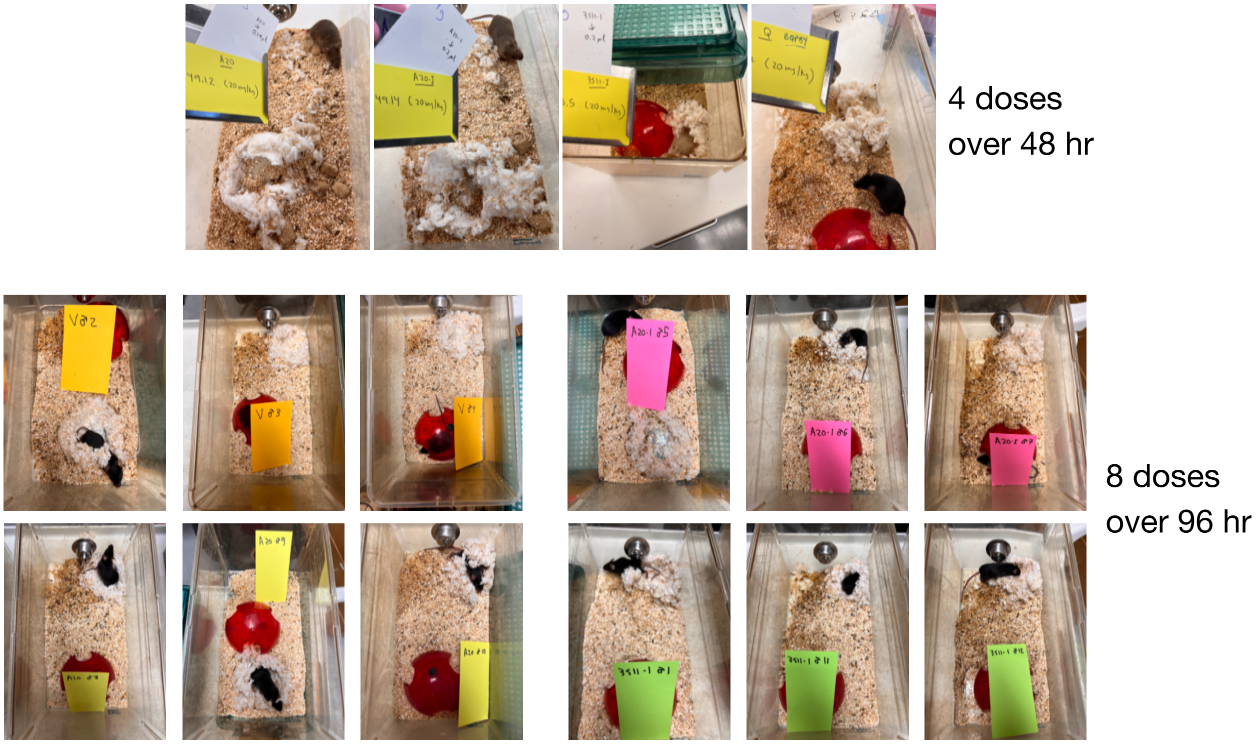

**Figure S5G.** Photos of mice in their cages either 48 hr after starting IP injections of 20 mg/kg doses just before their fifth dose of A20, A20-I, 3511-I, and BQPB4 salts (top), or 96 hr after starting IP injections of 20 mg/kg doses just before their ninth dose of A20, A20-I, and 3511-I salts. Note that the compounds were well tolerated as the dosed animals exhibited no sign of abnormalities or distress, as exhibited by their normal nest building activities. Vehicle for A20 and A20-I (100% PBS), for 3511-I (2% DMSO, 98% PBS), and for BQPB4 (4% DMSO, 96% sunflower seed oil).

PK studies show kinase inhibitor Demeclocycline exhibits impressive ADMET properties

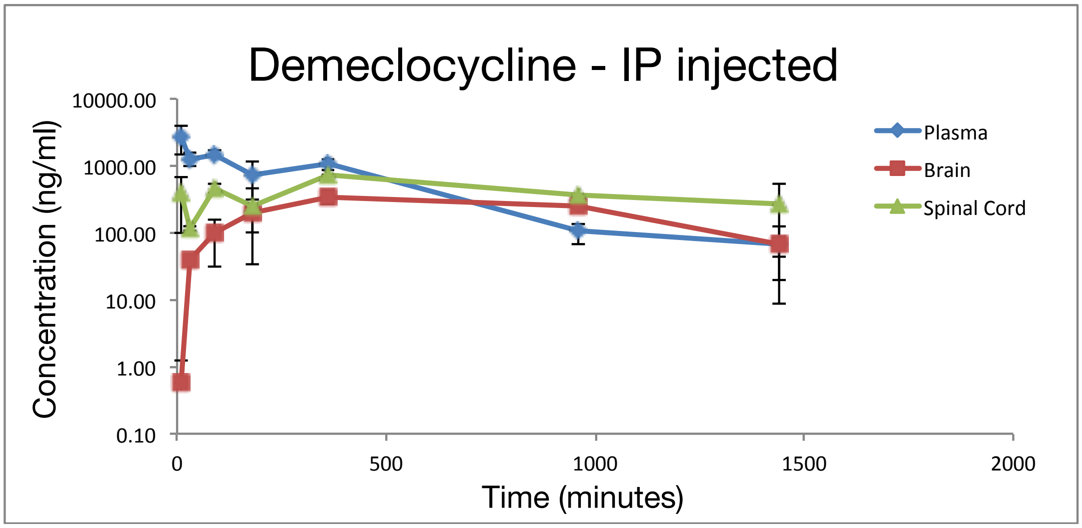

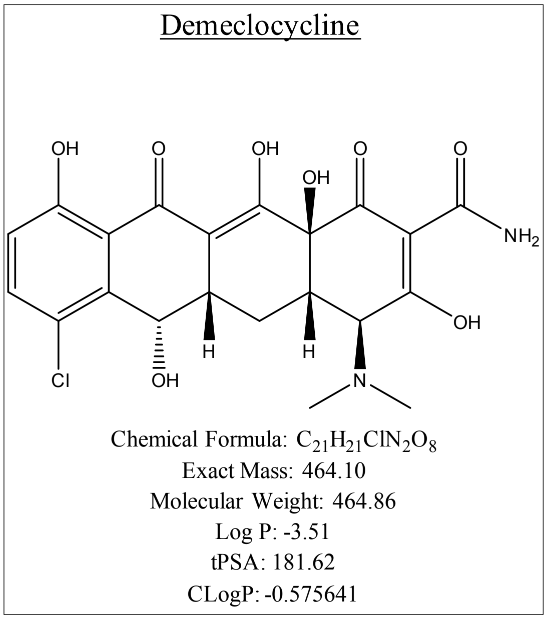

**Figure S5H.** The EphB kinase inhibitor Demeclocycline exhibited strong PK distribution following a single IP injection of 17 mg/kg (vehicle was PBS) into adult CD1 mice and followed for 10, 30, 90, 180, 360, 960, and 1440 min. High levels of Demeclocycline were observed to accumulate and persist in the plasma, brain, and spinal cord.

#### *In vitro* tyrosine kinase assay from adult brains

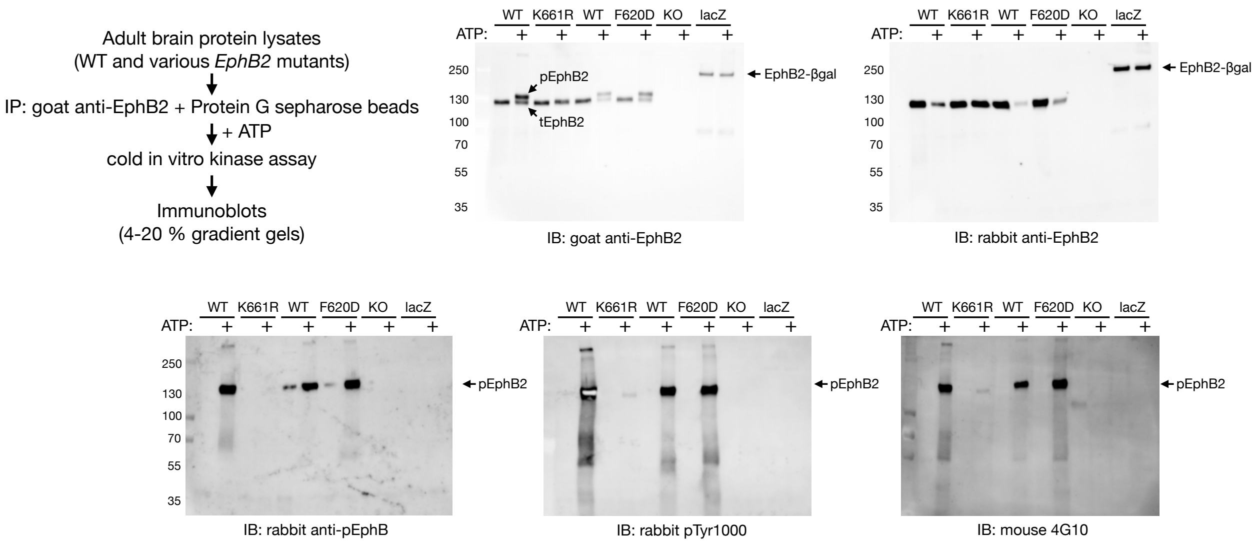

**Figure S6A (Related to Figure 2A).** *In vitro* EphB2 tyrosine kinase assay and validation of antibodies used to show the intracellular juxtamembrane tyrosine residue Y596 of EphB2 becomes autophosphorylated when the catalytic domain is activated by addition of ATP, and this results in reduced mobility of the protein in SDS-PAGE gels. EphB2 protein was immunoprecipitated (IP) by adding goat anti-EphB2 antibodies (R&D Systems, AF467) to adult mouse brain protein lysates made from *EphB2* +/+ wild-type (WT) and four different *EphB2* mutants (K661R kinase-dead point mutant, F620D kinase-overactive point mutant, protein-null knockout [KO], and kinase-deleted intracellular truncated EphB2-βgal fusion protein [lacZ]). Immunoprecipitated protein was then washed and subjected to a cold *in vitro* kinase assay by adding ATP and incubating for 15 min at room temp (RT). Proteins were then resolved under reducing conditions on five 4-20% gradient SDS-PAGE gels, transferred to membranes, immunoblotted (IB) with indicated antibodies, and detected using appropriate HRP-conjugated secondary antibodies and chemiluminescent imaging. The goat anti-EphB2 IB shows addition of ATP to the immunoprecipitates (+) resulted in a shift in migration of >50% the receptor protein from just under 130 KDa for the unphosphorylated tEphB2 protein to a phosphorylated pEphB2 band just above the 130 KDa protein marker, which was present only in the WT and F620D point mutant embryo lysates. Note that the goat anti-EphB2 antibody is raised against the extracellular region of the protein and recognizes both unphosphorylated tEphB2 and tyrosine phosphorylated pEphB2 species, and that no shift following addition of ATP was observed in the K661R kinase-dead point mutant or the larger sized 250 KDa lacZ truncated EphB2-βgal fusion protein that lacks the receptor's intracellular tyrosine kinase catalytic domain. The IB using a rabbit anti-EphB2 antibody raised against the unphosphorylated juxtamembrane segment of EphB2 (Cell Signaling, D2X2I) only recognized the faster migrating unphosphorylated tEphB2 protein (including the EphB2-βgal fusion protein which retains the juxtamembrane segment), whereas the rabbit anti-pEphB2 antibody (Abcam, 61791) raised against the same juxtamembrane peptide of EphB2 though with tyrosine residue 596 (YIDP) phosphorylated specifically recognized the pEphB2 protein which was present only in the WT and F620D IPs subjected to the *in vitro* kinase reaction. Likewise, rabbit pTyr1000 (Cell Signaling, #8954) and mouse 4G10 Platinum (Millipore Sigma, 05-1050), both pan phosphotyrosine antibodies, also detected the pEphB2 protein species, again only in the WT and F620D lysates subjected to the *in vitro* kinase reaction with ATP.

### EphrinB2 stimulated EphB2 forward signaling in Cos1 - time course of activation

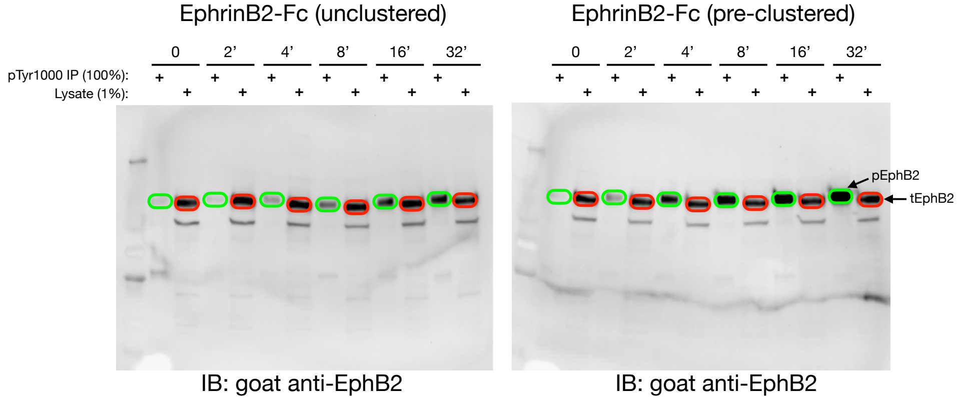

pEphB2 (100% of the IP)

tEphB2 (1% of lysate)

$$\% \text{ pEphB2} = \frac{\#\text{ phospho-photons}}{\#\text{ photons total}} = \frac{\text{pEphB2}}{\text{pEphB2} + (\text{tEphB2} \times 100)}$$

Gels 243A - tEphB2 band analysis

| No. | Label | Type | Volume (Int) | Adj. Vol. (Int) |
|---|---|---|---|---|
| 1 | tEphB2 - No EB2-Fc | Unknown | 230,780,879 | 180,137,258 |
| 2 | tEphB2 - 2' unclustered EB2-Fc | Unknown | 298,765,540 | 248,121,919 |
| 3 | tEphB2 - 4' unclustered EB2-Fc | Unknown | 299,576,807 | 248,933,186 |
| 4 | tEphB2 - 8' unclustered EB2-Fc | Unknown | 268,336,692 | 217,693,071 |
| 5 | tEphB2 - 16' unclustered EB2-Fc | Unknown | 278,743,313 | 228,099,692 |
| 6 | tEphB2 - 32' unclustered EB2-Fc | Unknown | 234,284,482 | 183,640,861 |
| 7 | B1 | Background | 50,643,621 | 0 |

Gels 243A - pEphB2 band analysis

| No. | Label | Type | Volume (Int) | Adj. Vol. (Int) |
|---|---|---|---|---|
| 1 | pEphB2 - No EB2-Fc | Unknown | 51,280,505 | 16,922,475 |
| 2 | pEphB2 - 2' unclustered EB2-Fc | Unknown | 46,147,107 | 11,789,077 |
| 3 | pEphB2 - 4' unclustered EB2-Fc | Unknown | 74,637,246 | 40,279,216 |
| 4 | pEphB2 - 8' unclustered EB2-Fc | Unknown | 104,108,534 | 69,750,504 |
| 5 | pEphB2 - 16' unclustered EB2-Fc | Unknown | 147,706,530 | 113,348,500 |
| 6 | pEphB2 - 32' unclustered EB2-Fc | Unknown | 170,328,088 | 135,970,058 |
| 7 | B1 | Background | 34,358,030 | 0 |

<u>Band analysis: EphrinB2-Fc (unclustered)</u>

| Gels 243A - % pEphB2 | fold increase % pEphB2 | fold increase pEphB2 levels |
|---|---|---|
| 0 = 0.094% | 0 = | 0 = |
| 2 = 0.048% | 2 = 0.5-fold | 2 = 0.7-fold |
| 4 = 0.161% | 4 = 1.7-fold | 4 = 2.4-fold |
| 8 = 0.319% | 8 = 3.4-fold | 8 = 4.1-fold |
| 16 = 0.500% | 16 = 5.3-fold | 16 = 6.7-fold |
| 32 = 0.735% | 32 = 7.8-fold | 32 = 8.0-fold |

Gels 243B - tEphB2 band analysis

| No. | Label | Type | Volume (Int) | Adj. Vol. (Int) |
|---|---|---|---|---|
| 1 | tEphB2 - No EB2-Fc | Unknown | 121,558,628 | 88,667,029 |
| 2 | tEphB2 - 2' preclustered EB2-Fc | Unknown | 110,314,833 | 77,423,234 |
| 3 | tEphB2 - 4' preclustered EB2-Fc | Unknown | 106,546,512 | 73,654,913 |
| 4 | tEphB2 - 8' preclustered EB2-Fc | Unknown | 104,442,701 | 71,551,102 |
| 5 | tEphB2 - 16' preclustered EB2-Fc | Unknown | 118,779,263 | 85,887,664 |
| 6 | tEphB2 - 132' preclustered EB2-Fc | Unknown | 125,633,491 | 92,741,892 |
| 7 | B1 | Background | 32,891,599 | 0 |

Gels 243B - pEphB2 band analysis

| No. | Label | Type | Volume (Int) | Adj. Vol. (Int) |
|---|---|---|---|---|
| 1 | tEphB2 - No EB2-Fc | Unknown | 42,052,145 | 4,548,427 |
| 2 | pEphB2 - 2' preclustered EB2-Fc | Unknown | 52,335,603 | 14,831,885 |
| 3 | pEphB2 - 4' preclustered EB2-Fc | Unknown | 90,503,109 | 52,999,391 |
| 4 | pEphB2 - 8' preclustered EB2-Fc | Unknown | 121,321,733 | 83,818,015 |
| 5 | pEphB2 - 16' preclustered EB2-Fc | Unknown | 151,978,815 | 114,475,097 |
| 6 | pEphB2 - 132' preclustered EB2-Fc | Unknown | 253,045,954 | 215,542,236 |
| 7 | B1 | Background | 37,503,718 | 0 |

<u>Band analysis: EphrinB2-Fc (pre-clustered)</u>

| Gels 243B - % pEphB2 | fold increase % pEphB2 | fold increase pEphB2 levels |
|---|---|---|
| 0 = 0.051% | 0 = | 0 = |
| 2 = 0.191% | 2 = 3.8-fold | 2 = 3.3-fold |
| 4 = 0.714% | 4 = 14.0-fold | 4 = 11.7-fold |
| 8 = 1.158% | 8 = 22.7-fold | 8 = 18.4-fold |
| 16 = 1.315% | 16 = 25.8-fold | 16 = 25.2-fold |
| 32 = 2.271% | 32 = 44.5-fold | 32 = 47.4-fold |

**<u>EphrinB2-Fc (unclustered)</u>**

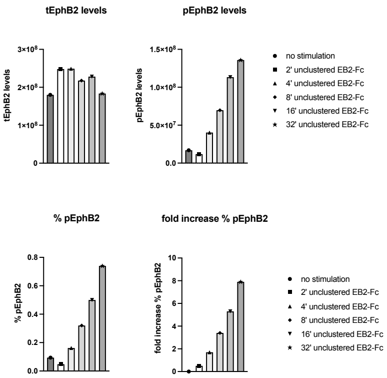

**<u>EphrinB2-Fc (pre-clustered)</u>**

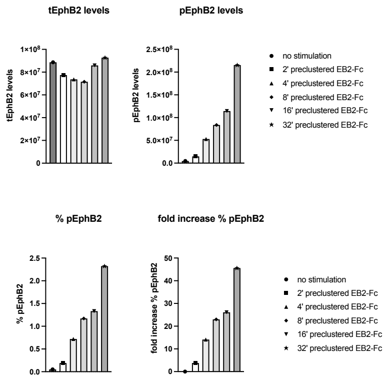

**Figure S6B (related to Figure 2B).** Band quantification details for data shown in **Figure 2B** to precisely quantify levels of activated, tyrosine autophosphorylated pEphB2

for the *in vitro* studies of EphB2 activation in Cos1 cells that endogenously express the receptor and stimulated for 0, 2, 4, 8, 16, and 32 min with 1.5 ug/ml (30 nM) unclustered or pre-clustered EphrinB2-Fc. Densely plated serum-starved Cos1 cells endogenously expressing EphB2 growing in 6-well plates were serum starved and then exposed for 0, 2, 4, 8, 16, or 32 min to 30 nM of EphrinB2-Fc ectodomain that was either unclustered or pre-clustered (with anti-human IgG) to bind the receptor and activate its tyrosine kinase catalytic domain. Resulting whole cell protein lysates were then mixed with pTyr1000 conjugated sepharose beads (Cell Signaling, 14005) to bind all the phosphotyrosine-containing proteins and, after washing beads and eluting proteins, the entire immunoprecipitate (100%) was loaded onto SDS-PAGE gels next to a lane containing 1% of the protein lysate collected before pTyr1000 bead addition. The resulting chemiluminescent images in the goat anti-EphB2 immunoblots shows addition of unclustered and to a greater extent pre-clustered EphrinB2-Fc resulted in a time-dependent progressive increase in the slower migrating pEphB2 protein species in the IP lanes. This increase in tyrosine phosphorylation is consistent with the ability of unclustered EphrinB2-Fc to form Fc-Fc dimers that bind the EphB2 receptor to stimulate dimerization/tetramerization events, and that this ability becomes supercharged by pre-clustering EphrinB2-Fc into a more potent stimulating reagent with anti-human IgG. Quantification of chemiluminescent signals enabled precise determination of the percent level of pEphB2 (% pEphB2) in any one lysate sample by simply dividing the number of phospho-specific pEphB2 photons in the pTyr1000 bead IP lane (which is from 100% of the lysate) by the total number of EphB2 photons which is calculated by multiplying the number of tEphB2 photons in the respective lysate lane by 100 and adding the pEphB2 number, and then plotting the data as % pEphB2 or fold increase in % EphB2. The unstimulated densely plated Cos1 cells exhibited an extremely low % pEphB2 (<0.1% of the tEphB2), that increased 8-fold and 45-fold (to 0.74% and 2.27%) after 32 min stimulation with unclustered or pre-clustered EphrinB2-Fc, respectively. Throughout the stimulations the pre-clustered EphrinB2-Fc reagent exhibited ~5 times greater ability to activate EphB2 and induce its autophosphorylation compared to unclustered EphrinB2-Fc.

EphrinB2 stimulated EphB2 forward signaling in Cos1 - testing MCD tyrosine kinase inhibitor

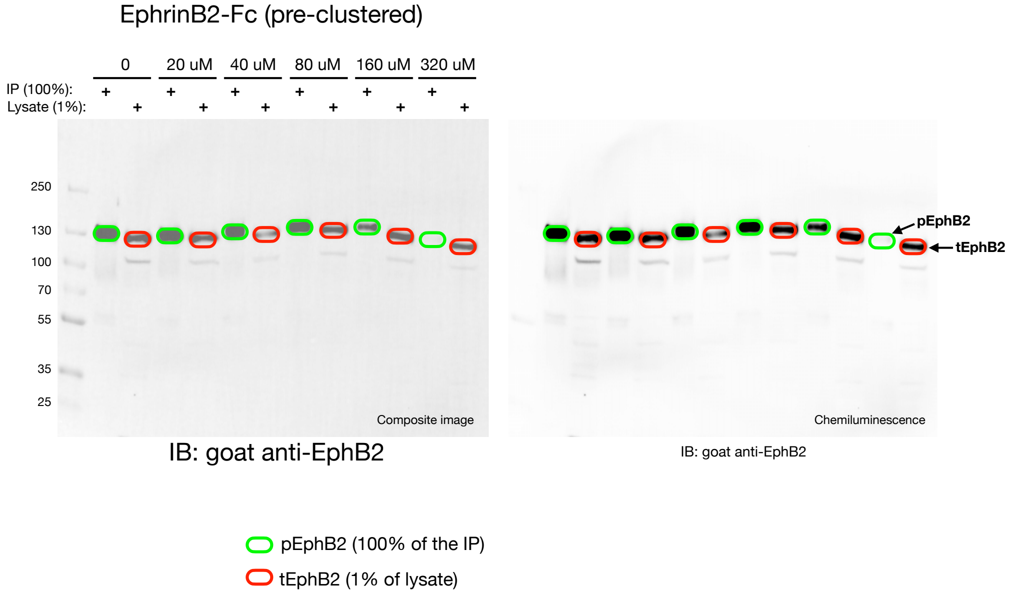

$$\% \text{ pEphB2} = \frac{\text{\# phospho-photons}}{\text{\# photons total}} = \frac{\text{pEphB2}}{\text{pEphB2} + (\text{tEphB2} \times 100)}$$

Gel 250C - 30 nM EB2-Fc for 32 minutes - tEphB2 testing MCD

| No. | Label | Type | Volume (Int) | Adj. Vol. (Int) |
|---|---|---|---|---|
| 1 | tEphB2 - no MCD | Unknown | 141,658,737 | 85,774,154 |
| 2 | tEphB2 - 20 uM MCD | Unknown | 137,235,731 | 81,351,148 |
| 3 | tEphB2 - 40 uM MCD | Unknown | 103,077,339 | 47,192,756 |
| 4 | tEphB2 - 80 uM MCD | Unknown | 127,621,142 | 71,736,559 |
| 5 | tEphB2 - 160 uM MCD | Unknown | 131,296,545 | 75,411,962 |
| 6 | tEphB2 - 320 uM MCD | Unknown | 110,539,622 | 54,655,039 |
| 7 | B1 | Background | 55,884,583 | 0 |

Gel 250C - 30 nM EB2-Fc for 32 minutes - pEphB2 testing MCD

| No. | Label | Type | Volume (Int) | Adj. Vol. (Int) |
|---|---|---|---|---|
| 1 | pEphB2 - no MCD | Unknown | 233,767,570 | 200,939,425 |
| 2 | pEphB2 - 20 uM MCD | Unknown | 199,374,191 | 166,546,046 |
| 3 | pEphB2 - 40 uM MCD | Unknown | 161,416,272 | 128,588,127 |
| 4 | pEphB2 - 80 uM MCD | Unknown | 148,516,104 | 115,687,959 |
| 5 | pEphB2 - 160 uM MCD | Unknown | 97,451,695 | 64,623,550 |
| 6 | pEphB2 - 320 uM MCD | Unknown | 50,242,410 | 17,414,265 |
| 7 | B1 | Background | 32,828,145 | 0 |

Gels 250C - % pEphB2

0 uM MCD = 2.29%
20 uM MCD = 2.00%
40 uM MCD = 2.65%
80 uM MCD = 1.59%
160 uM MCD = 0.85%
320 uM MCD = 0.32%

level of % pEphB2 (compared to no MCD)

0 = 100%
20 = 87%
40 = 116%
80 = 69%
160 = 37%
320 = 14%

reduction in pEphB2 levels (compared to no MCD)

0 = 100%
20 = 83%
40 = 64%
80 = 58%
160 = 32%
320 = 9%

**Figure S6C (related to Figure 2C).** Band quantification details for the data shown in **Figure 2C** that examined the effect of EphB2 tyrosine kinase inhibitor MCD on receptor activation in densely plated serum-starved Cos1 cells stimulated for 32 min with 1.5 ug/ml (30 nM) pre-clustered human EphrinB2-Fc. The tetracyclines minocycline/chlortetracycline/demeclocycline (MCD) when presented as an equal molar mix have previously been shown to act as a pan EphB tyrosine kinase inhibitor by binding within the receptor's intracellular ATP-binding domain to block catalytic activity. When tested at 20-320 μM, MCD resulted in a concentration-dependent reduction in the pEphB2 signal level in the immunoblot, that was further confirmed using the more accurate, internally controlled and calculated % pEphB2 based on the pEphB2/tEphB2 ratio.

### EphrinB2 stimulated EphB2 forward signaling in Cos1 - testing MCD tyrosine kinase inhibitor

#### EphrinB2-Fc (unclustered) - Testing MCD

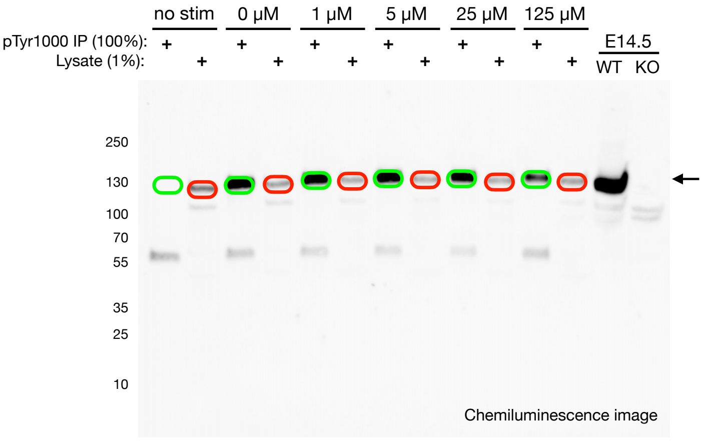

IB: goat anti-EphB2

pEphB2 (100% of the IP)
tEphB2 (1% of lysate)

$$\% \text{ pEphB2} = \frac{\text{\# phospho-photons}}{\text{\# photons total}} = \frac{\text{pEphB2}}{\text{pEphB2} + (\text{tEphB2} \times 100)}$$

12E - MCD

| | Label | Type | Volume (Int) | Adj. Vol. (Int) | % pEphB2 |
|---|---|---|---|---|---|
| 1 | 12E1 - pEphB2 | unstimulated | 9,548,702 | 937,580 | 0.027806 |
| 2 | 12E1 - tEphB2 | unstimulated | 42,320,744 | 33,709,622 | |
| 3 | 12E2 - pEphB2 | no MCD | 171,253,266 | 162,642,144 | 6.138051 |
| 4 | 12E2 - tEphB2 | no MCD | 33,482,061 | 24,870,939 | |
| 5 | 12E3 - pEphB2 | 1 uM MCD | 142,465,561 | 133,854,439 | 6.042352 |
| 6 | 12E3 - tEphB2 | 1 uM MCD | 29,425,283 | 20,814,161 | |
| 7 | 12E4 - pEphB2 | 5 uM MCD | 142,818,809 | 134,207,687 | 5.960429 |
| 8 | 12E4 - tEphB2 | 5 uM MCD | 29,785,491 | 21,174,369 | |
| 9 | 12E5 - pEphB2 | 25 uM MCD | 117,858,562 | 109,247,440 | 4.642468 |
| 10 | 12E5 - tEphB2 | 25 uM MCD | 31,050,839 | 22,439,717 | |
| 11 | 12E6 - pEphB2 | 125 uM MCD | 66,781,278 | 58,170,156 | 2.222925 |
| 12 | 12E6 - tEphB2 | 125 uM MCD | 34,197,716 | 25,586,594 | |

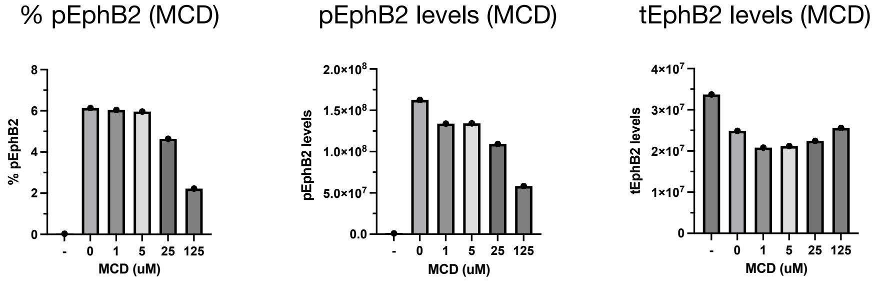

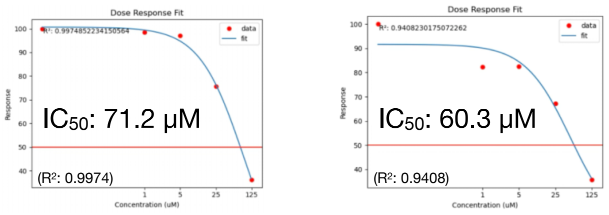

**Figure S6D (related to Figure 2D).** Band quantification details and $IC_{50}$ determination for data shown in **Figure 2D** that tested ability of MCD, A20-I (2xHCl salt), and BQPB4

(3xHCl salt) to disrupt EphrinB2 stimulated EphB2 forward signaling in Cos1 cells. Increasing concentrations (1, 5, 25, 125 μM) of the indicated compounds were added to serum-starved Cos1 cells plated at low density and stimulated with 1.5 ug/ml (30 nM) unclustered EphrinB2-Fc for 32 min resulted in a concentration-dependent reduction in the % pEphB2, with BQPB4 having the strongest effect ($IC_{50}$ = 3.2 μM), followed by A20-I ($IC_{50}$ = 8.5 μM), and then MCD ($IC_{50}$ = 71.2 μM).

### EphrinB2 stimulated EphB2 forward signaling in Cos1 - testing A20-I tetramer inhibitor

EphrinB2-Fc (unclustered) - Testing A20-I

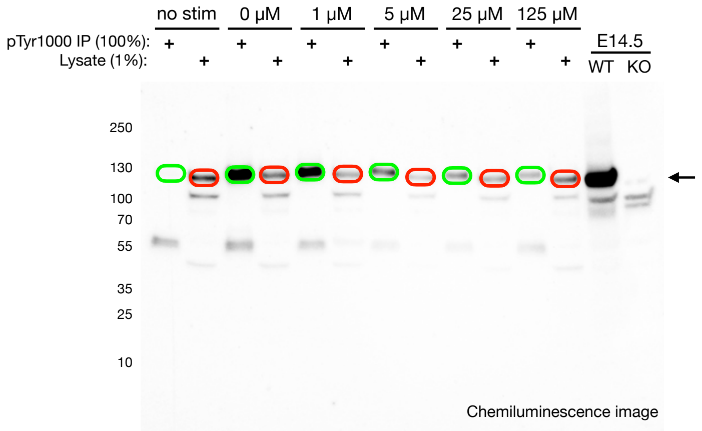

IB: goat anti-EphB2

pEphB2 (100% of the IP)

tEphB2 (1% of lysate)

$$\% \text{ pEphB2} = \frac{\text{\# phospho-photons}}{\text{\# photons total}} = \frac{\text{pEphB2}}{\text{pEphB2} + (\text{tEphB2} \times 100)}$$

| | Label | Type | Volume (Int) | Adj. Vol. (Int) | % pEphB2 |
|---|---|---|---|---|---|
| 1 | 12A1 - pEphB2 | unstimulated | 10,051,575 | 1,071,505 | 0.036054 |
| 2 | 12A1 - tEphB2 | unstimulated | 38,688,813 | 29,708,743 | |
| 3 | 12A2 - pEphB2 | no A20-I | 128,864,560 | 119,884,490 | 5.027043 |
| 4 | 12A2 - tEphB2 | no A20-I | 31,629,137 | 22,649,067 | |
| 5 | 12A3 - pEphB2 | 1 uM A20-I | 75,152,432 | 66,172,362 | 4.740761 |
| 6 | 12A3 - tEphB2 | 1 uM A20-I | 22,276,520 | 13,296,450 | |
| 7 | 12A4 - pEphB2 | 5 uM A20-I | 29,578,956 | 20,598,886 | 3.244750 |
| 8 | 12A4 - tEphB2 | 5 uM A20-I | 15,122,455 | 6,142,385 | |
| 9 | 12A5 - pEphB2 | 25 uM A20-I | 21,704,558 | 12,724,488 | 1.015808 |
| 10 | 12A5 - tEphB2 | 25 uM A20-I | 21,379,291 | 12,399,221 | |
| 11 | 12A6 - pEphB2 | 125 uM A20-I | 17,572,520 | 8,592,450 | 0.435094 |
| 12 | 12A6 - tEphB2 | 125 uM A20-I | 28,642,622 | 19,662,552 | |

% pEphB2 (A20-I)

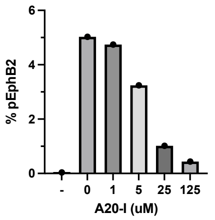

pEphB2 levels (A20-I)

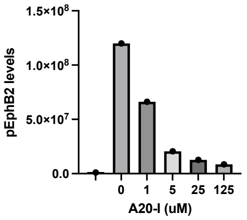

tEphB2 levels (A20-I)

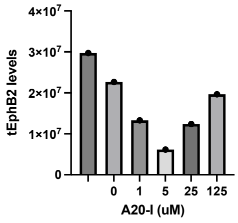

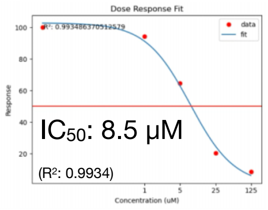

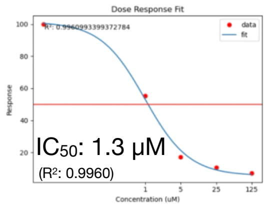

**Figure S6D (continued).**

#### EphrinB2 stimulated EphB2 forward signaling in Cos1 - testing BQPB4 tetramer inhibitor

##### EphrinB2-Fc (unclustered) - Testing BQPB4

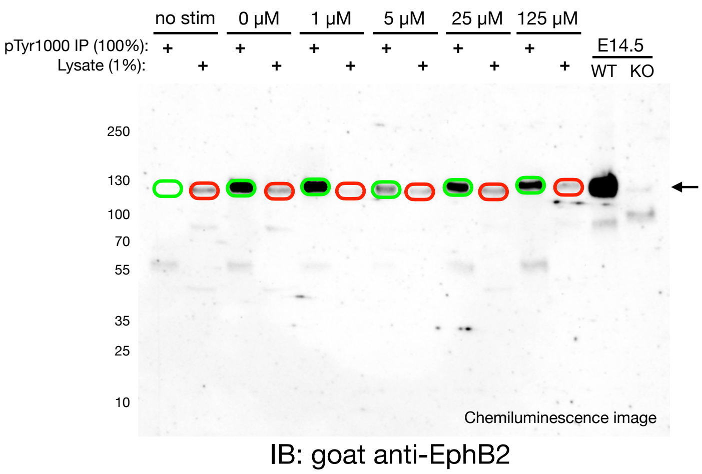

pEphB2 (100% of the IP)

tEphB2 (1% of lysate)

$$\% \text{ pEphB2} = \frac{\#\text{ phospho-photons}}{\#\text{ photons total}} = \frac{\text{pEphB2}}{\text{pEphB2} + (\text{tEphB2} \times 100)}$$

12B - BQPB4

| | Label | Type | Volume (Int) | Adj. Vol. (Int) | % pEphB2 |
|---|---|---|---|---|---|
| 1 | 12B1 - pEphB2 | unstimulated | 395,789 | 21145 | 0.0386 |
| 2 | 12B1 - tEphB2 | unstimulated | 922,833 | 548189 | |
| 3 | 12B2 - pEphB2 | no BQPB4 | 4,470,969 | 4096325 | 7.0477 |
| 4 | 12B2 - tEphB2 | no BQPB4 | 955,873 | 581229 | |
| 5 | 12B3 - pEphB2 | 1 uM BQPB4 | 3,852,338 | 3477694 | 8.604002 |
| 6 | 12B3 - tEphB2 | 1 uM BQPB4 | 544,062 | 369418 | |
| 7 | 12B4 - pEphB2 | 5 uM BQPB4 | 1,092,375 | 717731 | 2.193948 |
| 8 | 12B4 - tEphB2 | 5 uM BQPB4 | 694,608 | 319964 | |
| 9 | 12B5 - pEphB2 | 25 uM BQPB4 | 2,250,509 | 1875865 | 3.197225 |
| 10 | 12B5 - tEphB2 | 25 uM BQPB4 | 942,602 | 567958 | |
| 11 | 12B6 - pEphB2 | 125 uM BQPB4 | 2,046,019 | 1671375 | 3.593051 |
| 12 | 12B6 - tEphB2 | 125 uM BQPB4 | 823,099 | 448455 | |

% pEphB2 (BQPB4)

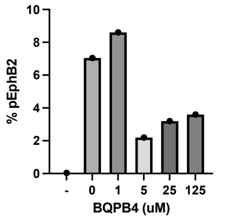

pEphB2 levels (BQPB4)

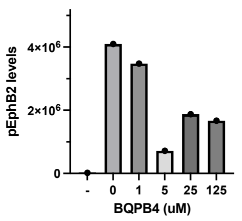

tEphB2 levels (BQPB4)

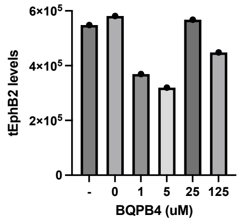

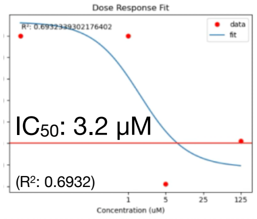

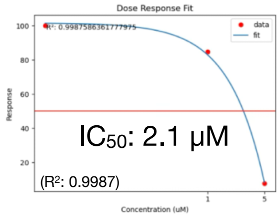

**Figure S6D (continued).**

A20 reduces ability of EphrinB2 to bind, cluster, and activate EphB2 in Cos1 cells

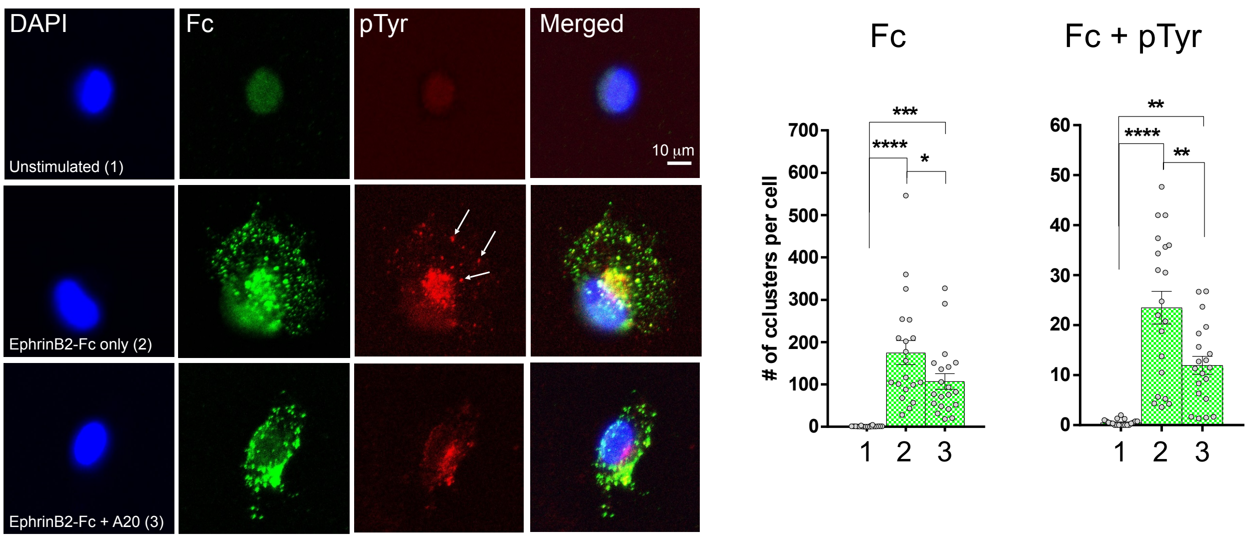

**Figure S6E.** A20 disrupts EphB2 clustering and forward signaling in live cells. Cos1 cells that endogenously express EphB2 growing on coverslips were either unstimulated or stimulated with 4 ug/ml pre-clustered EphrinB2-Fc ectodomain (80 nM) without or with 40 µM A20 (2xHCl salt) for 30 min. Cells were then fixed and subjected to immunofluorescence with anti-IgG antibodies to identify the pre-clustered EphrinB2-Fc ectodomain that bound to EphB2 and formed visible clusters/spots on the cell (green) and with pTyr1000 antibodies to determine if the spots contained detectable phosphotyrosine signal (red, arrows). Any spot sized between 1-5 µm was considered a cluster, and quantification of the data indicated A20 reduced the number of EphrinB2-EphB2 clusters per cell and fewer of the spots contained pTyr signal. Each data point in the scatter plots represent the average number of green and red spots counted from three or more cells in a selected field, and was obtained from 4 different experiments where five high magnification fields per coverslip was selected (n = 3 or more cells/field x 5 fields/experiment x 4 experiments = 60 or more cells per condition).

#### Detecting activated pEphB2 protein in whole embryo lysates

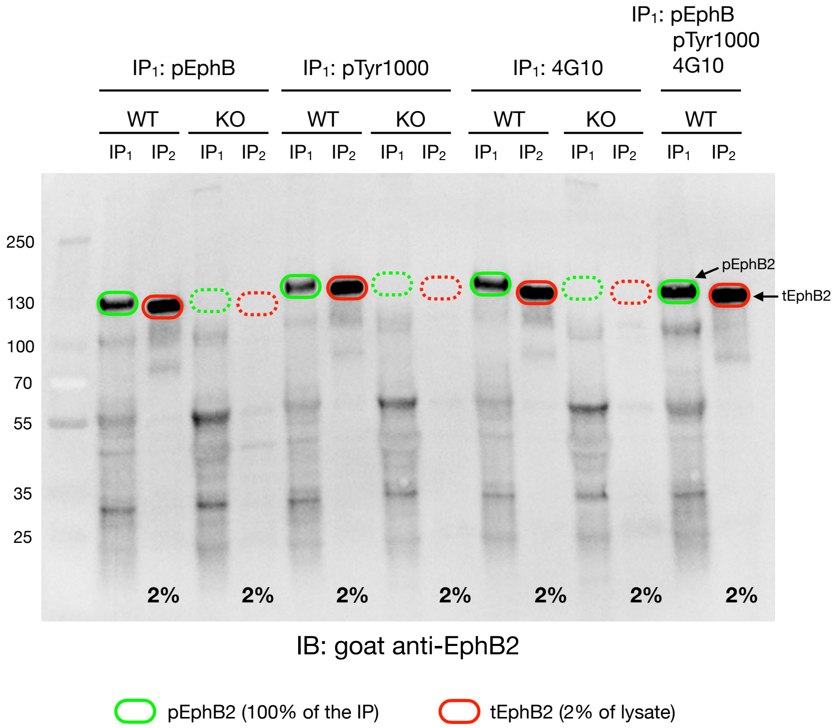

Gel 89 - E14.5 day embryos

| Lane | Label | Volume (Int) | Adj. Vol. (Int) |
|---|---|---|---|
| 1 | pEphB2 | 16,216,156 | 8,734,853 |
| 2 | EphB2 | 31,345,937 | 23,864,634 |
| 3 | pTyr1000 | 14,144,568 | 6,663,265 |
| 4 | EphB2 | 29,536,418 | 22,055,115 |
| 5 | 4G10 | 16,303,509 | 8,822,206 |
| 6 | EphB2 | 28,896,240 | 21,414,937 |
| 7 | pEphB2+pTyr1000+4G10 | 22,009,450 | 14,528,147 |
| 8 | EphB2 | 36,319,864 | 28,838,561 |
| 9 | B1 | 7,481,303 | 0 |

IP$_1$: 100% loaded: # pEphB2-specific photons

IP$_2$: 2% loaded x 50 = # tEphB2 photons

$$\% \text{ pEphB2} = \frac{\text{\# phospho-photons}}{\text{\# photons total}} = \frac{\text{pEphB2}}{\text{pEphB2} + (\text{tEphB2} \times 50)}$$

$$\% \text{ pEphB2} = \frac{\text{\# photons pEphB2}}{\text{\# photons total}} = \frac{8{,}734{,}853}{8{,}734{,}853 + (23{,}864{,}634 \times 50)} = 0.73 \text{ \% pEphB2 (IP with pEphB)}$$

$$\% \text{ pTyr1000} = \frac{\text{\# photons pTyr1000}}{\text{\# photons total}} = \frac{6{,}663{,}265}{6{,}663{,}265 + (22{,}055{,}115 \times 50)} = 0.60 \text{ \% pEphB2 (IP with pTyr1000)}$$

$$\% \text{ 4G10} = \frac{\text{\# photons 4G10}}{\text{\# photons total}} = \frac{8{,}822{,}206}{8{,}822{,}206 + (21{,}414{,}937 \times 50)} = 0.82 \text{ \% pEphB2 (IP with 4G10)}$$

$$\% \text{ pTriple} = \frac{\text{\# photons pTriple}}{\text{\# photons total}} = \frac{14{,}528{,}147}{14{,}528{,}147 + (28{,}838{,}561 \times 50)} = 1.00 \text{ \% pEphB2 (IP with pEphB+pTyr1000+4G10)}$$

**Figure S6F (related to Figure 3A).** Quantification of the band intensities for the experiment shown in **Figure 3A** showed the % of pEphB2 in E14.5 day embryos was not much different between the anti-phospho-EphB (0.73%), pTry1000 (0.60%), or 4G10 (0.82%) antibodies used in the immunoprecipitations, and that combining all three phosphotyrosine antibodies in IP1 only slightly increased the % pEphB2 signal (1.00%). Phosphotyrosine-containing proteins from whole mid-gestation E14.5 day mouse embryo lysates of wild-type (WT) and EphB2 knockout (KO) animals were first immunoprecipitated (IP1) using Protein G Sepharose beads bound to either (1) rabbit anti-pEphB specific antibodies (Abcam, 61791), (2) rabbit pTyr1000 pan phosphotyrosine antibodies (Cell Signaling, #8954), (3) mouse 4G10 Platinum pan phosphotyrosine antibodies (Millipore Sigma, 05-1050), or (4) a combination of all three antibodies. The remaining EphB2 (tEphB2) present in the phosphotyrosine cleared lysates was then immunoprecipitated (IP2) using Protein G Sepharose beads bound to goat anti-EphB2 antibodies (R&D Systems, AF467), and then 100% of the phosphotyrosine IP1 eluted proteins were loaded next to 2% of the EphB2 IP2 eluted proteins, run on a 4-20% gradient SDS-PAGE gel, and immunoblotted with goat anti-EphB2 antibodies. All three phosphotyrosine antibodies were found to immunoprecipitate similar amounts of pEphB2 protein in the WT embryo lysates, and whose migration in the gel was at or just above the 130 KDa marker, slightly higher in the gel than the unphosphorylated EphB2 protein in the IP2 lanes which migrated just below the 130 kDa marker. Note that the pEphB2 band intensities of IP1 lanes are less than the corresponding band intensities of IP2 lanes loaded with 2% of the tEphB2, indicating no more than 2% of the EphB2 protein in the E14.5 day embryo is in the active, tyrosine phosphorylated pEphB2 state. Note that no EphB2 signal was detected in the KO embryos demonstrating specificity of the goat anti-EphB2 antibody (stippled green and red ovals).

### Detecting activated pEphB2 protein in whole brain lysates

$IP_1$: pTyr1000 + 4G10 / $IP_2$: goat anti-EphB2

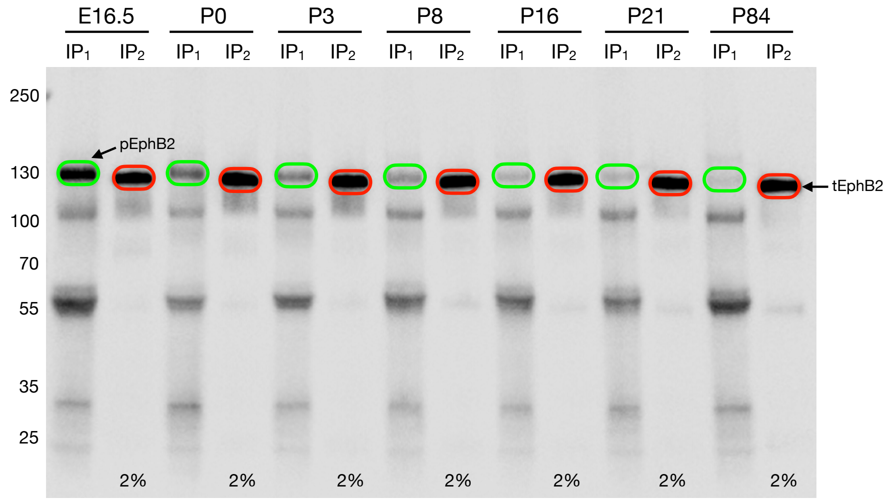

IB: goat anti-EphB2

pEphB2 (100% of the IP)

tEphB2 (2% of lysate)

Chemiluminescence image

Gel 103 - Embryonic and Postnatal Brains

| No. | Label | Type | Volume (Int) | Adj. Vol. (Int) |
|---|---|---|---|---|
| 1 | E16.5 | Unknown | 17,670,080 | 13,430,250 |
| 2 | pE16.5 | Unknown | 13,692,971 | 9,453,141 |
| 3 | P0 | Unknown | 22,547,623 | 18,307,793 |
| 4 | pP0 | Unknown | 8,658,026 | 4,418,196 |
| 5 | P3 | Unknown | 22,464,928 | 18,225,098 |
| 6 | pP3 | Unknown | 7,278,138 | 3,038,308 |
| 7 | P8 | Unknown | 20,077,322 | 15,837,492 |
| 8 | pP8 | Unknown | 6,600,915 | 2,361,085 |
| 9 | P16 | Unknown | 18,532,327 | 14,292,497 |
| 10 | pP16 | Unknown | 5,222,315 | 982,485 |
| 11 | P21 | Unknown | 19,583,282 | 15,343,452 |
| 12 | pP21 | Unknown | 5,125,717 | 885,887 |
| 13 | P84 | Unknown | 19,970,382 | 15,730,552 |
| 14 | pP84 | Unknown | 4,932,303 | 692,473 |
| 15 | B1 | Background | 4,239,830 | 0 |

| pEphB2 protein | tEphB2 protein |
|---|---|
| E16.5 = 9,453,141 | E16.5 = 13,430,250 |
| P0 = 4,418,196 | P0 = 18,307,793 |
| P3 = 3,038,308 | P3 = 18,225,098 |
| P8 = 2,361,085 | P8 = 15,837,492 |
| P16 = 982,485 | P16 = 14,292,497 |
| P21 = 885,887 | P21 = 15,343,452 |
| P84 = 692,473 | P84 = 15,730,552 |

$$\% \; p = \frac{\# \text{ phospho-photons}}{\# \text{ photons total}} = \frac{\text{pEphB2}}{\text{pEphB2} + (\text{tEphB2} \times 50)}$$

= 1.39% pEphB2 in E16.5 brain

= 0.48% pEphB2 in P0 brain

= 0.33% pEphB2 in P3 brain

= 0.30% pEphB2 in P8 brain

= 0.14% pEphB2 in P16 brain

= 0.12% pEphB2 in P21 brain

= 0.09% pEphB2 in P84 brain

**Figure S6G (related to Figure 3C).** Quantification of the band intensities for the experiment shown in **Figure 3C** indicated pEphB2 represented 1.39% of the total EphB2 protein in the E16.5 day embryonic brain, only 0.48% in the brain at birth (P0), and essentially undetectable in the 84 day adult brain (0.09%).

Administration of A20 or EDTA tetramer inhibitors reduced activated pEphB2 protein in the intestines

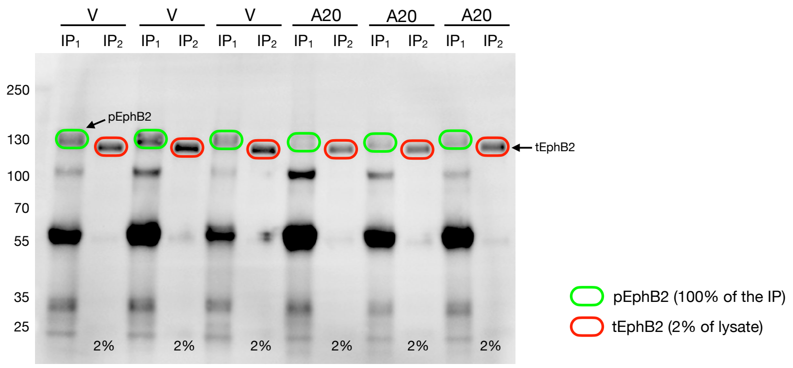

IB: goat anti-EphB2

Intestines: Vehicle vs A20

Gel 107C - A20 Treated P8 Intestines

| No. | Label | Type | Volume (Int) | Adj. Vol. (Int) |
|---|---|---|---|---|
| 1 | V-1 | Unknown | 61,321,051 | 28,039,215 |
| 2 | pV-1 | Unknown | 65,476,040 | 32,194,204 |
| 3 | V-2 | Unknown | 69,435,598 | 36,153,762 |
| 4 | pV-2 | Unknown | 73,518,419 | 40,236,583 |
| 5 | V-3 | Unknown | 62,712,950 | 29,431,114 |
| 6 | pV-3 | Unknown | 54,656,724 | 21,374,888 |
| 7 | A20-1 | Unknown | 47,023,746 | 13,741,910 |
| 8 | pA20-1 | Unknown | 40,293,473 | 7,011,637 |
| 9 | A20-2 | Unknown | 48,096,278 | 14,814,442 |
| 10 | pA20-2 | Unknown | 41,195,951 | 7,914,115 |
| 11 | A20-3 | Unknown | 55,823,129 | 22,541,293 |
| 12 | pA20-3 | Unknown | 46,339,399 | 13,057,563 |
| 13 | B1 | Background | 33,281,836 | 0 |

| tEphB2 | pEphB2 |
|---|---|
| V = 28,039,215 | V = 32,194,204 |
| V = 36,153,762 | V = 40,236,583 |
| V = 29,431,114 | V = 21,374,888 |
| A20 = 13,741,910 | A20 = 7,011,637 |
| A20 = 14,814,442 | A20 = 7,914,115 |
| A20 = 22,541,293 | A20 = 13,057,563 |

$$\% \text{ pEphB2} = \frac{\text{pEphB2}}{\text{pEphB2} + (\text{tEphB2} \times 50)}$$

= 2.25% pEphB2 in V (1)
= 2.18% pEphB2 in V (2)
= 1.43% pEphB2 in V (3)
= 1.01% pEphB2 in A20 (1)
= 1.06% pEphB2 in A20 (2)
= 1.15% pEphB2 in A20 (3)

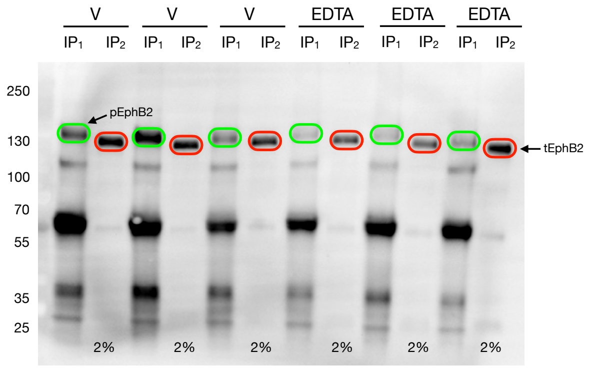

IB: goat anti-EphB2

Intestines: Vehicle vs EDTA

Gel 107D - EDTA Treated P8 Intestines

| No. | Label | Type | Volume (Int) | Adj. Vol. (Int) |
|---|---|---|---|---|
| 1 | V-1 | Unknown | 143,333,531 | 80,941,003 |
| 2 | pV-1 | Unknown | 132,570,131 | 70,177,603 |
| 3 | V-2 | Unknown | 128,941,445 | 66,548,917 |
| 4 | pV-2 | Unknown | 169,916,829 | 107,524,301 |
| 5 | V-3 | Unknown | 117,514,650 | 55,122,122 |
| 6 | pV-3 | Unknown | 108,461,124 | 46,068,596 |
| 7 | EDTA-1 | Unknown | 100,107,691 | 37,715,163 |
| 8 | pEDTA-1 | Unknown | 79,424,831 | 17,032,303 |
| 9 | EDTA-2 | Unknown | 99,095,669 | 36,703,141 |
| 10 | pEDTA-2 | Unknown | 82,954,007 | 20,561,479 |
| 11 | EDTA-3 | Unknown | 133,341,156 | 70,948,628 |
| 12 | pEDTA-3 | Unknown | 99,067,348 | 36,674,820 |
| 13 | B1 | Background | 62,392,528 | 0 |

| tEphB2 | pEphB2 |
|---|---|
| V = 80,941,003 | V = 70,177,603 |
| V = 66,548,917 | V = 107,524,301 |
| V = 55,122,122 | V = 46,068,596 |
| EDTA = 37,715,163 | EDTA = 17,032,303 |
| EDTA = 36,703,141 | EDTA = 20,561,479 |
| EDTA = 70,948,628 | EDTA = 36,674,820 |

$$\% \text{ pEphB2} = \frac{\text{pEphB2}}{\text{pEphB2} + (\text{tEphB2} \times 50)}$$

= 1.71% pEphB2 in V (1)
= 3.13% pEphB2 in V (2)
= 1.64% pEphB2 in V (3)
= 0.90% pEphB2 in EDTA (1)
= 1.11% pEphB2 in EDTA (2)
= 1.02% pEphB2 in EDTA (3)

**Figure S6H (related to Figure 3D).** Quantification of the band intensities for the experiment shown in **Figure 3D** that studied the effect of A20 and EDTA on activation state of EphB2 in P8 intestines. Intestine lysates were prepared from P8 pups after four intraperitoneal dosings with ~2 ul of a 50 mg/ml solution of either A20 (2xHCl salt), EDTA (ethylenediaminetetraacetic acid tetrasodium salt), or vehicle only (100% PBS) over a two day period starting at P6. Phosphotyrosine-containing proteins were then immunoprecipitated (IP1) using Protein G Sepharose beads bound to pTyr1000+4G10 antibodies, and then the phosphotyrosine cleared lysates were immunoprecipitated (IP2) using Protein G Sepharose beads bound to goat anti-EphB2 antibodies. 100% of the phosphotyrosine IP1 eluted proteins were loaded next to 2% of the anti-EphB2 IP2 eluted proteins, run on a gradient SDS-PAGE gel, and immunoblotted with goat anti-EphB2 antibodies.

Administration of A20 or EDTA tetramer inhibitors reduced activated pEphB2 protein in the kidneys

$IP_1$: pTyr1000-conjugated beads / $IP_2$: goat anti-EphB2

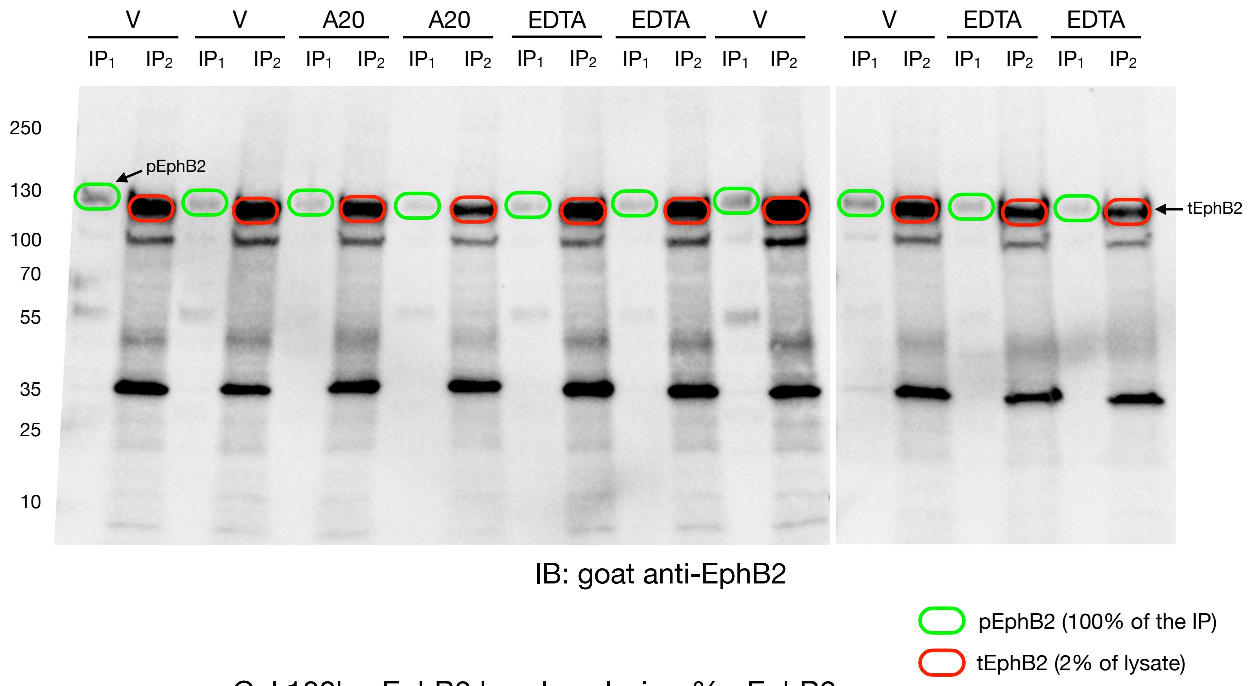

Gel 180b - EphB2 band analysis - % pEphB2

| No. | Label | Type | Volume (Int) | Adj. Vol. (Int) | % pEphB2 |
|---|---|---|---|---|---|
| 1 | pEphB2 - V #1 | Unknown | 4,484,068 | 1,963,139 | 0.163192 |
| 2 | tEphB2 - V #1 | Unknown | 15,587,005 | 12,029,608 | |
| 3 | pEphB2 - V #2 | Unknown | 3,948,193 | 1,427,264 | 0.118714 |
| 4 | tEphB2 - V #2 | Unknown | 15,580,131 | 12,022,734 | |
| 5 | pEphB2 - V #3 | Unknown | 4,204,938 | 1,684,009 | 0.092708 |
| 6 | tEphB2 - V #3 | Unknown | 21,722,118 | 18,164,721 | |
| 7 | pEphB2 - V #4 | Unknown | 4,220,865 | 1,699,936 | 0.154916 |
| 8 | tEphB2 - V #4 | Unknown | 14,530,702 | 10,973,305 | |
| 9 | pEphB2 - A20 #1 | Unknown | 3,451,335 | 930,406 | 0.089462 |
| 10 | tEphB2 - A20 #1 | Unknown | 13,957,438 | 10,400,041 | |
| 11 | pEphB2 - A20 #2 | Unknown | 2,944,602 | 423,673 | 0.066928 |
| 12 | tEphB2 - A20 #2 | Unknown | 9,887,637 | 6,330,240 | |
| 13 | pEphB2 - EDTA #1 | Unknown | 3,200,083 | 679,154 | 0.056066 |
| 14 | tEphB2 - EDTA #1 | Unknown | 15,670,830 | 12,113,433 | |
| 15 | pEphB2 - EDTA #2 | Unknown | 3,369,766 | 848,837 | 0.076356 |
| 16 | tEphB2 - EDTA #2 | Unknown | 14,674,188 | 11,116,791 | |
| 17 | pEphB2 - EDTA #3 | Unknown | 3,429,127 | 908,198 | 0.104517 |
| 18 | tEphB2 - EDTA #3 | Unknown | 12,246,852 | 8,689,455 | |
| 19 | pEphB2 - EDTA #4 | Unknown | 3,105,526 | 584,597 | 0.089145 |
| 20 | tEphB2 - EDTA #4 | Unknown | 10,115,184 | 6,557,787 | |
| 21 | B1 | Background | 2,520,929 | 0 | |
| 22 | B1 | Background | 3,557,397 | 0 | |

**Figure S6I (related to Figure 3E).** Quantification of the band intensities for the experiment shown in **Figure 3E** that studied the effect of A20 and EDTA on activation state of EphB2 in P8 kidneys. Kidney lysates were prepared from P8 pups after four intraperitoneal dosings with ~2 ul of a 50 mg/ml solution of either A20 (2xHCl salt), EDTA, or vehicle only (100% PBS) over a two day period starting at P6. Phosphotyrosine-containing proteins were then immunoprecipitated (IP1) using pTyr1000-conjugated beads, and then the phosphotyrosine cleared lysates were immunoprecipitated (IP2) using Protein G Sepharose beads bound to goat anti-EphB2 antibodies. 100% of the phosphotyrosine IP1 eluted proteins were loaded next to 1% of the anti-EphB2 IP2 eluted proteins, run on a gradient SDS-PAGE gel, and immunoblotted with goat anti-EphB2 antibodies.

#### Administration of A20 or EDTA reduced activated pEphB2 protein *in vivo* in the intestines

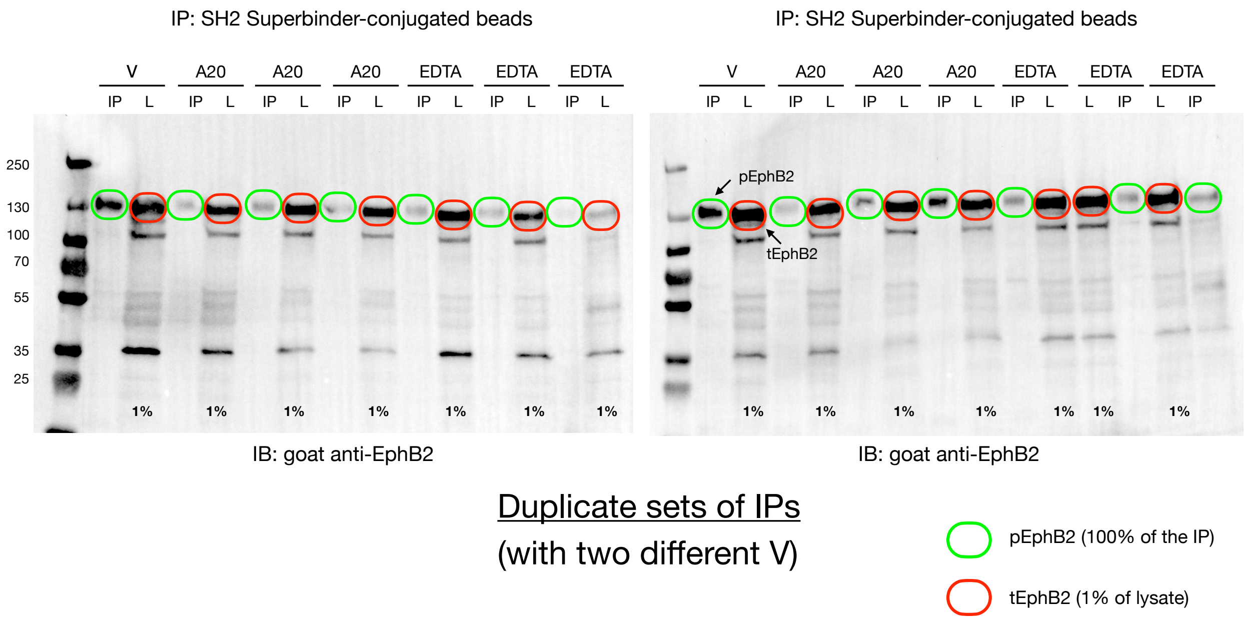

##### Gel 176a - A20/EDTA Treated P8 Intestines

| No. | Label | Type | Volume (Int) | Adj. Vol. (Int) | % pEphB2 |
|---|---|---|---|---|---|
| 1 | pVehicle #2 | Unknown | 25,554,704 | 16,903,088 | 0.637006 |
| 2 | tVehicle #2 | Unknown | 43,434,792 | 26,535,218 | |
| 3 | pA20 #1 | Unknown | 12,448,100 | 3,796,484 | 0.187454 |
| 4 | tA20 #1 | Unknown | 37,152,476 | 20,252,902 | |
| 5 | pA20 #2 | Unknown | 13,902,652 | 5,251,036 | 0.221457 |
| 6 | tA20 #2 | Unknown | 40,610,845 | 23,711,271 | |
| 7 | pA20 #3 | Unknown | 13,136,871 | 4,485,255 | 0.231365 |
| 8 | tA20 #3 | Unknown | 36,285,636 | 19,386,062 | |
| 9 | pEDTA #1 | Unknown | 12,801,608 | 4,149,992 | 0.164810 |
| 10 | tEDTA #1 | Unknown | 42,079,986 | 25,180,412 | |
| 11 | pEDTA #2 | Unknown | 12,589,937 | 3,938,321 | 0.275216 |
| 12 | tEDTA #2 | Unknown | 31,209,514 | 14,309,940 | |
| 13 | pEDTA #3 | Unknown | 9,483,255 | 831,639 | 0.184468 |
| 14 | tEDTA #3 | Unknown | 21,407,888 | 4,508,314 | |
| 15 | B1 | Backgrounc | 8,651,616 | 0 | |
| 16 | B1 | Backgrounc | 16,899,574 | 0 | |

##### Gel 176b - A20/EDTA Treated P8 Intestines

| No. | Label | Type | Volume (Int) | Adj. Vol. (Int) | % pEphB2 |
|---|---|---|---|---|---|
| 1 | pVehicle 3 | Unknown | 28,945,748 | 20,434,765 | 0.422586 |
| 2 | tVehicle 3 | Unknown | 64,225,997 | 48,356,485 | |
| 3 | pA20 #1 | Unknown | 12,721,540 | 4,210,557 | 0.136205 |
| 4 | tA20 #1 | Unknown | 46,782,969 | 30,913,457 | |
| 5 | pA20 #2 | Unknown | 15,990,371 | 7,479,388 | 0.223883 |
| 6 | tA20 #2 | Unknown | 49,277,077 | 33,407,565 | |
| 7 | pA20 #3 | Unknown | 21,159,544 | 12,648,561 | 0.436793 |
| 8 | tA20 #3 | Unknown | 44,827,335 | 28,957,823 | |
| 9 | pEDTA #1 | Unknown | 16,203,034 | 7,692,051 | 0.201398 |
| 10 | tEDTA #1 | Unknown | 54,062,845 | 38,193,333 | |
| 11 | pEDTA #2 | Unknown | 15,680,469 | 7,169,486 | 0.179920 |
| 12 | tEDTA #2 | Unknown | 55,716,103 | 39,846,591 | |
| 13 | pEDTA #3 | Unknown | 15,680,469 | 8,328,897 | 0.236169 |
| 14 | tEDTA #3 | Unknown | 51,136,207 | 35,266,695 | |
| 15 | B1 | Background | 8,510,983 | 0 | |
| 16 | B1 | Background | 15,869,512 | 0 | |

**Figure S6J (related to Figure 3F).** Quantification of the band intensities for the experiment shown in **Figure 3F** that studied the effect of A20 (2xHCl salt) and EDTA on activation state of EphB2 in P8 intestines in duplicate IPs done at same time using lysates from three A20-treated and three EDTA-treated animals (the two V samples are from different animals).

#### Methods to quantify levels of pEphrinB2 protein *in vivo* from E13.5 day mouse embryos

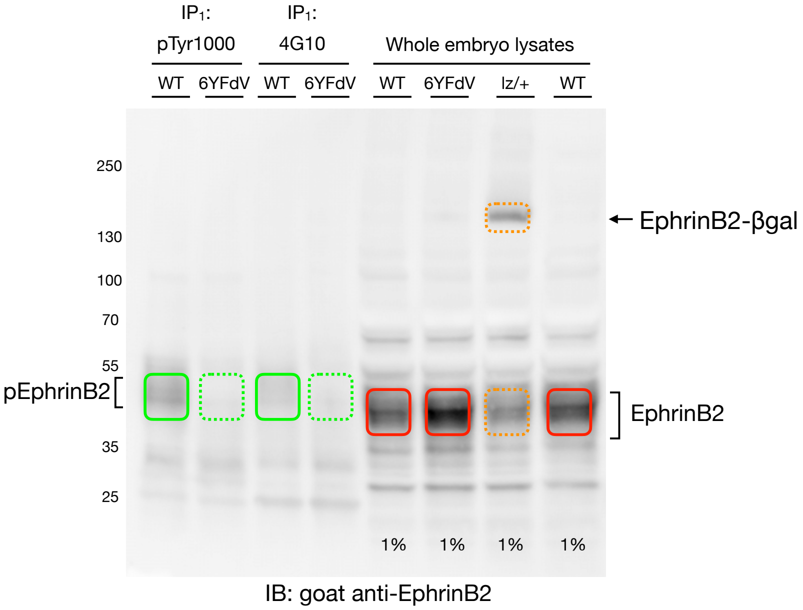

pEphrinB2 in WT (100% of the IP)

pEphrinB2 in 6YFdV (100% of the IP)

tEphrinB2 in WT and 6YFdV/6YFdV (1% of the lysate)

tEphrinB2 in lacZ/+ both WT and EphrinB2-βgal proteins (1% of the lysate)

##### EphrinB2 band volume analysis - E13.5 embryos

Gel 119 - E13.5 embryos

| No. | Label | Type | Volume (Int) | Adj. Vol. (Int) |
|---|---|---|---|---|
| 1 | WT+pTyr1000 | Unknown | 159,552,132 | 94,797,847 |
| 2 | 6YFdV-pTyr1000 | Unknown | 74,571,199 | 9,816,914 |
| 3 | WT+4G10 | Unknown | 112,594,539 | 47,840,254 |
| 4 | 6YFdV+4G10 | Unknown | 79,045,521 | 14,291,236 |
| 5 | EB2 +/+ (WT) | Unknown | 281,846,206 | 217,091,921 |
| 6 | EB2 6YFdV/6YFdV | Unknown | 360,727,475 | 295,973,190 |
| 7 | EB2 lz/+ (WT band) | Unknown | 215,803,415 | 151,049,130 |
| 8 | EB2 +/+ | Unknown | 308,911,450 | 244,157,165 |
| 9 | EB2 lz/+ (lz band) | Unknown | 119,347,508 | 54,593,223 |
| 10 | B1 | Background | 64,754,285 | 0 |

<u>IP: pTyr 1000</u>
pWT = 94,797,847
p6YFdV = 9,816,914

<u>IP: 4G10</u>
pWT = 47,840,254
p6YFdV = 14,291,236

<u>IB: goat EphrinB2</u>
WT = 217,091,921
6YFdV = 295,973,190
WT band (EB2-lz/+) = 151,049,130
LZ band (EB2-lz/+) = 54,593,223
WT = 244,157,165

$$\% \text{ pEphrinB2} = \frac{\#\text{ phospho-photons}}{\#\text{ photons total}} = \frac{94{,}797{,}847}{94{,}797{,}847 + (217{,}091{,}921 \times 100)} = 0.435\% \text{ pEphrinB2 in WT (with pTyr1000)}$$

>13-fold increase in signal

$$\frac{9{,}816{,}914}{9{,}816{,}914 + (295{,}973{,}190 \times 100)} = 0.033\% \text{ pEphrinB2 in 6YFdV (with pTyr1000)}$$

$$\frac{47{,}840{,}254}{47{,}840{,}254 + (217{,}091{,}921 \times 100)} = 0.220\% \text{ pEphrinB2 in WT (with 4G10)}$$

4.6-fold increase in signal

$$\frac{14{,}291{,}236}{9{,}816{,}914 + (295{,}973{,}190 \times 100)} = 0.048\% \text{ pEphrinB2 in 6YFdV (with 4G10)}$$

**Figure S7A (related to Figure 4A).** Quantification of the band intensities for the experiment shown in **Figure 4A** that investigated the phosphotyrosine state of EphrinB2 protein in E13.5 day mouse embryos as a readout for levels of reverse signaling *in vivo*. Phosphotyrosine-containing proteins from lysates of *EphrinB2* +/+ (WT) and *EphrinB2* (*Efnb2*) intracellular point mutant (6YFdV) embryos collected at E13.5 day were immunoprecipitated (IP) using Protein G Sepharose beads bound to either pTyr1000 or 4G10 antibodies, then beads were washed and 100% of the eluted proteins were loaded and run on a gradient SDS-PAGE gel along with 1% samples of the lysates collected prior to IP, and then immunoblotted with goat anti-EphrinB2 antibodies (R&D Systems, AF496). The whole embryo lysate lanes show WT EphrinB2 protein migrates in SDS-PAGE in a broad band of 35-50 kDa (red boxes), as does the homozygous EphrinB2-6YFdV/6YFdV point mutant in which the six intracellular tyrosine residues of EphrinB2 known to become phosphorylated during reverse signaling are mutated to phenylalanine, while the *EphrinB2* lacZ/+ (lz/+) heterozygote embryo lysate included in the immunoblot expresses a larger sized, intracellular truncated EphrinB2-βgal fusion protein of ~150 kDa and correspondingly less WT protein and serves as an excellent control for the specificity of the goat anti-EphrinB2 antibody used in the immunoblot (stippled orange boxes). The data indicate pTyr1000 was able to immunoprecipitate a small amount of phosphorylated pEphrinB2 protein in the WT embryo lysate (green box), whose migration was slightly retarded in the SDS-PAGE gel compared to the unphosphorylated WT protein in the lysate lanes (red box), while no pEphrinB2 protein was detected in the 6YFdV mutant as expected (stippled green boxes). Compared to pTyr1000 beads, the 4G10 antibody precipitated much less pEphrinB2 protein in the WT lysates. Quantification of the pTyr1000 data indicate 0.435% of the EphrinB2 protein is in the active tyrosine phosphorylated pEphrinB2 state in E13.5 day embryo lysates.

Methods to quantify levels of pEphrinB2 protein *in vivo* from E14.5 day mouse embryos

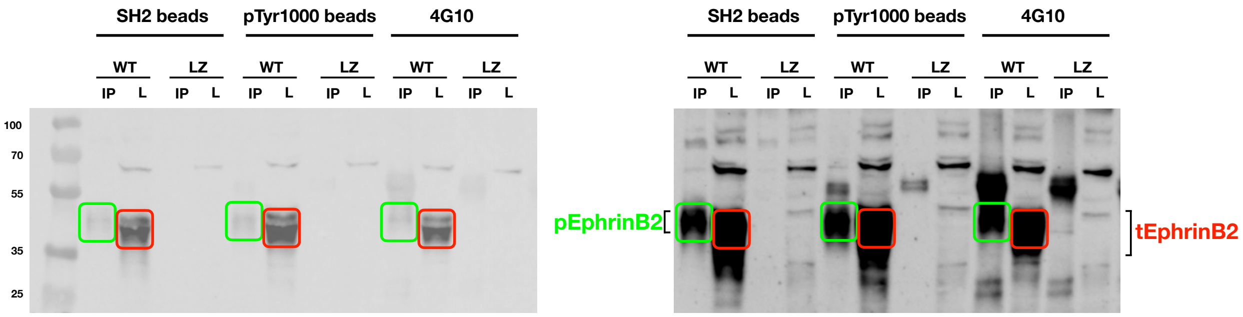

**Immunoblot: goat anti-EphrinB2 (multichannel)**

**Immunoblot: goat anti-EphrinB2 (chemiluminescence) (burn)**

**pEphrinB2 (100% of the IP)**

**tEphrinB2 (1% of the lysate)**

215a Band Analysis - pEphrinB2 in E14.5 embryo lysates

| Label | Type | Volume (Int) | Adj. Vol. (Int) | % pEphrinB2 |
|---|---|---|---|---|
| pEB2 - WT (225a) SH2 bead IP | Unknown | 3,651,163 | 2,996,229 | 0.140991 |
| tEB2 - WT (225a) Lysate | Unknown | 21,906,210 | 21,251,276 | |
| pEB2 - WT (225b) pTyr1000 bead IP | Unknown | 4,160,769 | 3,505,835 | 0.127245 |
| tEB2 - WT (225c) Lysates | Unknown | 28,206,683 | 27,551,749 | |
| pEB2 - WT (225c) 4G10 IP | Unknown | 3,868,210 | 3,213,276 | 0.166135 |
| tEB2 - WT (225c) Lysate | Unknown | 19,996,333 | 19,341,399 | |

**Figure S7B (related to Figure 4B).** Quantification of the band intensities for the experiment shown in **Figure 4B** that investigated the phosphotyrosine state of EphrinB2 protein in E14.5 day mouse embryos as a readout for levels of reverse signaling in vivo. Phosphotyrosine-containing proteins from lysates of *EphrinB2* +/+ (WT) and *EphrinB2* lacZ/lacZ homozygous mutant (LZ) embryos collected at E14.5 days were immunoprecipitated (IP) using SH2 domain superbinder-conjugated beads, pTyr1000-conjugated beads, or 4G10 antibodies bound to Protein G Sepharose beads. After washing beads, 100% of the eluted proteins were loaded and run on a gradient SDS-PAGE gel along with 1% samples of the lysates collected prior to IP, and then immunoblotted with goat anti-EphrinB2 antibodies (R&D Systems, AF496). Shown are two exposures of the anti-EphrinB2 immunoblot. The whole embryo lysate lanes (L) show WT EphrinB2 protein migrates in SDS-PAGE as a broad band of 35-50 kDa (red boxes), while the LZ homozygous mutant embryo lysate does not express a WT sized EphrinB2 protein and serves as an excellent control for the specificity of the goat anti-EphrinB2 antibody used in the immunoblot. The SH2 Domain Superbinder-conjugated beads and the pTyr1000-conjugated beads were both able to cleanly immunoprecipitate phosphorylated pEphrinB2 protein in the WT embryo lysate (green boxes), whose migration was slightly retarded in the SDS-PAGE gel compared to the unphosphorylated WT protein in the lysate lanes, while no pEphrinB2 protein was detected in the LZ mutant lysates as expected. While the 4G10 IP also showed pEphrinB2 protein (green box), its signal was masked by the presence of a strong background band corresponding to the mouse 4G10 antibody used in the IP that also is eluted and loaded onto the gel. The antibody background was greatly diminished in the pTyr1000-conjugated bead IPs, and absent in the SH2-bead IPs as no antibody fragments are present in this reagent. Quantification of signals in the SH2 Domain Superbinder IP indicated the active pEphrinB2 represented 0.14% of the total protein, and was similar to the 0.13% determined from the  pTyr1000 bead IP.

Tetramer inhibitor BQPB4 reduced EphrinB2 reverse signaling in stellate cells stimulated with EphB2

#### EphB2-Fc (unclustered) - Testing BQPB4

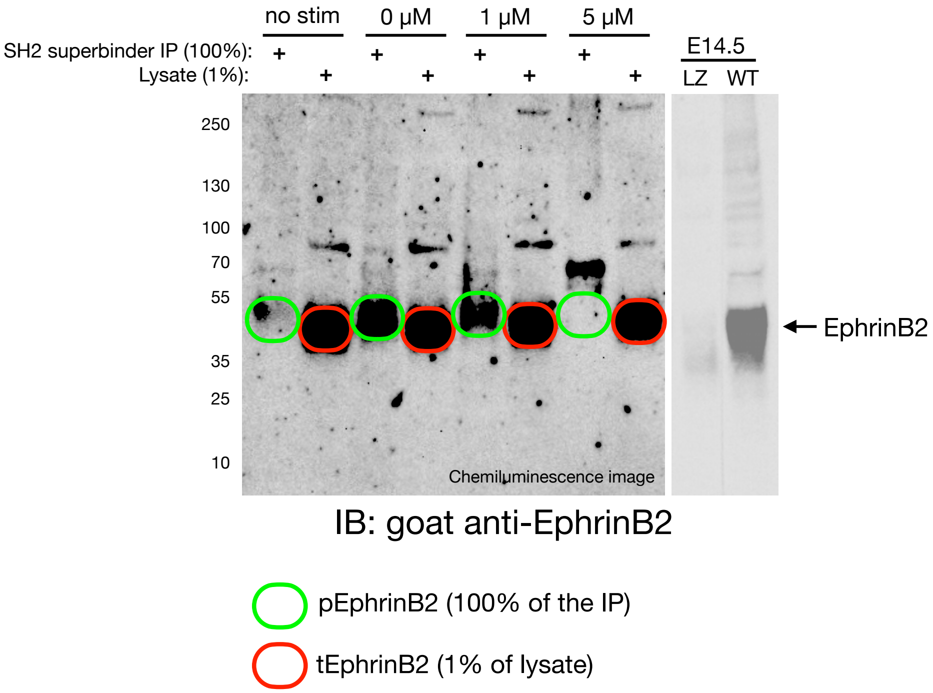

$$\% \text{ pEphrinB2} = \frac{\#\text{ phospho-photons}}{\#\text{ photons total}} = \frac{\text{pEphrinB2}}{\text{pEphrinB2} + (\text{tEphrinB2} \times 100)}$$

Cell Exp 16B - Testing BQPB4 (scan 16B_5)

| | Label | Type | Volume (Int) | Adj. Vol. (Int) | % pEphrinB2 |
|---|---|---|---|---|---|
| 1 | pEphrinB2: unstimulated | Unknown | 2,770,553 | 895394 | 0.084552 |
| 2 | tEphrinB2: unstimulated | Unknown | 12,456,065 | 10580906 | |
| 3 | pEphrinB2: + 30 nM unclustered EphB2-Fc | Unknown | 4,158,244 | 2283085 | 0.408999 |
| 4 | tEphrinB2: + 30 nM unclustered EphB2-Fc | Unknown | 7,434,457 | 5559298 | |
| 5 | pEphrinB2: + 30 nM unclustered EphB2-Fc + 1 uM BQPB4 | Unknown | 3,405,549 | 1530390 | 0.282720 |
| 6 | tEphrinB2: + 30 nM unclustered EphB2-Fc + 1 uM BQPB4 | Unknown | 7,272,945 | 5397786 | |
| 7 | pEphrinB2: + 30 nM unclustered EphB2-Fc + 5 uM BQPB4 | Unknown | 1,903,450 | 28291 | 0.005465 |
| 8 | tEphrinB2: + 30 nM unclustered EphB2-Fc + 5 uM BQPB4 | Unknown | 7,051,733 | 5176574 | |

**Figure S7C (related to Figure 4C).** Quantification of the band intensities for the experiment shown in **Figure 4C** that investigated the phosphotyrosine state of EphrinB2 protein as a readout for levels of reverse signaling *in vitro* in pancreatic stellate cells after stimulation with EphB2-Fc and testing BQPB4.

Tetramer inhibitor EDTA reduced EphrinB2 reverse signaling in stellate cells stimulated with EphB2

#### EphB2-Fc (unclustered) - Testing EDTA

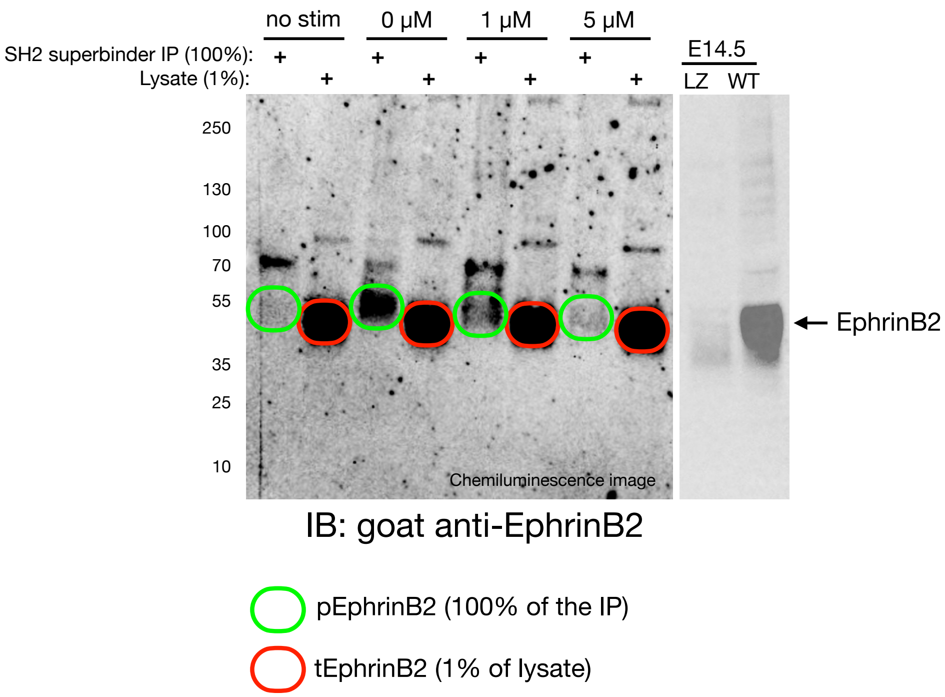

$$\% \text{ pEphrinB2} = \frac{\#\text{ phospho-photons}}{\#\text{ photons total}} = \frac{\text{pEphrinB2}}{\text{pEphrinB2} + (\text{tEphrinB2} \times 100)}$$

Cell Exp 16B - Testing EDTA (scan 16D_5)

| | Label | Type | Volume (Int) | Adj. Vol. (Int) | % pEphrinB2 |
|---|---|---|---|---|---|
| 1 | pEphrinB2: unstimulated | Unknown | 2,419,575 | 62809 | 0.021995 |
| 2 | tEphrinB2: unstimulated | Unknown | 5,211,685 | 2854919 | |
| 3 | pEphrinB2: + 30 nM unclustered EphB2-Fc | Unknown | 2,831,004 | 474238 | 0.179161 |
| 4 | tEphrinB2: + 30 nM unclustered EphB2-Fc | Unknown | 4,999,017 | 2642251 | |
| 5 | pEphrinB2: + 30 nM unclustered EphB2-Fc + 1 uM EDTA | Unknown | 2,664,893 | 308127 | 0.092045 |
| 6 | tEphrinB2: + 30 nM unclustered EphB2-Fc + 1 uM EDTA | Unknown | 5,701,253 | 3344487 | |
| 7 | pEphrinB2: + 30 nM unclustered EphB2-Fc + 5 uM EDTA | Unknown | 2,252,432 | 130973 | 0.032104 |
| 8 | tEphrinB2: + 30 nM unclustered EphB2-Fc + 5 uM EDTA | Unknown | 6,199,844 | 4078385 | |

**Figure S7D (related to Figure 4D).** Quantification of the band intensities for the experiment shown in **Figure 4D** that investigated the phosphotyrosine state of EphrinB2 protein as a readout for levels of reverse signaling *in vitro* in pancreatic stellate cells after stimulation with EphB2-Fc and testing EDTA.

#### Dosing animals with tetramer inhibitor EDTA reduced pEphrinB2 levels in P8 intestines

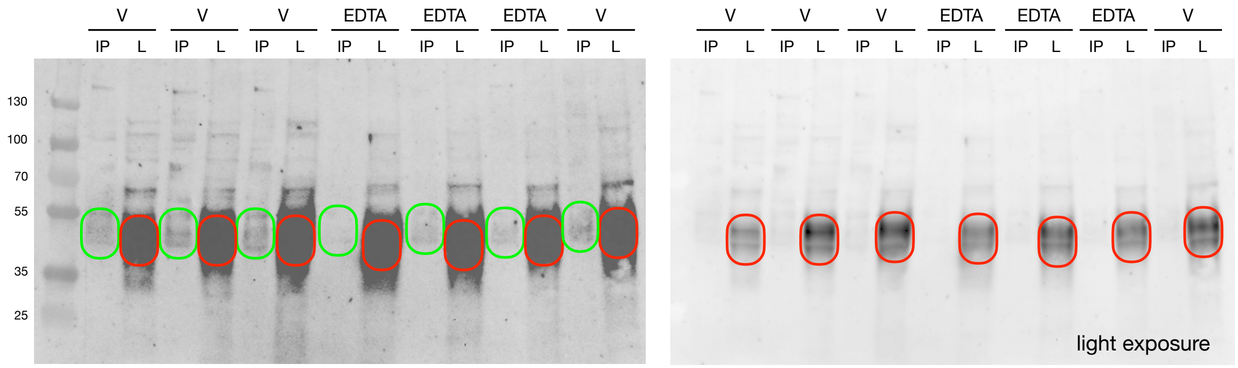

IB: goat anti-EphrinB2 (multichannel)

IB: goat anti-EphrinB2 (chemiluminescence)

$$\% \text{ pEphrinB2} = \frac{\text{\# phospho-photons}}{\text{\# photons total}} = \frac{\text{pEphrinB2}}{\text{pEphrinB2} + (\text{tEphrinB2} \times 100)}$$

pEphrinB2 (100% of the IP)

tEphrinB2 (1% of lysate)

Gel 217b - % pEphrinB2 band analysis

| Label | Type | Volume (Int) | Adj. Vol. (Int) | % pEphrinB2 |
|---|---|---|---|---|
| pEB2 - V1 | Unknown | 12,024,370 | 3,576,039 | 0.146801 |
| tEB2 - V1 | Unknown | 34,824,397 | 24,324,077 | |
| pEB2 - V4 | Unknown | 12,676,687 | 4,228,356 | 0.100014 |
| tEB2 - V4 | Unknown | 52,735,567 | 42,235,247 | |
| pEB2 - V7 | Unknown | 12,573,604 | 4,125,273 | 0.092215 |
| tEB2 - V7 | Unknown | 55,194,378 | 44,694,058 | |
| pEB2 - E1 | Unknown | 9,666,739 | 1,218,408 | 0.038103 |
| tEB2 - E1 | Unknown | 42,464,924 | 31,964,604 | |
| pEB2 - E3 | Unknown | 10,783,272 | 2,334,941 | 0.058570 |
| tEB2 - E3 | Unknown | 50,342,542 | 39,842,222 | |
| pEB2 - E4 | Unknown | 10,295,661 | 1,847,330 | 0.058920 |
| tEB2 - E4 | Unknown | 41,834,937 | 31,334,617 | |
| pEB2 - V8 | Unknown | 12,357,496 | 3,909,165 | 0.095628 |
| tEB2 - V8 | Unknown | 51,340,228 | 40,839,908 | |

**Figure S7E (related to Figure 4E).** Quantification of the band intensities for the experiment shown in **Figure 4E** that investigated the phosphotyrosine state of EphrinB2 protein as a readout for levels of reverse signaling *in vivo* in P8 intestines following dosing with EDTA.

#### Statistical results of the thermal and mechanical pain testing in *EphB1* mutant mice

##### Figure 5B

Thermal Pain Ratio = $\frac{\text{uninjured right paw response time (sec)}}{\text{injured left paw response time (sec)}}$

two-way ANOVA (**** *p* <0.0001)

| Source of Variation | % of total variation | P value | Summary |
|---|---|---|---|
| Interaction | 12.48 | <0.0001 | **** |
| Row Factor | 28.85 | <0.0001 | **** |
| Column Factor | 25.76 | <0.0001 | **** |

Tukey's post hoc (by genotype):
WT: Baseline vs CFA +1 (**** *p* <0.0001)
WT: Baseline vs CFA +2 (**** *p* <0.0001)
WT: Baseline vs CFA +4 (**** *p* <0.0001)
WT: Baseline vs CFA +6 (*** *p* = 0.0003)
EphB1-lacZ: Baseline vs CFA +1 (**** *p* <0.0001)
EphB1-lacZ: Baseline vs CFA +2 (* *p* = 0.0318)

Tukey's post hoc (by time):
CFA +1: WT vs KO (**** *p* <0.0001)
CFA +1: KO vs lacZ (**** *p* <0.0001)
CFA +2: WT vs KO (*** *p* = 0.0001)
CFA +4: WT vs KO (**** *p* <0.0001)
CFA +6: WT vs KO (** *p* = 0.0022)
CFA +6: WT vs lacZ (* *p* = 0.0392)
CFA +8: WT vs KO (*** *p* = 0.0005)
CFA +8: WT vs lacZ (* *p* = 0.0426)

##### Figure 5C

Mechanical Pain Ratio = $\frac{\text{uninjured right paw (adjusted filament)}}{\text{injured left paw (adjusted filament)}}$

two-way ANOVA

| Source of Variation | % of total variation | P value | Summary |
|---|---|---|---|
| Interaction | 25.22 | 0.0079 | ** |
| Row Factor | 20.11 | 0.0030 | ** |
| Column Factor | 9.439 | 0.0286 | * |

Tukey's post hoc (by genotype):
WT: Baseline -2 vs CFA +3 (* *p* = 0.0181)
WT: Baseline -2 vs CFA +8 (** *p* = 0.0012)
WT: Baseline -1 vs CFA +3 (* *p* = 0.0284)
WT: Baseline -1 vs CFA +8 (** *p* = 0.0019)
EphB1-lacZ: Baseline vs CFA +3 (** *p* = 0.0059)

Tukey's post hoc (by time):
CFA +3: WT vs KO (* *p* = 0.0284)
CFA +3: KO vs lacZ (** *p* = 0.0058)
CFA +8: WT vs KO (** *p* = 0.0028)
CFA +8: WT vs lacZ (* *p* = 0.0343)

**Figure S8A (related to Figures 5B and 5C).** Two-way ANOVA and Tukey's post hoc results of the thermal and mechanical pain ratios obtained from a group of female *EphB1* +/+ (WT), *EphB1*-KO, and *EphB1*-lacZ mutant mice at 3 months age in a mixed 129/B6/CD1 background.

Statistical results of the thermal pain testing after dosing mice with A20 compounds

$$\text{Thermal Pain Ratio} = \frac{\text{uninjured right paw response time (sec)}}{\text{injured left paw response time (sec)}}$$

#### Figure 5D

CFA +3: one-way ANOVA (* $p$ = 0.0183)
Dunnett's post hoc:
Vehicle vs A20 (* $p$ = 0.0274)
Vehicle vs A20+MCD (* $p$ = 0.0121)

#### Figure 5E

CFA +3: one-way ANOVA (*** $p$ = 0.0003)
Dunnett's post hoc:
Vehicle vs A20 (*** $p$ = 0.0009)
Vehicle vs QPB4-Bn (*** $p$ = 0.0005)

**Figure S8B (related to Figures 5D and 5E).** One-way ANOVA and Dunnett's post hoc results of thermal pain ratios obtained from mice that were assessed on +3 days post-CFA. Thermal pain ratios in **Figure 5D** were from a group of 20 week old CFA-injured WT female mice in a mixed 129/CD1/B6 background that were IP injected twice each day with either vehicle only (6% DMSO/94% sunflower seed oil), A20 free base (20 mg/kg in 6% DMSO/94% sunflower seed oil), MCD kinase inhibitor (10 mg/kg in PBS), or dual injected with both A20+MCD, starting two days prior to CFA injury and continuing for two additional days. Thermal pain ratios in **Figure 5E** were from a group of 8 week old CFA-injured WT male and female mice in a mixed 129/CD1/B6 background that were IP injected with either vehicle only (PBS), A20.2xHCl salt (20 mg/kg in PBS), or QPB4-Bn.2xHCl salt (20 mg/kg in PBS) starting 15 min after CFA was injected into the left hind paw to initiate inflammatory pain, followed by 6 more injections spaced every 12 hours before pain testing.

#### Statistical results of the thermal pain testing after PO and IP dosing mice with A20

Left Injured Paw | Right Uninjured Paw

Thermal Pain
(grouped by treatment)

Thermal Pain
(grouped by time)

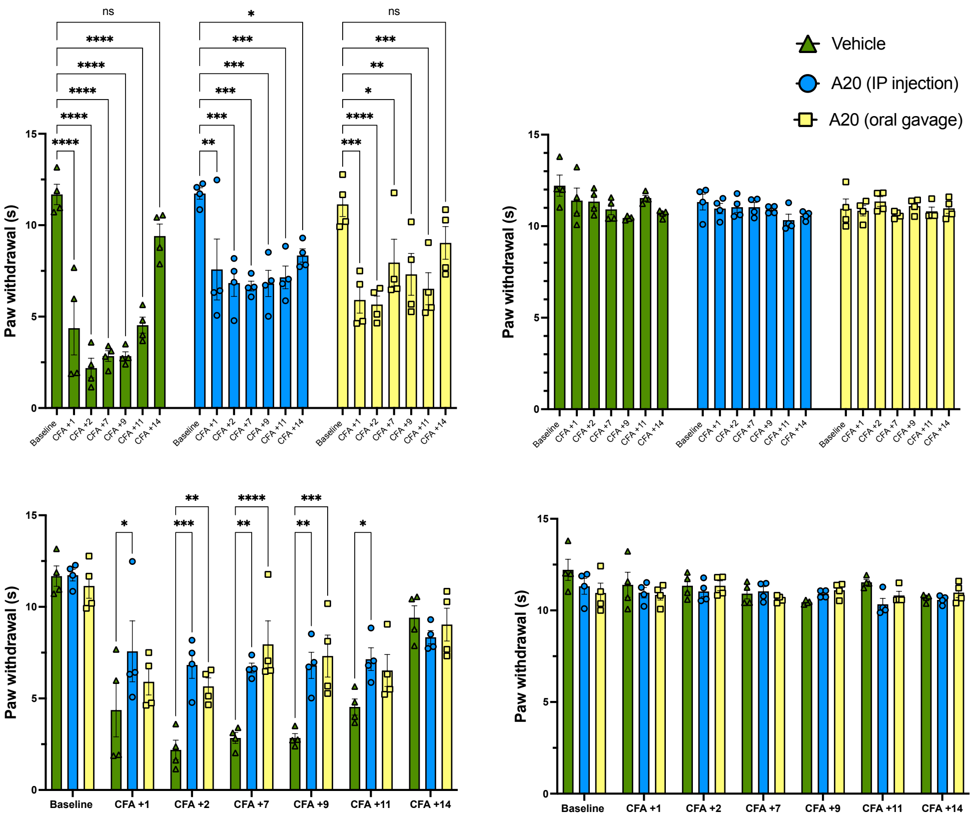

<u>two-way ANOVA (Thermal Pain Measurements)</u>

| Source of Variation | % of total variation | P value | Summary |
|---|---|---|---|
| Interaction | 11.34 | 0.0046 | ** |
| Row Factor | 53.33 | <0.0001 | **** |
| Column Factor | 13.68 | <0.0001 | **** |

<u>Tukey's post hoc (by treatment group):</u>

Vehicle
- Baseline vs. CFA +1 (**** *p* <0.0001)
- Baseline vs. CFA +2 (**** *p* <0.0001)
- Baseline vs. CFA +7 (**** *p* <0.0001)
- Baseline vs. CFA +9 (**** *p* <0.0001)
- Baseline vs. CFA +11 (**** *p* <0.0001)
- Baseline vs. CFA +14 (**** *p* <0.0001)

A20 (IP Injection)
- Baseline vs. CFA +1 (** *p* = 0.003)
- Baseline vs. CFA +2 (*** *p* = 0.0004)
- Baseline vs. CFA +7 (*** *p* = 0.0002)
- Baseline vs. CFA +9 (*** *p* = 0.0003)
- Baseline vs. CFA +11 (*** *p* = 0.0009)
- Baseline vs. CFA +14 (* *p* = 0.0214)

A20 (oral gavage)
- Baseline vs. CFA +1 (*** *p* = 0.0001)
- Baseline vs. CFA +2 (**** *p* <0.0001)
- Baseline vs. CFA +7 (* *p* = 0.0342)
- Baseline vs. CFA +9 (** *p* = 0.0072)
- Baseline vs. CFA +11 (*** *p* = 0.0008)

<u>Tukey's post hoc (by treatment time):</u>

CFA +1
- Vehicle vs. A20 (IP) (* *p* = 0.0126)

CFA +2
- Vehicle vs. A20 (IP) (*** *p* = 0.0003)
- Vehicle vs. A20 (PO) (** *p* = 0.0066)

CFA +7
- Vehicle vs. A20 (IP) (** *p* = 0.0027)
- Vehicle vs. A20 (PO) (**** *p* <0.0001)

CFA +9
- Vehicle vs. A20 (IP) (** *p* = 0.003)
- Vehicle vs. A20 (PO) (*** *p* = 0.0004)

CFA +11
- Vehicle vs. A20 (IP) (* *p* = 0.0474)

**Figure S8C (related to Figure 5F).** Thermal pain measurement data and two-way

ANOVA with Tukey's post hoc analysis of the mice shown in **Figure 5F/G** of a group of 11 week old WT male 129x1/SvJ mice mice that were assessed on multiple days post-CFA and dosed PO with vehicle only (3% DMSO/97% PBS) or IP or PO dosed with A20.2xHCl salt (20 mg/kg in 3% DMSO/97% PBS) starting 15 min after CFA was injected into the left hind paw to initiate inflammatory pain, followed by additional dosings every 12 or 24 hours using the schedule outlined in **Figure 5A**.

#### Statistical results of the thermal pain testing after PO and IP dosing mice with A20

$$\text{Thermal Pain Ratio} = \frac{\text{uninjured right paw response time (sec)}}{\text{injured left paw response time (sec)}}$$

**two-way ANOVA (Thermal Pain Ratios)**

| Source of Variation | % of total variation | P value | Summary |
|---|---|---|---|
| Interaction | 19.62 | 0.0006 | *** |
| Row Factor | 22.6 | <0.0001 | **** |
| Column Factor | 27.83 | <0.0001 | **** |

Dunnett's post hoc (by treatment group):

Vehicle
- Baseline vs. CFA +1 (*** $p = 0.0006$)
- Baseline vs. CFA +2 (**** $p < 0.0001$)
- Baseline vs. CFA +7 (*** $p = 0.0003$)
- Baseline vs. CFA +9 (** $p = 0.0011$)

Dunnett's post hoc (by treatment time):

CFA +1
- Vehicle vs. A20 (IP) (** $p = 0.0033$)
- Vehicle vs. A20 (PO) (* $p = 0.0108$)

CFA +2
- Vehicle vs. A20 (IP) (**** $p < 0.0001$)
- Vehicle vs. A20 (PO) (**** $p < 0.0001$)

CFA +7
- Vehicle vs. A20 (IP) (** $p = 0.002$)
- Vehicle vs. A20 (PO) (*** $p = 0.0006$)

CFA +9
- Vehicle vs. A20 (IP) (** $p = 0.0063$)
- Vehicle vs. A20 (PO) (** $p = 0.0055$)

**Figure S8D (related to Figure 5F).** Two-way ANOVA with Dunnett's post hoc analysis of the thermal pain ratios shown in **Figure 5F** of a group of 11 week old WT male 129x1/SvJ mice mice that were assessed on multiple days post-CFA and dosed PO with vehicle only (3% DMSO/97% PBS) or IP or PO dosed with A20.2xHCl salt (20 mg/kg in 3% DMSO/97% PBS) starting 15 min after CFA was injected into the left hind paw to initiate inflammatory pain, followed by additional dosings every 12 or 24 hours using the schedule outlined in **Figure 5A**.

Statistical results of the mechanical pain testing after PO and IP dosing mice with A20

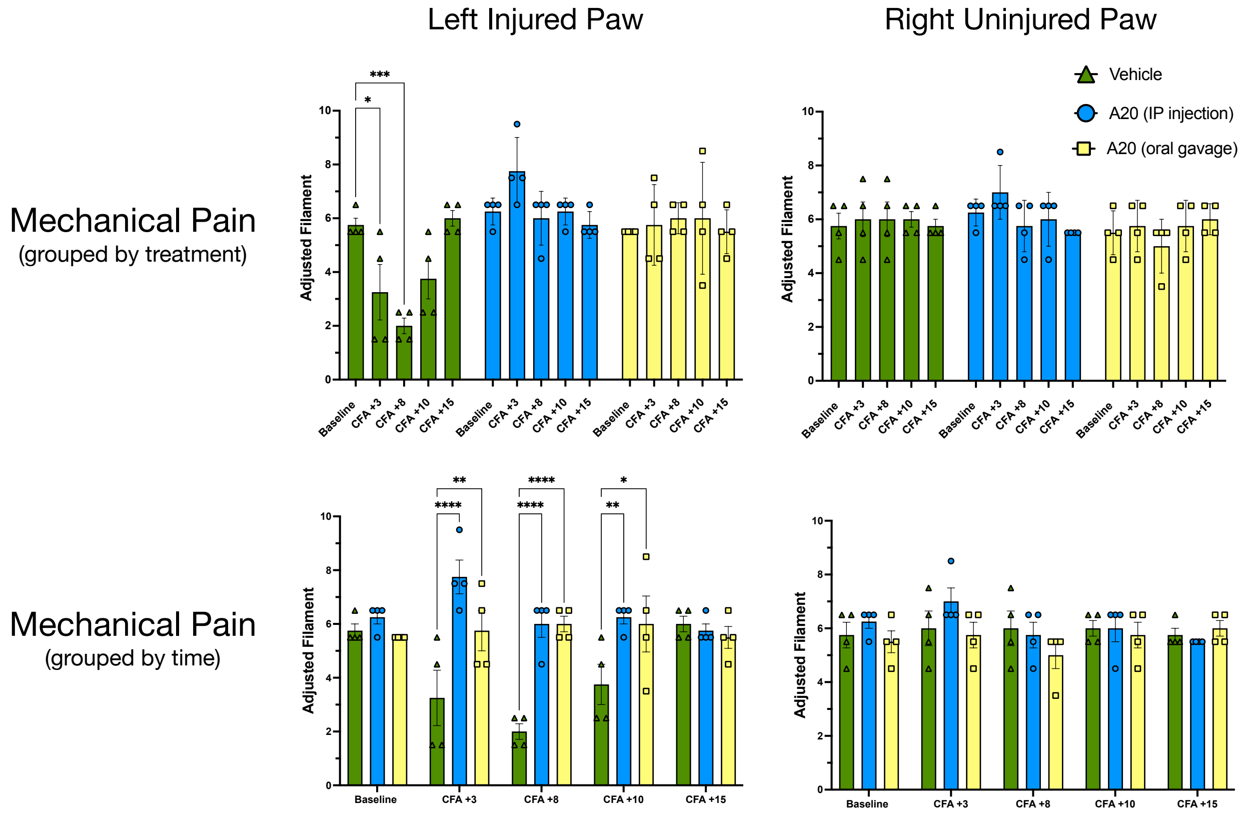

#### two-way ANOVA (Mechanical Pain Measurements)

| Source of Variation | % of total variation | P value | Summary |
|---|---|---|---|
| Interaction | 28.08 | 0.0003 | *** |
| Row Factor | 6.376 | 0.0887 | ns |
| Column Factor | 32.36 | <0.0001 | **** |

Tukey's post hoc (by treatment group):

Vehicle
- Baseline vs. CFA +3 (* $p = 0.0203$)
- Baseline vs. CFA +8 (*** $p = 0.0002$)

Tukey's post hoc (by treatment time):

CFA +3
- Vehicle vs. A20 (IP) (**** $p < 0.0001$)
- Vehicle vs. A20 (PO) (** $p = 0.007$)

CFA +8
- Vehicle vs. A20 (IP) (**** $p < 0.0001$)
- Vehicle vs. A20 (PO) (**** $p < 0.0001$)

**Figure S8E (related to Figure 5G).** Mechanical pain measurement data and two-way ANOVA with Tukey's post hoc analysis of the mice shown in **Figure 5F/G** of a group of 11 week old WT male 129x1/SvJ mice mice that were assessed on multiple days post-CFA and dosed PO with vehicle only (3% DMSO/97% PBS) or IP or PO dosed with

A20.2xHCl salt (20 mg/kg in 3% DMSO/97% PBS) starting 15 min after CFA was injected into the left hind paw to initiate inflammatory pain, followed by additional dosings every 12 or 24 hours using the schedule outlined in **Figure 5A**.

Statistical results of the mechanical pain testing after PO and IP dosing mice with A20

$$\text{Mechanical Pain Ratio} = \frac{\text{uninjured right paw (adjusted filament)}}{\text{injured left paw (adjusted filament)}}$$

two-way ANOVA (Mechanical Pain Ratios)

| Source of Variation | % of total variation | P value | Summary |
|---|---|---|---|
| Interaction | 25.62 | 0.0017 | ** |
| Row Factor | 10.09 | 0.0276 | * |
| Column Factor | 26.63 | <0.0001 | **** |

Dunnett's post hoc (by treatment group):

Vehicle
- Baseline vs. CFA +3 (** *p* = 0.0018)
- Baseline vs. CFA +8 (**** *p* <0.0001)

Dunnett's post hoc (by treatment time):

CFA +3
- Vehicle vs. A20 (IP) (*** *p* = 0.0006)
- Vehicle vs. A20 (PO) (** *p* = 0.0014)

CFA +8
- Vehicle vs. A20 (IP) (**** *p* <0.0001)
- Vehicle vs. A20 (PO) (**** *p* <0.0001)

**Figure S8F (related to Figure 5G).** Two-way ANOVA with Dunnett's post hoc analysis of the mechanical pain ratios shown in **Figure 5G** of a group of 11 week old WT male 129x1/SvJ mice mice that were assessed on multiple days post-CFA and dosed PO with vehicle only (3% DMSO/97% PBS) or IP or PO dosed with A20.2xHCl salt (20 mg/kg in 3% DMSO/97% PBS) starting 15 min after CFA was injected into the left hind paw to initiate inflammatory pain, followed by additional dosings every 12 or 24 hours using the schedule outlined in **Figure 5A**.

### Figure 5H

two-way ANOVA (Thermal Pain Measurements)

| Source of Variation | % of total variation | P value | Summary |
|---|---|---|---|
| Interaction | 1.134 | 0.3616 | ns |
| Row Factor | 486.91 | <0.0001 | **** |
| Column Factor | 0.5733 | 0.2032 | ns |

Uncorrected Fisher's LSD post hoc (by treatment time):
2-4 hr EphB1 KO vs 4-6 hr EphB1 HET (* $p$ = 0.0143)
6-8 hr EphB1 WT vs 4-6 hr EphB1 KO (* $p$ = 0.0325)
6-8 hr EphB1 WT vs 4-6 hr EphB1 HET (** $p$ = 0.0062)

$$\text{Thermal Pain Ratio} = \frac{\text{baseline/uninjured left paw response time (sec)}}{\text{injured left paw response time (sec)}}$$

two-way ANOVA (Thermal Pain Ratio)

| Source of Variation | % of total variation | P value | Summary |
|---|---|---|---|
| Interaction | 9.596 | 0.0662 | ns |
| Row Factor | 21.91 | 0.0008 | *** |
| Column Factor | 14.93 | 0.0173 | * |

Tukey's post hoc (by treatment time):
6-8 hr EphB1 WT vs 6-8 hr EphB1 KO (** $p$ = 0.0088)
6-8 hr EphB1 WT vs 6-8 hr EphB1 HET (** $p$ = 0.0045)

two-way ANOVA (Mechanical Pain Measurements)

| Source of Variation | % of total variation | P value | Summary |
|---|---|---|---|
| Interaction | 9.900 | 0.0115 | * |
| Row Factor | 47.22 | <0.0001 | **** |
| Column Factor | 11.43 | 0.0063 | ** |

Tukey's post hoc (by treatment time):
4-6 hr EphB1 WT vs 4-6 hr EphB1 KO (** $p$ = 0.0015)
4-6 hr EphB1 WT vs 4-6 hr EphB1 HET (* $p$ = 0.0149)

$$\text{Mechanical Pain Ratio} = \frac{\text{baseline/uninjured left paw (adjusted filament)}}{\text{injured left paw (adjusted filament)}}$$

one-way ANOVA (Mechanical Pain Ratio) (* $p$ = 0.0314)
Dunnett's post hoc:
4-6 hr WT vs 4-6 hr EphB1 KO (* $p$ = 0.0162)

**Figure S8G (related to Figure 5H).** One-way and two-way ANOVA of the thermal and mechanical pain testing data shown in **Figure 5H** using Zymosan model that assessed *EphB1* +/+ (WT), *EphB1* -/- (KO), and *EphB1* +/- (HET) male and female mice at 4-6 months age in a mixed 129/CD1/B6 background. Data from two independent experiments were pooled.

Statistical results of the mechanical pain in Zymoson injured mice - testing *EphrinB2* mutants

### Figure 5I

Mechanical Pain Ratio = $\frac{\text{baseline/uninjured left paw (adjusted filament)}}{\text{injured left paw (adjusted filament)}}$

two-way ANOVA

| Source of Variation | % of total variation | P value | Summary |
|---|---|---|---|
| Interaction | 9.50 | 0.0136 | * |
| Row Factor | 35.80 | <0.0001 | **** |
| Column Factor | 28.07 | 0.0002 | *** |

Tukey's post hoc:

Baseline WT vs Zymosan WT (**** *p* <0.0001)
Baseline MUT vs Zymosan WT (**** *p* <0.0001)
Zymosan WT vs Zymosan MUT (*** *p* = 0.0003)

unpaired t-test:
WT vs MUT (** *p* = 0.0039)

**Figure S8H (related to Figure 5I).** Two-way ANOVA with Tukey's post hoc analysis of the mechanical pain testing data shown in **Figure 5I** using Zymosan model that assessed *EphrinB2* lacZ/6YFdV reverse signaling mutant mice at 5-6 months age in a mixed 129/CD1/B6 background at 4-6 hr post-Zymosan.

Statistical results of the thermal and mechanical pain in Zymoson injured mice - testing 20 mg/kg oral dosing of 3511-I and BQPB4

### Figure 5J

$$\text{Thermal Pain Ratio} = \frac{\text{baseline/uninjured left paw response time (sec)}}{\text{injured left paw response time (sec)}}$$

two-way ANOVA (Thermal Pain Measurements)

| Source of Variation | % of total variation | P value | Summary |
|---|---|---|---|
| Interaction | 6.221 | 0.0176 | * |
| Row Factor | 83.25 | <0.0001 | **** |
| Column Factor | 4.061 | 0.0538 | ns |

Dunnett's post hoc (by treatment time):
4-6 hr: Vehicle vs 3511-I (* *p* = 0.0362)
4-6 hr: Vehicle vs BQPB4 (** *p* = 0.0021)

one-way ANOVA (Mechanical Pain Ratio) (** *p* = 0.0054)
Dunnett's post hoc:
4-6 hr: Vehicle vs 3511-I (* *p* = 0.0162)
4-6 hr: Vehicle vs BQPB4 (** *p* = 0.0039)

$$\text{Mechanical Pain Ratio} = \frac{\text{baseline/uninjured left paw (adjusted filament)}}{\text{injured left paw (adjusted filament)}}$$

two-way ANOVA (Mechanical Pain Measurements)

| Source of Variation | % of total variation | P value | Summary |
|---|---|---|---|
| Interaction | 13.90 | 0.2979 | ns |
| Row Factor | 24.33 | 0.0236 | * |
| Column Factor | 14.62 | 0.0880 | ns |

Dunnett's post hoc (by treatment group):
Vehicle: Baseline vs 6-8 hr (** *p* = 0.0278)
3511-I:: Baseline vs 2-4 hr (* *p* = 0.0354)

one-way ANOVA (Mechanical Pain Ratio) (* *p* = 0.0385)
Dunnett's post hoc:
6-8 hr: Vehicle vs BQPB4 (* *p* = 0.0303)

**Figure S8I (related to Figure 5J).** One-way and two-way ANOVA with Dunnett's post hoc analysis of the thermal and mechanical pain testing data shown in **Figure 5J** using Zymosan model that assessed a group of 5-6 month old WT adult male mice in a mixed 129/CD1 background that were orally dosed 15 min post-Zymosan with either vehicle only (3% DMSO/97% PBS) or with 20 mg/kg of 3511-I (2xHCl salt in 3% DMSO/97%

PBS) or 20 mg/kg BQPB4 (3xHCl salt in 3% DMSO/97% sunflower seed oil). Two hours after Zymosan injection, the injured left hind paws were assessed for mechanical pain (2-4 hr), then mice were orally dosed again with vehicle only or 3511-I/BQPB4 compounds, and then left hind paws tested for thermal pain (4-6 hr), and then again for mechanical pain (6-8 hr).

Thermal and mechanical pain in Zymoson injured mice - testing 5, 10, and 20 mg/kg oral dosing of A20

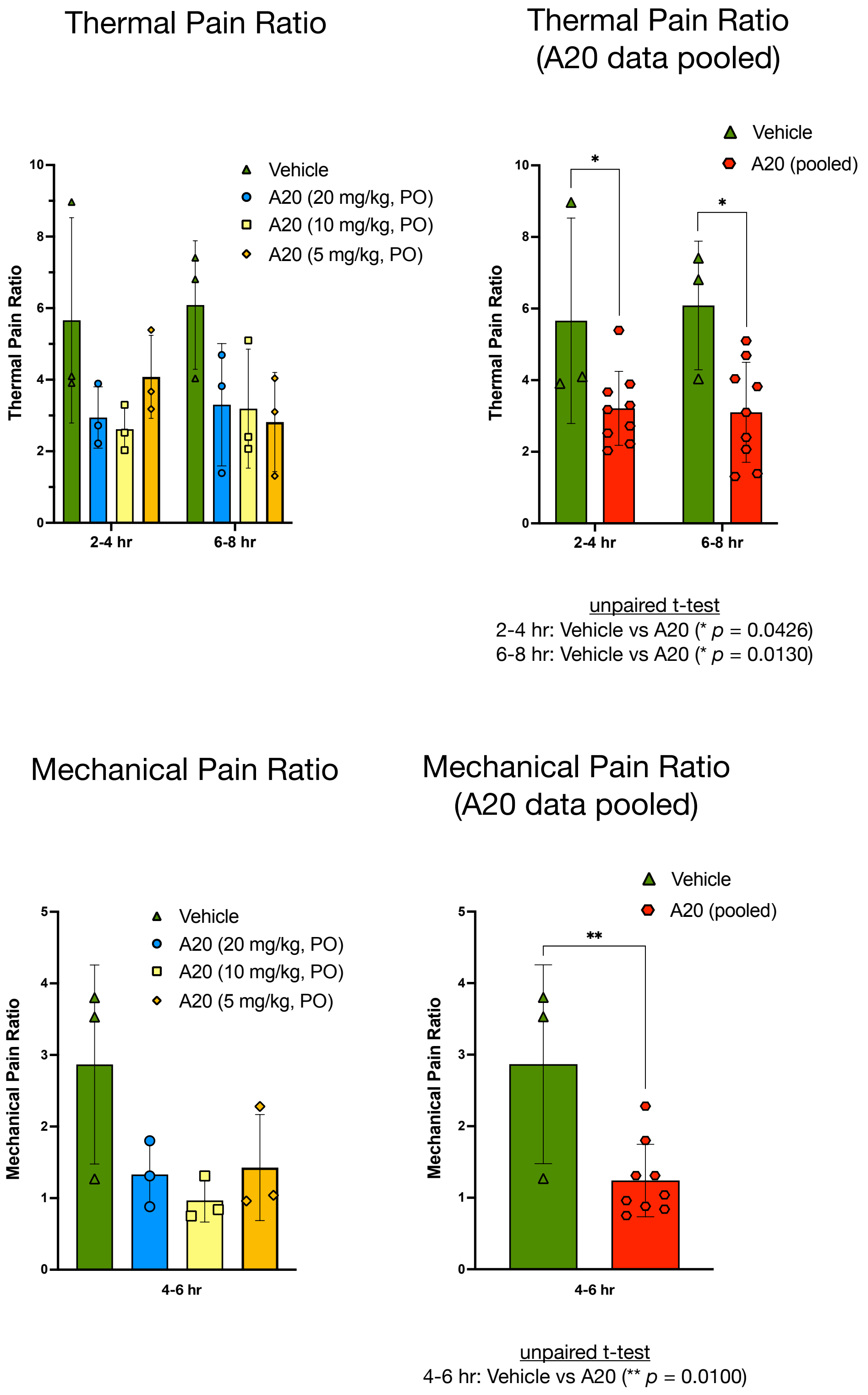

**Figure S8J (related to Figure 5K).** A20 oral dosing (PO) blunts Zymosan-induced thermal and mechanical pain in WT mice. Thermal and mechanical pain measurements and calculated pain ratios obtained from a group of Zymosan-injured 5-6 month old WT adult male mice in a 129x1/SvJ background that were orally dosed 15 min post-Zymosan with either vehicle (3% DMSO/97% PBS) or with 20, 10, or 5 mg/kg of A20 (2xHCl salt in 3% DMSO/97% PBS). Two hours after Zymosan injection, the hind paws were assessed for thermal pain using Hargreaves device (2-4 hr), then mice were orally dosed again with vehicle or A20, and then hind paws tested for mechanical pain (4-6 hr) and again for thermal pain (6-8 hr). The results of unpaired t-tests with significant values are presented below the relevant bar graph.

Statistical results of mechanical pain in Zymoson injured mice - testing 5, 10, and 20 mg/kg oral dosing of BQPB4

### Figure 5K

two-way ANOVA (Thermal Pain Measurements)

| Source of Variation | % of total variation | P value | Summary |
|---|---|---|---|
| Interaction | 3.770 | 0.0150 | * |
| Row Factor | 69.29 | <0.0001 | **** |
| Column Factor | 3.286 | 0.0253 | * |

Dunnett's post hoc (by treatment time):

4-6 hr: Vehicle vs 20 mg/kg BQPB4 (*** $p$ = 0.0004)
4-6 hr: Vehicle vs 10 mg/kg BQPB4 (* $p$ = 0.0143)

one-way ANOVA (Thermal Pain Ratio) (** $p$ = 0.0017)

Dunnett's post hoc:
4-6 hr: Vehicle vs 20 mg/kg BQPB4 (*** $p$ = 0.0010)
4-6 hr: Vehicle vs 10 mg/kg BQPB4 (** $p$ = 0.0078)
4-6 hr: Vehicle vs 5 mg/kg BQPB4 (* $p$ = 0.0289)

unpaired t-test
4-6 hr: Vehicle vs BQPB4 (pooled) (** $p$ <0.0001)

**Figure S8K (related to Figure 5K).** Thermal pain measurements and one-way and two-way ANOVA and unpaired t-test analysis from the mice shown in **Figure 5K** of a group of Zymosan-injured 3-5 month old WT adult male mice in a mixed 129/CD1 background that were orally dosed 15 min post-Zymosan with either vehicle only (3% DMSO/97% sunflower seed oil) or with 5, 10, or 20 mg/kg of BQPB4 (3xHCl salt in 3% DMSO/97% sunflower seed oil). Two hours after Zymosan injection, the hind paws were assessed for mechanical pain (2-4 hr), then mice were orally dosed again with vehicle only or BQPB4, and then hind paws tested for thermal pain (4-6 hr), and then again for mechanical pain (6-8 hr). Shown are the thermal pain results.

Example of the methods used to quantify Trap2+/Tom+ red fluorescent neurons in the dorsal horn of the lumbar spinal cord following CFA injury into the left hind paw

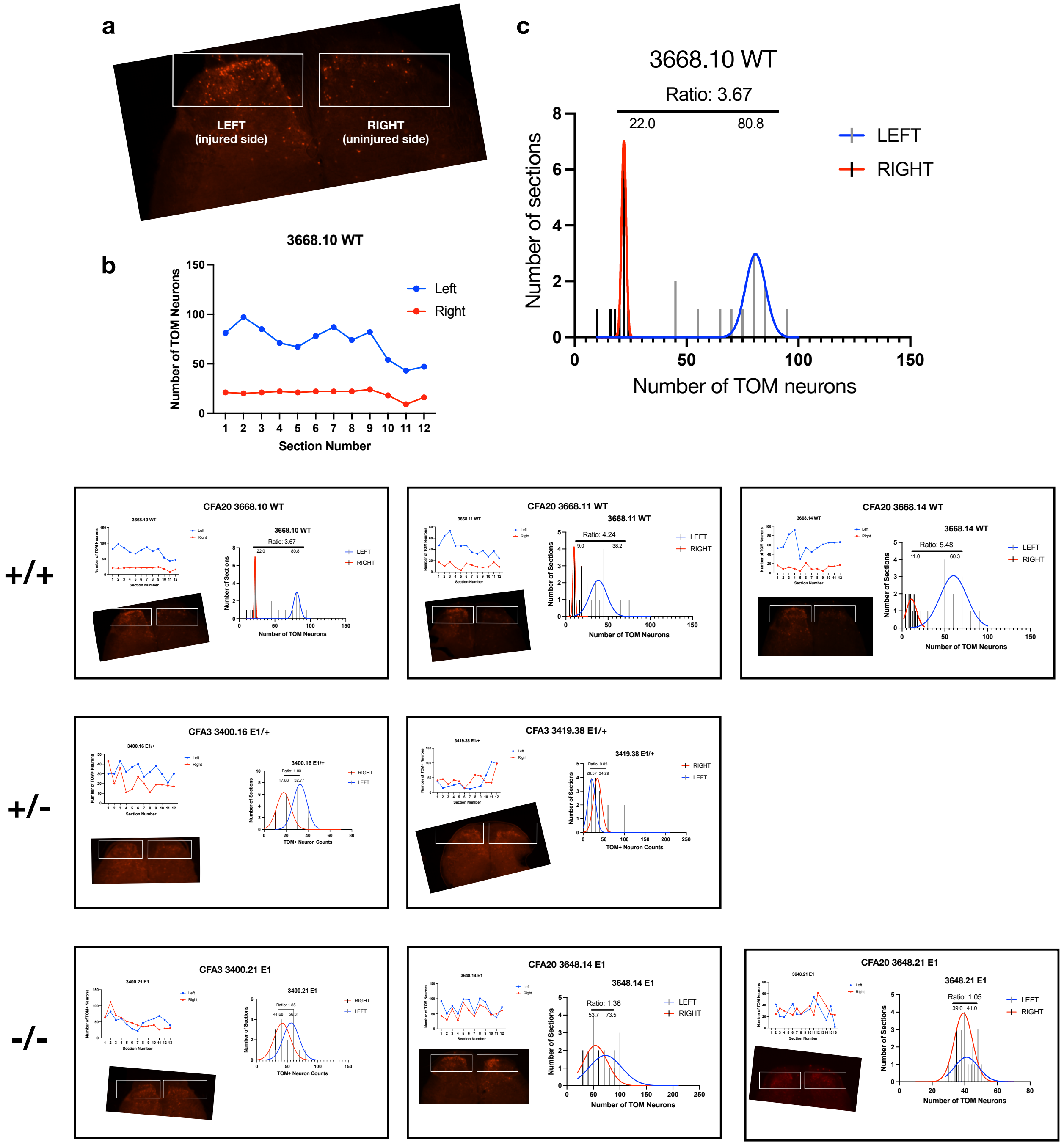

**Figure S9A (related to Figure 6).** Quantification methods using Trap2/Ai9-mediated Cre recombination following i.p. injection of 4-OHT at 4 hr post-CFA to permanently label with dtTomato red fluorescence those dorsal horn (DH) neurons that were activated in the lumbar spinal cord a short period after the inflammatory pain-generating insult in *EphB1* +/+ (WT), +/- (HET), and -/- (KO) mutant mice. In early analysis as shown here, 50 µm thick serial sections of perfusion-fixed spinal cords were cut using a vibratome in the transverse plane throughout the entire length of the lumbar cord of each animal to capture all sections. As serial sections were cut, they were placed into 10 section bins (500 µm per bin) and a total of 12 bins were collected spanning a 6 mm total length encompassing the lumbar spinal cord and regions above (rostral) and below (caudal). In these early studies, one section per each bin was mounted onto a slide and imaged to quantify Trap2/Ai9 driven red neurons, typically observing especially in WT (and vehicle-treated WT mice) that a consecutive group of 4 to 5 of the 12 bins usually contained the majority of sections that exhibited strong differences in the numbers of labeled neurons on the left (injured) side compared to the right (uninjured) side. We assumed these 4-5 bins represented the key ~2 mm segment of the lumbar spinal cord that was innervated by the sciatic nerve. Focusing in on this “sweet spot” of 4-5 bins of the lumbar cord, additional sections were then mounted and imaged, aiming for at least 12 sections to confirm neuron counts with many more slides. Shown here is a representative example of the data collected and analyzed, with (a) a typical fluorescent image taken using a Zeiss Axioscan to rapidly capture red labeled neurons in the lumbar spinal cord, (b) plotting the number of hand counted red fluorescent DH neurons from the left (injured) and right (uninjured) sides of 12 different “sweet spot” sections centered on the lumbar region, and (c) histogram plotting of the data as a function of the number of sections with X number of Trap2/Ai9 labeled red DH neurons, and dividing the mean of the number of labeled neurons on injured side by the mean of the number of labeled neurons on the control, uninjured side. The example shown gave a DH neuron activation ratio of 3.67 in the injured side, with additional quantification examples of *EphB1* WT, HET, and KO animals provided below.

A new computer program used to count and quantify red fluorescent labeled neurons in the dorsal horn of the lumbar spinal cord following CFA injury into the left hind paw

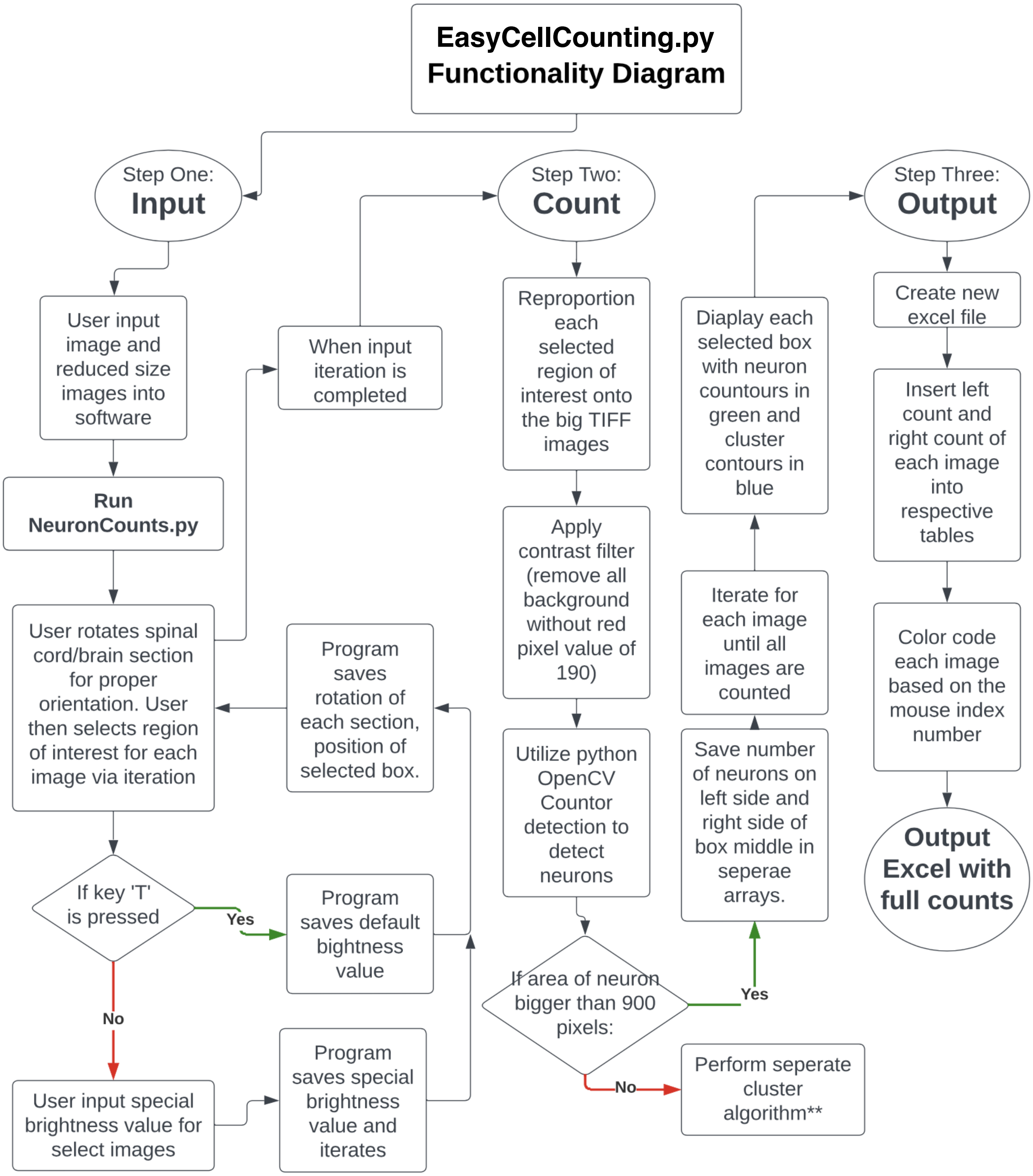

**Figure S9B (related to Figure 6).** Shown is the general program steps of a new computer program, EasyCellCounting.py, used to count and quantify labeled cells that are imaged within a selected area of interest to the end user. EasyCellCounting.py was

developed to replace the labor-intensive manual process of counting tdTomato red fluorescent neurons in the spinal cord induced by Trap2/Ai9/4-OHT with a highly optimized and efficient program and with the aim to minimize bias and increase the overall precision of the data. EasyCellCounting.py can be used for analysis of any type of cell labeling system, as long as signal-to-noise ratios are discernible over background levels, and is particularly useful for counting images of fluorescent labeled cells.

**A new computer program used to count and quantify red fluorescent labeled neurons in the dorsal horn of the lumbar spinal cord following CFA injury into the left hind paw**

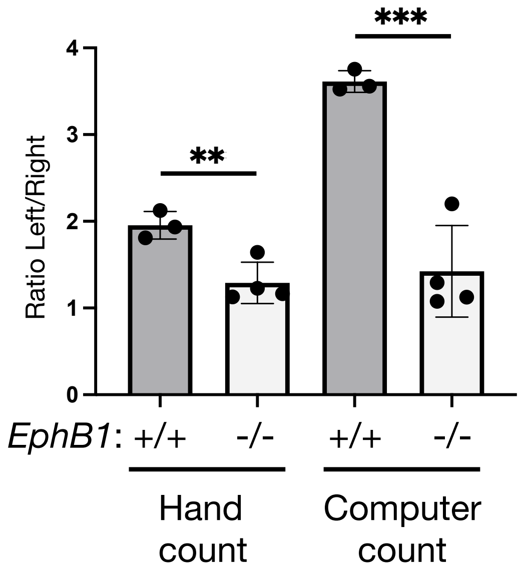

**Figure S9C (related to Figure 6).** Shown is the original test of EasyCellCounting.py, comparing results of the program to those obtained by manual counting of the same images. The t-tests indicate computer program is as good, if not better than hand counting at detecting differences in Trap2/Ai9 tdTomato-labeled lumbar DH neurons in *EphB1* +/+ (WT) and -/- (KO) mice subjected to the CFA pain model.

A new computer program used to count and quantify red fluorescent labeled neurons in the dorsal horn of the lumbar spinal cord - testing of 3511-I and BQPB4

Red neurons: injured side/uninjured side (average ratio for each section plotted)

Red neurons: injured side/uninjured side (average ratio for each animal plotted)

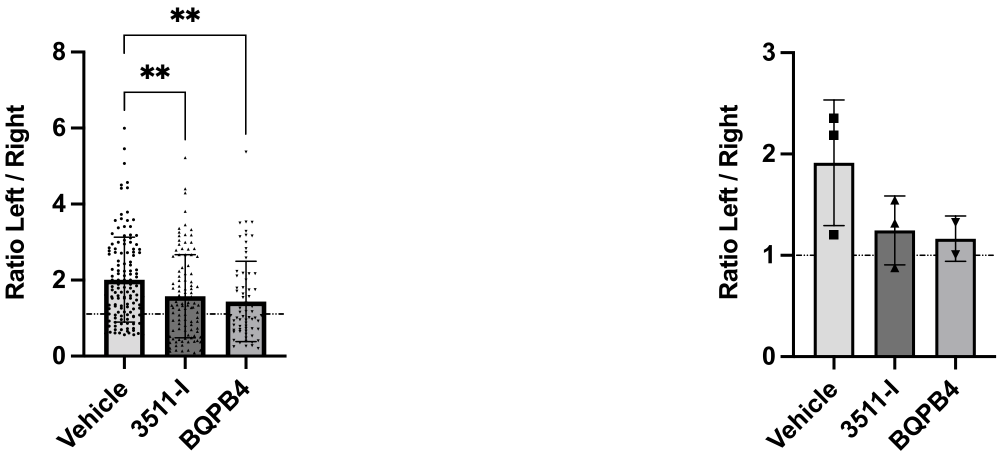

one-way ANOVA (*** *p* = 0.0005)
Tukey's multiple comparison:
1. Vehicle vs 3511-I (** *p* = 0.0071)
2. Vehicle vs BQPB4 (** *p* = 0.0014)

No significance found (n value too low)

**Figure S9D (related to Figure 6).** EasyCellCounting.py was used to quickly count and quantify the ratios of Trap2/Ai9/4-OHT red fluorescent neurons in sections of the spinal cord from Zymosan injured WT mice that were orally dosed with either vehicle, 3511-I, or BQPB4. Here, 307 Axioscan imaged spinal cord sections from eight +/+ (WT) mice subjected to Zymosan pain model and oral dosing of either vehicle (n=3), 3511-I (n=3), or BQPB4 (n=2) were inputted and analyzed. Data is plotted as the ratio of tdTomato red DH neurons (injured side/uninjured side) for each individual section (left), and as the average ratio for all sections of an individual spinal cord (right). Note that when all sections were individually plotted, those from the 3511-I and BQPB4 treated mice exhibited significantly lower ratios of red neurons compared to the vehicle treated animals, though even the vehicles showed a fairly mild ratio of 2. We suspect this may be due to the Zymosan model not strongly activating DH pain neurons as much as the CFA-induced pain model. Consistent with this reduced activation in Zymosan model, there was also a trend towards lower ratios in the 3511-I and BQPB4 treated mice when the sections from each mouse were averaged and plotted as a single value, though the numbers of mice analyzed here were too low to reveal statistical significance.

### Supplementary Chemistry Information

#### Materials and Methods

All chemicals for synthesis were obtained from commercial suppliers (Thermo Scientific, Sigma-Aldrich, or Ambeed) in “reagent grade” purity or better and used as received. Deuterated solvents used for NMR experiments were purchased from Cambridge Isotope Laboratories.

For purification of reaction products by flash column chromatography, silica gel from Sigma-Aldrich (Silica gel, high-purity grade, pore size 60 Å, 220-440 mesh particle size, 35-75 µm particle size) was used. Thin layer chromatography was performed using aluminum sheets pre-coated with silica gel 60 matrix and fluorescent indicator for 254 nm, purchased from Millipore Sigma. Visualization of the compounds occurred under UV light of 254 nm wavelength.

NMR spectra were recorded on a Bruker Avance NEO instrument operating at either 400 or 600 MHz proton frequency. Samples were recorded at either 295 k or 323 K in 5 mm diameter tubes. Coupling patterns are abbreviated with s (singlet), d (doublet), dd (doublet of doublets), ddd (doublet of doublet of doublets), t (triplet), hept (heptet) and m (multiplet) with indicated coupling constants J in Hertz (Hz).

High-resolution mass spectra (HRMS) were recorded by Dr. Hamid Baniasadi (UT Southwestern Medical Center, Metabolomics Core) on Sciex TripleTOF® 6600 System. The data was processed by Sciex OS software.

### Synthetic Schemes

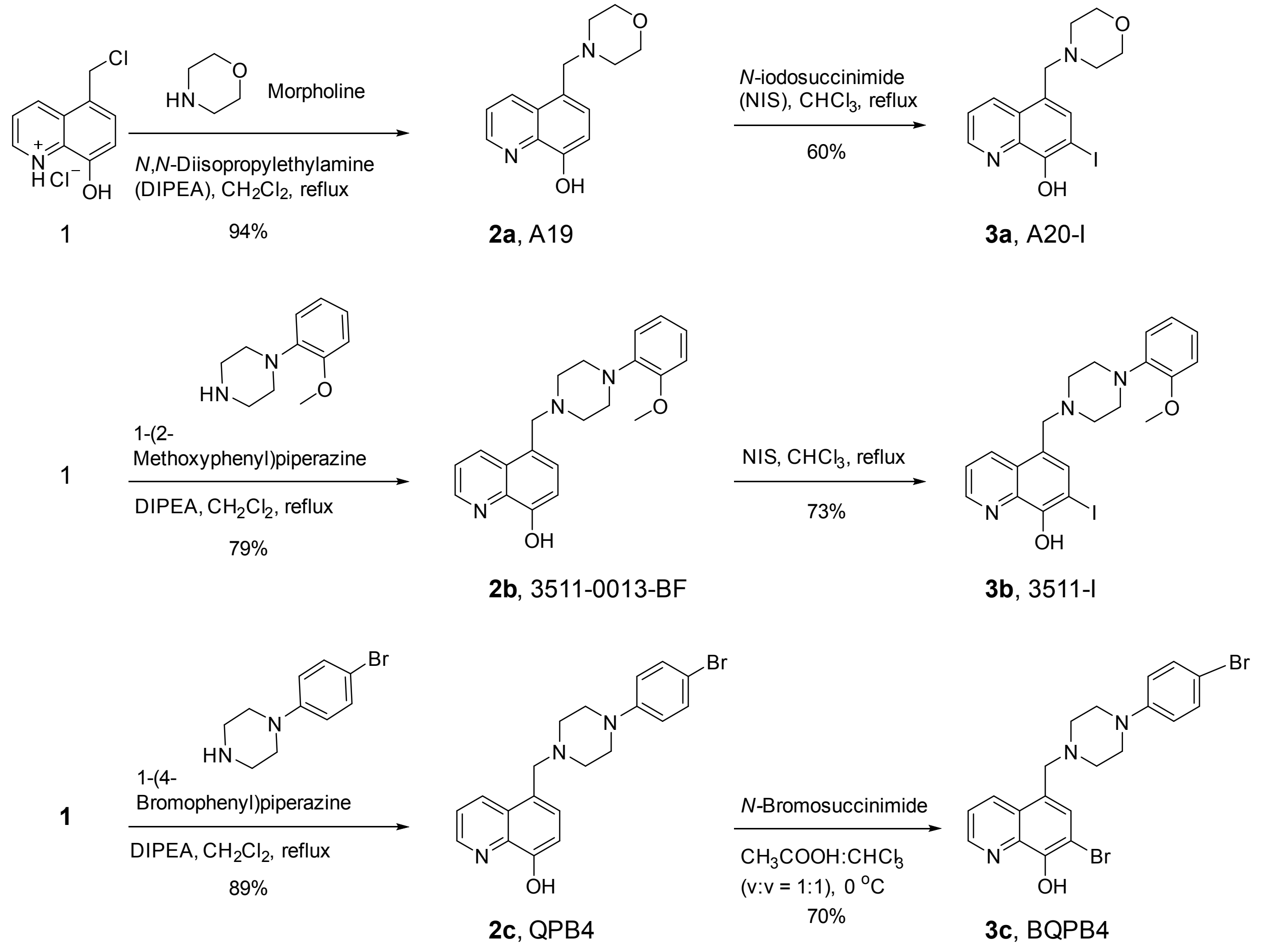

Synthetic route to **3a** (A20-I), **3b** (3511-I), and **3c** (BQPB4).

### Synthetic Procedures

Compound **1** was prepared according to published procedures with modification.[1] Intermediate compounds **2a**-**2c** (i.e., A19, 3511-0013-BF, and QPB4) were synthesized according to published procedures with modification.[2,3] Final products **3a**-**3c** (i.e., A20-I, 3511-I, and BQPB4) were prepared according to published procedures with modification.[3]

## 2a (A19)

To a 500 mL round bottom flask, compound A (10.0 g, 0.04346 mol, 1.0 eq.) and morpholine (4.12 mL, 0.04781 mol, 1.1 eq.) as well as 160 mL dichloromethane were added. While stirring, *N,N*-diisopropylethylamine (DIPEA, 15.14 mL, 0.08692 mol, 2.0 eq.) was added dropwise to the suspension generating a clear solution. The reaction mixture was refluxed overnight. Following completion, the organic solvent was removed under reduced pressure. The crude mixture was further solidified in hexane, and the organic solvent was removed under reduced pressure. The crude solid was redispersed in 30 mL cold DI water, filtered on a sintered glass crucible under vacuum and washed with minimum amount of cold acetone. The product was obtained as a white solid in 94% yield (**A19**, 10.0 g).

$^{1}$**H NMR** (400 MHz, 298 K, $CDCl_3$) δ 8.78 (dd, *J* = 4.2, 1.6 Hz, 1H), 8.67 (dd, *J* = 8.6, 1.6 Hz, 1H), 7.46 (dd, *J* = 8.5, 4.2 Hz, 1H), 7.33 (d, *J* = 7.7 Hz, 1H), 7.07 (d, *J* = 7.7 Hz, 1H), 3.79 (s, 2H), 3.66 (t, *J* = 4.6 Hz, 4H), 2.45 (t, *J* = 4.6 Hz, 4H).

$^{13}$**C NMR** (101 MHz, 298 K, $CDCl_3$) δ 151.9, 147.6, 138.7, 134.1, 129.1, 127.9, 124.1, 121.5, 108.6, 67.1, 61.1, 53.6.

## 3a (A20-I)

In a 50 mL round bottom flask, A19 (300 mg, 1.2280 mmol, 1.0 eq.) was dissolved in 10 mL chloroform at room temperature. *N*-iodosuccinimide (NIS, 290.1 mg, 1.2894 mmol, 1.05 eq.) was added to the flask. The reaction mixture was stirred overnight at 50 $^{o}$C. Following completion, the organic solvent was removed under vacuum. The crude solid was taken up in ethyl acetate and washed with saturated sodium bicarbonate, brine, and DI water. The organic fractions were collected, dried over anhydrous magnesium sulfate, filtered and concentrated. The crude solid was purified with column chromatography on silica gel (ethyl acetate:hexane, 1:1, v:v). The produce was obtained as an off-white solid in 60% yield (**A20-I**, 275.0 mg).

$^{1}$**H NMR** (400 MHz, 298 K, $CDCl_3$) δ 8.75 (dd, *J* = 4.3, 1.5 Hz, 1H), 8.64 (dd, *J* = 8.6, 1.6 Hz, 1H), 7.66 (s, 1H), 7.48 (dd, *J* = 8.5, 4.2 Hz, 1H), 3.74 (s, 2H), 3.66 (t, *J* = 4.6 Hz, 4H), 2.45 (t, *J* = 4.7 Hz, 4H).

$^{13}$**C NMR** (101 MHz, 298 K, CDCl3) δ 152.7, 148.3, 137.8, 136.9, 134.4, 127.6, 125.9, 121.9, 75.5, 67.0, 60.5, 53.5.

**HRMS** (ESI) m/z: [M+H$^{+}$] Calculated for C14H16IN2O2 371.02510; Found 371.02473.

## 2b (3511-0013-BF)

To a 100 mL round bottom flask, compound A (2.0 g, 0.008692 mol, 1.0 eq.) and 1-(2-methoxyphenyl)piperazine) (1.7 g, 0.008866 mol, 1.02 eq.) as well as 35 mL dichloromethane were added. While stirring, *N,N*-diisopropylethylamine (DIPEA, 3.79 mL, 0.02173 mol, 2.5 eq.) was added dropwise to the suspension generating a clear solution. The reaction mixture was refluxed overnight. Following completion, the organic solvent was removed under reduced pressure. The crude mixture was taken up in chloroform and washed with saturated sodium bicarbonate (aq.) and DI water. The organic fractions were collected, dried over anhydrous magnesium sulfate, filtered and concentrated. The crude solid was redispersed in 100 mL DI water, filtered on a sintered glass crucible under vacuum and washed with DI water and minimum amount of cold acetone. The product was obtained as a white to pale green solid in 79% yield (**3511-0013-BF**, 2.4 g).

$^{1}$**H NMR** (400 MHz, 298 K, $CDCl_3$) δ 8.79 (dd, *J* = 4.2, 1.5 Hz, 1H), 8.72 (dd, *J* = 8.6, 1.6 Hz, 1H), 7.47 (dd, *J* = 8.5, 4.2 Hz, 1H), 7.38 (d, *J* = 7.7 Hz, 1H), 7.09 (d, *J* = 7.7 Hz, 1H), 6.98 (ddd, *J* = 8.3, 5.4, 3.6 Hz, 1H), 6.90 (d, *J* = 3.7 Hz, 2H), 6.85 (d, *J* = 7.9 Hz, 1H), 3.87 (d, *J* = 5.4 Hz, 5H), 3.04 (s, 4H), 2.67 (t, *J* = 4.9 Hz, 4H).

$^{13}$**C NMR** (101 MHz, 298 K, $CDCl_3$) δ 152.3, 151.8, 147.6, 141.42, 138.7, 134.2, 129.0, 128.0, 124.7, 122.8, 121.5, 120.9, 118.2, 111.2, 108.7, 60.8, 55.4, 53.4, 50.8.

## 3b (3511-I)

In a 50 mL round bottom flask, 3511-0013-BF (200.0 mg, 0.5724 mmol, 1.0 eq.) was dissolved in 10 mL chloroform at room temperature. *N*-iodosuccinimide (NIS, 134.0 mg, 0.5956 mmol, 1.04 eq.) was added to the flask. The reaction mixture was stirred overnight at 50 $^{o}$C. Following completion, the organic solvent was removed under vacuum. The crude solid was taken up in ethyl acetate and washed with saturated sodium bicarbonate, brine, and DI water. The organic fractions were collected, dried over anhydrous magnesium sulfate, filtered and concentrated. The crude solid was purified with column chromatography on silica gel (ethyl acetate:hexane, 1:1, v:v). The produce was obtained as an off-white solid in 73% yield (**3511-I**, 198.3 mg).

$^{1}$**H NMR** (400 MHz, 298 K, $CDCl_3$) δ 8.75 (d, *J* = 4.1 Hz, 1H), 8.70 (dd, *J* = 8.6, 1.5 Hz, 1H), 7.71 (s, 1H), 7.49 (dd, *J* = 8.5, 4.2 Hz, 1H), 7.04 – 6.94 (m, 1H), 6.90 (d, *J* = 4.4 Hz, 2H), 6.85 (d, *J* = 7.9 Hz, 1H), 3.84 (d, *J* = 12.4 Hz, 5H), 3.05 (s, 4H), 2.69 (d, *J* = 4.9 Hz, 4H).

$^{13}$**C NMR** (101 MHz, 298 K, $CDCl_3$) δ 152.7, 152.3, 148.3, 141.3, 137.8, 136.9, 134.6, 127.7, 126.4, 122.9, 121.9, 121.0, 118.2, 111.1, 75.6, 60.1, 55.4, 53.3, 50.7.

**HRMS** (ESI) m/z: [M+H$^{+}$] Calculated for C21H23IN3O2 476.08295; Found 476.08296.

#### 2c (QPB4)

To a 250 mL round bottom flask, compound A (5.6 g, 0.02438 mol, 1.0 eq.) and 1-(4-bromophenyl)piperazine (6.0 g, 0.02488 mol, 1.02 eq.) as well as 100 mL dichloromethane and 30 mL chloroform were added. While stirring, *N,N*-diisopropylethylamine (DIPEA, 12.74 mL, 0.07312 mol, 3.0 eq.) was added dropwise to the suspension. The reaction mixture was refluxed overnight. Following completion, the organic solvent was removed under reduced pressure. The crude mixture was further solidified in hexane, and the organic solvent was removed under reduced pressure. The crude solid was redispersed in 40 mL DI water, filtered on a sintered glass crucible under vacuum and washed with DI water, cold acetone, and cold dichloromethane. The product was obtained as a white to pale green solid in 89% yield (**QPB4**, 8.6 g).

$^{1}$**H NMR** (400 MHz, 298 K, $CDCl_3$) δ 8.79 (dd, *J* = 4.2, 1.6 Hz, 1H), 8.69 (dd, *J* = 8.6, 1.6 Hz, 1H), 7.46 (dd, *J* = 8.5, 4.2 Hz, 1H), 7.39 – 7.27 (m, 3H), 7.09 (d, *J* = 7.7 Hz, 1H), 6.80 – 6.70 (m, 2H), 3.85 (s, 2H), 3.16 – 3.06 (m, 4H), 2.66 – 2.53 (m, 4H).

$^{13}$**C NMR** (101 MHz, 298 K, $CDCl_3$) δ 151.9, 150.4, 147.6, 138.8, 134.1, 131.8, 129.0, 127.9, 124.3, 121.5, 117.6, 111.7, 108.7, 60.7, 52.9, 49.0.

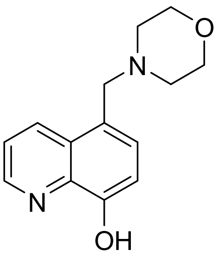

#### 3c (BQPB4)

In a 100 mL round bottom flask, QPB4 (150.0 mg, 0.3766 mmol, 1.0 eq.) was dissolved in a mixture of chloroform and acetic acid (v:v, 1:1, 7 ml : 7 mL). The mixture was cooled down to 0 $^{o}$C in ice bath for 5-10 min. *N*-bromosuccinimide (69.0 mg, 0.3877 mmol, 1.03 eq.) was added to the mixture portion by portion and avoiding light. The reaction mixture was stirred overnight. Following completion, the mixture was transferred to a beaker in an ice batch and 10 mL of DI water was added. After adjusting pH to 5-6 using 6 M sodium hydroxide (aq.), the mixture was extracted with ethyl acetate. The organic fraction was washed with saturated sodium bicarbonate (aq.), brine, and DI water. The organic fraction was then dried over anhydrous magnesium sulfate, filtered, and concentrated. The crude solid was further purified with column chromatography on silica gel (ethyl acetate:hexane, 1:1, v:v). The product was obtained as an off-white solid in 70% yield (**BQPB4**, 126.1 mg)

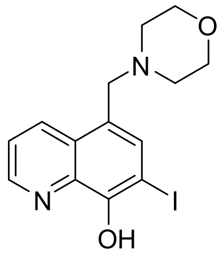

**$^{1}$H NMR** (600 MHz, 323 K, $CDCl_3$) δ 8.79 (dd, *J* = 4.3, 1.6 Hz, 1H), 8.68 (d, *J* = 8.5 Hz, 1H), 7.60 – 7.54 (m, 1H), 7.52 – 7.47 (m, 1H), 7.34 – 7.30 (m, 2H), 6.77 – 6.73 (m, 2H), 3.85 (s, 2H), 3.14 (s, 4H), 2.71 – 2.57 (m, 4H).

**$^{13}$C NMR** (151 MHz, 323 K, $CDCl_3$) δ 150.3, 149.5, 148.4, 138.9, 134.2, 132.1, 131.9, 126.9, 125.5, 121.6, 117.63, 111.8, 102.8, 60.1, 52.8, 49.0.

**HRMS** (ESI) m/z: [M+H$^+$] Calculated for C20H20Br2N3O 475.99676; Found 475.99466.

#### General Procedure for Preparing Salts

In a round bottom flask, a free base (1.0 eq.) was dissolved in dichloromethane or chloroform. The solution was cooled down to 0 $^oC$ in an ice batch for 5-10 min. A solution of 2 M hydrochloride in diethyl ether (2.0 or 3.0 e.q.) was added dropwise and a precipitation immediately formed. The reaction mixture was stirred for a couple of hours, filtered under vacuum, and washed with dichloromethane. The solid was transferred to a scintillation vial and kept under vacuum to give dried salt.

##### A20-I hydrochloride salt (A20-I·2xHCl)

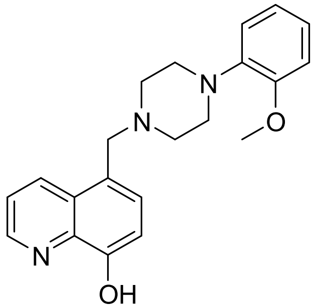

A20-I·2xHCl was prepared according to General Procedure for Preparing Salts using free base A20-I (200.0 mg, 0.5403 mmol, 1.0 eq.) and 2 M hydrochloride in diethyl ether (540.3 µL, 1.0806 mmol, 2.0 eq.). The product was obtained as a light-yellow solid in 91% yield (217.8 mg).

**$^{1}$H NMR** (400 MHz, 298 K, DMSO-*d6*) δ 11.30 (s, 1H), 9.04 (d, J = 8.6 Hz, 1H), 8.94 (d, J = 4.2 Hz, 1H), 8.35 (s, 1H), 7.76 (dd, J = 8.6, 4.3 Hz, 1H), 4.74 (d, J = 4.0 Hz, 2H), 3.96 – 3.75 (m, 4H), 3.24 (s, 4H).

**$^{13}$C NMR** (101 MHz, 298 K, DMSO-*d6*) δ 155.2, 148.8, 142.2, 136.8, 135.6, 128.8, 123.2, 117.9, 81.0, 63.5, 54.2, 51.1.

##### 3511-I hydrochloride salt (3511-I·2xHCl)

3511-I·2xHCl was prepared according to General Procedure for Preparing Salts using free base 3511-I (132.1 mg, 0.2779 mmol, 1.0 eq.) and 2 M hydrochloride in diethyl ether (277.9 µL, 0.5558 mmol, 2.0 eq.). The product was obtained as a light-yellow solid in 93% yield (142.2 mg).

**$^{1}$H NMR** (400 MHz, 298 K, DMSO-*d6*) δ 11.01 (s, 1H), 9.02 (dd, *J* = 8.7, 1.5 Hz, 1H), 8.95 (dd, *J* = 4.3, 1.4 Hz, 1H), 8.37 (s, 1H), 7.78 (dd, *J* = 8.6, 4.3 Hz, 1H), 6.99 (dtd, *J* = 16.2, 8.3, 2.0 Hz, 2H), 6.94 – 6.85 (m, 2H), 4.77 (s, 2H), 3.78 (s, 3H), 3.49 – 3.32 (m, 6H), 3.15 – 3.02 (m, 2H).

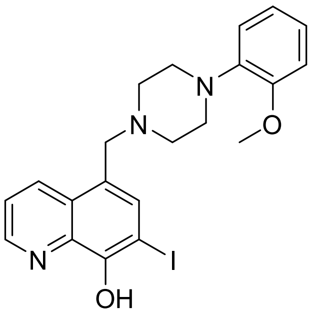

**$^{13}$C NMR** (101 MHz, 298 K, DMSO-*d6*) δ 155.4, 152.3, 149.0, 142.1, 139.7, 137.1, 135.1, 128.7, 124.1, 123.3, 121.3, 118.8, 118.1, 112.4, 80.6, 55.8, 53.9, 51.3, 47.3.

#### BQPB4 hydrochloride salt (BQPB4·3xHCl)

BQPB4·3xHCl was prepared according to General Procedure for Preparing Salts using free base BQPB4 (60.1 mg, 0.1259 mmol, 1.0 eq.) and 2 M hydrochloride in diethyl ether (188.9 µL, 0.3777 mmol, 3.0 eq.). The product was obtained as a light-yellow solid in 98% yield (72 mg).

**$^{1}$H NMR** (400 MHz, 298 K, DMSO-*d6*) δ 11.19 (s, 1H), 9.04 (d, *J* = 8.6 Hz, 1H), 8.99 (d, *J* = 4.2 Hz, 1H), 8.25 (s, 1H), 7.78 (dd, *J* = 8.6, 4.3 Hz, 1H), 7.45 – 7.33 (m, 2H), 7.00 – 6.86 (m, 2H), 4.81 (s, 2H), 3.80 (d, *J* = 12.9 Hz, 2H), 3.40 (d, *J* = 11.6 Hz, 2H), 3.33 (dd, *J* = 13.1, 7.5 Hz, 2H), 3.25 – 3.14 (m, 2H).

**$^{13}$C NMR** (101 MHz, 298 K, DMSO-*d6*) δ 152.2, 149.1, 149.0, 138.0, 137.1, 135.7, 132.2, 128.3, 123.1, 118.3, 117.6, 111.6, 105.8, 53.8, 50.5, 45.5.

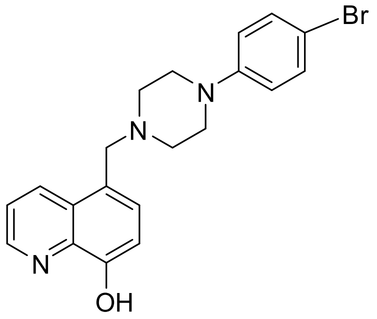

### NMR Spectra

## 2a (A19)

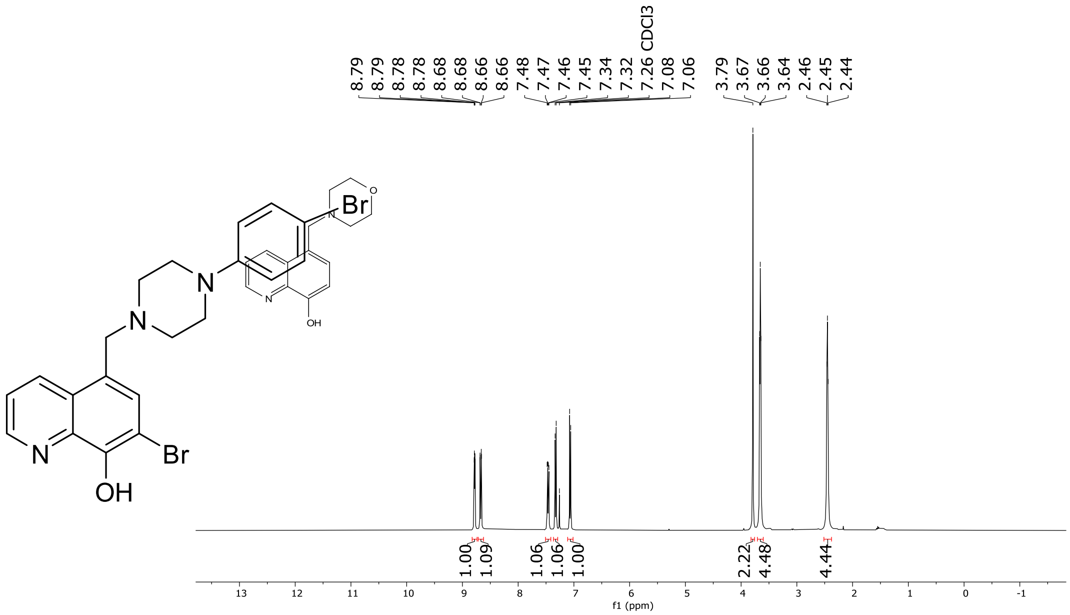

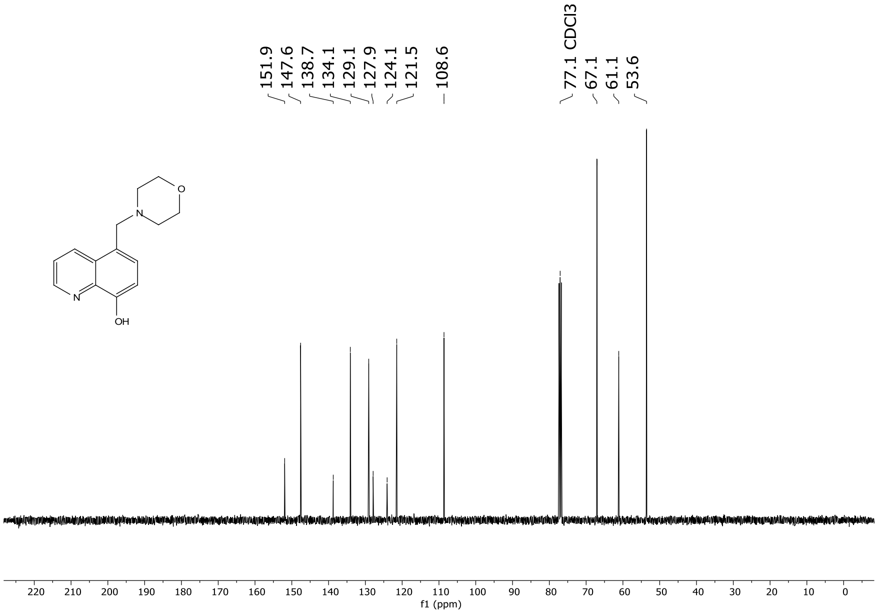

## 2a (A20-I)

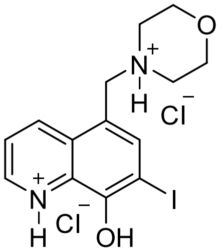

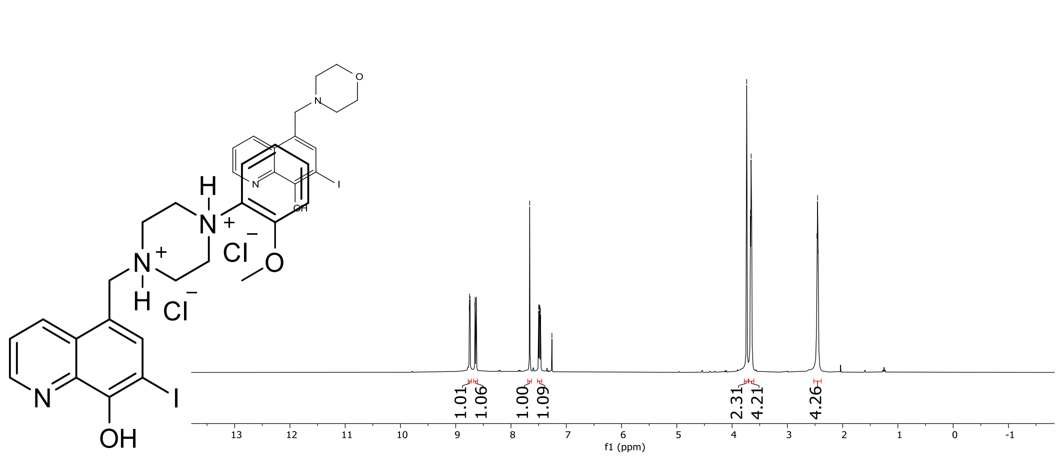
8.75
8.75
8.74
8.74
8.65
8.65
8.63
8.63
7.66
7.50
7.49
7.48
7.47
7.26 CDCl3
3.74
3.67
3.66
3.64
2.46
2.45
2.44
1.01
1.06
1.00
1.09
2.31
4.21
4.26
f1 (ppm)

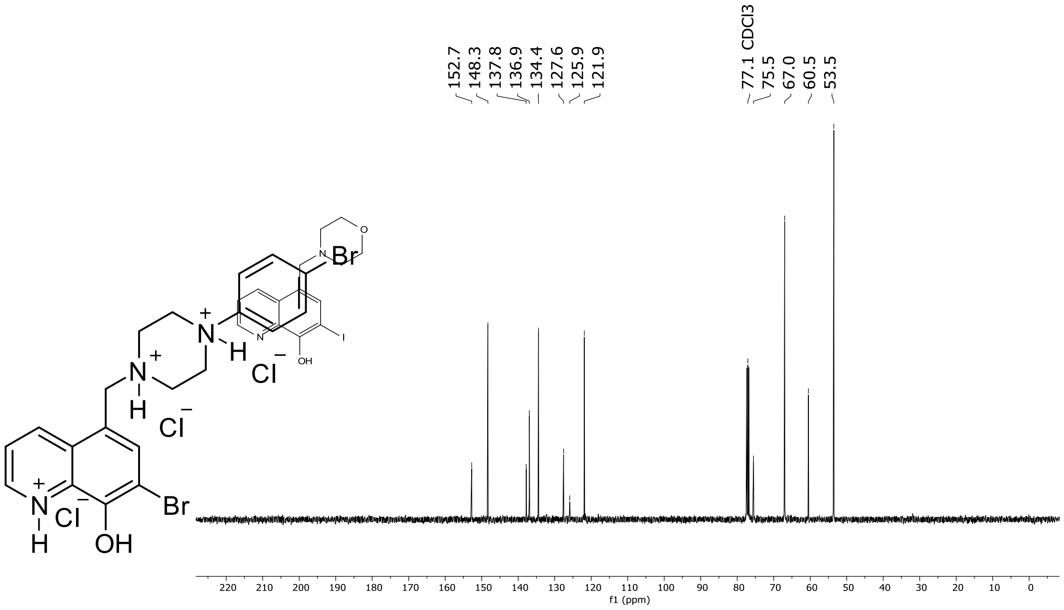
152.7
148.3
137.8
136.9
134.4
127.6
125.9
121.9
77.1 CDCl3
75.5
67.0
60.5
53.5
f1 (ppm)

## 2b (3511-0013-BF)

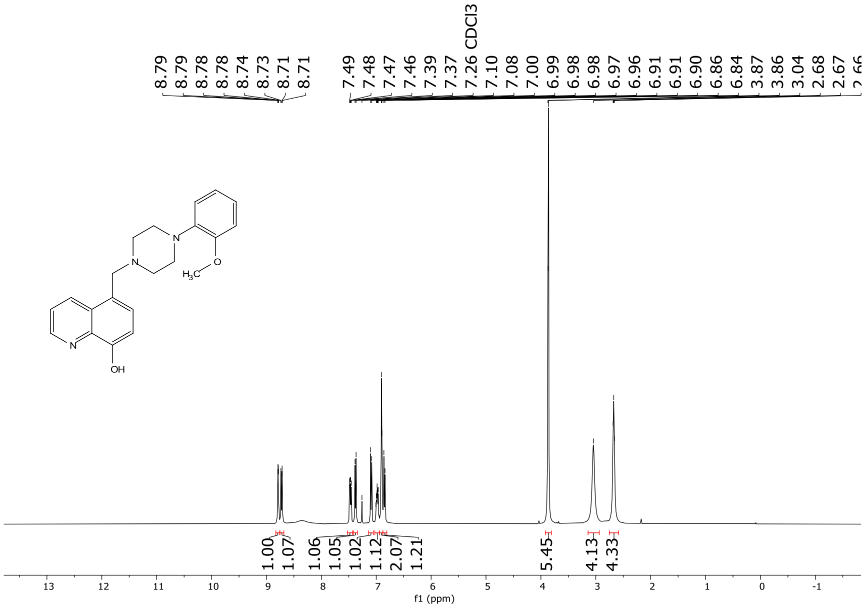

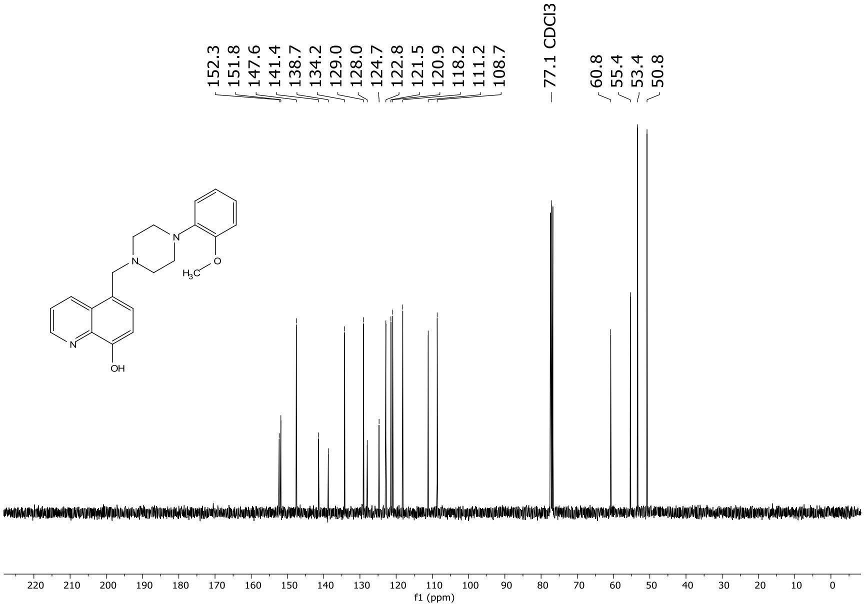

**3b (3511-I)**

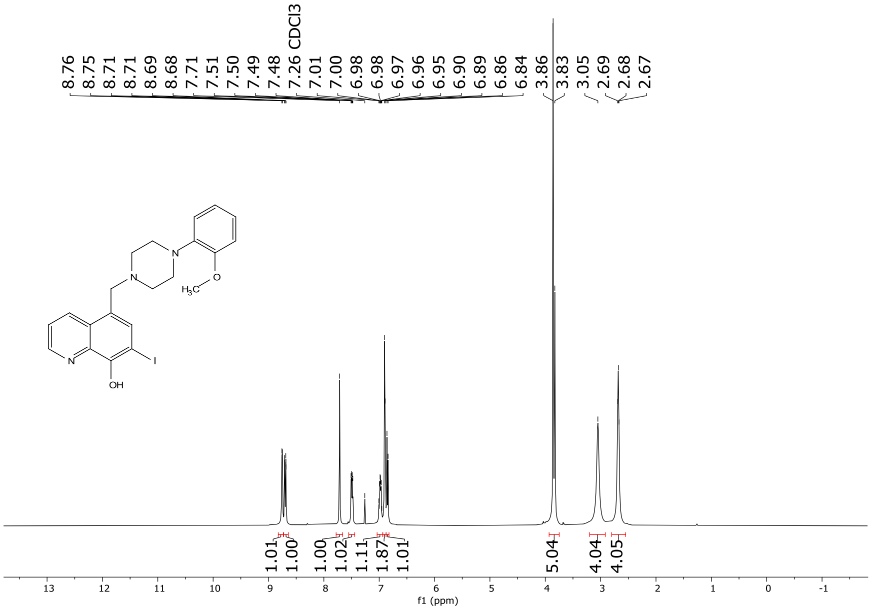

8.76
8.75
8.71
8.71
8.69
8.68
7.71
7.51
7.50
7.49
7.48
7.26 CDCl3
7.01
7.00
6.98
6.98
6.97
6.96
6.95
6.90
6.89
6.86
6.84
3.86
3.83
3.05
2.69
2.68
2.67
1.01
1.00
1.00
1.02
1.11
1.87
1.01
5.04
4.04
4.05
f1 (ppm)

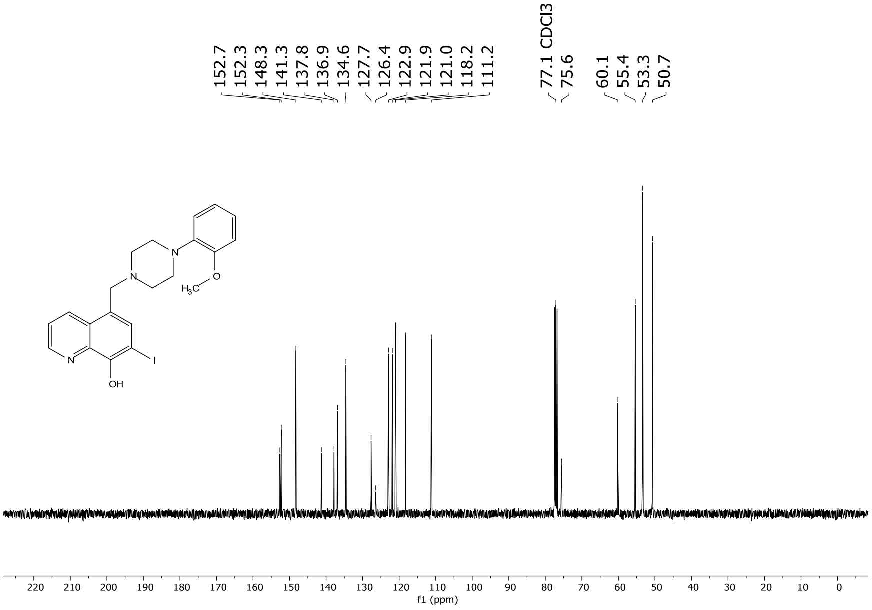

152.7
152.3
148.3
141.3
137.8
136.9
134.6
127.7
126.4
122.9
121.9
121.0
118.2
111.2
77.1 CDCl3
75.6
60.1
55.4
53.3
50.7
f1 (ppm)

#### 3a (QPB4)

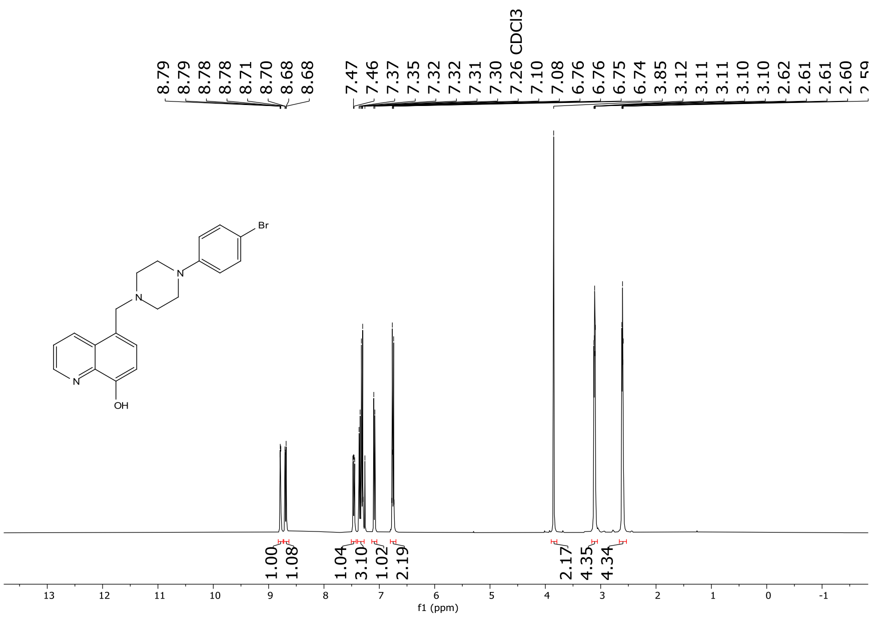
8.79
8.79
8.78
8.78
8.71
8.70
8.68
8.68
7.47
7.46
7.37
7.35
7.32
7.32
7.31
7.30
7.26 CDCl3
7.10
7.08
6.76
6.76
6.75
6.74
3.85
3.12
3.11
3.11
3.10
3.10
2.62
2.61
2.61
2.60
Br
N
N
N
OH
1.00
1.08
1.04
3.10
1.02
2.19
2.17
4.35
4.34
f1 (ppm)

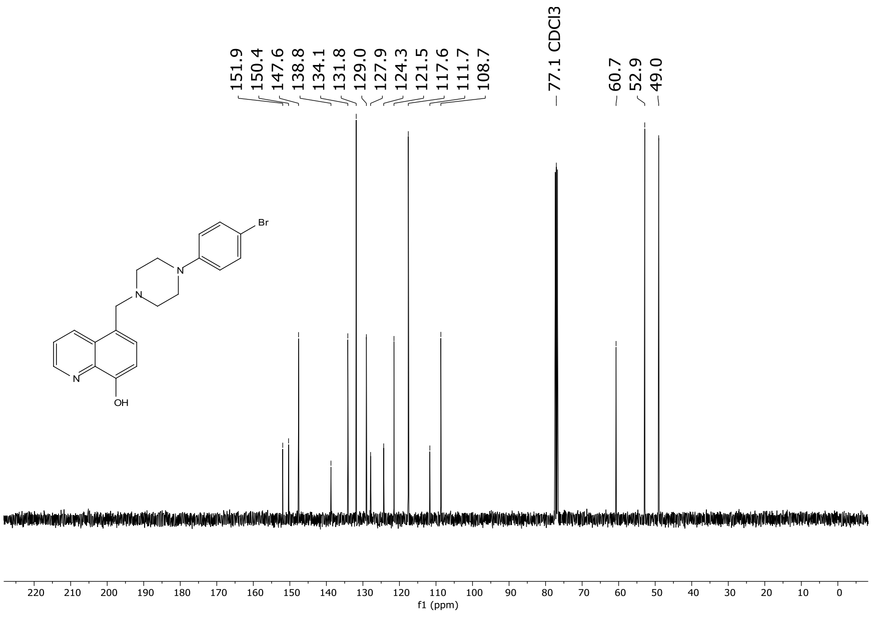
151.9
150.4
147.6
138.8
134.1
131.8
129.0
127.9
124.3
121.5
117.6
111.7
108.7
77.1 CDCl3
60.7
52.9
49.0
Br
N
N
N
OH
f1 (ppm)

### 3c (BQPB4)

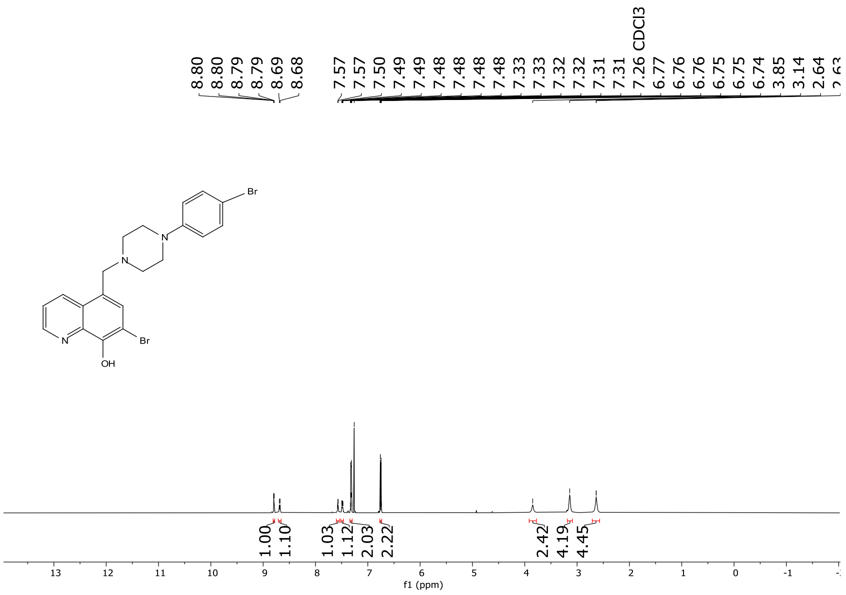
8.80
8.80
8.79
8.79
8.69
8.68
7.57
7.57
7.50
7.49
7.49
7.48
7.48
7.48
7.48
7.33
7.33
7.32
7.32
7.31
7.31
7.26 CDCl3
6.77
6.76
6.76
6.75
6.75
6.74
3.85
3.14
2.64
Br
N
N
N
Br
OH
1.00
1.10
1.03
1.12
2.03
2.22
2.42
4.19
4.45
f1 (ppm)

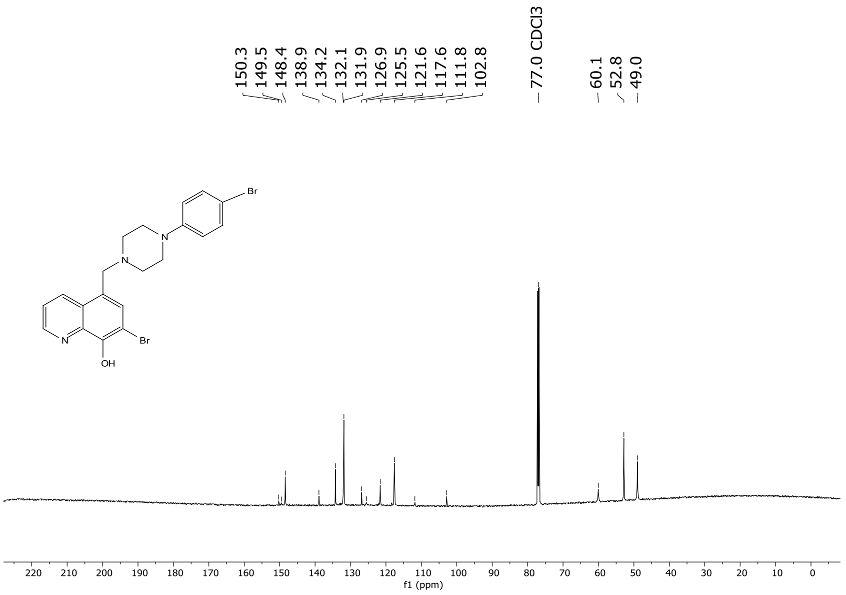
150.3
149.5
148.4
138.9
134.2
132.1
131.9
126.9
125.5
121.6
117.6
111.8
102.8
77.0 CDCl3
60.1
52.8
49.0
Br
N
N
N
Br
OH
f1 (ppm)

### A20-I hydrochloride salt (A20-I·2xHCl)

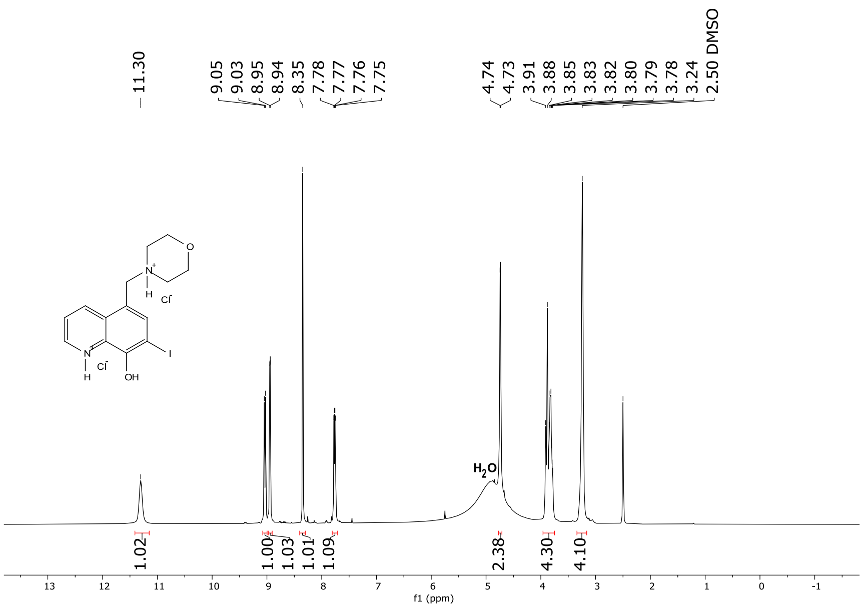

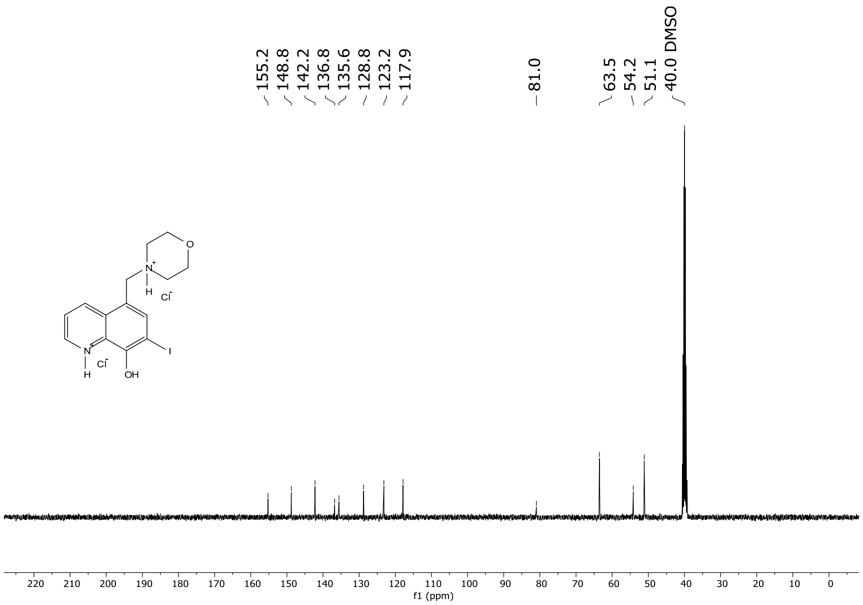

### 3511-I hydrochloride salt (3511-I·2xHCl)

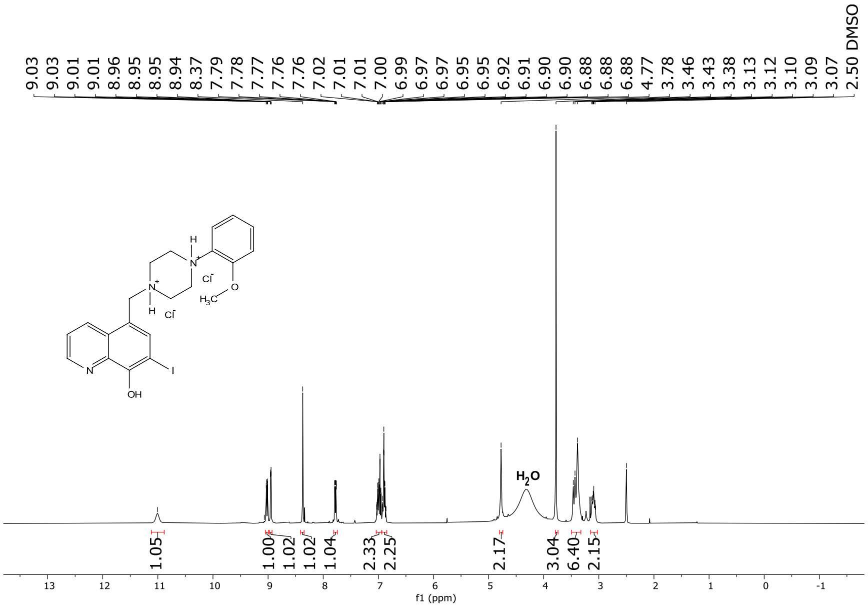

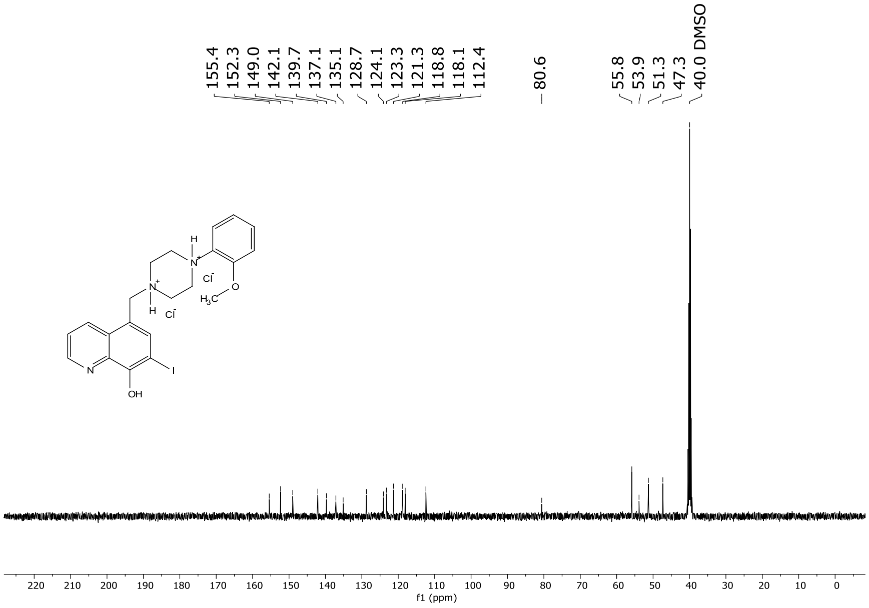

### BQPB4 hydrochloride salt (BQPB4·3xHCl)

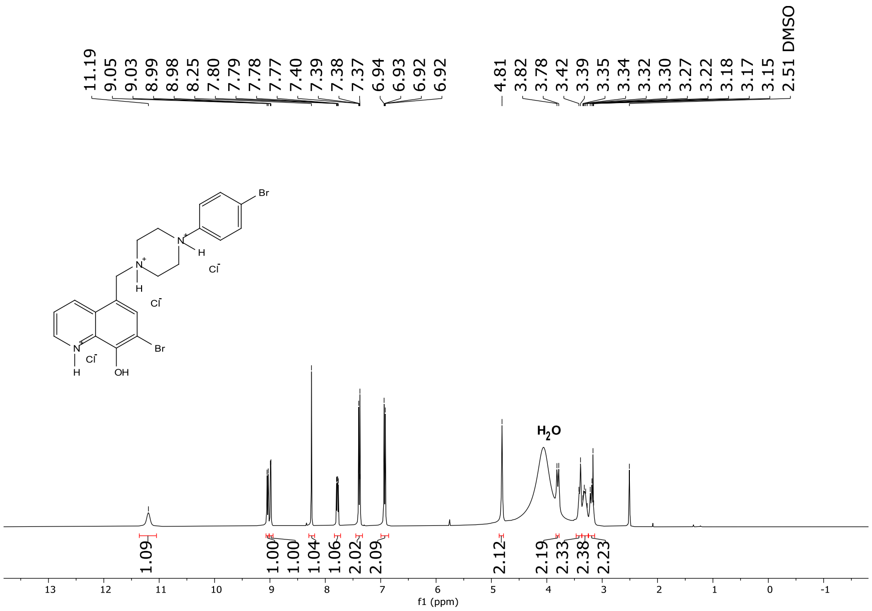

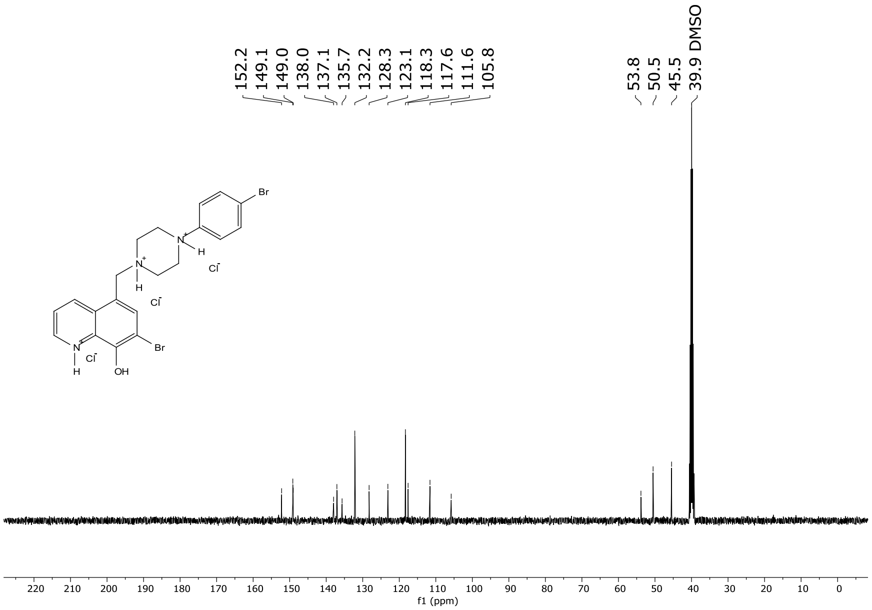

#### HRMS Spectra

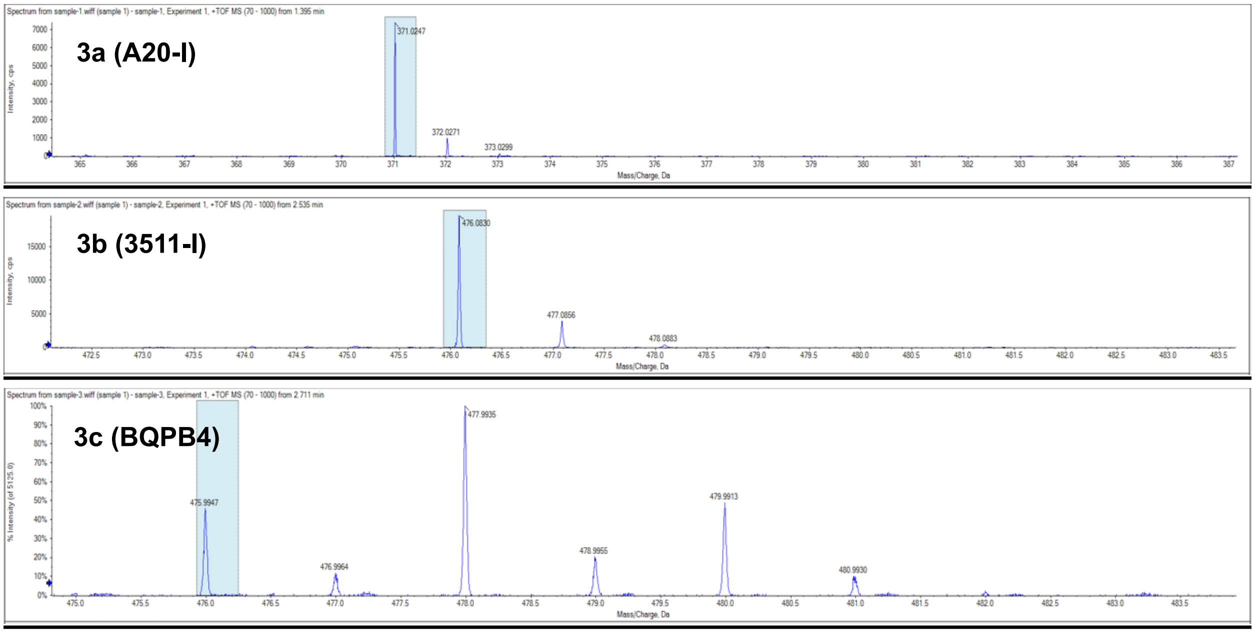

### Supplementary Computer Code Information

**Octet_DimerTetramer.py**
A Python algorithm for analyzing biolayer interferometry (BLI) data. It separates two co-existing binding species in Ephrin × Eph receptor interactions: a fast-dissociating dimer and a slow-dissociating tetramer. The script imports Octet sensorgram data and fits the dissociation phase with a global biexponential model. The fast component is assigned to the dimer and the slow component to the tetramer. It then fits the association phase with a two-phase model across all analyte concentrations to get association rate constants for each species. From these fits it calculates dissociation rates, equilibrium dissociation constants ($K_D$), and half-lives for the dimer and tetramer. Quality-control checks exclude poor sensors and fits, and flag tetramer rates measured over short dissociation windows as approximate. Outputs include annotated fit plots (PDF) and summary tables of kinetic constants. The goodness-of-fit is quantified by the coefficient of determination ($R^2$), and a sigmoidal dose–response plot with $R^2$ values automatically generated. Accuracy is lower when the slow phase is small or the dissociation window is short. Written in Python using open-source libraries (NumPy, SciPy, pandas, and Matplotlib), the script provides a transparent and reproducible framework for dimer–tetramer kinetic analysis of Octet data.

**Find50.py**
A Python algorithm for estimating half-maximal inhibitory concentration ($IC_{50}$) values from dose-dependent bioassay data. The script is optimized for kinetic binding data obtained from Octet instruments but can also perform generalized sigmoidal regression for any dataset consisting of paired concentration–response points. When used with Octet BLI data, the script imports Excel files containing time series measurements, isolates association and dissociation phases, and computes both standard $IC_{50}$ values and time-integrated AUC $IC_{50}$ information for analyses. Input data are read from user-specified Excel sheets, where signal values at defined time points (e.g., 75 sec association and 130 sec dissociation) are extracted and processed to generate concentration–response curves for each sample or compound. Responses are normalized to a 0–100 scale, outliers are filtered to ensure monotonic signal decay, and a three-parameter logistic model is fit via nonlinear least squares (scipy.optimize.leastsq) to determine the inflection point corresponding to the $IC_{50}$. The goodness-of-fit is quantified by the coefficient of determination ($R^2$), and a sigmoidal dose–response plot with annotated $IC_{50}$ and $R^2$ values is automatically generated. When applied to Octet datasets, Find50.py outputs $IC_{50}$ values for both association and dissociation phases, as well as baseline-corrected (relative) fits for each, enabling users to distinguish kinetic binding affinities across conditions. The program also supports simpler single-point (concentration, signal) datasets for non- Octet experiments, allowing it to function as a general tool for dose–response analysis in medicinal chemistry or pharmacology. Outputs include printed $IC_{50}$ estimates, $R^2$ values, and optional graphical plots for each fit. While robust for monotonic data, accuracy may decrease with noisy or incomplete concentration–response curves, and smoothing or re-parameterization may be required for datasets exhibiting non-

sigmoidal trends. Written entirely in Python using open-source libraries (NumPy, SciPy, pandas, and Matplotlib), Find50.py provides a transparent, reproducible, and easily modifiable framework for rapid $IC_{50}$ determination across biochemical and biophysical assay formats.

**EasyCellCounting.py**

A Python algorithm developed for rapid, semi-automated quantification of fluorescently labeled neurons in confocal microscopy images. Designed for neuroscientists and biomedical researchers, it enables efficient identification and counting of fluorescent cells—such as tdTomato-positive neurons—across large datasets with minimal coding experience. The program accepts both full-resolution TIFF images and corresponding downsampled preview images (PNG or TIFF) for user-guided region-of-interest (ROI) selection, rotation, and brightness threshold adjustment. After selecting ROIs, the algorithm automatically maps these to the full-resolution images and applies red-channel intensity filtering to isolate fluorescent pixels above user-defined brightness thresholds. Cell boundaries are then detected using OpenCV contour analysis, with contours between 40 and 900 pixels identified as single cells (outlined in green) and larger regions classified as clusters (outlined in blue), where cluster size is estimated by dividing total cluster area by the 99th percentile of single-cell area. Minimum cell size and cluster size can be adjusted by the user. Detected cells are separated into left and right subregions for hemispheric quantification, and results are automatically inserted into a pre-formatted Excel workbook, producing color-coded tables of left and right cell counts per sample and region. EasyCellCounting.py outputs both annotated images and an Excel file summarizing counts by animal ID and region, supporting streamlined, reproducible cell quantification across experiments. While highly efficient for moderately dense fluorescence signals, performance may decrease with heavy clustering or overlapping cells, and brightness or area thresholds may require manual tuning. Written entirely in Python using open-source libraries (OpenCV, PIL, NumPy, pandas, and openpyxl), EasyCellCounting offers a transparent, customizable, and reproducible framework for automated cell counting in fluorescence microscopy.

**Code Availability Statement:**

The full source codes for **Octet_DimerTetramer.py**, **Find50.py**, and **EasyCellCounting.py** along with usage documentation, is freely available on GitHub at:

https://github.com/ack41704

Scripts are distributed under an open-source license and can be modified or extended for specific research applications.

### Supplementary Tables

**Table S1:** Global Fit Kinetic Binding Information and Equilibrium Dissociation Constants ($K_D$) for Indicated Ephrin-Eph Interaction Pairs Using Octet Studio Analysis 13.1 software 1:1 and 2:1 Binding Models.

**Table S2:** Global Fit Kinetic Binding Information and Equilibrium Dissociation Constants ($K_D$) and Dissociation Half-Lifes for Indicated Ephrin-Eph Interaction Pairs Using In-House Developed Dimer:Tetramer Binding Model.

**Table S3:** High-Throughput Screen Identified 8-Hydroxyquinoline Compounds and $IC_{50}$ Information.

**Table S4:** MedChem Synthesized 8-Hydroxyquinoline Compounds and $IC_{50}$ Information.

**Table S5:** Statistics for Figures 3 and 4.

**Table S6:** Statistics for Figure 5.

**Table S7:** Statistics for Figure 6.

**Table S8:** Statistics for Figure 7.

### Table S1: Global Fit Kinetic Binding Information and Equilibrium Dissociation Constants (KD) for Indicated Ephrin-Eph Interaction Pairs Using Octet Studio Analysis 13.1 software 1:1 and 2:1 Binding Models.

| Interaction | Octet Experiment | Global Fit # | KD | KD Error | ka (1/Ms) | ka Error | kdis (1/s) | kdis Error | $R^2$ | $KD_1$ | $KD_2$ | $KD_1$ Error | $KD_2$ Error | $ka_1$ (1/Ms) | $ka_2$ | $ka_1$ Error | $ka_2$ Error | $kdis_1$ (1/s) | $kdis_2$ | $kdis_1$ Error | $kdis_2$ Error | $R^2$ |
|---|---|---|---|---|---|---|---|---|---|---|---|---|---|---|---|---|---|---|---|---|---|---|
| **EphrinB1-Fc + EphB1-His** | 2026.48a | 24a | 2.600E-12 | 3.359E-14 | 1.629E+06 | 9.672E+03 | 4.224E-06 | 4.864E-08 | 0.9575 | 1.000E-12 | 1.790E-11 | 1.000E-12 | 1.735E-12 | 1.833E+06 | 1.358E+06 | 3.333E+04 | 3.910E+04 | 1.000E-07 | 2.434E-05 | 1.000E-07 | 2.250E-06 | 0.9571 |
| | 2026.48b | 24b | 6.900E-12 | 3.373E-14 | 7.707E+05 | 2.463E+03 | 5.277E-06 | 1.978E-08 | 0.9895 | 1.000E-12 | 1.540E-11 | 1.000E-12 | 1.835E-12 | 6.697E+05 | 8.203E+05 | 6.038E+03 | 1.887E+04 | 1.000E-07 | 1.264E-05 | 1.000E-07 | 1.477E-06 | 0.9902 |
| | | | **4.750E-12** | **3.366E-14** | **1.200E+06** | **6.068E+03** | **4.751E-06** | **3.421E-08** | **0.9735** | **1.000E-12** | **1.665E-11** | **1.000E-12** | **1.785E-12** | **1.251E+06** | **1.089E+06** | **1.968E+04** | **2.899E+04** | **1.000E-07** | **1.849E-05** | **1.000E-07** | **1.864E-06** | **0.9737** |
| **EphrinB1-Fc + EphB2-His** | 2026.49 | 25 | 2.090E-11 | 1.163E-13 | 1.179E+06 | 6.362E+03 | 2.465E-05 | 3.336E-08 | 0.9751 | 1.000E-12 | 1.230E-07 | 1.000E-12 | 2.392E-12 | 4.591E+05 | 5.616E+05 | 8.016E+03 | 9.177E+03 | 1.000E-07 | 6.911E-05 | 1.000E-07 | 7.267E-07 | 0.9776 |
| | | | **2.090E-11** | **1.163E-13** | **1.179E+06** | **6.362E+03** | **2.465E-05** | **3.336E-08** | **0.9751** | **1.000E-12** | **1.230E-07** | **1.000E-12** | **2.392E-12** | **4.591E+05** | **5.616E+05** | **8.016E+03** | **9.177E+03** | **1.000E-07** | **6.911E-05** | **1.000E-07** | **7.267E-07** | **0.9776** |
| **EphrinB1-Fc + EphA4-His** | 2026.50 | 26 | 2.250E-10 | 3.565E-12 | 1.962E+05 | 3.072E+03 | 4.413E-05 | 1.095E-07 | 0.7128 | 1.820E-09 | 7.510E-08 | 1.188E-11 | 3.480E-10 | 6.568E+03 | 8.125E+04 | 3.947E+01 | 3.574E+02 | 1.194E-05 | 6.105E-03 | 3.073E-08 | 8.844E-06 | 0.9837 |
| | | | **2.250E-10** | **3.565E-12** | **1.962E+05** | **3.072E+03** | **4.413E-05** | **1.095E-07** | **0.7128** | **1.820E-09** | **7.510E-08** | **1.188E-11** | **3.480E-10** | **6.568E+03** | **8.125E+04** | **3.947E+01** | **3.574E+02** | **1.194E-05** | **6.105E-03** | **3.073E-08** | **8.844E-06** | **0.9837** |
| **EphrinB2-Fc + EphB1-His** | 2026.42 | 19 | 1.000E-12 | 1.000E-12 | 8.628E+05 | 3.029E+03 | 1.000E-07 | 1.000E-07 | 0.9813 | 1.000E-12 | 3.719E-07 | 1.000E-12 | 1.744E-07 | 5.218E+04 | 3.150E+05 | 5.657E+02 | 1.473E+05 | 1.000E-07 | 1.171E-01 | 1.000E-07 | 3.705E-03 | 0.9462 |
| | 2026.53 | 29 | 3.080E-11 | 2.875E-13 | 1.185E+06 | 9.841E+03 | 3.647E-05 | 1.565E-07 | 0.9792 | 8.500E-12 | 2.330E-08 | 8.163E-14 | 3.239E-10 | 9.561E+05 | 1.343E+05 | 3.139E+03 | 1.801E+03 | 8.164E-06 | 3.126E-03 | 7.330E-08 | 1.165E-05 | 0.9979 |
| | 2026.63a | 35 | 1.500E-12 | 8.746E-14 | 7.397E+05 | 2.725E+03 | 1.125E-06 | 6.457E-08 | 0.9964 | 1.000E-12 | 1.810E-11 | 1.000E-12 | 1.479E-12 | 1.157E+05 | 4.129E+05 | 6.464E+03 | 1.417E+04 | 1.000E-07 | 1.005E-05 | 1.000E-07 | 5.039E-07 | 0.9934 |
| | | | **1.110E-11** | **4.583E-13** | **9.292E+05** | **5.198E+03** | **1.257E-05** | **1.070E-07** | **0.9856** | **3.500E-12** | **1.317E-07** | **6.939E-13** | **5.824E-08** | **3.747E+05** | **2.874E+05** | **3.390E+03** | **5.442E+04** | **2.788E-06** | **4.008E-02** | **9.110E-08** | **1.239E-03** | **0.9792** |
| **EphrinB2-Fc + EphB2-His** | 2026.18 | 4 | 4.850E-11 | 3.507E-13 | 5.419E+05 | 3.557E+03 | 2.628E-05 | 7.979E-08 | 0.6102 | 1.000E-12 | 1.370E-10 | 1.000E-12 | 7.749E-12 | 5.194E+05 | 6.522E+05 | 1.574E+04 | 3.474E+04 | 1.000E-07 | 8.909E-05 | 1.000E-07 | 1.738E-06 | 0.6637 |
| | 2026.40 | 18 | 1.000E-12 | 1.000E-12 | 5.261E+05 | 7.995E+02 | 1.000E-07 | 1.000E-07 | 0.9923 | 1.000E-12 | 6.230E-07 | 1.000E-12 | 3.423E-06 | 3.701E+05 | 7.023E+05 | 1.171E+03 | 3.860E+06 | 1.000E-07 | 4.371E-01 | 1.000E-07 | 6.934E-02 | 0.9858 |
| | 2026.51 | 27 | 5.030E-11 | 3.794E-13 | 7.889E+05 | 5.367E+03 | 3.965E-05 | 1.298E-07 | 0.9382 | 1.020E-11 | 5.800E-09 | 1.347E-13 | 5.599E-11 | 3.428E+05 | 1.596E+06 | 1.045E+03 | 1.500E+04 | 3.491E-06 | 9.199E-03 | 4.493E-08 | 2.267E-05 | 0.9952 |
| | 2026.80 | 45 | 6.712E-10 | 1.358E-11 | 4.027E+05 | 7.808E+03 | 2.703E-04 | 1.558E-06 | 0.9631 | 1.000E-12 | 1.254E-09 | 1.000E-12 | 3.915E-11 | 3.618E+04 | 8.129E+05 | 5.626E+02 | 2.307E+04 | 1.000E-07 | 1.019E-03 | 1.000E-07 | 1.326E-05 | 0.9779 |
| | | | **1.928E-10** | **3.828E-12** | **5.649E+05** | **4.383E+03** | **8.408E-05** | **4.669E-07** | **0.8760** | **3.300E-12** | **1.575E-07** | **7.837E-13** | **8.558E-07** | **3.171E+05** | **9.409E+05** | **4.630E+03** | **9.832E+05** | **9.478E-07** | **1.119E-01** | **8.623E-08** | **1.734E-02** | **0.9057** |
| **EphrinB2-Fc + EphB4-His** | 2026.37 | 17 | 4.300E-12 | 1.075E-14 | 1.027E+06 | 1.323E+03 | 4.392E-06 | 9.479E-09 | 0.9959 | 1.000E-12 | 5.500E-12 | 1.000E-12 | 3.202E-14 | 2.648E+05 | 1.095E+06 | 1.437E+04 | 4.501E+03 | 1.000E-07 | 6.032E-06 | 1.000E-07 | 2.477E-08 | 0.9955 |
| | 2026.52 | 28 | 4.340E-11 | 4.812E-13 | 1.064E+06 | 1.069E+04 | 4.622E-05 | 2.162E-07 | 0.9502 | 1.000E-12 | 5.180E-10 | 1.000E-12 | 1.353E-11 | 9.501E+04 | 5.247E+05 | 2.277E+03 | 1.221E+04 | 1.000E-07 | 2.716E-04 | 1.000E-07 | 3.232E-06 | 0.9540 |
| | 2026.65a | 37 | 1.000E-12 | 1.000E-12 | 1.095E+06 | 3.174E+03 | 1.000E-07 | 1.000E-07 | 0.9937 | 1.000E-12 | 4.600E-10 | 1.000E-12 | 1.682E-10 | 5.981E+04 | 6.646E+04 | 1.048E+04 | 1.009E+04 | 1.000E-07 | 3.055E-05 | 1.000E-07 | 1.017E-05 | 0.9580 |
| | | | **1.623E-11** | **4.973E-13** | **1.062E+06** | **5.062E+03** | **1.690E-05** | **1.086E-07** | **0.9799** | **1.000E-12** | **3.278E-10** | **1.000E-12** | **6.059E-11** | **1.399E+05** | **5.621E+05** | **9.042E+03** | **8.934E+03** | **1.000E-07** | **1.027E-04** | **1.000E-07** | **4.476E-06** | **0.9692** |
| **EphrinB2-Fc + EphA4-His** | 2026.44 | 20 | 2.620E-10 | 1.661E-12 | 7.512E+04 | 4.432E+02 | 1.967E-05 | 4.579E-08 | 0.9431 | 1.200E-12 | 8.100E-10 | 5.777E-12 | 1.486E-11 | 3.966E+04 | 8.030E+04 | 5.286E+02 | 1.087E+03 | 1.000E-07 | 6.508E-05 | 1.000E-07 | 8.055E-07 | 0.9616 |
| | 2026.66b | 38a | 8.260E-10 | 9.275E-12 | 3.231E+04 | 2.516E+02 | 2.670E-05 | 2.158E-07 | 0.9797 | 1.000E-12 | 1.900E-09 | 1.000E-12 | 1.717E-10 | 2.641E+04 | 4.755E+04 | 6.685E+02 | 2.079E+03 | 1.000E-07 | 8.832E-05 | 1.000E-07 | 7.191E-06 | 0.9817 |
| | 2026.66c | 38b | 4.980E-10 | 6.595E-12 | 9.366E+04 | 1.112E+03 | 4.666E-05 | 2.731E-07 | 0.9715 | 1.900E-09 | 8.600E-08 | 1.923E-11 | 2.174E-09 | 1.403E+04 | 8.482E+04 | 1.242E+02 | 2.061E+03 | 2.623E-05 | 7.316E-03 | 1.378E-07 | 4.904E-05 | 0.9952 |
| | | | **5.287E-10** | **5.844E-12** | **6.703E+04** | **6.023E+02** | **3.101E-05** | **1.782E-07** | **0.9648** | **6.341E-10** | **2.957E-08** | **8.669E-12** | **7.869E-10** | **2.670E+04** | **7.089E+04** | **4.404E+02** | **1.742E+03** | **8.810E-06** | **2.490E-03** | **1.126E-07** | **1.901E-05** | **0.9795** |
| **EphrinB3-Fc + EphB1-His** | 2026.29 | 13 | 6.138E-11 | 8.052E-13 | 5.775E+05 | 7.386E+03 | 3.544E-05 | 1.036E-07 | 0.6146 | 3.060E-11 | 1.621E-07 | 1.559E-13 | 8.133E-10 | 3.614E+05 | 3.006E+04 | 1.533E+03 | 1.448E+02 | 1.106E-05 | 4.874E-03 | 3.122E-08 | 6.841E-06 | 0.9735 |
| | 2026.46 | 22 | 1.090E-11 | 1.031E-13 | 1.185E+06 | 1.041E+04 | 1.291E-05 | 4.503E-08 | 0.9244 | 1.000E-12 | 1.200E-10 | 1.000E-12 | 3.326E-12 | 2.200E+05 | 5.778E+05 | 4.495E+03 | 1.255E+04 | 1.000E-07 | 6.905E-05 | 1.000E-07 | 1.201E-06 | 0.9112 |
| | | | **3.614E-11** | **4.542E-13** | **8.813E+05** | **8.898E+03** | **2.418E-05** | **7.432E-08** | **0.7695** | **1.580E-11** | **8.111E-08** | **5.780E-13** | **4.083E-10** | **2.907E+05** | **3.039E+05** | **3.014E+03** | **6.347E+03** | **5.580E-06** | **2.472E-03** | **6.561E-08** | **4.021E-06** | **0.9424** |
| **EphrinB3-Fc + EphB2-His** | 2026.21 | 6 | 1.740E-11 | 2.272E-13 | 9.333E+05 | 1.195E+04 | 1.625E-05 | 4.093E-08 | 0.9611 | 1.000E-12 | 3.540E-10 | 1.000E-12 | 9.286E-12 | 1.926E+06 | 1.969E+05 | 5.240E+04 | 4.500E+03 | 1.000E-07 | 6.962E-05 | 1.000E-07 | 9.011E-07 | 0.9651 |
| | 2026.47a | 23a | 1.040E-11 | 1.686E-13 | 1.365E+06 | 2.146E+04 | 1.421E-05 | 5.514E-08 | 0.9530 | 1.000E-12 | 1.680E-10 | 1.000E-12 | 1.332E-11 | 1.553E+05 | 1.736E+05 | 1.224E+04 | 8.960E+03 | 1.000E-07 | 2.922E-05 | 1.000E-07 | 1.752E-06 | 0.9506 |
| | 2026.47b | 23b | 5.100E-11 | 1.142E-12 | 1.068E+06 | 2.386E+04 | 5.447E-05 | 8.811E-08 | 0.9081 | 3.220E-11 | 9.340E-08 | 1.704E-13 | 9.334E-08 | 6.861E+05 | 1.593E+03 | 3.565E+03 | 4.910E+01 | 2.208E-05 | 4.821E-03 | 2.229E-08 | 4.865E-06 | 0.9959 |
| | | | **2.627E-11** | **5.126E-13** | **1.122E+06** | **1.909E+04** | **2.831E-05** | **6.139E-08** | **0.9407** | **1.140E-11** | **3.131E-08** | **7.235E-13** | **3.112E-08** | **9.225E+05** | **1.240E+05** | **2.274E+04** | **4.503E+03** | **7.427E-06** | **1.640E-03** | **7.410E-08** | **2.506E-06** | **0.9705** |
| **EphrinB3-Fc + EphA4-His** | 2026.45 | 21 | 1.000E-11 | 7.043E-13 | 1.853E+05 | 1.233E+03 | 1.852E-05 | 4.269E-08 | 0.9560 | 1.100E-12 | 7.810E-10 | 5.221E-12 | 1.506E-11 | 4.411E+04 | 1.147E+05 | 5.516E+02 | 1.936E+03 | 1.000E-07 | 8.959E-05 | 1.000E-07 | 8.355E-07 | 0.9612 |
| | | | **1.000E-11** | **7.043E-13** | **1.853E+05** | **1.233E+03** | **1.852E-05** | **4.269E-08** | **0.9560** | **1.100E-12** | **7.810E-10** | **5.221E-12** | **1.506E-11** | **4.411E+04** | **1.147E+05** | **5.516E+02** | **1.936E+03** | **1.000E-07** | **8.959E-05** | **1.000E-07** | **8.355E-07** | **0.9612** |

If Octet Experiment number is underlined, the data are from using AHC2 biosensors, if not underlined AHC biosensors were used.

Human EphrinB1-Fc is a 49.5 KDa protein containing extracellular amino acids Leu28-Ser236 with human Fc at the C-terminus (from R&D Systems).
Human EphrinB2-Fc is a 48.8 KDa protein containing extracellular amino acids Met1-Ala229 with human Fc at the C-terminus (from R&D Systems).
Human EphrinB3-Fc is a 48.3 KDa protein containing extracellular amino acids Leu28-Ser224 with human Fc at the C-terminus (from R&D Systems).

Human EphB1-His is a 60 KDa protein containing extracellular amino acids Met1-Pro540 with a polyhistidine tag at the C-terminus (from Sino Biological).
Human EphB2-His is a 59.7 KDa protein containing extracellular amino acids Met1-Leu543 with a polyhistidine tag at the C-terminus (from Sino Biological).
Human EphB4-His is a 58.5 KDa protein containing extracellular amino acids Met1-Ala539 with a polyhistidine tag at the C-terminus (from Sino Biological).
Human EphA4-His is a 60 KDa protein containing extracellular amino acids Met1-Thr547 with a polyhistidine tag at the C-terminus (from Sino Biological).

#### Table S2: Global Fit Kinetic Binding Information and Equilibrium Dissociation Constants (KD) and Dissociation Half-Lifes for Indicated Ephrin-Eph Interaction Pairs Using In-House Developed Dimer:Tetramer Binding Model.

| Interaction | Octet Experiment | Global Fit # | Buffer | Association (min) | Dissociation (h) | Ka1 tetramer (1/Ms) | Kdis1 tetramer (1/s) | KD1 tetramer (nM) | Ka2 dimer (1/Ms) | Kdis2 dimer (1/s) | KD2 dimer (nM) | Tetramer half-life (days) | Dimer half-life (min) | R² association, top conc | R² association, all curves | R² dissociation |
|---|---|---|---|---|---|---|---|---|---|---|---|---|---|---|---|---|
| **EphrinB1-Fc + EphB1-His** | 2026.48a | 24a | PBST | 13.1 | 4.00 | 1.252E+05 | 3.401E-06 | 0.0272 | 3.656E+05 | 5.692E-03 | 15.571 | 2.36 | 2.0 | 0.9956 | 0.9991 | 0.9914 |
| | 2026.48b | 24b | PBST | 13.3 | 4.00 | 2.424E+04 | 6.880E-07 | 0.0284 | 5.721E+05 | 9.835E-03 | 17.190 | 11.66 | 1.2 | 0.9984 | 0.9997 | 0.9968 |
| | | | | | | **7.473E+04** | **2.044E-06** | **2.777E-02** | **4.688E+05** | **7.763E-03** | **1.638E+01** | **7.010E+00** | **1.602E+00** | **9.970E-01** | **9.994E-01** | **9.941E-01** |
| **EphrinB1-Fc + EphB2-His** | 2026.49 | 25 | PBST | 13.2 | 4.00 | 2.082E+04 | 1.698E-05 | 0.8154 | 8.856E+05 | 7.078E-03 | 7.993 | 0.47 | 1.6 | 0.9981 | 0.9992 | 0.9938 |
| | | | | | | **2.082E+04** | **1.698E-05** | **8.154E-01** | **8.856E+05** | **7.078E-03** | **7.993E+00** | **4.726E-01** | **1.632E+00** | **9.981E-01** | **9.992E-01** | **9.938E-01** |
| **EphrinB1-Fc + EphA4-His** | 2026.50 | 26 | PBST | 13.3 | 4.00 | 6.832E+03 | 1.923E-05 | 2.8146 | 1.427E+05 | 7.995E-03 | 56.048 | 0.42 | 1.4 | 0.9954 | 0.9993 | 0.9824 |
| **EphrinB2-Fc + EphB1-His** | 2026.42 | 19 | | | | | | | | | | | | | | |
| | 2026.53 | 29 | PBST | 13.3 | 2.00 | 7.589E+03 | 9.887E-06 | 1.3029 | 7.200E+05 | 3.578E-03 | 4.970 | 0.81 | 3.2 | 0.9933 | 0.9992 | 0.9974 |
| | 2026.63a | 35 | PBST | 13.3 | 2.00 | 7.155E+03 | — | — | 6.161E+05 | 8.224E-03 | 13.348 | — | 1.4 | 0.9888 | 0.9971 | 0.9906 |
| | | | | | | **7.372E+03** | **9.887E-06** | **1.303E+00** | **6.681E+05** | **5.901E-03** | **9.159E+00** | **8.114E-01** | **2.317E+00** | **9.911E-01** | **9.982E-01** | **9.940E-01** |
| **EphrinB2-Fc + EphB2-His** | 2026.18 | 4 | PBST | 13.4 | 3.00 | 1.095E+04 | 1.346E-05 | 1.2288 | 6.839E+05 | 6.863E-03 | 10.035 | 0.60 | 1.7 | 0.9966 | 0.9999 | 0.9987 |
| | 2026.40 | 18 | PBST | 119.3 | 2.23 | — | 6.935E-06 | — | 1.405E+06 | — | — | 1.16 | — | 0.5982 | 0.9951 | 1.0000 |
| | 2026.51 | 27 | PBST | 13.3 | 2.00 | 8.378E+03 | 6.025E-06 | 0.7191 | 8.795E+05 | 1.189E-02 | 13.522 | 1.33 | 1.0 | 0.9980 | 0.9996 | 0.9921 |
| | 2026.64c | 36c | PBST | 13.3 | 2.00 | 8.721E+03 | 3.822E-07 | 0.0438 | 9.201E+05 | 3.371E-03 | 3.664 | 20.99 | 3.4 | 0.9975 | 0.9975 | 0.9974 |
| | | | | | | **8.550E+03** | **3.203E-06** | **3.815E-01** | **8.998E+05** | **7.632E-03** | **8.593E+00** | **1.116E+01** | **2.199E+00** | **9.977E-01** | **9.986E-01** | **9.947E-01** |
| **EphrinB2-Fc + EphB4-His** | 2026.37 | 17 | PBST | 13.2 | 4.00 | 2.596E+04 | 2.621E-06 | 0.1010 | 1.005E+06 | 4.364E-03 | 4.344 | 3.06 | 2.6 | 0.9927 | 0.9985 | 0.9988 |
| | 2026.52 | 28 | PBST | 13.3 | 2.00 | 8.535E+03 | 1.506E-05 | 1.7648 | 9.678E+05 | 1.953E-03 | 2.018 | 0.53 | 5.9 | 0.9913 | 0.9990 | 0.9977 |
| | 2026.65a | 37 | PBST | 13.4 | 2.00 | 1.785E+04 | — | — | 9.330E+05 | 9.859E-03 | 10.568 | — | 1.2 | 0.9883 | 0.9935 | 0.9717 |
| | | | | | | **1.319E+04** | **1.506E-05** | **1.765E+00** | **9.504E+05** | **5.906E-03** | **6.293E+00** | **5.326E-01** | **3.543E+00** | **9.898E-01** | **9.963E-01** | **9.847E-01** |
| **EphrinB2-Fc + EphA4-His** | 2026.44 | 20 | PBST | 13.3 | 4.00 | 6.818E+03 | 1.441E-05 | 2.1141 | 1.498E+05 | 3.809E-03 | 25.429 | 0.56 | 3.0 | 0.9956 | 0.9984 | 0.9883 |
| | 2026.66b | 38 | PBST | 13.3 | 2.00 | 1.585E+04 | 1.813E-05 | 1.1440 | 2.257E+05 | 7.144E-03 | 31.659 | 0.44 | 1.6 | 0.9995 | 0.9999 | 0.9983 |
| | 2026.66c | 38 | PBST | 13.3 | 2.00 | 1.388E+04 | 3.069E-05 | 2.2119 | 2.045E+05 | 1.310E-02 | 64.042 | 0.26 | 0.9 | 1.0000 | 0.9984 | 0.9519 |
| | | | | | | **1.486E+04** | **2.441E-05** | **1.678E+00** | **2.151E+05** | **1.012E-02** | **4.785E+01** | **3.520E-01** | **1.250E+00** | **9.997E-01** | **9.991E-01** | **9.751E-01** |
| **EphrinB3-Fc + EphB1-His** | 2026.29 | 13 | PBST | 13.3 | 4.00 | 1.722E+04 | 1.481E-05 | 0.8605 | 3.225E+05 | 7.624E-03 | 23.641 | 0.54 | 1.5 | 0.9993 | 0.9999 | 0.9861 |
| | 2026.46 | 22 | PBST | 13.3 | 4.00 | 7.047E+03 | 4.177E-06 | 0.5927 | 2.530E+05 | 1.387E-02 | 54.819 | 1.92 | 0.8 | 0.9981 | 0.9968 | 0.9888 |
| | | | | | | **7.047E+03** | **4.177E-06** | **5.927E-01** | **2.530E+05** | **1.387E-02** | **5.482E+01** | **1.921E+00** | **8.330E-01** | **9.981E-01** | **9.968E-01** | **9.888E-01** |
| **EphrinB3-Fc + EphB2-His** | 2026.21 | 6 | PBST | 13.3 | 4.00 | 9.077E+03 | 6.396E-06 | 0.7046 | 3.011E+05 | 8.281E-03 | 27.498 | 1.25 | 1.4 | 0.9975 | 0.9995 | 0.9969 |
| | 2026.47a | 23a | PBST | 13.2 | 4.00 | 9.775E+03 | 4.767E-06 | 0.4877 | 3.715E+05 | 1.251E-02 | 33.669 | 1.68 | 0.9 | 0.9978 | 0.9994 | 0.9965 |
| | 2026.47b | 23b | PBST | 13.3 | 4.00 | 9.418E+03 | 2.868E-05 | 3.0449 | 2.835E+05 | 7.767E-03 | 27.396 | 0.28 | 1.5 | 0.9991 | 0.9998 | 0.9955 |
| | | | | | | **9.418E+03** | **2.868E-05** | **3.045E+00** | **2.835E+05** | **7.767E-03** | **2.740E+01** | **2.798E-01** | **1.487E+00** | **9.991E-01** | **9.998E-01** | **9.955E-01** |
| **EphrinB3-Fc + EphA4-His** | 2026.45 | 21 | PBST | 13.3 | 4.00 | 5.337E+03 | 8.553E-06 | 1.6026 | 1.299E+05 | 5.706E-03 | 43.937 | 0.94 | 2.0 | 0.9954 | 0.9995 | 0.9913 |

#### Global Fit Equilibrium Dissociation Constants (KD) and Dissociation Half-Life for Indicated Ephrin-Eph Interaction Pairs with BSA blocking agent

| Interaction | Octet Experiment | Global Fit # | Buffer | Association (min) | Dissociation (h) | Ka1 tetramer (1/Ms) | Kdis1 tetramer (1/s) | KD1 tetramer (nM) | Ka2 dimer (1/Ms) | Kdis2 dimer (1/s) | KD2 dimer (nM) | Tetramer half-life (days) | Dimer half-life (min) | R² association, top conc | R² association, all curves | R² dissociation |
|---|---|---|---|---|---|---|---|---|---|---|---|---|---|---|---|---|
| **EphrinB2-Fc + EphB2-His** | 2026.90 | 59 | PBSTB | 19.9 | 16.00 | 5.661E+04 | 1.883E-06 | 0.0333 | 5.541E+05 | 1.541E-03 | 2.781 | 4.26 | 7.5 | 0.9985 | 0.9999 | 0.9977 |
| | 2026.94 | 64 | PBSTB | 89.9 | 15.00 | 1.691E+04 | 1.813E-06 | 0.1073 | 1.320E+05 | 1.502E-03 | 11.379 | 4.42 | 7.7 | 0.9957 | 0.9993 | 0.9993 |
| | | | | | | **3.676E+04** | **1.848E-06** | **7.026E-02** | **3.431E+05** | **1.521E-03** | **7.080E+00** | **4.342E+00** | **7.594E+00** | **9.971E-01** | **9.996E-01** | **9.985E-01** |
| **EphrinB3-Fc + EphB2-His** | 2026.96 | 65 | PBSTB | 90.0 | 15.00 | 3.259E+03 | 8.756E-07 | 0.2687 | 1.359E+05 | 1.793E-04 | 1.320 | 9.16 | 64.4 | 1.0000 | 1.0000 | 0.9999 |
| | 2026.97 | 66 | PBSTB | 89.9 | 15.00 | 2.799E+03 | 1.384E-06 | 0.4945 | 6.038E+04 | 1.638E-04 | 2.712 | 5.79 | 70.5 | 0.9998 | 1.0000 | 0.9990 |
| | | | | | | **3.029E+03** | **1.130E-06** | **3.816E-01** | **9.812E+04** | **1.715E-04** | **2.016E+00** | **7.479E+00** | **6.749E+01** | **9.999E-01** | **1.000E+00** | **9.995E-01** |

### Table S3: High-Throughput Screen Identified 8-Hydroxyquinoline Compounds and IC50 Information.

| | | | EphB1-EphrinB2 AlphaScreen HTS Results | $IC_{50}$ Values for BLI Interaction Between Immobilized Human EphrinB2 Ectodomain Binding to 25 nM Soluble Human EphB1 Ectodomain with 0.1-3.2 $\mu$M Compound (AUC analysis of 400 second association + 500 second dissociation) | | | $IC_{50}$ Values and Average Relative $K_i$ (calculated using $K_d$ of 0.28 nM) for BLI Interaction Between Immobilized Human EphrinB2 Ectodomain Binding to 25 nM Soluble Human EphB2 Ectodomain with 0.1-3.2 $\mu$M Compound (AUC analysis of 400 second association + 500 second dissociation) | | | | | |
|---|---|---|---|---|---|---|---|---|---|---|---|---|
| **Number** | **Common Name** | **Chemical Structure** | **% reduction in Alpha signal** | **EphrinB2-EphB1 $IC_{50}$ (Absolute, $\mu$M)** | **EphrinB2-EphB1 $IC_{50}$ (Relative, $\mu$M)** | | **EphrinB2-EphB2 $IC_{50}$ (Absolute, $\mu$M)** | **EphrinB2-EphB2 $IC_{50}$ (Relative, $\mu$M)** | **EphrinB2-EphB2 $K_i$ (Relative, nM)** | | **Sample Name (UTSW Barcode)** | **SMILES** |
| **1** | **G15** | 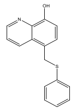 | -29% | 0.55 / 0.33 $\mu$M | 0.37 / 0.26 $\mu$M | | 0.49 / 0.26 / 0.33 $\mu$M | 0.48 / 0.23 / 0.30 $\mu$M | 3.73 nM | | **SW160569** | OC1=C2N=CC=CC2=C(CSC2=CC=CC=C2)C=C1 |
| **2** | | 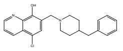 | -29% | | | | 0.44 $\mu$M | 0.38 $\mu$M | 4.21 nM | | **SW195701** | OC1=C2N=CC=CC2=C(Cl)C=C1CN1CCC(CC2=CC=CC=C2)CC1 |
| **3** | | 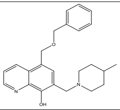 | -28% | | | | 0.45 $\mu$M | 0.40 $\mu$M | 4.43 nM | | **SW036036** | CC1CCN(CC2=CC(COCC3=CC=CC=C3)=C3C=CC=NC3=C2O)CC1 |
| **4** | | 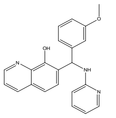 | -27% | 0.68 $\mu$M | 0.52 $\mu$M | | 0.61 $\mu$M | 0.59 $\mu$M | 6.54 nM | | **SW132689** | COC1=CC(=CC=C1)C(NC1=CC=CC=N1)C1=CC=C2C=CC=NC2=C1O |
| **5** | **3511-0013** | 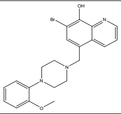 | -27% | | | | 0.39 $\mu$M | 0.34 $\mu$M | 3.77 nM | | **SW134218** | COC1=CC=CC=C1N1CCN(CC2=CC(Br)=C(O)C3=NC=CC=C23)CC1 |
| **6** | | 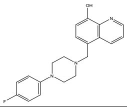 | -25% | | | | 3.24 $\mu$M | 2.46 $\mu$M | 27.25 nM | | **SW036222** | OC1=CC=C(CN2CCN(CC2)C2=CC=C(F)C=C2)C2=C1N=CC=C2 |
| **7** | | 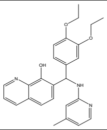 | -24% | | | | 0.48 $\mu$M | 0.40 $\mu$M | 4.43 nM | | **SW174816** | CCOC1=CC=C(C=C1OCC)C(NC1=CC(C)=CC=N1)C1=CC=C2C=CC=NC2=C1O |
| **8** | **SW022101** | 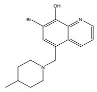 | -23% | | | | 0.85 $\mu$M | 0.63 $\mu$M | 6.98 nM | | **SW022101** | CC1CCN(CC2=CC(Br)=C(O)C3=NC=CC=C23)CC1 |
| **9** | | 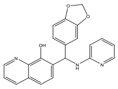 | -22% | 1.38 $\mu$M | 0.98 $\mu$M | | 0.93 $\mu$M | 0.83 $\mu$M | 9.19 nM | | **SW039486** | OC1=C2N=CC=CC2=CC=C1C(NC1=CC=CC=N1)C1=CC=C2OCOC2=C1 |
| **10** | | 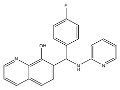 | -21% | | | | 0.41 $\mu$M | 0.34 $\mu$M | 3.77 nM | | **SW164687** | OC1=C2N=CC=CC2=CC=C1C(NC1=CC=CC=N1)C1=CC=C(F)C=C1 |
| **11** | | 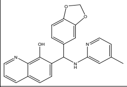 | -21% | | | | 0.34 $\mu$M | 0.25 $\mu$M | 2.77 nM | | **SW164600** | CC1=CC=NC(NC(C2=CC=C3OCOC3=C2)C2=CC=C3C=CC=NC3=C2O)=C1 |
| **12** | | 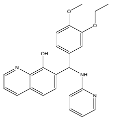 | -20.0% | 1.45 $\mu$M | 0.89 $\mu$M | | 0.98 $\mu$M | 0.72 $\mu$M | 7.98 nM | | **SW039682** | CCOC1=CC(=CC=C1OC)C(NC1=CC=CC=N1)C1=CC=C2C=CC=NC2=C1O |
| **13** | | 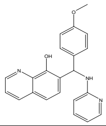 | -20.0% | 0.65 $\mu$M | 0.51 $\mu$M | | 0.26 $\mu$M | 0.20 $\mu$M | 2.22 nM | | **SW135386** | COC1=CC=C(C=C1)C(NC1=CC=CC=N1)C1=CC=C2C=CC=NC2=C1O |
| **14** | | 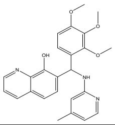 | -20.0% | | | | 0.28 $\mu$M | 0.17 $\mu$M | 1.88 nM | | **SW164761** | COC1=CC=C(C(NC2=CC(C)=CC=N2)C2=CC=C3C=CC=NC3=C2O)C(OC)=C1OC |
| **15** | | 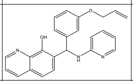 | -20.0% | 0.51 $\mu$M | 0.39 $\mu$M | | 0.52 $\mu$M | 0.49 $\mu$M | 5.43 nM | | **SW132690** | OC1=C2N=CC=CC2=CC=C1C(NC1=CC=CC=N1)C1=CC=CC(OCC=C)=C1 |
| **16** | | 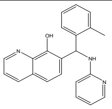 | -20.0% | 0.63 $\mu$M | 0.49 $\mu$M | | 0.56 $\mu$M | 0.54 $\mu$M | 5.98 nM | | **SW167596** | CC1=CC=CC=C1C(NC1=CC=CC=N1)C1=CC=C2C=CC=NC2=C1O |

| | | | | | | | | | | | | |
|---|---|---|---|---|---|---|---|---|---|---|---|---|
| **17** | | 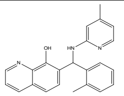 | -20.0% | | | | 0.32 μM | 0.29 μM | 3.21 nM | | **SW164644** | CC1=CC=NC(NC(C2=CC=CC=C2C)C2=CC=C3C=CC=NC3=C2O)=C1 |
| **18** | | 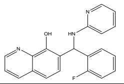 | -20.0% | 0.48 μM | 0.41 μM | | 0.54 μM | 0.51 μM | 5.65 nM | | **SW164805** | OC1=C2N=CC=CC2=CC=C1C(NC1=CC=CC=N1)C1=CC=CC=C1F |
| **19** | | 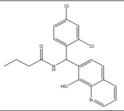 | -19% | 1.05 μM | 0.15 μM | | 0.59 μM | 0.54 μM | 5.98 nM | | **SW137279** | CCCC(=O)NC(C1=CC=C(Cl)C=C1Cl)C1=CC=C2C=CC=NC2=C1O |
| **20** | | 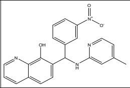 | -19% | 0.68 μM | 0.59 μM | | >3.2 μM | 0.62 μM | 6.87 nM | | **SW039339** | CC1=CC=NC(NC(C2=CC=CC(=C2)[N+]([O-])=O)C2=CC=C3C=CC=NC3=C2O)=C1 |
| **21** | | 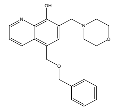 | -17% | | | | 1.95 μM | 1.70 μM | 18.83 nM | | **SW036037** | OC1=C2N=CC=CC2=C(COCC2=CC=CC=C2)C=C1CN1CCOCC1 |
| **22** | | 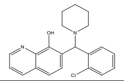 | -17% | | | | 1.02 μM | 0.74 μM | 8.20 nM | | **SW043125** | OC1=C2N=CC=CC2=CC=C1C(N1CCCCC1)C1=CC=CC=C1Cl |
| **23** | | 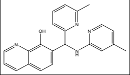 | -17% | | | | 0.71 μM | 0.60 μM | 6.65 nM | | **SW164492** | CC1=CC=NC(NC(C2=CC=CC(C)=N2)C2=CC=C3C=CC=NC3=C2O)=C1 |
| **24** | | 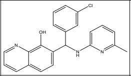 | -17% | | | | 0.32 μM | 0.28 μM | 3.10 nM | | **SW164635** | CC1=NC(NC(C2=CC=CC(Cl)=C2)C2=CC=C3C=CC=NC3=C2O)=CC=C1 |
| **25** | | 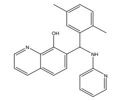 | -16% | | | | 0.48 μM | 0.37 μM | 4.10 nM | | **SW132706** | CC1=CC=C(C)C(=C1)C(NC1=CC=CC=N1)C1=CC=C2C=CC=NC2=C1O |
| **26** | | 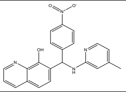 | -16% | | | | 0.40 μM | 0.28 μM | 3.10 nM | | **SW167614** | CC1=CC=NC(NC(C2=CC=C(C=C2)[N+]([O-])=O)C2=CC=C3C=CC=NC3=C2O)=C1 |
| **27** | | 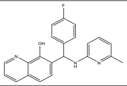 | -16% | 1.15 μM | 0.87 μM | | | | | | **SW039542** | CC1=NC(NC(C2=CC=C(F)C=C2)C2=CC=C3C=CC=NC3=C2O)=CC=C1 |
| **28** | | 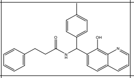 | -15% | 1.86 μM | 1.32 μM | | | | | | **SW042650** | CC1=CC=C(C=C1)C(NC(=O)CCC1=CC=CC=C1)C1=CC=C2C=CC=NC2=C1O |
| **29** | | 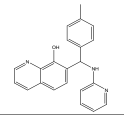 | -15% | 0.68 μM | 0.55 μM | | 0.48 μM | 0.46 μM | 5.10 nM | | **SW039492** | CC1=CC=C(C=C1)C(NC1=CC=CC=N1)C1=CC=C2C=CC=NC2=C1O |
| **30** | | 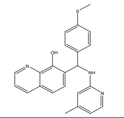 | -15% | | | | 0.58 μM | 0.51 μM | 5.65 nM | | **SW167684** | CSC1=CC=C(C=C1)C(NC1=CC(C)=CC=N1)C1=CC=C2C=CC=NC2=C1O |
| **31** | | 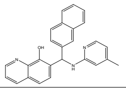 | -15% | | | | 0.31 μM | 0.28 μM | 3.10 nM | | **SW167607** | CC1=CC=NC(NC(C2=CC=C3C=CC=CC3=C2)C2=CC=C3C=CC=NC3=C2O)=C1 |
| **32** | | 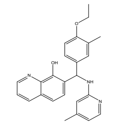 | -15% | 0.44 μM | 0.23 μM | | 0.48 μM | 0.43 μM | 4.76 nM | | **SW164571** | CCOC1=CC=C(C=C1C)C(NC1=CC(C)=CC=N1)C1=CC=C2C=CC=NC2=C1O |
| **33** | | 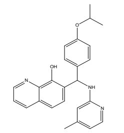 | -15% | | | | 0.44 μM | 0.37 μM | 4.10 nM | | **SW174839** | CC(C)OC1=CC=C(C=C1)C(NC1=CC(C)=CC=N1)C1=CC=C2C=CC=NC2=C1O |
| **34** | | 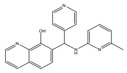 | -14% | 0.62 μM | 0.35 μM | | 0.20 μM | 0.16 μM | 1.77 nM | | **SW132682** | CC1=NC(NC(C2=CC=NC=C2)C2=CC=C3C=CC=NC3=C2O)=CC=C1 |
| **35** | | 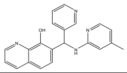 | -14% | | | | 1.17 μM | 0.93 μM | 10.30 nM | | **SW132685** | CC1=CC=NC(NC(C2=CC=CN=C2)C2=CC=C3C=CC=NC3=C2O)=C1 |

| | | | | | | | | | | | | |
|---|---|---|---|---|---|---|---|---|---|---|---|---|
| **36** | | 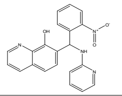 | -14% | 1.10 $\mu$M | 0.74 $\mu$M | | | | | | **SW039416** | OC1=C2N=CC=CC2=CC=C1C(NC1=CC=CC=N1)C1=CC=CC=C1[N+]([O-])=O |
| **37** | | 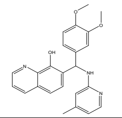 | -14% | >3.2 $\mu$M | 2.29 $\mu$M | | | | | | **SW039515** | COC1=CC=C(C=C1OC)C(NC1=CC(C)=CC=N1)C1=CC=C2C=CC=NC2=C1O |
| **38** | | 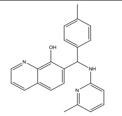 | -14% | | | | 0.81 $\mu$M | 0.60 $\mu$M | 6.65 nM | | **SW164504** | CC1=CC=C(C=C1)C(NC1=CC=CC(C)=N1)C1=CC=C2C=CC=NC2=C1O |
| **39** | | 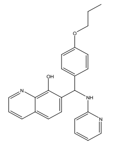 | -14% | 0.71 $\mu$M | 0.59 $\mu$M | | 0.55 $\mu$M | 0.50 $\mu$M | 5.54 nM | | **SW039607** | CCCOC1=CC=C(C=C1)C(NC1=CC=CC=N1)C1=CC=C2C=CC=NC2=C1O |
| **40** | | 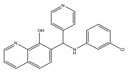 | -14% | 0.65 $\mu$M | 0.55 $\mu$M | | 0.47 $\mu$M | 0.43 $\mu$M | 4.76 nM | | **SW135270** | OC1=C2N=CC=CC2=CC=C1C(NC1=CC=CC(Cl)=C1)C1=CC=NC=C1 |
| **41** | | 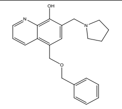 | -13% | | | | 0.5 $\mu$M | 0.4 $\mu$M | 4.43 nM | | **SW036035** | OC1=C2N=CC=CC2=C(COCC2=CC=CC=C2)C=C1CN1CCCC1 |
| **42** | | 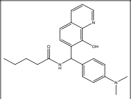 | -13% | 3.02 $\mu$M | 2.20 $\mu$M | | | | | | **SW021559** | CCCCC(=O)NC(C1=CC=C(C=C1)N(C)C)C1=CC=C2C=CC=NC2=C1O |
| **43** | | 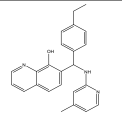 | -13% | | | | 0.52 $\mu$M | 0.49 $\mu$M | 5.43 nM | | **SW167620** | CCC1=CC=C(C=C1)C(NC1=CC(C)=CC=N1)C1=CC=C2C=CC=NC2=C1O |
| **44** | | 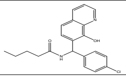 | -13% | 1.66 $\mu$M | 0.78 $\mu$M | | | | | | **SW021576** | CCCCC(=O)NC(C1=CC=C(Cl)C=C1)C1=CC=C2C=CC=NC2=C1O |
| **45** | | 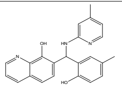 | -13% | 0.60 $\mu$M | 0.37 $\mu$M | | 0.98 $\mu$M | 0.90 $\mu$M | 9.97 nM | | **SW039544** | CC1=CC=NC(NC(C2=CC(C)=CC=C2O)C2=CC=C3C=CC=NC3=C2O)=C1 |
| **46** | | 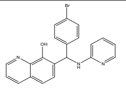 | -13% | | | | 0.48 $\mu$M | 0.44 $\mu$M | 4.87 nM | | **SW164378** | OC1=C2N=CC=CC2=CC=C1C(NC1=CC=CC=N1)C1=CC=C(Br)C=C1 |
| **47** | | 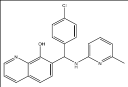 | -13% | 0.63 $\mu$M | 0.47 $\mu$M | | 0.59 $\mu$M | 0.53 $\mu$M | 5.87 nM | | **SW039592** | CC1=NC(NC(C2=CC=C(Cl)C=C2)C2=CC=C3C=CC=NC3=C2O)=CC=C1 |
| **48** | | 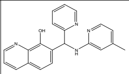 | -13% | 1.6 $\mu$M | 1.2 $\mu$M | | | | | | **SW039361** | CC1=CC=NC(NC(C2=CC=CC=N2)C2=CC=C3C=CC=NC3=C2O)=C1 |
| **49** | | 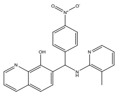 | -13% | 0.66 $\mu$M | 0.56 $\mu$M | | 0.7 $\mu$M | 0.66 $\mu$M | 7.31 nM | | **SW018955** | CC1=C(NC(C2=CC=C(C=C2)[N+]([O-])=O)C2=CC=C3C=CC=NC3=C2O)N=CC=C1 |
| **50** | | 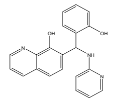 | -13% | | | | 0.28 $\mu$M | 0.25 $\mu$M | 2.77 nM | | **SW011803** | OC1=CC=CC=C1C(NC1=CC=CC=N1)C1=CC=C2C=CC=NC2=C1O |
| **51** | | 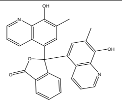 | -13% | | | | 0.53 $\mu$M | 0.49 $\mu$M | 5.43 nM | | **SW102817** | CC1=C(O)C2=C(C=CC=N2)C(=C1)C1(OC(=O)C2=C1C=CC=C2)C1=CC(C)=C(O)C2=C1C=CC=N2 |
| **52** | | 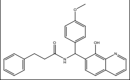 | -13% | | | | 1.02 $\mu$M | 0.96 $\mu$M | 10.63 nM | | **SW138235** | COC1=CC=C(C=C1)C(NC(=O)CCC1=CC=CC=C1)C1=CC=C2C=CC=NC2=C1O |
| **53** | | 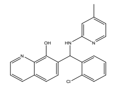 | -13% | | | | 0.59 $\mu$M | 0.54 $\mu$M | 5.98 nM | | **SW039502** | CC1=CC=NC(NC(C2=CC=CC=C2Cl)C2=CC=C3C=CC=NC3=C2O)=C1 |

| | | | | | | | | | | | | |
|---|---|---|---|---|---|---|---|---|---|---|---|---|
| **54** | | 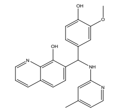 | -13% | | | | | | | | **SW132693** | COC1=CC(=CC=C1O)C(NC1=CC(C)=CC=N1)C1=CC=C2C=CC=NC2=C1O |
| **55** | | 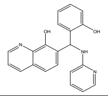 | -12% | >3.2 μM | 0.62 μM | | | | | | **SW018013** | OC1=C2N=CC=CC2=CC=C1C(NC1=CC=CC=N1)C1=CC=C(OC(F)F)C=C1 |
| **56** | | 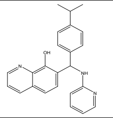 | -12% | 1.00 μM | 0.54 μM | | 1.15 μM | 1.10 μM | 12.18 nM | | **SW018899** | CC(C)C1=CC=C(C=C1)C(NC1=CC=CC=N1)C1=CC=C2C=CC=NC2=C1O |
| **57** | | 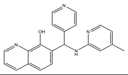 | -12% | | | | 0.66 μM | 0.60 μM | 6.65 nM | | **SW132701** | CC1=CC=NC(NC(C2=CC=NC=C2)C2=CC=C3C=CC=NC3=C2O)=C1 |
| **58** | | 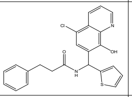 | -12% | | | | 0.55 μM | 0.51 μM | 5.65 nM | | **SW042657** | OC1=C2N=CC=CC2=C(Cl)C=C1C(NC(=O)CCC1=CC=CC=C1)C1=CC=CS1 |
| **59** | | 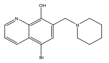 | -12% | 1.26 μM | 0.95 μM | | | | | | **SW036032** | OC1=C2N=CC=CC2=C(Br)C=C1CN1CCCCC1 |
| **60** | | 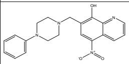 | -12% | 0.65 μM | 0.50 μM | | 0.79 μM | 0.66 μM | 7.31 nM | | **SW170388** | OC1=C(CN2CCN(CC2)C2=CC=CC=C2)C=C(C2=CC=CN=C12)[N+]([O-])=O |
| **61** | | 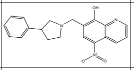 | -12% | | | | 0.49 μM | 0.44 μM | 4.87 nM | | **SW170404** | OC1=C(CN2CCC(C2)C2=CC=CC=C2)C=C(C2=CC=CN=C12)[N+]([O-])=O |
| **62** | | 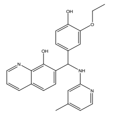 | -12% | | | | >3.2 μM | >3.2 μM | ND | | **SW039498** | CCOC1=CC(=CC=C1O)C(NC1=CC(C)=CC=N1)C1=CC=C2C=CC=NC2=C1O |
| **63** | **A20** | 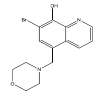 | -12% | 1.07 μM | 0.66 μM | | 0.47 μM | 0.39 μM | 4.32 nM | | **SW056428** | OC1=C(Br)C=C(CN2CCOCC2)C2=CC=CN=C12 |
| **64** | | 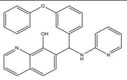 | -12% | 0.7 μM | | | 0.56 μM | 0.49 μM | 5.43 nM | | **SW047016** | OC1=C2N=CC=CC2=CC=C1C(NC1=CC=CC=N1)C1=CC=CC(OC2=CC=CC=C2)=C1 |
| **65** | | 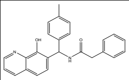 | -11% | 0.79 μM | 0.56 μM | | | | | | **SW021569** | CC1=CC=C(C=C1)C(NC(=O)CC1=CC=CC=C1)C1=CC=C2C=CC=NC2=C1O |
| **66** | | 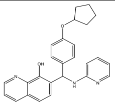 | -11% | 0.80 μM | 0.65 μM | | | | | | **SW039751** | OC1=C2N=CC=CC2=CC=C1C(NC1=CC=CC=N1)C1=CC=C(OC2CCCC2)C=C1 |
| **67** | | 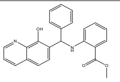 | -11% | | | | 1.82 μM | 1.55 μM | 17.17 nM | | **SW103129** | COC(=O)C1=CC=CC=C1NC(C1=CC=CC=C1)C1=CC=C2C=CC=NC2=C1O |

### Table S4: MedChem Synthesized 8-Hydroxyquinoline Compounds and IC50 Information.

| # | Chemical Structure | Common Name | IUPAC Nomenclature | Chemical Formula | Molecular Weight (g/mol) | Percent Inhibition of BLI Interaction Between Immobilized Human EphrinB2 Ectodomain Binding to 25 nM Soluble Human EphB1, EphB2, and EphB4 Ectodomains (AUC analysis of 80 second association +/- 80 second dissociation)<br>Percent inhibited = $AUC_{no\ compound} - AUC_{with\ compound} / AUC_{no\ compound}$ | | | | | |
|---|---|---|---|---|---|---|---|---|---|---|---|
| | | | | | | Inhibition During Association ONLY (80") | | | Inhibition During Association + Dissociation (160") | | |
| | | | | | | EphB1 Percent Inhibition (at 3.2 μM) | EphB2 Percent Inhibition (at 3.2 μM) | EphB4 Percent Inhibition (at 3.2 μM) | EphB1 Percent Inhibition (at 3.2 μM) | EphB2 Percent Inhibition (at 3.2 μM) | EphB4 Percent Inhibition (at 3.2 μM) |
| 3 | 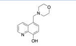 | A19 | 5-(morpholinomethyl)quinolin-8-ol | C14H16N2O2 | 244.29 | 19.0 / 18.7 / 5% | 31.8 / 12.9 / 0% | 10.7 / 0.1 / 0% | 33.3 / 25.4 / 14.7% | 42.8 / 26.2 / 10.3% | 22.0 / 8.6 / 0% |
| 4 | 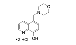 | A19 . 2 HCl | 4-((8-hydroxyquinolin-1-ium-5-yl)methyl)morpholin-4-ium chloride | C14H18Cl2N2O8 | 317.21 | | | | | | |
| 5 | 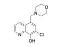 | 8009-9255 | 7-chloro-5-(morpholinomethyl)quinolin-8-ol | C14H15ClN2O2 | 278.74 | | | | | | |
| 6 | 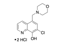 | 8009-9255 . 2 HCl | 4-((7-chloro-8-hydroxyquinolin-1-ium-5-yl)methyl)morpholin-4-ium chloride | C14H17Cl3N2O2 | 351.65 | 34.2% | 64.1% | 16.8% | 48.4% | 78.1% | 35.5% |
| 1 | 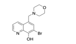 | A20 | 7-bromo-5-(morpholinomethyl)quinolin-8-ol | C14H15BrN2O2 | 323.19 | | | | | | |
| 2 | 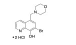 | A20 . 2 HCl | 4-((7-bromo-8-hydroxyquinolin-1-ium-5-yl)methyl)morpholin-4-ium chloride | C14H17BrCl2N2O2 | 396.11 | 33.5% | 61.0% | 20.0% | 48.7% | 76.8% | 35.9% |
| 7 | 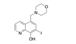 | A20-I | 7-iodo-5-(morpholinomethyl)quinolin-8-ol | C14H15IN2O2 | 370.19 | | | | | | |
| 8 | 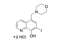 | A20-I . 2 HCl | 4-((8-hydroxy-7-iodoquinolin-1-ium-5-yl)methyl)morpholin-4-ium chloride | C14H17Cl2IN2O2 | 443.11 | 37.0 / 30.2 / 29.5 / 32.6 / 24.6 / 33.3 / 35.4 / 35.9 / 32.0 / 34.2 / 32.9 / 36.6% | 66.1 / 62.2 / 59.6 / 56.4 / 61.2 / 56.7 / 43.9 / 61.8 / 61.0 / 54.8 / 62.5 / 61.8% | 18.8 / 29.6 / 13.3 / 15.4 /9.7 /14.3 / 9.8 / 28.3 / 22.8 / 13.2 / 20.2 / 13.9% | 53.1 / 46.8 / 49.4 / 44.3 / 52.4 / 51.0 / 53.2 / 50.4 / 53.2 / 48.3 / 41.9 / 48.5% | 80.6 / 75.3 / 74.7 / 77.6 / 74.3 / 64.2 / 73.6 / 76.4 / 73.6 / 76.9 / 77.2 / 76.5% | 35.1 / 32.9 / 37.0 / 30.8 / 34.1 / 28.8 / 32.0 / 40.8 / 30.2 / 33.5 / 34.5 / 33.9% |
| 89 | 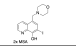 | A20-I . 2 MSA | 4-((8-hydroxy-7-iodoquinolin-1-ium-5-yl)methyl)morpholin-4-ium methanesulfonate | $C_{16}H_{23}IN_2O_8S_2$ | 562.39 | 28.4% | 55.1% | 1.7% | 46.3% | 74.6% | 25.4% |
| 9 | 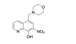 | A19-NO2 | 5-(morpholinomethyl)-7-nitroquinolin-8-ol | C14H15N3O4 | 289.29 | 22.5 / 16.1 / 15.7% | 40.4 / 17.9 / 49.2% | 0.6 / 0* / 8.2% | 36.9 / 29.8 / 22.8% | 58.0 / 38.9 / 58.0% | 10.2 / 0* / 8.7% |
| 10 | 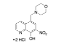 | A19-NO2 . 2 HCl | 4-((8-hydroxy-7-nitroquinolin-1-ium-5-yl)methyl)morpholin-4-ium chloride | C14H17Cl2N3O4 | 362.21 | 15.9 / 5.1 / 18.4% | 9.2 / 11.6 / 16.5% | 0.1 / 0 / 4.1% | 23.1 / 14.2 / 20.2% | 24.5 / 24.2 / 21.0% | 9.0 / 0 / 5.9% |
| 14 | 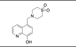 | QTM | 4-((8-hydroxyquinolin-5-yl)methyl)thiomorpholine 1,1-dioxide | C14H16N2O3S | 292.35 | | | | | | |
| 15 | 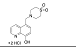 | QTM . 2 HCl | 4-((8-hydroxyquinolin-1-ium-5-yl)methyl)thiomorpholin-4-ium 1,1-dioxide chloride | C14H18Cl2N2O3S | 365.27 | 16.8% | 33.8% | 10.8% | 28.6% | 46.4% | 19.0% |
| 16 | 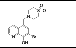 | QTM-Br | 4-((7-bromo-8-hydroxyquinolin-5-yl)methyl)thiomorpholine 1,1-dioxide | C14H15BrN2O3S | 371.25 | | | | | | |
| 17 | 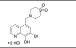 | QTM-Br . 2 HCl | 4-((7-bromo-8-hydroxyquinolin-1-ium-5-yl)methyl)thiomorpholin-4-ium 1,1-dioxide chloride | C14H17BrCl2N2O3S | 444.17 | 33.0% | 59.5% | 18.1% | 48.0% | 74.5% | 37.2% |
| 18 | 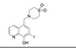 | QTM-I | 4-((8-hydroxy-7-iodoquinolin-5-yl)methyl)thiomorpholine 1,1-dioxide | C14H15IN2O2 | 418.25 | | | | | | |
| 19 | 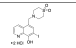 | QTM-I . 2 HCl | 4-((8-hydroxy-7-iodoquinolin-1-ium-5-yl)methyl)thiomorpholin-4-ium 1,1-dioxide chloride | C14H17Cl2IN2O3S | 491.17 | 33.9% | 53.7% | 12.9% | 50.1% | 71.4% | 36.2% |
| 20 | 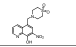 | QTM-NO2 | 4-((8-hydroxy-7-nitroquinolin-5-yl)methyl)thiomorpholine 1,1-dioxide | | 337.35 | | | | | | |
| 21 | 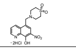 | QTM-NO2.2HCl | 4-((8-hydroxy-7-nitroquinolin-1-ium-5-yl)methyl)thiomorpholin-4-ium 1,1-dioxide chloride | | 410.27 | 24.0 / 23.8% | 36.7 / 57.0% | 0* / 4.3% | 36.0 / 28.9% | 51.7 / 69.2% | 0* / 10.6% |
| 22 | 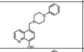 | QPPh | 5-((4-phenylpiperazin-1-yl)methyl)quinolin-8-ol | C20H21N3O | 319.41 | | | | | | |
| 23 | 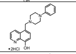 | QPPh . 2HCl | 8-hydroxy-5-((4-phenylpiperazin-1-ium-1-yl)methyl)quinolin-1-ium chloride | C20H23Cl2N3O | 392.32 | 30.8 / 16.2% | 66.0 / 38.5% | 23.1 / 12.1% | 41.2 / 24.8% | 79.1 / 49% | 26.7 / 17.9% |
| 69 | 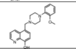 | 3511-0013-BF | 5-((4-(2-methoxyphenyl)piperazin-1-yl)methyl)quinolin-8-ol | | 349.43 | 11.9 / 24.2% | 3.6 / 0*% | 0* / 0*% | 18.7 / 45.2% | 19.2 / 15.5% | 2.2 / 0*% |
| 70 | 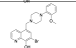 | 3511-0013 | 7-bromo-5-((4-(2-methoxyphenyl)piperazin-1-yl)methyl)quinolin-8-ol | C21H22BrN3O2 | 428.33 | 47.2% | 53.4% | 15.1% | 58.8% | 73.0% | 36.3% |
| 71 | 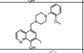 | 3511-I | 7-iodo-5-((4-(2-methoxyphenyl)piperazin-1-yl)methyl)quinolin-8-ol | | 475.33 | | | | | | |
| | 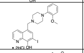 | 3511-I . 2 HCl | 7-iodo-5-((4-(2-methoxyphenyl)piperazin-1-yl)methyl)quinolin-8-ol dihydrochloride | C21H22IN3O2Cl2 | 548.35 | 39.6% | 58.7% | 17.5% | 52.3% | 73.2% | 35.6% |
| 72 | 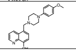 | 3511-4OMe-BF | 5-((4-(4-methoxyphenyl)piperazin-1-yl)methyl)quinolin-8-ol | | 349.43 | 30.0% | 39.4% | 19.8% | 44.5% | 60.2% | 33.2% |

| # | Common Name | Individual $IC_{50}$ Values and Average Relative $K_i$ (calculated using $K_D$ of 0.28 nM) for BLI Interaction Between Immobilized Human EphrinB2 Ectodomain Binding to 25 nM Soluble Human EphB2 Ectodomain (AUC analysis of 400 second association + 500 second dissociation) | | | $IC_{50}$ Values for BLI Interaction Between Immobilized Rat EphB1 Ectodomain Binding to 50 nM Soluble Mouse EphrinB2 Ectodomain (AUC analysis of 400 second association + 500 second dissociation) | | $IC_{50}$ Values for BLI Interaction Between Immobilized Mouse EphB2 Ectodomain Binding to 50 nM Soluble Mouse EphrinB2 Ectodomain (AUC analysis of 400 second association + 500 second dissociation) | | $IC_{50}$ Values for BLI Interaction Between Immobilized Mouse EphB2 Ectodomain Binding to 50 nM Soluble Human EphrinA5 Ectodomain (AUC analysis of 400 second association + 500 second dissociation) | | $IC_{50}$ Values for BLI Interaction Between Immobilized Human EphA3 Ectodomain Binding to 50 nM Soluble Human EphrinA5 Ectodomain (AUC analysis of 400 second association + 500 second dissociation) | | $IC_{50}$ Values for BLI Interaction Between Immobilized Human EphA3 Ectodomain Binding to 50 nM Soluble Human EphrinA1 Ectodomain (AUC analysis of 400 second association + 500 second dissociation) | |
|---|---|---|---|---|---|---|---|---|---|---|---|---|---|---|
| | | EphrinB2-EphB2 $IC_{50}$ (Absolute, μM) | EphrinB2-EphB2 $IC_{50}$ (Relative, μM) | EphrinB2-EphB2 $K_i$ (Relative, μM) | EphB1-EphrinB2 $IC_{50}$ (Absolute, μM) | EphB1-EphrinB2 $IC_{50}$ (Relative, μM) | EphB2-EphrinB2 $IC_{50}$ (Absolute, μM) | EphB2-EphrinB2 $IC_{50}$ (Relative, μM) | EphB2-EphrinA5 $IC_{50}$ (Absolute, μM) | EphB2-EphrinA5 $IC_{50}$ (Relative, μM) | EphA3-EphrinA5 $IC_{50}$ (Absolute, μM) | EphA3-EphrinA5 $IC_{50}$ (Relative, μM) | EphA3-EphrinA1 $IC_{50}$ (Absolute, μM) | EphA3-EphrinA1 $IC_{50}$ (Relative, μM) |
| 3 | A19 | >>1.6 μM | >>1.6 μM | ND | >100 μM | 3.29 μM | 9.9 μM | 0.90 μM | | | N/A | 3.22 μM | 8.24 / 6.29 μM | 6.73 / 2.39 μM |
| 4 | A19 . 2 HCl | > 6.4 μM | 2.29 μM | 25.37 nM | | | | | | | | | | |
| 5 | 8009-9255 | 2.40 μM | 1.12 μM | 12.41 nM | | | | | | | | | | |
| 6 | 8009-9255 . 2 HCl | 0.93 μM | 0.51 μM | 5.65 nM | 0.83 μM | 0.64 μM | 0.90 μM | 0.63 μM | <1 μM | <1 μM | N/A | 0.54 μM | 1.84 μM | 0.90 μM |
| 1 | A20 | | | ND | | | | | | | | | | |
| 2 | A20 . 2 HCl | 0.78 μM | 0.48 μM | 5.32 nM | 0.73 μM | 0.54 μM | 0.68 μM | 0.50 μM | <1 μM | <1 μM | N/A | 0.37 μM | 7.24 μM | 1.00 μM |
| 7 | A20-I | >1.6 μM | 0.44 μM | 4.87 nM | | | | | | | | | | |
| 8 | A20-I . 2 HCl | 1.32 / 1.00 / 1.48 μM | 0.63 / 0.58 / 0.24 μM | 5.35 nM | 0.71 / 0.51 μM | 0.51 μM / 0.45 μM | 0.47 / 0.51 μM | 0.25 / 0.51 μM | 0.42 / 0.22 μM | 0.09 / 0.08 μM | N/A | 0.36 μM | 1.20 / 1.52 μM | 0.46 μM / 1.14 μM |
| 89 | A20-I . 2 MSA | | | ND | | | | | | | | | | |
| 9 | A19-NO2 | >3.2 μM | >3.2 μM | ND | | | | | | | | | | |
| 10 | A19-NO2 . 2 HCl | >>1.6 μM | >>1.6 μM | ND | 2.48 μM | 1.99 μM | <10 μM | <10 μM | | | N/A | 4.48 μM | 14.0 μM | 6.3 μM |
| 14 | QTM | | | ND | | | | | | | | | | |
| 15 | QTM . 2 HCl | >>1.6 μM | 0.81 μM | 8.97 nM | >100 μM | 0.90 μM | 6.53 μM | 0.99 μM | | | N/A | 2.10 μM | 12.3 μM | 6.0 μM |
| 16 | QTM-Br | | | ND | | | | | | | | | | |
| 17 | QTM-Br . 2 HCl | 1.48 μM | 0.62 μM | 6.87 nM | | | | | | | | | | |
| 18 | QTM-I | | | ND | | | | | | | | | | |
| 19 | QTM-I . 2 HCl | >>1.6 μM | 0.69 μM | 7.64 nM | | | | | | | | | | |
| 20 | QTM-NO2 | >3.2 μM | 2.09 μM | 23.15 | | | | | | | | | | |
| 21 | QTM-NO2.2HCl | 1.15 / 1.95 μM | 1.00 / 0.89 μM | 10.47 nM | | | | | | | | | | |
| 22 | QPPh | >3.2 μM | ND | ND | | | | | | | | | | |
| 23 | QPPh . 2HCl | >>1.6 μM | >>1.6 μM | ND | | | | | | | | | | |
| 69 | 3511-0013-BF | >3.2 μM | 2.01 μM | 22.26 nM | | | | | | | | | | |
| 70 | 3511-0013 | >>1.6 μM | 0.17 μM | 1.88 nM | | | | | | | | | | |
| 71 | 3511-I | 0.42 / 0.53 μM | 0.38 / 0.41 μM | 4.38 nM | | | | | | | | | | |
| | 3511-I . 2 HCl | | | ND | | | | | | | | | | |
| 72 | 3511-4OMe-BF | 2.29 μM | 1.20 μM | 13.29 nM | | | | | | | | | | |

| | | | | | | | | | | | | | | | | | | | | | | | | | | | | | | | | |
|---|---|---|---|---|---|---|---|---|---|---|---|---|---|---|---|---|---|---|---|---|---|---|---|---|---|---|---|---|---|---|---|---|
| 73 | 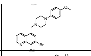 | 3511-4OMe | 7-bromo-5-((4-(4-methoxyphenyl)piperazin-1-yl)methyl)quinolin-8-ol | | 428.33 | | | | | | | | | | 2.29 µM | 1.20 µM | 13.29 nM | | | | | | | | | | | | | | | |
| 74 | 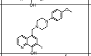 | 3511-4OMe-I | 7-iodo-5-((4-(4-methoxyphenyl)piperazin-1-yl)methyl)quinolin-8-ol | | 475.33 | | 47.2% | 50.6% | 21.0% | | 58.2% | 70.0% | 43.4% | | | | ND | | | | | | | | | | | | | | | |
| 62 | 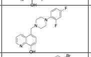 | QPDF | 5-((4-(2,4-difluorophenyl)piperazin-1-yl)methyl)quinolin-8-ol | C20H19F2N3O | 355.39 | | 17.9% | 0*% | 5.0% | | 14.7% | 0*% | 0*% | | >>1.6 µM | >>1.6 µM | NA | | | | | | | | | | | | | | | |
| 26 | 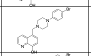 | QPB4 | 5-((4-(4-bromophenyl)piperazin-1-yl)methyl)quinolin-8-ol | C20H20BrN3O | 398.3 | | 11.2% | 35.9% | 0*% | | 36.6% | 60.1% | 0*% | | | | ND | | | | | | | | | | | | | | | |
| 27 | 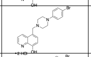 | QPB4 . 2 HCl | 5-((4-(4-bromophenyl)piperazin-1-ium-1-yl)methyl)-8-hydroxyquinolin-1-ium chloride | C20H22BrCl2N3O | 471.22 | | 24.2 / 33.8% | 16.8 / 29.2% | 11.4 / 11.4% | | 57.1 / 43.8% | 60.9 / 43.7% | 19.5 / 12.6% | | >>1.6 µM | 0.42 µM | 4.65 nM | | | | | | | | | | | | | | | |
| 28 | 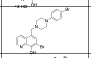 | BQPB4 | 7-bromo-5-((4-(4-bromophenyl)piperazin-1-yl)methyl)quinolin-8-ol | C20H19Br2N3O | 477.2 | | | | | | | | | | 0.36 / 0.53 / 0.59 / 0.51 µM | 0.22 / 0.37 / 0.23 / 0.32 µM | 3.16 nM | | | | | | | | | | | | | | | |
| 30 | 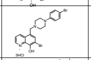 | BQPB4 . 3 HCl | 7-bromo-5-((4-(4-bromophenyl)piperazin-1-ium-1-yl)methyl)-8-hydroxyquinolin-1-ium chloride | C20H22Br2Cl3N3O | 586.52 | | 24.7 / 39.1% | 57.7 / 62.5% | 16.6 / 12.5% | | 38.6 / 53.1% | 72.9 / 78.7% | 29.3 / 36.4% | | 0.26 / 0.20 / 0.20 µM | 0.29 / 0.26 / 0.18 µM | 2.69 nM | | | | | | | | | | | | | | | |
| 32 | 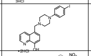 | QPI4 . 2HCl | 8-hydroxy-5-((4-(4-iodophenyl)piperazin-1-ium-1-yl)methyl)quinolin-1-ium chloride | C20H22Cl2IN3O | 518.22 | | 24.5% | 63.9% | 33.8% | | 40.5% | 78.2% | 42.5% | | >1.6 µM | 0.79 µM | 8.75 nM | | | | | | | | | | | | | | | |
| 38 | 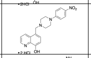 | BFNO4 . 2 HCl | 8-hydroxy-5-((4-(4-nitrophenyl)piperazin-1-ium-1-yl)methyl)quinolin-1-ium chloride | C20H22Cl2N4O3 | 437.32 | | 24.8% | 54.3% | 9.6% | | 43.2% | 72.1% | 17.5% | | 2.18 µM | 1.20 µM | 13.29 nM | | | | | | | | | | | | | | | |
| 39 | 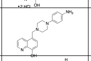 | QPA | 5-((4-(4-aminophenyl)piperazin-1-yl)methyl)quinolin-8-ol | C19H21N5O | 335.41 | | 12.7 / 0*% | 33.9 / 31.2% | 0.4 / 0*% | | 16.9 / 8.0% | 46.4 / 45.6% | 6.9 / 5.5% | | 3.09 µM | 1.70 µM | 18.83 nM | | | | | | | | | | | | | | | |
| 50 | 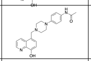 | QPA-Ac | N-(4-(4-((8-hydroxyquinolin-5-yl)methyl)piperazin-1-yl)phenyl)acetamide | C22H24N4O2 | 376.46 | | | | | | | | | | >3.2 µM | 1.95 µM | 21.60 nM | | | | | | | | | | | | | | | |
| 51 | 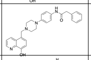 | QPA-PAc | N-(4-(4-((8-hydroxyquinolin-5-yl)methyl)piperazin-1-yl)phenyl)-2-phenylacetamide | C28H28N4O2 | 452.56 | | 52.7 / 13.8 / 3.7% | 64.8 / 15.4 / 47.8% | 18.1 / 0* / 19.4% | | 61.2 / 24.3 / 40.5% | 78.8 / 24.6 / 72.8% | 37.2 / 0.6 / 41.0% | | 1.51 µM | 1.02 µM | 11.30 nM | | | | | | | | | | | | | | | |
| 52 | 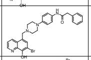 | BQPA-PAc | N-(4-(4-((7-bromo-8-hydroxyquinolin-5-yl)methyl)piperazin-1-yl)phenyl)-2-phenylacetamide | | 531.45 | | 19.0 / 12.2 / 25.4% | 6.2 / 0* / 0*% | 12.6 / 0* / 0*% | | 14.1 /15.8 / 32.1% | 19.4 / 7.6 / 0*% | 9.4 / 2.7 / 0*% | | >3.2 µM | >3.2 µM | ND | | | | | | | | | | | | | | | |
| 33 | 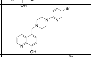 | QPP | 5-((4-(5-bromopyridin-2-yl)piperazin-1-yl)methyl)quinolin-8-ol | C19H19BrN4O | 399.29 | | | | | | | | | | 2.51 µM | 2.19 µM | 24.26 nM | | | | | | | | | | | | | | | |
| 34 | 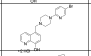 | QPP. 2 HCl | 5-((4-(5-bromopyridin-2-yl)piperazin-1-ium-1-yl)methyl)-8-hydroxyquinolin-1-ium chloride | C19H21BrCl2N4O | 472.21 | | 46.4% | 69.4% | 20.9% | | 58.6% | 81.6% | 36.9% | | >> 3.2 µM | 1.70 µM | 18.83 nM | | | | | | | | | | | | | | | |
| 83 | 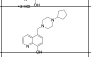 | QPcP | 5-((4-cyclopentylpiperazin-1-yl)methyl)quinolin-8-ol | C19H25N3O | 311.43 | | 21.4% | 20.9% | 0.5% | | 23.6% | 21.2%% | 0.5%% | | | | ND | | | | | | | | | | | | | | | |
| 41 | 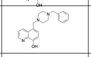 | QP-Bn | 5-((4-benzylpiperazin-1-yl)methyl)quinolin-8-ol | C21H23N3O | 333.44 | | | | | | | | | | 2.09 / >3.2 µM | NA /2.14 µM | 23.70 nM | | | | | | | | | | | | | | | |
| 48 | 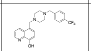 | QPCF34-Bn | 5-((4-(4-(trifluoromethyl)benzyl)piperazin-1-yl)methyl)quinolin-8-ol | C22H22F3N3O | 401.43 | | | | | | | | | | >>1.6 µM | >>1.6 µM | ND | | 15.1 µM | 2.37 µM | | 8.12 µM | 2.12 µM | | 8.24/>1 µM | 1.00/1.00 µM | | N/A | 4.33 µM | | >100 µM | 5.3 µM |
| 49 | 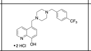 | QPCF34-Bn . 2 HCl | 8-hydroxy-5-((4-(4-(trifluoromethyl)benzyl)piperazin-1-ium-1-yl)methyl)quinolin-1-ium chloride | C22H24Cl2F3N3O | 474.35 | | 2.6 / 32.1% | 21.3 / 24.9% | 0* / 0*% | | 19.8 / 39.0% | 46.2 / 36.2% | 0* / 2.8% | | | | ND | | | | | | | | | | | | | | | |
| 42 | 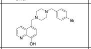 | QPB4-Bn | 5-((4-(4-bromobenzyl)piperazin-1-yl)methyl)quinolin-8-ol | C21H22BrN3O | 412.33 | | | | | | | | | | 2.51 µM | 0.93 µM | 10.30 nM | | | | | | | | | | | | | | | |
| 43 | 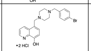 | QPB4-Bn . 2 HCl | 5-((4-(4-bromobenzyl)piperazin-1-ium-1-yl)methyl)-8-hydroxyquinolin-1-ium chloride | C21H24BrCl2N3O | 485.25 | | 19.9 / 33.5% | 37.8 / 62.3% | 0* / 22.9% | | 36.5 / 47.3% | 56.3 / 75.4% | 0* / 32.5% | | 3.09 / 3.24 µM | NA / 2.24 µM | 24.81 nM | | | | | | | | | | | | | | | |
| 44 | 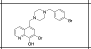 | BQPB4-Bn | 7-bromo-5-((4-(4-bromobenzyl)piperazin-1-yl)methyl)quinolin-8-ol | C21H21Br2N3O | 491.23 | | | | | | | | | | >>3.2 µM | 0.30 µM | 3.32 nM | | | | | | | | | | | | | | | |
| 45 | 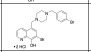 | BQPB4-Bn . 2 HCl | 7-bromo-5-((4-(4-bromobenzyl)piperazin-1-ium-1-yl)methyl)-8-hydroxyquinolin-1-ium chloride | C21H23Br2Cl2N3O | 564.14 | | 37.3% | 62.1% | 4.9% | | 49.9% | 77.1% | 28.3% | | 0.76 µM | 0.56 µM | 6.20 nM | | | | | | | | | | | | | | | |
| 46 | 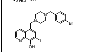 | IQPB4-Bn | 5-((4-(4-bromobenzyl)piperazin-1-yl)methyl)-7-iodoquinolin-8-ol | | 538.23 | | 51.6% | 53.0% | 18.1% | | 57.2% | 69.6% | 32.0% | | 0.63 µM | 0.52 µM | 5.76 nM | | | | | | | | | | | | | | | |
| 12 | 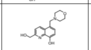 | 2OH-A19 | 2-(hydroxymethyl)-5-(morpholinomethyl)quinolin-8-ol | | 274.32 | | 17.4% | 48.1% | 11.0% | | 17.4% | 48.1% | 11.0% | | >3.2 µM | >3.2 µM | ND | | | | | | | | | | | | | | | |
| 13 | 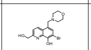 | 2OH-A20 | 7-bromo-2-(hydroxymethyl)-5-(morpholinomethyl)quinolin-8-ol | | 353.22 | | 12.4% | 50.5% | 0*% | | 29.1% | 68.4% | 7.6% | | >3.2 µM | 1.45 µM | 16.06 nM | | | | | | | | | | | | | | | |
| 57 | 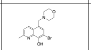 | 2-MeA20 | 7-bromo-2-methyl-5-(morpholinomethyl)quinolin-8-ol | C15H17BrN2O2 | 337.22 | | 0*% | 1.4% | 0*% | | 15.3% | 20.3% | 8.7% | | >3.2 µM | 2.24 µM | 24.81 nM | | | | | | | | | | | | | | | |
| 60 | 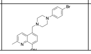 | 2-MeQPB4 | 5-((4-(4-bromophenyl)piperazin-1-yl)methyl)-2-methylquinolin-8-ol | C21H22BrN3O | 412.33 | | | | | | | | | | >>3.2 µM | >>3.2 µM | NA | | | | | | | | | | | | | | | |

| | | | | | | | | | | | | | | | | | |
|---|---|---|---|---|---|---|---|---|---|---|---|---|---|---|---|---|---|
| 61 | 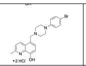 | 2-MeQPB4 . 2 HCl | 5-((4-(4-bromophenyl)piperazin-1-ium-1-yl)methyl)-8-hydroxy-2-methylquinolin-1-ium chloride | C21H24BrCl2N3O | 485.25 | | 8.1% | 0*% | 0*% | | 11.3% | 0*% | 0*% | | >>1.6 µM | >>1.6 µM | NA |
| 53 | 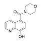 | A19-L | (8-hydroxyquinolin-5-yl)(morpholino)methanone | C14H13BrN2O3 | 337.17 | | 31.7 / 9.9% | 55.2 / 20.3% | 20.1 / 5.8% | | 38.7 / 26.3% | 65.3 / 40.6% | 16.8 / 12.5% | | 2.82 µM | 1.86 µM | 20.60 nM |
| 68 | 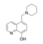 | D014-0004 | 5-(piperidin-1-ylmethyl)quinolin-8-ol | C15H18N2O | 242.32 | | 28.3% | 54.9% | 0*% | | 46.8% | 72.9% | 5.8% | | | | ND |
| 76 | 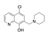 | F235-1974 | 5-chloro-7-(piperidin-1-ylmethyl)quinolin-8-ol | C15H17ClN2O | 276.76 | | 34.2 / 16.6% | 59.7 / 46.7% | 17.9 / 11.6% | | 51.5 / 43.1% | 77.8 / 71.3% | 34.5 / 29.6% | | | | ND |
| 77 | 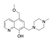 | 8002-0823 | 5-(methoxymethyl)-7-((4-methylpiperazin-1-yl)methyl)quinolin-8-ol | C17H23N3O2 | 301.39 | | 16.9% | 5.4% | 0*% | | 35.0% | 35.1% | 2.7% | | | | ND |
| 63 | 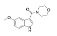 | IM | (5-methoxy-1H-indol-3-yl)(morpholino)methanone | C14H16N2O3 | 260.29 | | 1.2% | 4.3% | 0*% | | 0*% | 0*% | 0*% | | >>3.2 µM (no activity) | >>3.2 µM (no activity) | NA |
| 64 | 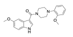 | IMP2 | (5-methoxy-1H-indol-3-yl)(4-(2-methoxyphenyl)piperazin-1-yl)methanone | C21H23N3O3 | 365.43 | | 0*% | 0*% | 0*% | | 0*% | 0*% | 0*% | | >>1.6 µM (no activity) | >>1.6 µM (no activity) | NA |
| 65 | 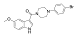 | IPB4 | (4-(4-bromophenyl)piperazin-1-yl)(5-methoxy-1H-indol-3-yl)methanone | C20H20BrN3O2 | 414.3 | | 22.2% | 5.6% | 20.2% | | 25.2% | 14.6% | 19.6% | | >>1.6 µM (no activity) | >>1.6 µM (no activity) | NA |

#### Table S5: Statistics for Figures 3 and 4.

Fig. 3D - t-test - %pEphB2

| | |
|---|---|
| Table Analyzed | phospho-EphB2 |
| Column B | pWT (A20) |
| vs. | vs. |
| Column A | pWT (Vehicle) |
| Unpaired t test | |
| P value | 0.0296 |
| P value summary | * |
| Significantly different (P < 0.05)? | Yes |
| One- or two-tailed P value? | Two-tailed |
| t, df | t=3.313, df=4 |
| How large is the effect? | |
| Mean of column A | 1.953 |
| Mean of column B | 1.073 |
| Difference between means (B - A) ± SEM | -0.8800 ± 0.2656 |
| 95% confidence interval | -1.617 to -0.1425 |
| R squared (eta squared) | 0.7329 |
| F test to compare variances | |
| F, DFn, Dfd | 41.05, 2, 2 |
| P value | 0.0476 |
| P value summary | * |
| Significantly different (P < 0.05)? | Yes |
| Data analyzed | |
| Sample size, column A | 3 |
| Sample size, column B | 3 |

Fig. 3D - t-test - pEphB2

| | |
|---|---|
| Table Analyzed | pEphB2 protein levels |
| Column B | WT (A20) |
| vs. | vs. |
| Column A | WT (Vehicle) |
| Unpaired t test | |
| P value | 0.0192 |
| P value summary | * |
| Significantly different (P < 0.05)? | Yes |
| One- or two-tailed P value? | Two-tailed |
| t, df | t=3.797, df=4 |
| How large is the effect? | |
| Mean of column A | 31.27 |
| Mean of column B | 9.327 |
| Difference between means (B - A) ± SEM | -21.94 ± 5.780 |
| 95% confidence interval | -37.99 to -5.896 |
| R squared (eta squared) | 0.7828 |
| F test to compare variances | |
| F, DFn, Dfd | 8.405, 2, 2 |
| P value | 0.2127 |
| P value summary | ns |
| Significantly different (P < 0.05)? | No |
| Data analyzed | |
| Sample size, column A | 3 |
| Sample size, column B | 3 |

Fig. 3D - t-test - tEphB2

| | |
|---|---|
| Table Analyzed | EphB2 protein levels |
| Column B | WT (A20) |
| vs. | vs. |
| Column A | WT (Vehicle) |
| Unpaired t test | |
| P value | 0.0192 |
| P value summary | * |
| Significantly different (P < 0.05)? | Yes |
| One- or two-tailed P value? | Two-tailed |
| t, df | t=3.795, df=4 |
| How large is the effect? | |
| Mean of column A | 31.21 |
| Mean of column B | 17.03 |
| Difference between means (B - A) ± SEM | -14.18 ± 3.736 |
| 95% confidence interval | -24.55 to -3.805 |
| R squared (eta squared) | 0.7826 |
| F test to compare variances | |
| F, DFn, Dfd | 1.226, 2, 2 |
| P value | 0.8986 |
| P value summary | ns |
| Significantly different (P < 0.05)? | No |
| Data analyzed | |
| Sample size, column A | 3 |
| Sample size, column B | 3 |

Fig. 3D - one-way ANOVA

| | | | | | |
|---|---|---|---|---|---|
| Table Analyzed | phospho-EphB2 | | | | |
| Data sets analyzed | A-C | | | | |
| Distribution assumption | Normal (Gaussian) | | | | |
| ANOVA summary | | | | | |
| F | 7.245 | | | | |
| P value | 0.0133 | | | | |
| P value summary | * | | | | |
| Significant diff. among means (P < 0.05)? | Yes | | | | |
| R squared | 0.6169 | | | | |
| Brown-Forsythe test | | | | | |
| F (DFn, DFd) | 3.216 (2, 9) | | | | |
| P value | 0.0884 | | | | |
| P value summary | ns | | | | |
| Are SDs significantly different (P < 0.05)? | No | | | | |
| Bartlett's test | | | | | |
| Bartlett's statistic (corrected) | | | | | |
| P value | | | | | |
| P value summary | | | | | |
| Are SDs significantly different (P < 0.05)? | | | | | |
| ANOVA table | SS | DF | MS | F (DFn, DFd) | P value |
| Treatment (between columns) | 3.097 | 2 | 1.548 | F (2, 9) = 7.245 | P=0.0133 |
| Residual (within columns) | 1.923 | 9 | 0.2137 | | |
| Total | 5.02 | 11 | | | |
| Data summary | | | | | |
| Number of treatments (columns) | 3 | | | | |
| Number of values (total) | 12 | | | | |

Fig. 3D - post-hoc

| | | | | | | | | |
|---|---|---|---|---|---|---|---|---|
| Number of families | 1 | | | | | | | |
| Number of comparisons per family | 2 | | | | | | | |
| Alpha | 0.05 | | | | | | | |
| Dunnett's multiple comparisons test | Mean diff. | 95.00% CI of diff. | Below threshold? | Summary | Adjusted P Value | A-? | | |
| pWT (Vehicle) vs. pWT (A20) | 0.9833 | 0.1207 to 1.846 | Yes | * | 0.0277 | B | pWT (A20) | |
| pWT (Vehicle) vs. pWT (EDTA) | 1.047 | 0.1841 to 1.909 | Yes | * | 0.0203 | C | pWT (EDTA) | |
| Test details | Mean 1 | Mean 2 | Mean diff. | SE of diff. | n1 | n2 | q | DF |
| pWT (Vehicle) vs. pWT (A20) | 2.057 | 1.073 | 0.9833 | 0.3269 | 6 | 3 | 3.008 | 9 |
| pWT (Vehicle) vs. pWT (EDTA) | 2.057 | 1.01 | 1.047 | 0.3269 | 6 | 3 | 3.202 | 9 |

Fig. 3E - one-way-ANOVA %pEphB2

| | | | | | |
|---|---|---|---|---|---|
| Table Analyzed | phospho-EphB2 | | | | |
| Data sets analyzed | A-C | | | | |
| ANOVA summary | | | | | |
| F | 4.856 | | | | |
| P value | 0.0476 | | | | |
| P value summary | * | | | | |
| Significant diff. among means (P < 0.05)? | Yes | | | | |
| R squared | 0.5812 | | | | |
| Brown-Forsythe test | | | | | |
| F (DFn, DFd) | 1.771 (2, 7) | | | | |
| P value | 0.2386 | | | | |
| P value summary | ns | | | | |
| Are SDs significantly different (P < 0.05)? | No | | | | |
| Bartlett's test | | | | | |
| Bartlett's statistic (corrected) | | | | | |
| P value | | | | | |
| P value summary | | | | | |
| Are SDs significantly different (P < 0.05)? | | | | | |
| ANOVA table | SS | DF | MS | F (DFn, DFd) | P value |
| Treatment (between columns) | 0.006622 | 2 | 0.003311 | F (2, 7) = 4.856 | P=0.0476 |
| Residual (within columns) | 0.004772 | 7 | 0.0006818 | | |
| Total | 0.01139 | 9 | | | |
| Data summary | | | | | |
| Number of treatments (columns) | 3 | | | | |
| Number of values (total) | 10 | | | | |

Fig. 3E - post-hoc %pEphB2

| | | | | | | | | |
|---|---|---|---|---|---|---|---|---|
| Number of families | 1 | | | | | | | |
| Number of comparisons per family | 2 | | | | | | | |
| Alpha | 0.05 | | | | | | | |
| Dunnett's multiple comparisons test | Mean Diff. | 95.00% CI of diff. | Below threshold? | Summary | Adjusted P Value | A-? | | |
| Vehicle (PBS) vs. A20 | 0.054 | -0.008616 to 0.1166 | No | ns | 0.0859 | B | A20 | |
| Vehicle (PBS) vs. EDTA | 0.05175 | 0.0006239 to 0.1029 | Yes | * | 0.0477 | C | EDTA | |
| Test details | Mean 1 | Mean 2 | Mean Diff. | SE of diff. | n1 | n2 | q | DF |
| Vehicle (PBS) vs. A20 | 0.1325 | 0.0785 | 0.054 | 0.02261 | 4 | 2 | 2.388 | 7 |
| Vehicle (PBS) vs. EDTA | 0.1325 | 0.08075 | 0.05175 | 0.01846 | 4 | 4 | 2.803 | 7 |

Fig. 3E - one-way-ANOVA pEphB2

| | | | | | |
|---|---|---|---|---|---|
| Table Analyzed | phospho-EphB2 | | | | |
| Data sets analyzed | A-C | | | | |
| ANOVA summary | | | | | |
| F | 23.21 | | | | |
| P value | 0.0008 | | | | |
| P value summary | *** | | | | |
| Significant diff. among means (P < 0.05)? | Yes | | | | |
| R squared | 0.869 | | | | |
| Brown-Forsythe test | | | | | |
| F (DFn, DFd) | 1.252 (2, 7) | | | | |
| P value | 0.3429 | | | | |
| P value summary | ns | | | | |
| Are SDs significantly different (P < 0.05)? | No | | | | |
| Bartlett's test | | | | | |
| Bartlett's statistic (corrected) | | | | | |
| P value | | | | | |
| P value summary | | | | | |
| Are SDs significantly different (P < 0.05)? | | | | | |
| ANOVA table | SS | DF | MS | F (DFn, DFd) | P value |
| Treatment (between columns) | 2.232 | 2 | 1.116 | F (2, 7) = 23.21 | P=0.0008 |
| Residual (within columns) | 0.3366 | 7 | 0.04809 | | |
| Total | 2.569 | 9 | | | |
| Data summary | | | | | |
| Number of treatments (columns) | 3 | | | | |
| Number of values (total) | 10 | | | | |

Fig. 3E - post-hoc pEphB2

| | | | | | | | | |
|---|---|---|---|---|---|---|---|---|
| Number of families | 1 | | | | | | | |
| Number of comparisons per family | 2 | | | | | | | |
| Alpha | 0.05 | | | | | | | |
| Dunnett's multiple comparisons test | Mean Diff. | 95.00% CI of diff. | Below threshold? | Summary | Adjusted P Value | A-? | | |
| Vehicle (PBS) vs. A20 | 1.018 | 0.4916 to 1.543 | Yes | ** | 0.002 | B | A20 | |
| Vehicle (PBS) vs. EDTA | 0.935 | 0.5056 to 1.364 | Yes | *** | 0.001 | C | EDTA | |
| Test details | Mean 1 | Mean 2 | Mean Diff. | SE of diff. | n1 | n2 | q | DF |
| Vehicle (PBS) vs. A20 | 1.693 | 0.675 | 1.018 | 0.1899 | 4 | 2 | 5.358 | 7 |
| Vehicle (PBS) vs. EDTA | 1.693 | 0.7575 | 0.935 | 0.1551 | 4 | 4 | 6.03 | 7 |

Fig. 3E - one-way-ANOVA tEphB2

| | | | | | |
|---|---|---|---|---|---|
| Table Analyzed | total EphB2 | | | | |
| Data sets analyzed | A-C | | | | |
| ANOVA summary | | | | | |
| F | 2.498 | | | | |
| P value | 0.1518 | | | | |
| P value summary | ns | | | | |
| Significant diff. among means (P < 0.05)? | No | | | | |
| R squared | 0.4165 | | | | |
| Brown-Forsythe test | | | | | |
| F (DFn, DFd) | 0.01255 (2, 7) | | | | |
| P value | 0.9876 | | | | |
| P value summary | ns | | | | |
| Are SDs significantly different (P < 0.05)? | No | | | | |
| Bartlett's test | | | | | |
| Bartlett's statistic (corrected) | | | | | |
| P value | | | | | |
| P value summary | | | | | |
| Are SDs significantly different (P < 0.05)? | | | | | |
| ANOVA table | SS | DF | MS | F (DFn, DFd) | P value |
| Treatment (between columns) | 42.36 | 2 | 21.18 | F (2, 7) = 2.498 | P=0.1518 |
| Residual (within columns) | 59.36 | 7 | 8.48 | | |
| Total | 101.7 | 9 | | | |
| Data summary | | | | | |
| Number of treatments (columns) | 3 | | | | |
| Number of values (total) | 10 | | | | |

Fig. 3E - post-hoc tEphB2

| | | | | | | | | |
|---|---|---|---|---|---|---|---|---|
| Number of families | 1 | | | | | | | |
| Number of comparisons per family | 2 | | | | | | | |
| Alpha | 0.05 | | | | | | | |
| Dunnett's multiple comparisons test | Mean Diff. | 95.00% CI of diff. | Below threshold? | Summary | Adjusted P Value | A-? | | |
| Vehicle (PBS) vs. A20 | 4.933 | -2.051 to 11.92 | No | ns | 0.1588 | B | A20 | |
| Vehicle (PBS) vs. EDTA | 3.678 | -2.024 to 9.379 | No | ns | 0.2013 | C | EDTA | |
| Test details | Mean 1 | Mean 2 | Mean Diff. | SE of diff. | n1 | n2 | q | DF |
| Vehicle (PBS) vs. A20 | 13.3 | 8.365 | 4.933 | 2.522 | 4 | 2 | 1.956 | 7 |
| Vehicle (PBS) vs. EDTA | 13.3 | 9.62 | 3.678 | 2.059 | 4 | 4 | 1.786 | 7 |

Fig. 3F - 1-way-ANOVA %pEphB2a

| | |
|---|---|
| Table Analyzed | phospho-EphB2 |
| Data sets analyzed | A-C |
| Distribution assumption | Normal (Gaussian) |
| ANOVA summary | |
| F | 10.71 |
| P value | 0.0026 |
| P value summary | ** |
| Significant diff. among means (P < 0.05)? | Yes |
| R squared | 0.6607 |
| Brown-Forsythe test | |
| F (DFn, DFd) | 1.284 (2, 11) |
| P value | 0.3154 |
| P value summary | ns |
| Are SDs significantly different (P < 0.05)? | No |
| Bartlett's test | |
| Bartlett's statistic (corrected) | |
| P value | |
| P value summary | |
| Are SDs significantly different (P < 0.05)? | |

| ANOVA table | SS | DF | MS | F (DFn, DFd) | P value |
|---|---|---|---|---|---|
| Treatment (between columns) | 0.1645 | 2 | 0.08224 | F (2, 11) = 10.71 | P=0.0026 |
| Residual (within columns) | 0.08447 | 11 | 0.007679 | | |
| Total | 0.2489 | 13 | | | |

| | |
|---|---|
| Data summary | |
| Number of treatments (columns) | 3 |
| Number of values (total) | 14 |

Fig. 3F - post-hoc %pEphB2a

| | | | | | | | | |
|---|---|---|---|---|---|---|---|---|
| Number of families | 1 | | | | | | | |
| Number of comparisons per family | 2 | | | | | | | |
| Alpha | 0.05 | | | | | | | |
| Dunnett's multiple comparisons test | Mean diff. | 95.00% CI of diff. | Below threshold? | Summary | Adjusted P Value | A-? | | |
| Vehicle (PBS) vs. A20 | 0.2905 | 0.1140 to 0.4670 | Yes | ** | 0.0032 | B | A20 | |
| Vehicle (PBS) vs. EDTA | 0.323 | 0.1465 to 0.4995 | Yes | ** | 0.0015 | C | EDTA | |
| Test details | Mean 1 | Mean 2 | Mean diff. | SE of diff. | n1 | n2 | q | DF |
| Vehicle (PBS) vs. A20 | 0.53 | 0.2395 | 0.2905 | 0.07155 | 2 | 6 | 4.06 | 11 |
| Vehicle (PBS) vs. EDTA | 0.53 | 0.207 | 0.323 | 0.07155 | 2 | 6 | 4.514 | 11 |

Fig. 3F - 1-way-ANOVA pEphB2

| | | | | | |
|---|---|---|---|---|---|
| Table Analyzed | phospho-EphB2 | | | | |
| Data sets analyzed | A-C | | | | |
| ANOVA summary | | | | | |
| F | 15.01 | | | | |
| P value | 0.0007 | | | | |
| P value summary | *** | | | | |
| Significant diff. among means (P < 0.05)? | Yes | | | | |
| R squared | 0.7318 | | | | |
| Brown-Forsythe test | | | | | |
| F (DFn, DFd) | 0.06475 (2, 11) | | | | |
| P value | 0.9377 | | | | |
| P value summary | ns | | | | |
| Are SDs significantly different (P < 0.05)? | No | | | | |
| Bartlett's test | | | | | |
| Bartlett's statistic (corrected) | | | | | |
| P value | | | | | |
| P value summary | | | | | |
| Are SDs significantly different (P < 0.05)? | | | | | |
| ANOVA table | SS | DF | MS | F (DFn, DFd) | P value |
| Treatment (between columns) | 285.1 | 2 | 142.5 | F (2, 11) = 15.01 | P=0.0007 |
| Residual (within columns) | 104.5 | 11 | 9.497 | | |
| Total | 389.5 | 13 | | | |
| Data summary | | | | | |
| Number of treatments (columns) | 3 | | | | |
| Number of values (total) | 14 | | | | |

Fig. 3F - post-hoc pEphB2

| | |
|---|---|
| Number of families | 1 |
| Number of comparisons per family | 2 |
| Alpha | 0.05 |

| Dunnett's multiple comparisons test | Mean Diff. | 95.00% CI of diff. | Below threshold? | Summary | Adjusted P Value | A-? | |
|---|---|---|---|---|---|---|---|
| Vehicle (PBS) vs. A20 | 12.35 | 6.143 to 18.56 | Yes | *** | 0.0008 | B | A20 |
| Vehicle (PBS) vs. EDTA | 13.31 | 7.105 to 19.52 | Yes | *** | 0.0004 | C | EDTA |

| Test details | Mean 1 | Mean 2 | Mean Diff. | SE of diff. | n1 | n2 | q | DF |
|---|---|---|---|---|---|---|---|---|
| Vehicle (PBS) vs. A20 | 18.67 | 6.313 | 12.35 | 2.516 | 2 | 6 | 4.909 | 11 |
| Vehicle (PBS) vs. EDTA | 18.67 | 5.352 | 13.31 | 2.516 | 2 | 6 | 5.291 | 11 |

Fig. 3F - 1-way-ANOVA tEphB2

| | |
|---|---|
| Table Analyzed | total-EphB2 |
| Data sets analyzed | A-C |
| ANOVA summary | |
| F | 0.8386 |
| P value | 0.4582 |
| P value summary | ns |
| Significant diff. among means (P < 0.05)? | No |
| R squared | 0.1323 |
| Brown-Forsythe test | |
| F (DFn, DFd) | 2.273 (2, 11) |
| P value | 0.1492 |
| P value summary | ns |
| Are SDs significantly different (P < 0.05)? | No |
| Bartlett's test | |
| Bartlett's statistic (corrected) | |
| P value | |
| P value summary | |
| Are SDs significantly different (P < 0.05)? | |

| ANOVA table | SS | DF | MS | F (DFn, DFd) | P value |
|---|---|---|---|---|---|
| Treatment (between columns) | 218.5 | 2 | 109.2 | F (2, 11) = 0.8386 | P=0.4582 |
| Residual (within columns) | 1433 | 11 | 130.3 | | |
| Total | 1651 | 13 | | | |

| | |
|---|---|
| Data summary | |
| Number of treatments (columns) | 3 |
| Number of values (total) | 14 |

Fig. 3F - post-hoc tEphB2

| | |
|---|---|
| Number of families | 1 |
| Number of comparisons per family | 2 |
| Alpha | 0.05 |

| Dunnett's multiple comparisons test | Mean Diff. | 95.00% CI of diff. | Below threshold? | Summary | Adjusted P Value | A-? | |
|---|---|---|---|---|---|---|---|
| Vehicle (PBS) vs. A20 | 11.35 | -11.65 to 34.34 | No | ns | 0.3598 | B | A20 |
| Vehicle (PBS) vs. EDTA | 11.23 | -11.76 to 34.23 | No | ns | 0.3658 | C | EDTA |

| Test details | Mean 1 | Mean 2 | Mean Diff. | SE of diff. | n1 | n2 | q | DF |
|---|---|---|---|---|---|---|---|---|
| Vehicle (PBS) vs. A20 | 37.45 | 26.11 | 11.35 | 9.319 | 2 | 6 | 1.217 | 11 |
| Vehicle (PBS) vs. EDTA | 37.45 | 26.22 | 11.23 | 9.319 | 2 | 6 | 1.205 | 11 |

Fig. 3F - 1-way-ANOVA %pEphB2b

| | |
|---|---|
| Table Analyzed | phospho-EphB2 |
| Data sets analyzed | A-C |
| ANOVA summary | |
| F | 9.104 |
| P value | 0.0215 |
| P value summary | * |
| Significant diff. among means (P < 0.05)? | Yes |
| R squared | 0.7846 |
| Brown-Forsythe test | |
| F (DFn, DFd) | 3.903 (2, 5) |
| P value | 0.0953 |
| P value summary | ns |
| Are SDs significantly different (P < 0.05)? | No |
| Bartlett's test | |
| Bartlett's statistic (corrected) | |
| P value | |
| P value summary | |
| Are SDs significantly different (P < 0.05)? | |

| ANOVA table | SS | DF | MS | F (DFn, DFd) | P value |
|---|---|---|---|---|---|
| Treatment (between columns) | 0.1427 | 2 | 0.07136 | F (2, 5) = 9.104 | P=0.0215 |
| Residual (within columns) | 0.03919 | 5 | 0.007839 | | |
| Total | 0.1819 | 7 | | | |

| | |
|---|---|
| Data summary | |
| Number of treatments (columns) | 3 |
| Number of values (total) | 8 |

Fig. 3F - post-hoc %pEphB2b

| | | | | | | | | |
|---|---|---|---|---|---|---|---|---|
| Number of families | 1 | | | | | | | |
| Number of comparisons per family | 2 | | | | | | | |
| Alpha | 0.05 | | | | | | | |
| Dunnett's multiple comparisons test | Mean Diff. | 95.00% CI of diff. | Below threshold? | Summary | Adjusted P Value | A-? | | |
| Vehicle (PBS) vs. A20 | 0.2905 | 0.04804 to 0.5330 | Yes | * | 0.0263 | B | A20 | |
| Vehicle (PBS) vs. EDTA | 0.323 | 0.08054 to 0.5655 | Yes | * | 0.0175 | C | EDTA | |
| Test details | Mean 1 | Mean 2 | Mean Diff. | SE of diff. | n1 | n2 | q | DF |
| Vehicle (PBS) vs. A20 | 0.53 | 0.2395 | 0.2905 | 0.08082 | 2 | 3 | 3.594 | 5 |
| Vehicle (PBS) vs. EDTA | 0.53 | 0.207 | 0.323 | 0.08082 | 2 | 3 | 3.996 | 5 |

Fig. 3F - 1-way-ANOVA pEphB2b

| | | | | | |
|---|---|---|---|---|---|
| Table Analyzed | phospho-EphB2 | | | | |
| Data sets analyzed | A-C | | | | |
| ANOVA summary | | | | | |
| F | 35.21 | | | | |
| P value | 0.0011 | | | | |
| P value summary | ** | | | | |
| Significant diff. among means (P < 0.05)? | Yes | | | | |
| R squared | 0.9337 | | | | |
| Brown-Forsythe test | | | | | |
| F (DFn, DFd) | 1.666 (2, 5) | | | | |
| P value | 0.279 | | | | |
| P value summary | ns | | | | |
| Are SDs significantly different (P < 0.05)? | No | | | | |
| Bartlett's test | | | | | |
| Bartlett's statistic (corrected) | | | | | |
| P value | | | | | |
| P value summary | | | | | |
| Are SDs significantly different (P < 0.05)? | | | | | |
| ANOVA table | SS | DF | MS | F (DFn, DFd) | P value |
| Treatment (between columns) | 248.4 | 2 | 124.2 | F (2, 5) = 35.21 | P=0.0011 |
| Residual (within columns) | 17.64 | 5 | 3.527 | | |
| Total | 266.1 | 7 | | | |
| Data summary | | | | | |
| Number of treatments (columns) | 3 | | | | |
| Number of values (total) | 8 | | | | |

Fig. 3F - post-hoc pEphB2b

| | | | | | | | | |
|---|---|---|---|---|---|---|---|---|
| Number of families | 1 | | | | | | | |
| Number of comparisons per family | 2 | | | | | | | |
| Alpha | 0.05 | | | | | | | |
| Dunnett's multiple comparisons test | Mean Diff. | 95.00% CI of diff. | Below threshold? | Summary | Adjusted P Value | A-? | | |
| Vehicle (PBS) vs. A20 | 12.35 | 7.210 to 17.50 | Yes | ** | 0.0014 | B | A20 | |
| Vehicle (PBS) vs. EDTA | 13.31 | 8.170 to 18.46 | Yes | *** | 0.001 | C | EDTA | |
| Test details | Mean 1 | Mean 2 | Mean Diff. | SE of diff. | n1 | n2 | q | DF |
| Vehicle (PBS) vs. A20 | 18.67 | 6.312 | 12.35 | 1.714 | 2 | 3 | 7.205 | 5 |
| Vehicle (PBS) vs. EDTA | 18.67 | 5.352 | 13.31 | 1.714 | 2 | 3 | 7.765 | 5 |

Fig. 3F - 1-way-ANOVA tEphB2b

| | | | | | |
|---|---|---|---|---|---|
| Table Analyzed | total-EphB2 | | | | |
| Data sets analyzed | A-C | | | | |
| ANOVA summary | | | | | |
| F | 1.342 | | | | |
| P value | 0.3415 | | | | |
| P value summary | ns | | | | |
| Significant diff. among means (P < 0.05)? | No | | | | |
| R squared | 0.3493 | | | | |
| Brown-Forsythe test | | | | | |
| F (DFn, DFd) | 9.241 (2, 5) | | | | |
| P value | 0.0209 | | | | |
| P value summary | * | | | | |
| Are SDs significantly different (P < 0.05)? | Yes | | | | |
| Bartlett's test | | | | | |
| Bartlett's statistic (corrected) | | | | | |
| P value | | | | | |
| P value summary | | | | | |
| Are SDs significantly different (P < 0.05)? | | | | | |
| ANOVA table | SS | DF | MS | F (DFn, DFd) | P value |
| Treatment (between columns) | 169.1 | 2 | 84.57 | F (2, 5) = 1.342 | P=0.3415 |
| Residual (within columns) | 315.1 | 5 | 63.02 | | |
| Total | 484.2 | 7 | | | |
| Data summary | | | | | |
| Number of treatments (columns) | 3 | | | | |
| Number of values (total) | 8 | | | | |

Fig. 3F - post-hoc tEphB2b

| | |
|---|---|
| Number of families | 1 |
| Number of comparisons per family | 2 |
| Alpha | 0.05 |

| Dunnett's multiple comparisons test | Mean Diff. | 95.00% CI of diff. | Below threshold? | Summary | Adjusted P Value | A-? | |
|---|---|---|---|---|---|---|---|
| Vehicle (PBS) vs. A20 | 9.817 | -11.92 to 31.56 | No | ns | 0.3571 | B | A20 |
| Vehicle (PBS) vs. EDTA | 11.23 | -10.51 to 32.97 | No | ns | 0.2831 | C | EDTA |

| Test details | Mean 1 | Mean 2 | Mean Diff. | SE of diff. | n1 | n2 | q | DF |
|---|---|---|---|---|---|---|---|---|
| Vehicle (PBS) vs. A20 | 37.45 | 27.63 | 9.817 | 7.247 | 2 | 3 | 1.355 | 5 |
| Vehicle (PBS) vs. EDTA | 37.45 | 26.22 | 11.23 | 7.247 | 2 | 3 | 1.55 | 5 |

Fig. 4E - t-test - %pEphrinB2

| | |
|---|---|
| Table Analyzed | phospho-EB2 |
| Column B | EDTA |
| vs. | vs. |
| Column A | Vehicle |
| Unpaired t test | |
| P value | 0.0172 |
| P value summary | * |
| Significantly different (P < 0.05)? | Yes |
| One- or two-tailed P value? | Two-tailed |
| t, df | t=3.503, df=5 |
| How big is the difference? | |
| Mean of column A | 0.1087 |
| Mean of column B | 0.05186 |
| Difference between means (B - A) ± SEM | -0.05680 ± 0.01622 |
| 95% confidence interval | -0.09848 to -0.01512 |
| R squared (eta squared) | 0.7105 |
| F test to compare variances | |
| F, DFn, Dfd | 4.622, 3, 2 |
| P value | 0.366 |
| P value summary | ns |
| Significantly different (P < 0.05)? | No |
| Data analyzed | |
| Sample size, column A | 4 |
| Sample size, column B | 3 |

Fig. 4E - t-test - pEphrinB2

| | |
|---|---|
| Table Analyzed | phospho-EB2 |
| Column B | EDTA |
| vs. | vs. |
| Column A | Vehicle |
| Unpaired t test | |
| P value | 0.0011 |
| P value summary | ** |
| Significantly different (P < 0.05)? | Yes |
| One- or two-tailed P value? | Two-tailed |
| t, df | t=6.776, df=5 |
| How big is the difference? | |
| Mean of column A | 3.97 |
| Mean of column B | 1.8 |
| Difference between means (B - A) ± SEM | -2.170 ± 0.3203 |
| 95% confidence interval | -2.993 to -1.347 |
| R squared (eta squared) | 0.9018 |
| F test to compare variances | |
| F, DFn, Dfd | 3.584, 2, 3 |
| P value | 0.3205 |
| P value summary | ns |
| Significantly different (P < 0.05)? | No |
| Data analyzed | |
| Sample size, column A | 4 |
| Sample size, column B | 3 |

Fig. 4E - t-test - tEphrinB2

| | |
|---|---|
| Table Analyzed | total-EB1 |
| Column B | EDTA |
| vs. | vs. |
| Column A | Vehicle |
| Unpaired t test | |
| P value | 0.5665 |
| P value summary | ns |
| Significantly different (P < 0.05)? | No |
| One- or two-tailed P value? | Two-tailed |
| t, df | t=0.6133, df=5 |
| How big is the difference? | |
| Mean of column A | 38.02 |
| Mean of column B | 34.38 |
| Difference between means (B - A) ± SEM | -3.646 ± 5.945 |
| 95% confidence interval | -18.93 to 11.64 |
| R squared (eta squared) | 0.06996 |
| F test to compare variances | |
| F, DFn, Dfd | 3.824, 3, 2 |
| P value | 0.4284 |
| P value summary | ns |
| Significantly different (P < 0.05)? | No |
| Data analyzed | |
| Sample size, column A | 4 |
| Sample size, column B | 3 |

### Table S6: Statistics for Figure 5.

Fig. 5B - two-way ANOVA Results

| | | | | | |
|---|---|---|---|---|---|
| Table Analyzed | CFA - thermal Ratio for WT-E1-tFB1 for bar graph | | | | |
| Two-way ANOVA | Ordinary | | | | |
| Alpha | 0.05 | | | | |
| Source of Variation | % of total variation | P value | P value summary | Significant? | |
| Interaction | 12.48 | <0.0001 | **** | Yes | |
| Row Factor | 28.85 | <0.0001 | **** | Yes | |
| Column Factor | 25.76 | <0.0001 | **** | Yes | |
| ANOVA table | SS (Type III) | DF | MS | F (DFn, DFd) | P value |
| Interaction | 96.74 | 10 | 9.674 | F (10, 114) = 4.059 | P<0.0001 |
| Row Factor | 223.7 | 5 | 44.74 | F (5, 114) = 18.77 | P<0.0001 |
| Column Factor | 199.7 | 2 | 99.86 | F (2, 114) = 41.90 | P<0.0001 |
| Residual | 271.7 | 114 | 2.383 | | |
| Data summary | | | | | |
| Number of columns (Column Factor) | 3 | | | | |
| Number of rows (Row Factor) | 6 | | | | |
| Number of values | 132 | | | | |

Within each column, compare rows (simple effects within columns)

| | |
|---|---|
| Number of families | 3 |
| Number of comparisons per family | 15 |
| Alpha | 0.05 |

| Tukey's multiple comparisons test | Predicted (LS) Mean diff. | 95.00% CI of diff. | Below threshold? | Summary | Adjusted P Value |
|---|---|---|---|---|---|
| WT | | | | | |
| BL vs. 1 | -6.904 | -9.141 to -4.666 | Yes | **** | <0.0001 |
| BL vs. 2 | -3.984 | -6.221 to -1.746 | Yes | **** | <0.0001 |
| BL vs. 4 | -3.934 | -6.171 to -1.696 | Yes | **** | <0.0001 |
| BL vs. 6 | -3.214 | -5.451 to -0.9762 | Yes | *** | 0.0009 |
| BL vs. 9 | -1.945 | -4.183 to 0.2926 | No | ns | 0.127 |
| 1 vs. 2 | 2.92 | 0.6824 to 5.158 | Yes | ** | 0.0033 |
| 1 vs. 4 | 2.97 | 0.7324 to 5.208 | Yes | ** | 0.0027 |
| 1 vs. 6 | 3.69 | 1.452 to 5.928 | Yes | **** | <0.0001 |
| 1 vs. 9 | 4.959 | 2.721 to 7.196 | Yes | **** | <0.0001 |
| 2 vs. 4 | 0.05 | -2.188 to 2.288 | No | ns | >0.9999 |
| 2 vs. 6 | 0.77 | -1.468 to 3.008 | No | ns | 0.9179 |
| 2 vs. 9 | 2.039 | -0.1988 to 4.276 | No | ns | 0.0958 |
| 4 vs. 6 | 0.72 | -1.518 to 2.958 | No | ns | 0.9372 |
| 4 vs. 9 | 1.989 | -0.2488 to 4.226 | No | ns | 0.1116 |
| 6 vs. 9 | 1.269 | -0.9688 to 3.506 | No | ns | 0.5716 |
| EphB1-KO | | | | | |
| BL vs. 1 | -0.9122 | -3.022 to 1.197 | No | ns | 0.8093 |
| BL vs. 2 | -0.7711 | -2.881 to 1.338 | No | ns | 0.8962 |
| BL vs. 4 | -0.4467 | -2.556 to 1.663 | No | ns | 0.9898 |
| BL vs. 6 | -0.5933 | -2.703 to 1.516 | No | ns | 0.9642 |
| BL vs. 9 | -0.3189 | -2.428 to 1.791 | No | ns | 0.9979 |
| 1 vs. 2 | 0.1411 | -1.968 to 2.251 | No | ns | >0.9999 |
| 1 vs. 4 | 0.4656 | -1.644 to 2.575 | No | ns | 0.9877 |
| 1 vs. 6 | 0.3189 | -1.791 to 2.428 | No | ns | 0.9979 |
| 1 vs. 9 | 0.5933 | -1.516 to 2.703 | No | ns | 0.9642 |
| 2 vs. 4 | 0.3244 | -1.785 to 2.434 | No | ns | 0.9977 |
| 2 vs. 6 | 0.1778 | -1.932 to 2.287 | No | ns | 0.9999 |
| 2 vs. 9 | 0.4522 | -1.657 to 2.562 | No | ns | 0.9892 |
| 4 vs. 6 | -0.1467 | -2.256 to 1.963 | No | ns | >0.9999 |
| 4 vs. 9 | 0.1278 | -1.982 to 2.237 | No | ns | >0.9999 |
| 6 vs. 9 | 0.2744 | -1.835 to 2.384 | No | ns | 0.999 |
| EphB1-lacZ | | | | | |
| BL vs. 1 | -5.256 | -8.086 to -2.426 | Yes | **** | <0.0001 |
| BL vs. 2 | -2.658 | -5.488 to 0.1723 | No | ns | 0.0787 |
| BL vs. 4 | -2.052 | -4.882 to 0.7783 | No | ns | 0.2939 |
| BL vs. 6 | -1.024 | -3.854 to 1.806 | No | ns | 0.9002 |
| BL vs. 9 | -0.874 | -3.704 to 1.956 | No | ns | 0.947 |
| 1 vs. 2 | 2.598 | -0.2323 to 5.428 | No | ns | 0.0914 |
| 1 vs. 4 | 3.204 | 0.3737 to 6.034 | Yes | * | 0.0168 |
| 1 vs. 6 | 4.232 | 1.402 to 7.062 | Yes | *** | 0.0004 |
| 1 vs. 9 | 4.382 | 1.552 to 7.212 | Yes | *** | 0.0002 |
| 2 vs. 4 | 0.606 | -2.224 to 3.436 | No | ns | 0.9893 |
| 2 vs. 6 | 1.634 | -1.196 to 4.464 | No | ns | 0.552 |
| 2 vs. 9 | 1.784 | -1.046 to 4.614 | No | ns | 0.4527 |
| 4 vs. 6 | 1.028 | -1.802 to 3.858 | No | ns | 0.8987 |
| 4 vs. 9 | 1.178 | -1.652 to 4.008 | No | ns | 0.8329 |
| 6 vs. 9 | 0.15 | -2.680 to 2.980 | No | ns | >0.9999 |

| Test details | Predicted (LS) Mean 1 | Predicted (LS) Mean 2 | Predicted (LS) Mean diff. | SE of diff. | N1 | N2 | q | DF |
|---|---|---|---|---|---|---|---|---|
| WT | | | | | | | | |
| BL vs. 1 | 0.975 | 7.879 | -6.904 | 0.7719 | 8 | 8 | 12.65 | 114 |
| BL vs. 2 | 0.975 | 4.959 | -3.984 | 0.7719 | 8 | 8 | 7.299 | 114 |
| BL vs. 4 | 0.975 | 4.909 | -3.934 | 0.7719 | 8 | 8 | 7.207 | 114 |
| BL vs. 6 | 0.975 | 4.189 | -3.214 | 0.7719 | 8 | 8 | 5.888 | 114 |
| BL vs. 9 | 0.975 | 2.92 | -1.945 | 0.7719 | 8 | 8 | 3.563 | 114 |
| 1 vs. 2 | 7.879 | 4.959 | 2.92 | 0.7719 | 8 | 8 | 5.35 | 114 |
| 1 vs. 4 | 7.879 | 4.909 | 2.97 | 0.7719 | 8 | 8 | 5.441 | 114 |
| 1 vs. 6 | 7.879 | 4.189 | 3.69 | 0.7719 | 8 | 8 | 6.76 | 114 |
| 1 vs. 9 | 7.879 | 2.92 | 4.959 | 0.7719 | 8 | 8 | 9.085 | 114 |
| 2 vs. 4 | 4.959 | 4.909 | 0.05 | 0.7719 | 8 | 8 | 0.09161 | 114 |
| 2 vs. 6 | 4.959 | 4.189 | 0.77 | 0.7719 | 8 | 8 | 1.411 | 114 |
| 2 vs. 9 | 4.959 | 2.92 | 2.039 | 0.7719 | 8 | 8 | 3.735 | 114 |
| 4 vs. 6 | 4.909 | 4.189 | 0.72 | 0.7719 | 8 | 8 | 1.319 | 114 |
| 4 vs. 9 | 4.909 | 2.92 | 1.989 | 0.7719 | 8 | 8 | 3.644 | 114 |
| 6 vs. 9 | 4.189 | 2.92 | 1.269 | 0.7719 | 8 | 8 | 2.324 | 114 |
| EphB1-KO | | | | | | | | |
| BL vs. 1 | 0.9978 | 1.91 | -0.9122 | 0.7278 | 9 | 9 | 1.773 | 114 |
| BL vs. 2 | 0.9978 | 1.769 | -0.7711 | 0.7278 | 9 | 9 | 1.498 | 114 |
| BL vs. 4 | 0.9978 | 1.444 | -0.4467 | 0.7278 | 9 | 9 | 0.868 | 114 |
| BL vs. 6 | 0.9978 | 1.591 | -0.5933 | 0.7278 | 9 | 9 | 1.153 | 114 |
| BL vs. 9 | 0.9978 | 1.317 | -0.3189 | 0.7278 | 9 | 9 | 0.6197 | 114 |
| 1 vs. 2 | 1.91 | 1.769 | 0.1411 | 0.7278 | 9 | 9 | 0.2742 | 114 |
| 1 vs. 4 | 1.91 | 1.444 | 0.4656 | 0.7278 | 9 | 9 | 0.9047 | 114 |
| 1 vs. 6 | 1.91 | 1.591 | 0.3189 | 0.7278 | 9 | 9 | 0.6197 | 114 |
| 1 vs. 9 | 1.91 | 1.317 | 0.5933 | 0.7278 | 9 | 9 | 1.153 | 114 |
| 2 vs. 4 | 1.769 | 1.444 | 0.3244 | 0.7278 | 9 | 9 | 0.6305 | 114 |
| 2 vs. 6 | 1.769 | 1.591 | 0.1778 | 0.7278 | 9 | 9 | 0.3455 | 114 |
| 2 vs. 9 | 1.769 | 1.317 | 0.4522 | 0.7278 | 9 | 9 | 0.8788 | 114 |
| 4 vs. 6 | 1.444 | 1.591 | -0.1467 | 0.7278 | 9 | 9 | 0.285 | 114 |
| 4 vs. 9 | 1.444 | 1.317 | 0.1278 | 0.7278 | 9 | 9 | 0.2483 | 114 |
| 6 vs. 9 | 1.591 | 1.317 | 0.2744 | 0.7278 | 9 | 9 | 0.5333 | 114 |
| EphB1-lacZ | | | | | | | | |
| BL vs. 1 | 0.988 | 6.244 | -5.256 | 0.9764 | 5 | 5 | 7.613 | 114 |
| BL vs. 2 | 0.988 | 3.646 | -2.658 | 0.9764 | 5 | 5 | 3.85 | 114 |
| BL vs. 4 | 0.988 | 3.04 | -2.052 | 0.9764 | 5 | 5 | 2.972 | 114 |
| BL vs. 6 | 0.988 | 2.012 | -1.024 | 0.9764 | 5 | 5 | 1.483 | 114 |
| BL vs. 9 | 0.988 | 1.862 | -0.874 | 0.9764 | 5 | 5 | 1.266 | 114 |
| 1 vs. 2 | 6.244 | 3.646 | 2.598 | 0.9764 | 5 | 5 | 3.763 | 114 |
| 1 vs. 4 | 6.244 | 3.04 | 3.204 | 0.9764 | 5 | 5 | 4.641 | 114 |
| 1 vs. 6 | 6.244 | 2.012 | 4.232 | 0.9764 | 5 | 5 | 6.13 | 114 |
| 1 vs. 9 | 6.244 | 1.862 | 4.382 | 0.9764 | 5 | 5 | 6.347 | 114 |
| 2 vs. 4 | 3.646 | 3.04 | 0.606 | 0.9764 | 5 | 5 | 0.8777 | 114 |
| 2 vs. 6 | 3.646 | 2.012 | 1.634 | 0.9764 | 5 | 5 | 2.367 | 114 |
| 2 vs. 9 | 3.646 | 1.862 | 1.784 | 0.9764 | 5 | 5 | 2.584 | 114 |
| 4 vs. 6 | 3.04 | 2.012 | 1.028 | 0.9764 | 5 | 5 | 1.489 | 114 |
| 4 vs. 9 | 3.04 | 1.862 | 1.178 | 0.9764 | 5 | 5 | 1.706 | 114 |
| 6 vs. 9 | 2.012 | 1.862 | 0.15 | 0.9764 | 5 | 5 | 0.2173 | 114 |

Within each row, compare columns (simple effects within rows)

| | |
|---|---|
| Number of families | 6 |
| Number of comparisons per family | 3 |
| Alpha | 0.05 |

| Tukey's multiple comparisons test | Predicted (LS) Mean diff. | 95.00% CI of diff. | Below threshold? | Summary | Adjusted P Value |
|---|---|---|---|---|---|
| Baseline (-4 days) | | | | | |
| WT vs. EphB1-KO | -0.02278 | -1.804 to 1.759 | No | ns | 0.9995 |
| WT vs. EphB1-lacZ | -0.013 | -2.103 to 2.077 | No | ns | 0.9999 |
| EphB1-KO vs. EphB1-lacZ | 0.009778 | -2.035 to 2.055 | No | ns | >0.9999 |
| CFA +1 | | | | | |
| WT vs. EphB1-KO | 5.969 | 4.187 to 7.750 | Yes | **** | <0.0001 |
| WT vs. EphB1-lacZ | 1.635 | -0.4552 to 3.725 | No | ns | 0.1559 |
| EphB1-KO vs. EphB1-lacZ | -4.334 | -6.379 to -2.289 | Yes | **** | <0.0001 |
| CFA +2 | | | | | |
| WT vs. EphB1-KO | 3.19 | 1.408 to 4.971 | Yes | *** | 0.0001 |
| WT vs. EphB1-lacZ | 1.313 | -0.7772 to 3.403 | No | ns | 0.2988 |
| EphB1-KO vs. EphB1-lacZ | -1.877 | -3.922 to 0.1677 | No | ns | 0.079 |
| CFA +3/4 | | | | | |
| WT vs. EphB1-KO | 3.464 | 1.683 to 5.246 | Yes | **** | <0.0001 |
| WT vs. EphB1-lacZ | 1.869 | -0.2212 to 3.959 | No | ns | 0.0896 |
| EphB1-KO vs. EphB1-lacZ | -1.596 | -3.640 to 0.4493 | No | ns | 0.1573 |
| CFA +6/7 | | | | | |
| WT vs. EphB1-KO | 2.598 | 0.8162 to 4.379 | Yes | ** | 0.0022 |
| WT vs. EphB1-lacZ | 2.177 | 0.08675 to 4.267 | Yes | * | 0.0392 |
| EphB1-KO vs. EphB1-lacZ | -0.4209 | -2.466 to 1.624 | No | ns | 0.8768 |
| CFA +8/9 | | | | | |
| WT vs. EphB1-KO | 1.603 | -0.1781 to 3.385 | No | ns | 0.0869 |
| WT vs. EphB1-lacZ | 1.058 | -1.032 to 3.148 | No | ns | 0.4544 |
| EphB1-KO vs. EphB1-lacZ | -0.5453 | -2.590 to 1.500 | No | ns | 0.8021 |

| Test details | Predicted (LS) Mean 1 | Predicted (LS) Mean 2 | Predicted (LS) Mean diff. | SE of diff. | N1 | N2 | q | DF |
|---|---|---|---|---|---|---|---|---|
| Baseline (-4 days) | | | | | | | | |
| WT vs. EphB1-KO | 0.975 | 0.9978 | -0.02278 | 0.7502 | 8 | 9 | 0.04294 | 114 |
| WT vs. EphB1-lacZ | 0.975 | 0.988 | -0.013 | 0.8801 | 8 | 5 | 0.02089 | 114 |
| EphB1-KO vs. EphB1-lacZ | 0.9978 | 0.988 | 0.009778 | 0.8611 | 9 | 5 | 0.01606 | 114 |
| CFA +1 | | | | | | | | |
| WT vs. EphB1-KO | 7.879 | 1.91 | 5.969 | 0.7502 | 8 | 9 | 11.25 | 114 |
| WT vs. EphB1-lacZ | 7.879 | 6.244 | 1.635 | 0.8801 | 8 | 5 | 2.627 | 114 |
| EphB1-KO vs. EphB1-lacZ | 1.91 | 6.244 | -4.334 | 0.8611 | 9 | 5 | 7.118 | 114 |
| CFA +2 | | | | | | | | |
| WT vs. EphB1-KO | 4.959 | 1.769 | 3.19 | 0.7502 | 8 | 9 | 6.014 | 114 |
| WT vs. EphB1-lacZ | 4.959 | 3.646 | 1.313 | 0.8801 | 8 | 5 | 2.109 | 114 |
| EphB1-KO vs. EphB1-lacZ | 1.769 | 3.646 | -1.877 | 0.8611 | 9 | 5 | 3.083 | 114 |
| CFA +3/4 | | | | | | | | |
| WT vs. EphB1-KO | 4.909 | 1.444 | 3.464 | 0.7502 | 8 | 9 | 6.531 | 114 |
| WT vs. EphB1-lacZ | 4.909 | 3.04 | 1.869 | 0.8801 | 8 | 5 | 3.003 | 114 |
| EphB1-KO vs. EphB1-lacZ | 1.444 | 3.04 | -1.596 | 0.8611 | 9 | 5 | 2.62 | 114 |
| CFA +6/7 | | | | | | | | |
| WT vs. EphB1-KO | 4.189 | 1.591 | 2.598 | 0.7502 | 8 | 9 | 4.897 | 114 |
| WT vs. EphB1-lacZ | 4.189 | 2.012 | 2.177 | 0.8801 | 8 | 5 | 3.498 | 114 |
| EphB1-KO vs. EphB1-lacZ | 1.591 | 2.012 | -0.4209 | 0.8611 | 9 | 5 | 0.6912 | 114 |
| CFA +8/9 | | | | | | | | |
| WT vs. EphB1-KO | 2.92 | 1.317 | 1.603 | 0.7502 | 8 | 9 | 3.023 | 114 |
| WT vs. EphB1-lacZ | 2.92 | 1.862 | 1.058 | 0.8801 | 8 | 5 | 1.7 | 114 |
| EphB1-KO vs. EphB1-lacZ | 1.317 | 1.862 | -0.5453 | 0.8611 | 9 | 5 | 0.8956 | 114 |

Fig. 5C - two-way ANOVA Results

| | | | | | |
|---|---|---|---|---|---|
| Table Analyzed | Ratio of Right/Left Paw | | | | |
| Two-way ANOVA | Ordinary | | | | |
| Alpha | 0.05 | | | | |
| Source of Variation | % of total variation | P value | P value summary | Significant? | |
| Interaction | 25.22 | 0.0079 | ** | Yes | |
| Row Factor | 20.11 | 0.003 | ** | Yes | |
| Column Factor | 9.439 | 0.0286 | * | Yes | |
| ANOVA table | SS (Type III) | DF | MS | F (DFn, DFd) | P value |
| Interaction | 10.7 | 6 | 1.783 | F (6, 36) = 3.499 | P=0.0079 |
| Row Factor | 8.529 | 3 | 2.843 | F (3, 36) = 5.579 | P=0.0030 |
| Column Factor | 4.004 | 2 | 2.002 | F (2, 36) = 3.929 | P=0.0286 |
| Residual | 18.34 | 36 | 0.5096 | | |
| Data summary | | | | | |
| Number of columns (Column Factor) | 3 | | | | |
| Number of rows (Row Factor) | 4 | | | | |
| Number of values | 48 | | | | |

Fig. 5C - post-hoc - genotype

Within each column, compare rows (simple effects within columns)

| | |
|---|---|
| Number of families | 3 |
| Number of comparisons per family | 6 |
| Alpha | 0.05 |

| Tukey's multiple comparisons test | Predicted (LS) Mean diff. | 95.00% CI of diff. | Below threshold? | Summary | Adjusted P Value |
|---|---|---|---|---|---|
| EphB1-KO | | | | | |
| Baseline -2 vs. Baseline -1 | -0.123 | -1.482 to 1.236 | No | ns | 0.9948 |
| Baseline -2 vs. CFA +3 | 0.064 | -1.295 to 1.423 | No | ns | 0.9993 |
| Baseline -2 vs. CFA +8 | 0.03975 | -1.320 to 1.399 | No | ns | 0.9998 |
| Baseline -1 vs. CFA +3 | 0.187 | -1.172 to 1.546 | No | ns | 0.9824 |
| Baseline -1 vs. CFA +8 | 0.1628 | -1.197 to 1.522 | No | ns | 0.9882 |
| CFA +3 vs. CFA +8 | -0.02425 | -1.384 to 1.335 | No | ns | >0.9999 |
| WT | | | | | |
| Baseline -2 vs. Baseline -1 | -0.08 | -1.296 to 1.136 | No | ns | 0.998 |
| Baseline -2 vs. CFA +3 | -1.406 | -2.622 to -0.1903 | Yes | * | 0.0181 |
| Baseline -2 vs. CFA +8 | -1.859 | -3.075 to -0.6435 | Yes | ** | 0.0012 |
| Baseline -1 vs. CFA +3 | -1.326 | -2.542 to -0.1103 | Yes | * | 0.0281 |
| Baseline -1 vs. CFA +8 | -1.779 | -2.995 to -0.5635 | Yes | ** | 0.0019 |
| CFA +3 vs. CFA +8 | -0.4532 | -1.669 to 0.7627 | No | ns | 0.7482 |
| EphB1-lacZ | | | | | |
| Baseline -2 vs. Baseline -1 | 0.7217 | -0.8481 to 2.291 | No | ns | 0.6072 |
| Baseline -2 vs. CFA +3 | -1.342 | -2.912 to 0.2277 | No | ns | 0.1164 |
| Baseline -2 vs. CFA +8 | 0.08433 | -1.485 to 1.654 | No | ns | 0.9989 |
| Baseline -1 vs. CFA +3 | -2.064 | -3.633 to -0.4939 | Yes | ** | 0.0059 |
| Baseline -1 vs. CFA +8 | -0.6373 | -2.207 to 0.9324 | No | ns | 0.6957 |
| CFA +3 vs. CFA +8 | 1.426 | -0.1434 to 2.996 | No | ns | 0.0862 |

| Test details | Predicted (LS) Mean 1 | Predicted (LS) Mean 2 | Predicted (LS) Mean diff. | SE of diff. | N1 | N2 | q | DF |
|---|---|---|---|---|---|---|---|---|
| EphB1-KO | | | | | | | | |
| Baseline -2 vs. Baseline -1 | 1.064 | 1.187 | -0.123 | 0.5048 | 4 | 4 | 0.3446 | 36 |
| Baseline -2 vs. CFA +3 | 1.064 | 1 | 0.064 | 0.5048 | 4 | 4 | 0.1793 | 36 |
| Baseline -2 vs. CFA +8 | 1.064 | 1.024 | 0.03975 | 0.5048 | 4 | 4 | 0.1114 | 36 |
| Baseline -1 vs. CFA +3 | 1.187 | 1 | 0.187 | 0.5048 | 4 | 4 | 0.5239 | 36 |
| Baseline -1 vs. CFA +8 | 1.187 | 1.024 | 0.1628 | 0.5048 | 4 | 4 | 0.456 | 36 |
| CFA +3 vs. CFA +8 | 1 | 1.024 | -0.02425 | 0.5048 | 4 | 4 | 0.06794 | 36 |
| WT | | | | | | | | |
| Baseline -2 vs. Baseline -1 | 0.8814 | 0.9614 | -0.08 | 0.4515 | 5 | 5 | 0.2506 | 36 |
| Baseline -2 vs. CFA +3 | 0.8814 | 2.288 | -1.406 | 0.4515 | 5 | 5 | 4.405 | 36 |
| Baseline -2 vs. CFA +8 | 0.8814 | 2.741 | -1.859 | 0.4515 | 5 | 5 | 5.825 | 36 |
| Baseline -1 vs. CFA +3 | 0.9614 | 2.288 | -1.326 | 0.4515 | 5 | 5 | 4.154 | 36 |
| Baseline -1 vs. CFA +8 | 0.9614 | 2.741 | -1.779 | 0.4515 | 5 | 5 | 5.574 | 36 |
| CFA +3 vs. CFA +8 | 2.288 | 2.741 | -0.4532 | 0.4515 | 5 | 5 | 1.42 | 36 |
| EphB1-lacZ | | | | | | | | |
| Baseline -2 vs. Baseline -1 | 1.465 | 0.7437 | 0.7217 | 0.5828 | 3 | 3 | 1.751 | 36 |
| Baseline -2 vs. CFA +3 | 1.465 | 2.807 | -1.342 | 0.5828 | 3 | 3 | 3.256 | 36 |
| Baseline -2 vs. CFA +8 | 1.465 | 1.381 | 0.08433 | 0.5828 | 3 | 3 | 0.2046 | 36 |
| Baseline -1 vs. CFA +3 | 0.7437 | 2.807 | -2.064 | 0.5828 | 3 | 3 | 5.007 | 36 |
| Baseline -1 vs. CFA +8 | 0.7437 | 1.381 | -0.6373 | 0.5828 | 3 | 3 | 1.546 | 36 |
| CFA +3 vs. CFA +8 | 2.807 | 1.381 | 1.426 | 0.5828 | 3 | 3 | 3.461 | 36 |

Within each row, compare columns (simple effects within rows)

| | |
|---|---|
| Number of families | 4 |
| Number of comparisons per family | 3 |
| Alpha | 0.05 |

| Tukey's multiple comparisons test | Predicted (LS) Mean diff. | 95.00% CI of diff. | Below threshold? | Summary | Adjusted P Value |
|---|---|---|---|---|---|
| Baseline -2 | | | | | |
| EphB1-KO vs. WT | 0.1826 | -0.9879 to 1.353 | No | ns | 0.9231 |
| EphB1-KO vs. EphB1-lacZ | -0.4013 | -1.734 to 0.9313 | No | ns | 0.7438 |
| WT vs. EphB1-lacZ | -0.5839 | -1.858 to 0.6903 | No | ns | 0.5081 |
| Baseline -1 | | | | | |
| EphB1-KO vs. WT | 0.2256 | -0.9449 to 1.396 | No | ns | 0.8852 |
| EphB1-KO vs. EphB1-lacZ | 0.4433 | -0.8893 to 1.776 | No | ns | 0.6974 |
| WT vs. EphB1-lacZ | 0.2177 | -1.057 to 1.492 | No | ns | 0.9086 |
| CFA +3 | | | | | |
| EphB1-KO vs. WT | -1.288 | -2.458 to -0.1171 | Yes | * | 0.0284 |
| EphB1-KO vs. EphB1-lacZ | -1.807 | -3.140 to -0.4747 | Yes | ** | 0.0058 |
| WT vs. EphB1-lacZ | -0.5197 | -1.794 to 0.7545 | No | ns | 0.5835 |
| CFA +8 | | | | | |
| EphB1-KO vs. WT | -1.717 | -2.887 to -0.5461 | Yes | ** | 0.0028 |
| EphB1-KO vs. EphB1-lacZ | -0.3568 | -1.689 to 0.9759 | No | ns | 0.7911 |
| WT vs. EphB1-lacZ | 1.36 | 0.08556 to 2.634 | Yes | * | 0.0343 |

| Test details | Predicted (LS) Mean 1 | Predicted (LS) Mean 2 | Predicted (LS) Mean diff. | SE of diff. | N1 | N2 | q | DF |
|---|---|---|---|---|---|---|---|---|
| Baseline -2 | | | | | | | | |
| EphB1-KO vs. WT | 1.064 | 0.8814 | 0.1826 | 0.4789 | 4 | 5 | 0.5393 | 36 |
| EphB1-KO vs. EphB1-lacZ | 1.064 | 1.465 | -0.4013 | 0.5452 | 4 | 3 | 1.041 | 36 |
| WT vs. EphB1-lacZ | 0.8814 | 1.465 | -0.5839 | 0.5213 | 5 | 3 | 1.584 | 36 |
| Baseline -1 | | | | | | | | |
| EphB1-KO vs. WT | 1.187 | 0.9614 | 0.2256 | 0.4789 | 4 | 5 | 0.6663 | 36 |
| EphB1-KO vs. EphB1-lacZ | 1.187 | 0.7437 | 0.4433 | 0.5452 | 4 | 3 | 1.15 | 36 |
| WT vs. EphB1-lacZ | 0.9614 | 0.7437 | 0.2177 | 0.5213 | 5 | 3 | 0.5907 | 36 |
| CFA +3 | | | | | | | | |
| EphB1-KO vs. WT | 1 | 2.288 | -1.288 | 0.4789 | 4 | 5 | 3.803 | 36 |
| EphB1-KO vs. EphB1-lacZ | 1 | 2.807 | -1.807 | 0.5452 | 4 | 3 | 4.688 | 36 |
| WT vs. EphB1-lacZ | 2.288 | 2.807 | -0.5197 | 0.5213 | 5 | 3 | 1.41 | 36 |
| CFA +8 | | | | | | | | |
| EphB1-KO vs. WT | 1.024 | 2.741 | -1.717 | 0.4789 | 4 | 5 | 5.07 | 36 |
| EphB1-KO vs. EphB1-lacZ | 1.024 | 1.381 | -0.3568 | 0.5452 | 4 | 3 | 0.9254 | 36 |
| WT vs. EphB1-lacZ | 2.741 | 1.381 | 1.36 | 0.5213 | 5 | 3 | 3.689 | 36 |

Fig. 5D - one-way ANOVA Results

| | | | | | |
|---|---|---|---|---|---|
| Table Analyzed | Thermal Pain Ratio | | | | |
| Data sets analyzed | A-D | | | | |
| Distribution assumption | Normal (Gaussian) | | | | |
| ANOVA summary | | | | | |
| F | 6.099 | | | | |
| P value | 0.0183 | | | | |
| P value summary | * | | | | |
| Significant diff. among means (P < 0.05)? | Yes | | | | |
| R squared | 0.6958 | | | | |
| Brown-Forsythe test | | | | | |
| F (DFn, DFd) | 0.5324 (3, 8) | | | | |
| P value | 0.6728 | | | | |
| P value summary | ns | | | | |
| Are SDs significantly different (P < 0.05)? | No | | | | |
| Bartlett's test | | | | | |
| Bartlett's statistic (corrected) | | | | | |
| P value | | | | | |
| P value summary | | | | | |
| Are SDs significantly different (P < 0.05)? | | | | | |
| ANOVA table | SS | DF | MS | F (DFn, DFd) | P value |
| Treatment (between columns) | 3.903 | 3 | 1.301 | F (3, 8) = 6.099 | P=0.0183 |
| Residual (within columns) | 1.707 | 8 | 0.2133 | | |
| Total | 5.61 | 11 | | | |
| Data summary | | | | | |
| Number of treatments (columns) | 4 | | | | |
| Number of values (total) | 12 | | | | |

Fig. 5D - post-hoc

| | |
|---|---|
| Number of families | 1 |
| Number of comparisons per family | 3 |
| Alpha | 0.05 |

| Dunnett's multiple comparisons test | Mean diff. | 95.00% CI of diff. | Below threshold? | Summary | Adjusted P Value | A-? | |
|---|---|---|---|---|---|---|---|
| Vehicle vs. A20 | 1.24 | 0.1540 to 2.326 | Yes | * | 0.0274 | B | A20 |
| Vehicle vs. MCD | 0.6033 | -0.4827 to 1.689 | No | ns | 0.3184 | C | MCD |
| Vehicle vs. A20 + MCD | 1.457 | 0.3707 to 2.543 | Yes | * | 0.0121 | D | \20 + MCD |

| Test details | Mean 1 | Mean 2 | Mean diff. | SE of diff. | n1 | n2 | q | DF |
|---|---|---|---|---|---|---|---|---|
| Vehicle vs. A20 | 2.223 | 0.9833 | 1.24 | 0.3771 | 3 | 3 | 3.288 | 8 |
| Vehicle vs. MCD | 2.223 | 1.62 | 0.6033 | 0.3771 | 3 | 3 | 1.6 | 8 |
| Vehicle vs. A20 + MCD | 2.223 | 0.7667 | 1.457 | 0.3771 | 3 | 3 | 3.863 | 8 |

Fig. 5E - one-way ANOVA Results

| | |
|---|---|
| Table Analyzed | Thermal Pain Ratio |
| Data sets analyzed | A-C |
| Distribution assumption | Normal (Gaussian) |
| ANOVA summary | |
| F | 12.93 |
| P value | 0.0003 |
| P value summary | *** |
| Significant diff. among means (P < 0.05)? | Yes |
| R squared | 0.5764 |
| Brown-Forsythe test | |
| F (DFn, DFd) | 1.745 (2, 19) |
| P value | 0.2016 |
| P value summary | ns |
| Are SDs significantly different (P < 0.05)? | No |
| Bartlett's test | |
| Bartlett's statistic (corrected) | 3.185 |
| P value | 0.2034 |
| P value summary | ns |
| Are SDs significantly different (P < 0.05)? | No |

| ANOVA table | SS | DF | MS | F (DFn, DFd) | P value |
|---|---|---|---|---|---|
| Treatment (between columns) | 16.14 | 2 | 8.07 | F (2, 19) = 12.93 | P=0.0003 |
| Residual (within columns) | 11.86 | 19 | 0.6242 | | |
| Total | 28 | 21 | | | |

| | |
|---|---|
| Data summary | |
| Number of treatments (columns) | 3 |
| Number of values (total) | 22 |

Fig. 5E - post-hoc

| | |
|---|---|
| Number of families | 1 |
| Number of comparisons per family | 2 |
| Alpha | 0.05 |

| Dunnett's multiple comparisons test | Mean diff. | 95.00% CI of diff. | Below threshold? | Summary | Adjusted P Value | A-? | |
|---|---|---|---|---|---|---|---|
| Vehicle vs. A20 | 1.716 | 0.7378 to 2.695 | Yes | *** | 0.0009 | B | A20 |
| Vehicle vs. QPB4-Bn | 1.839 | 0.8607 to 2.818 | Yes | *** | 0.0005 | C | QPB4-Bn |

| Test details | Mean 1 | Mean 2 | Mean diff. | SE of diff. | n1 | n2 | q | DF |
|---|---|---|---|---|---|---|---|---|
| Vehicle vs. A20 | 3.246 | 1.53 | 1.716 | 0.4089 | 8 | 7 | 4.197 | 19 |
| Vehicle vs. QPB4-Bn | 3.246 | 1.407 | 1.839 | 0.4089 | 8 | 7 | 4.498 | 19 |

Fig. 5F - two-way ANOVA Results

| Table Analyzed | R/L Paw - pain ratio for bar graph | | | | |
|---|---|---|---|---|---|
| Two-way ANOVA | Ordinary | | | | |
| Alpha | 0.05 | | | | |
| Source of Variation | % of total variation | P value | P value summary | Significant? | |
| Interaction | 19.62 | 0.0006 | *** | Yes | |
| Row Factor | 22.6 | <0.0001 | **** | Yes | |
| Column Factor | 27.83 | <0.0001 | **** | Yes | |
| ANOVA table | SS | DF | MS | F (DFn, DFd) | P value |
| Interaction | 38.63 | 12 | 3.22 | F (12, 63) = 3.441 | P=0.0006 |
| Row Factor | 44.49 | 6 | 7.415 | F (6, 63) = 7.924 | P<0.0001 |
| Column Factor | 54.79 | 2 | 27.4 | F (2, 63) = 29.28 | P<0.0001 |
| Residual | 58.95 | 63 | 0.9358 | | |
| Data summary | | | | | |
| Number of columns (Column Factor) | 3 | | | | |
| Number of rows (Row Factor) | 7 | | | | |
| Number of values | 84 | | | | |

Fig. 5F - post-hoc - genotype

Within each column, compare rows (simple effects within columns)

| | |
|---|---|
| Number of families | 3 |
| Number of comparisons per family | 6 |
| Alpha | 0.05 |

| Dunnett's multiple comparisons test | Mean diff. | 95.00% CI of diff. | Below threshold? | Summary | Adjusted P Value |
|---|---|---|---|---|---|
| A20 (IP Injection) | | | | | |
| Baseline vs. CFA +1 | -0.6675 | -2.472 to 1.137 | No | ns | 0.8351 |
| Baseline vs. CFA +2 | -0.705 | -2.509 to 1.099 | No | ns | 0.8016 |
| Baseline vs. CFA +7 | -0.7 | -2.504 to 1.104 | No | ns | 0.8062 |
| Baseline vs. CFA +9 | -0.6925 | -2.497 to 1.112 | No | ns | 0.813 |
| Baseline vs. CFA +11 | -0.515 | -2.319 to 1.289 | No | ns | 0.94 |
| Baseline vs. CFA +14 | -0.31 | -2.114 to 1.494 | No | ns | 0.9949 |
| A20 (oral gavage) | | | | | |
| Baseline vs. CFA +1 | -0.925 | -2.729 to 0.8795 | No | ns | 0.5761 |
| Baseline vs. CFA +2 | -1.058 | -2.862 to 0.7470 | No | ns | 0.4423 |
| Baseline vs. CFA +7 | -0.4325 | -2.237 to 1.372 | No | ns | 0.9727 |
| Baseline vs. CFA +9 | -0.6375 | -2.442 to 1.167 | No | ns | 0.86 |
| Baseline vs. CFA +11 | -0.7525 | -2.557 to 1.052 | No | ns | 0.756 |
| Baseline vs. CFA +14 | -0.2675 | -2.072 to 1.537 | No | ns | 0.9977 |
| Vehicle | | | | | |
| Baseline vs. CFA +1 | -2.825 | -4.629 to -1.021 | Yes | *** | 0.0006 |
| Baseline vs. CFA +2 | -5.14 | -6.944 to -3.336 | Yes | **** | <0.0001 |
| Baseline vs. CFA +7 | -2.975 | -4.779 to -1.171 | Yes | *** | 0.0003 |
| Baseline vs. CFA +9 | -2.7 | -4.504 to -0.8955 | Yes | ** | 0.0011 |
| Baseline vs. CFA +11 | -1.568 | -3.372 to 0.2370 | No | ns | 0.1118 |
| Baseline vs. CFA +14 | -0.105 | -1.909 to 1.699 | No | ns | >0.9999 |

| Test details | Mean 1 | Mean 2 | Mean diff. | SE of diff. | N1 | N2 | q | DF |
|---|---|---|---|---|---|---|---|---|
| A20 (IP Injection) | | | | | | | | |
| Baseline vs. CFA +1 | 0.965 | 1.633 | -0.6675 | 0.684 | 4 | 4 | 0.9759 | 63 |
| Baseline vs. CFA +2 | 0.965 | 1.67 | -0.705 | 0.684 | 4 | 4 | 1.031 | 63 |
| Baseline vs. CFA +7 | 0.965 | 1.665 | -0.7 | 0.684 | 4 | 4 | 1.023 | 63 |
| Baseline vs. CFA +9 | 0.965 | 1.658 | -0.6925 | 0.684 | 4 | 4 | 1.012 | 63 |
| Baseline vs. CFA +11 | 0.965 | 1.48 | -0.515 | 0.684 | 4 | 4 | 0.7529 | 63 |
| Baseline vs. CFA +14 | 0.965 | 1.275 | -0.31 | 0.684 | 4 | 4 | 0.4532 | 63 |
| A20 (oral gavage) | | | | | | | | |
| Baseline vs. CFA +1 | 0.9875 | 1.913 | -0.925 | 0.684 | 4 | 4 | 1.352 | 63 |
| Baseline vs. CFA +2 | 0.9875 | 2.045 | -1.058 | 0.684 | 4 | 4 | 1.546 | 63 |
| Baseline vs. CFA +7 | 0.9875 | 1.42 | -0.4325 | 0.684 | 4 | 4 | 0.6323 | 63 |
| Baseline vs. CFA +9 | 0.9875 | 1.625 | -0.6375 | 0.684 | 4 | 4 | 0.932 | 63 |
| Baseline vs. CFA +11 | 0.9875 | 1.74 | -0.7525 | 0.684 | 4 | 4 | 1.1 | 63 |
| Baseline vs. CFA +14 | 0.9875 | 1.255 | -0.2675 | 0.684 | 4 | 4 | 0.3911 | 63 |
| Vehicle | | | | | | | | |
| Baseline vs. CFA +1 | 1.048 | 3.873 | -2.825 | 0.684 | 4 | 4 | 4.13 | 63 |
| Baseline vs. CFA +2 | 1.048 | 6.188 | -5.14 | 0.684 | 4 | 4 | 7.514 | 63 |
| Baseline vs. CFA +7 | 1.048 | 4.023 | -2.975 | 0.684 | 4 | 4 | 4.349 | 63 |
| Baseline vs. CFA +9 | 1.048 | 3.748 | -2.7 | 0.684 | 4 | 4 | 3.947 | 63 |
| Baseline vs. CFA +11 | 1.048 | 2.615 | -1.568 | 0.684 | 4 | 4 | 2.292 | 63 |
| Baseline vs. CFA +14 | 1.048 | 1.153 | -0.105 | 0.684 | 4 | 4 | 0.1535 | 63 |

Within each row, compare columns (simple effects within rows)

| | |
|---|---|
| Number of families | 7 |
| Number of comparisons per family | 2 |
| Alpha | 0.05 |

| Dunnett's multiple comparisons test | Mean diff. | 95.00% CI of diff. | Below threshold? | Summary | Adjusted P Value |
|---|---|---|---|---|---|
| Baseline | | | | | |
| Vehicle vs. A20 (IP Injection) | 0.0825 | -1.465 to 1.630 | No | ns | 0.9894 |
| Vehicle vs. A20 (oral gavage) | 0.06 | -1.488 to 1.608 | No | ns | 0.9944 |
| CFA +1 | | | | | |
| Vehicle vs. A20 (IP Injection) | 2.24 | 0.6922 to 3.788 | Yes | ** | 0.0033 |
| Vehicle vs. A20 (oral gavage) | 1.96 | 0.4122 to 3.508 | Yes | * | 0.0108 |
| CFA +2 | | | | | |
| Vehicle vs. A20 (IP Injection) | 4.518 | 2.970 to 6.065 | Yes | **** | <0.0001 |
| Vehicle vs. A20 (oral gavage) | 4.143 | 2.595 to 5.690 | Yes | **** | <0.0001 |
| CFA +7 | | | | | |
| Vehicle vs. A20 (IP Injection) | 2.358 | 0.8097 to 3.905 | Yes | ** | 0.002 |
| Vehicle vs. A20 (oral gavage) | 2.603 | 1.055 to 4.150 | Yes | *** | 0.0006 |
| CFA +9 | | | | | |
| Vehicle vs. A20 (IP Injection) | 2.09 | 0.5422 to 3.638 | Yes | ** | 0.0063 |
| Vehicle vs. A20 (oral gavage) | 2.123 | 0.5747 to 3.670 | Yes | ** | 0.0055 |
| CFA +11 | | | | | |
| Vehicle vs. A20 (IP Injection) | 1.135 | -0.4128 to 2.683 | No | ns | 0.1781 |
| Vehicle vs. A20 (oral gavage) | 0.875 | -0.6728 to 2.423 | No | ns | 0.341 |
| CFA +14 | | | | | |
| Vehicle vs. A20 (IP Injection) | -0.1225 | -1.670 to 1.425 | No | ns | 0.9768 |
| Vehicle vs. A20 (oral gavage) | -0.1025 | -1.650 to 1.445 | No | ns | 0.9837 |

| Test details | Mean 1 | Mean 2 | Mean diff. | SE of diff. | N1 | N2 | q | DF |
|---|---|---|---|---|---|---|---|---|
| Baseline | | | | | | | | |
| Vehicle vs. A20 (IP Injection) | 1.048 | 0.965 | 0.0825 | 0.684 | 4 | 4 | 0.1206 | 63 |
| Vehicle vs. A20 (oral gavage) | 1.048 | 0.9875 | 0.06 | 0.684 | 4 | 4 | 0.08772 | 63 |
| CFA +1 | | | | | | | | |
| Vehicle vs. A20 (IP Injection) | 3.873 | 1.633 | 2.24 | 0.684 | 4 | 4 | 3.275 | 63 |
| Vehicle vs. A20 (oral gavage) | 3.873 | 1.913 | 1.96 | 0.684 | 4 | 4 | 2.865 | 63 |
| CFA +2 | | | | | | | | |
| Vehicle vs. A20 (IP Injection) | 6.188 | 1.67 | 4.518 | 0.684 | 4 | 4 | 6.604 | 63 |
| Vehicle vs. A20 (oral gavage) | 6.188 | 2.045 | 4.143 | 0.684 | 4 | 4 | 6.056 | 63 |
| CFA +7 | | | | | | | | |
| Vehicle vs. A20 (IP Injection) | 4.023 | 1.665 | 2.358 | 0.684 | 4 | 4 | 3.447 | 63 |
| Vehicle vs. A20 (oral gavage) | 4.023 | 1.42 | 2.603 | 0.684 | 4 | 4 | 3.805 | 63 |
| CFA +9 | | | | | | | | |
| Vehicle vs. A20 (IP Injection) | 3.748 | 1.658 | 2.09 | 0.684 | 4 | 4 | 3.055 | 63 |
| Vehicle vs. A20 (oral gavage) | 3.748 | 1.625 | 2.123 | 0.684 | 4 | 4 | 3.103 | 63 |
| CFA +11 | | | | | | | | |
| Vehicle vs. A20 (IP Injection) | 2.615 | 1.48 | 1.135 | 0.684 | 4 | 4 | 1.659 | 63 |

Fig. 5G - two-way ANOVA Results

| | | | | | |
|---|---|---|---|---|---|
| Table Analyzed | Ratio R/L Paw | | | | |
| Two-way ANOVA | Ordinary | | | | |
| Alpha | 0.05 | | | | |
| Source of Variation | % of total variation | P value | P value summary | Significant? | |
| Interaction | 25.62 | 0.0017 | ** | Yes | |
| Row Factor | 10.09 | 0.0276 | * | Yes | |
| Column Factor | 26.63 | <0.0001 | **** | Yes | |
| ANOVA table | SS | DF | MS | F (DFn, DFd) | P value |
| Interaction | 11.33 | 8 | 1.416 | F (8, 45) = 3.827 | P=0.0017 |
| Row Factor | 4.46 | 4 | 1.115 | F (4, 45) = 3.013 | P=0.0276 |
| Column Factor | 11.77 | 2 | 5.887 | F (2, 45) = 15.91 | P<0.0001 |
| Residual | 16.65 | 45 | 0.3701 | | |
| Data summary | | | | | |
| Number of columns (Column Factor) | 3 | | | | |
| Number of rows (Row Factor) | 5 | | | | |
| Number of values | 60 | | | | |

Fig. 5G - post-hoc - genotype

Within each column, compare rows (simple effects within columns)

| | |
|---|---|
| Number of families | 3 |
| Number of comparisons per family | 4 |
| Alpha | 0.05 |

| Dunnett's multiple comparisons test | Mean diff. | 95.00% CI of diff. | Below threshold? | Summary | Adjusted P Value |
|---|---|---|---|---|---|
| A20 (IP Injection) | | | | | |
| Baseline vs. VF +3 | 0.0675 | -1.021 to 1.156 | No | ns | 0.9996 |
| Baseline vs. VF +8 | 0.005 | -1.084 to 1.094 | No | ns | >0.9999 |
| Baseline vs. VF +10 | 0.0325 | -1.056 to 1.121 | No | ns | >0.9999 |
| Baseline vs. VF +15 | 0.0375 | -1.051 to 1.126 | No | ns | >0.9999 |
| A20 (oral gavage) | | | | | |
| Baseline vs. VF +3 | -0.065 | -1.154 to 1.024 | No | ns | 0.9996 |
| Baseline vs. VF +8 | 0.165 | -0.9239 to 1.254 | No | ns | 0.9861 |
| Baseline vs. VF +10 | -0.0125 | -1.101 to 1.076 | No | ns | >0.9999 |
| Baseline vs. VF +15 | -0.1175 | -1.206 to 0.9714 | No | ns | 0.9961 |
| Vehicle | | | | | |
| Baseline vs. VF +3 | -1.623 | -2.711 to -0.5336 | Yes | ** | 0.0018 |
| Baseline vs. VF +8 | -2.2 | -3.289 to -1.111 | Yes | **** | <0.0001 |
| Baseline vs. VF +10 | -0.855 | -1.944 to 0.2339 | No | ns | 0.1616 |
| Baseline vs. VF +15 | 0.0375 | -1.051 to 1.126 | No | ns | >0.9999 |

| Test details | Mean 1 | Mean 2 | Mean diff. | SE of diff. | N1 | N2 | q | DF |
|---|---|---|---|---|---|---|---|---|
| A20 (IP Injection) | | | | | | | | |
| Baseline vs. VF +3 | 1 | 0.9325 | 0.0675 | 0.4301 | 4 | 4 | 0.1569 | 45 |
| Baseline vs. VF +8 | 1 | 0.995 | 0.005 | 0.4301 | 4 | 4 | 0.01162 | 45 |
| Baseline vs. VF +10 | 1 | 0.9675 | 0.0325 | 0.4301 | 4 | 4 | 0.07556 | 45 |
| Baseline vs. VF +15 | 1 | 0.9625 | 0.0375 | 0.4301 | 4 | 4 | 0.08718 | 45 |
| A20 (oral gavage) | | | | | | | | |
| Baseline vs. VF +3 | 1 | 1.065 | -0.065 | 0.4301 | 4 | 4 | 0.1511 | 45 |
| Baseline vs. VF +8 | 1 | 0.835 | 0.165 | 0.4301 | 4 | 4 | 0.3836 | 45 |
| Baseline vs. VF +10 | 1 | 1.013 | -0.0125 | 0.4301 | 4 | 4 | 0.02906 | 45 |
| Baseline vs. VF +15 | 1 | 1.118 | -0.1175 | 0.4301 | 4 | 4 | 0.2732 | 45 |
| Vehicle | | | | | | | | |
| Baseline vs. VF +3 | 1 | 2.623 | -1.623 | 0.4301 | 4 | 4 | 3.772 | 45 |
| Baseline vs. VF +8 | 1 | 3.2 | -2.2 | 0.4301 | 4 | 4 | 5.115 | 45 |
| Baseline vs. VF +10 | 1 | 1.855 | -0.855 | 0.4301 | 4 | 4 | 1.988 | 45 |
| Baseline vs. VF +15 | 1 | 0.9625 | 0.0375 | 0.4301 | 4 | 4 | 0.08718 | 45 |

Fig. 5G - post-hoc - time

Within each row, compare columns (simple effects within rows)

| | |
|---|---|
| Number of families | 5 |
| Number of comparisons per family | 2 |
| Alpha | 0.05 |

| Dunnett's multiple comparisons test | Mean diff. | 95.00% CI of diff. | Below threshold? | Summary | Adjusted P Value |
|---|---|---|---|---|---|
| Baseline | | | | | |
| Vehicle vs. A20 (IP Injection) | 0 | -0.9823 to 0.9823 | No | ns | >0.9999 |
| Vehicle vs. A20 (oral gavage) | 0 | -0.9823 to 0.9823 | No | ns | >0.9999 |
| VF +3 | | | | | |
| Vehicle vs. A20 (IP Injection) | 1.69 | 0.7077 to 2.672 | Yes | *** | 0.0006 |
| Vehicle vs. A20 (oral gavage) | 1.558 | 0.5752 to 2.540 | Yes | ** | 0.0014 |
| VF +8 | | | | | |
| Vehicle vs. A20 (IP Injection) | 2.205 | 1.223 to 3.187 | Yes | **** | <0.0001 |
| Vehicle vs. A20 (oral gavage) | 2.365 | 1.383 to 3.347 | Yes | **** | <0.0001 |
| VF +10 | | | | | |
| Vehicle vs. A20 (IP Injection) | 0.8875 | -0.09478 to 1.870 | No | ns | 0.0812 |
| Vehicle vs. A20 (oral gavage) | 0.8425 | -0.1398 to 1.825 | No | ns | 0.1011 |
| VF +15 | | | | | |
| Vehicle vs. A20 (IP Injection) | 0 | -0.9823 to 0.9823 | No | ns | >0.9999 |
| Vehicle vs. A20 (oral gavage) | -0.155 | -1.137 to 0.8273 | No | ns | 0.91 |

| Test details | Mean 1 | Mean 2 | Mean diff. | SE of diff. | N1 | N2 | q | DF |
|---|---|---|---|---|---|---|---|---|
| Baseline | | | | | | | | |
| Vehicle vs. A20 (IP Injection) | 1 | 1 | 0 | 0.4301 | 4 | 4 | 0 | 45 |
| Vehicle vs. A20 (oral gavage) | 1 | 1 | 0 | 0.4301 | 4 | 4 | 0 | 45 |
| VF +3 | | | | | | | | |
| Vehicle vs. A20 (IP Injection) | 2.623 | 0.9325 | 1.69 | 0.4301 | 4 | 4 | 3.929 | 45 |
| Vehicle vs. A20 (oral gavage) | 2.623 | 1.065 | 1.558 | 0.4301 | 4 | 4 | 3.621 | 45 |
| VF +8 | | | | | | | | |
| Vehicle vs. A20 (IP Injection) | 3.2 | 0.995 | 2.205 | 0.4301 | 4 | 4 | 5.126 | 45 |
| Vehicle vs. A20 (oral gavage) | 3.2 | 0.835 | 2.365 | 0.4301 | 4 | 4 | 5.498 | 45 |
| VF +10 | | | | | | | | |
| Vehicle vs. A20 (IP Injection) | 1.855 | 0.9675 | 0.8875 | 0.4301 | 4 | 4 | 2.063 | 45 |
| Vehicle vs. A20 (oral gavage) | 1.855 | 1.013 | 0.8425 | 0.4301 | 4 | 4 | 1.959 | 45 |
| VF +15 | | | | | | | | |
| Vehicle vs. A20 (IP Injection) | 0.9625 | 0.9625 | 0 | 0.4301 | 4 | 4 | 0 | 45 |
| Vehicle vs. A20 (oral gavage) | 0.9625 | 1.118 | -0.155 | 0.4301 | 4 | 4 | 0.3603 | 45 |

Fig. 5H- two-way ANOVA (therm)

| | | | | | |
|---|---|---|---|---|---|
| Table Analyzed | Thermal - Injured Paw at 2-4 and 4-6 hr post-Zymosan (seconds) | | | | |
| Two-way RM ANOVA | Matching: Stacked | | | | |
| Assume sphericity? | No | | | | |
| Alpha | 0.05 | | | | |
| Source of Variation | % of total variation | P value | P value summary | Significant? | isser-Greenhouse's epsilon |
| Time x Column Factor | 1.134 | 0.3616 | ns | No | 0.7461 |
| Time | 86.91 | <0.0001 | **** | Yes | 0.7461 |
| Column Factor | 0.5733 | 0.2032 | ns | No | |
| Subject | 2.781 | 0.8328 | ns | No | |
| ANOVA table | SS | DF | MS | F (DFn, DFd) | P value |
| Time x Column Factor | 12.2 | 4 | 3.051 | F (2.984, 25.37) = 1.114 | P=0.3616 |
| Time | 935.7 | 2 | 467.9 | F (1.492, 25.37) = 170.8 | P<0.0001 |
| Column Factor | 6.173 | 2 | 3.086 | F (2, 17) = 1.753 | P=0.2032 |
| Subject | 29.94 | 17 | 1.761 | F (17, 34) = 0.6430 | P=0.8328 |
| Residual | 93.12 | 34 | 2.739 | | |
| Data summary | | | | | |
| Number of columns (Column Factor) | 3 | | | | |
| Number of rows (Time) | 3 | | | | |
| Number of subjects (Subject) | 20 | | | | |
| Number of missing values | 0 | | | | |

Compare cell means with others in its row and its column

| | |
|---|---|
| Number of families | 6 |
| Number of comparisons per row family | 3 |
| Number of comparisons per column family | 3 |
| Alpha | 0.05 |

| Uncorrected Fisher's LSD | Mean diff. | 95.00% CI of diff. | Below threshold? | Summary | Individual P Value |
|---|---|---|---|---|---|
| BL | | | | | |
| +/+ vs. E1/E1 | 0.03095 | -2.368 to 2.430 | No | ns | 0.9773 |
| +/+ vs. E1/+ | -0.2205 | -1.942 to 1.501 | No | ns | 0.7828 |
| E1/E1 vs. E1/+ | -0.2514 | -2.750 to 2.247 | No | ns | 0.8277 |
| 2-4 hr | | | | | |
| +/+ vs. E1/E1 | -1.392 | -4.266 to 1.482 | No | ns | 0.2906 |
| +/+ vs. E1/+ | 0.5224 | -2.327 to 3.371 | No | ns | 0.6735 |
| E1/E1 vs. E1/+ | 1.914 | 0.4606 to 3.368 | Yes | * | 0.0143 |
| 6-8 hr | | | | | |
| +/+ vs. E1/E1 | -0.9683 | -1.839 to -0.09724 | Yes | * | 0.0325 |
| +/+ vs. E1/+ | -1.09 | -1.796 to -0.3836 | Yes | ** | 0.0062 |
| E1/E1 vs. E1/+ | -0.1214 | -0.9550 to 0.7121 | No | ns | 0.7534 |
| +/+ | | | | | |
| BL vs. 2-4 hr | 7.35 | 4.402 to 10.30 | Yes | ** | 0.0014 |
| BL vs. 6-8 hr | 9.885 | 8.750 to 11.02 | Yes | **** | <0.0001 |
| 2-4 hr vs. 6-8 hr | 2.535 | -0.3540 to 5.424 | No | ns | 0.0738 |
| E1/E1 | | | | | |
| BL vs. 2-4 hr | 5.927 | 2.703 to 9.151 | Yes | ** | 0.0041 |
| BL vs. 6-8 hr | 8.886 | 6.381 to 11.39 | Yes | *** | 0.0001 |
| 2-4 hr vs. 6-8 hr | 2.959 | 1.224 to 4.693 | Yes | ** | 0.0059 |
| E1/+ | | | | | |
| BL vs. 2-4 hr | 8.093 | 5.857 to 10.33 | Yes | *** | 0.0001 |
| BL vs. 6-8 hr | 9.016 | 7.430 to 10.60 | Yes | **** | <0.0001 |
| 2-4 hr vs. 6-8 hr | 0.9229 | 0.04184 to 1.804 | Yes | * | 0.0427 |

| Test details | Mean 1 | Mean 2 | Mean diff. | SE of diff. | N1 | N2 | t | DF |
|---|---|---|---|---|---|---|---|---|
| BL | | | | | | | | |
| +/+ vs. E1/E1 | 11.14 | 11.11 | 0.03095 | 1.056 | 6 | 7 | 0.0293 | 8.775 |
| +/+ vs. E1/+ | 11.14 | 11.36 | -0.2205 | 0.7798 | 6 | 7 | 0.2827 | 10.72 |
| E1/E1 vs. E1/+ | 11.11 | 11.36 | -0.2514 | 1.127 | 7 | 7 | 0.2232 | 10.34 |
| 2-4 hr | | | | | | | | |
| +/+ vs. E1/E1 | 3.787 | 5.179 | -1.392 | 1.219 | 6 | 7 | 1.142 | 7.106 |
| +/+ vs. E1/+ | 3.787 | 3.264 | 0.5224 | 1.184 | 6 | 7 | 0.4411 | 6.453 |
| E1/E1 vs. E1/+ | 5.179 | 3.264 | 1.914 | 0.6645 | 7 | 7 | 2.881 | 11.58 |
| 6-8 hr | | | | | | | | |
| +/+ vs. E1/E1 | 1.252 | 2.22 | -0.9683 | 0.3944 | 6 | 7 | 2.455 | 10.7 |
| +/+ vs. E1/+ | 1.252 | 2.341 | -1.09 | 0.3188 | 6 | 7 | 3.418 | 10.45 |
| E1/E1 vs. E1/+ | 2.22 | 2.341 | -0.1214 | 0.3764 | 7 | 7 | 0.3226 | 10.48 |
| +/+ | | | | | | | | |
| BL vs. 2-4 hr | 11.14 | 3.787 | 7.35 | 1.147 | 6 | 6 | 6.409 | 5 |
| BL vs. 6-8 hr | 11.14 | 1.252 | 9.885 | 0.4417 | 6 | 6 | 22.38 | 5 |
| 2-4 hr vs. 6-8 hr | 3.787 | 1.252 | 2.535 | 1.124 | 6 | 6 | 2.256 | 5 |
| E1/E1 | | | | | | | | |
| BL vs. 2-4 hr | 11.11 | 5.179 | 5.927 | 1.317 | 7 | 7 | 4.499 | 6 |
| BL vs. 6-8 hr | 11.11 | 2.22 | 8.886 | 1.024 | 7 | 7 | 8.681 | 6 |
| 2-4 hr vs. 6-8 hr | 5.179 | 2.22 | 2.959 | 0.7088 | 7 | 7 | 4.174 | 6 |
| E1/+ | | | | | | | | |
| BL vs. 2-4 hr | 11.36 | 3.264 | 8.093 | 0.9137 | 7 | 7 | 8.857 | 6 |
| BL vs. 6-8 hr | 11.36 | 2.341 | 9.016 | 0.6482 | 7 | 7 | 13.91 | 6 |
| 2-4 hr vs. 6-8 hr | 3.264 | 2.341 | 0.9229 | 0.3601 | 7 | 7 | 2.563 | 6 |

Fig. 5H- two-way ANOVA (thermR)

| | | | | | |
|---|---|---|---|---|---|
| Table Analyzed | Thermal - Injured Paw Ratio at 2-4 and 4-6 hr post-Zymosan (BL/Zymosan) | | | | |
| Two-way ANOVA | Ordinary | | | | |
| Alpha | 0.05 | | | | |
| Source of Variation | % of total variation | P value | P value summary | Significant? | |
| Interaction | 9.596 | 0.0662 | ns | No | |
| Row Factor | 21.91 | 0.0008 | *** | Yes | |
| Column Factor | 14.93 | 0.0173 | * | Yes | |
| ANOVA table | SS (Type III) | DF | MS | F (DFn, DFd) | P value |
| Interaction | 73.66 | 2 | 36.83 | F (2, 34) = 2.944 | P=0.0662 |
| Row Factor | 168.1 | 1 | 168.1 | F (1, 34) = 13.44 | P=0.0008 |
| Column Factor | 114.6 | 2 | 57.31 | F (2, 34) = 4.581 | P=0.0173 |
| Residual | 425.4 | 34 | 12.51 | | |
| Difference between row means | | | | | |
| Predicted (LS) mean of +2-4 hr | 3.404 | | | | |
| Predicted (LS) mean of +6-8 hr | 7.515 | | | | |
| Difference between predicted means | -4.111 | | | | |
| SE of difference | 1.121 | | | | |
| 95% CI of difference | -6.390 to -1.832 | | | | |
| Data summary | | | | | |
| Number of columns (Column Factor) | 3 | | | | |
| Number of rows (Row Factor) | 2 | | | | |
| Number of values | 40 | | | | |

Within each row, compare columns (simple effects within rows)

| | |
|---|---|
| Number of families | 2 |
| Number of comparisons per family | 3 |
| Alpha | 0.05 |

| Tukey's multiple comparisons test | Predicted (LS) Mean diff. | 95.00% CI of diff. | Below threshold? | Summary | Adjusted P Value |
|---|---|---|---|---|---|
| +2-4 hr | | | | | |
| EphB1 +/+ vs. EphB1 -/- | 1.57 | -3.253 to 6.392 | No | ns | 0.707 |
| EphB1 +/+ vs. EphB1 +/- | 0.1195 | -4.703 to 4.942 | No | ns | 0.998 |
| EphB1 -/- vs. EphB1 +/- | -1.45 | -6.083 to 3.183 | No | ns | 0.7256 |
| +6-8 hr | | | | | |
| EphB1 +/+ vs. EphB1 -/- | 6.242 | 1.420 to 11.06 | Yes | ** | 0.0088 |
| EphB1 +/+ vs. EphB1 +/- | 6.743 | 1.921 to 11.57 | Yes | ** | 0.0045 |
| EphB1 -/- vs. EphB1 +/- | 0.5014 | -4.132 to 5.134 | No | ns | 0.962 |

| Test details | Predicted (LS) Mean 1 | Predicted (LS) Mean 2 | Predicted (LS) Mean diff. | SE of diff. | N1 | N2 | q | DF |
|---|---|---|---|---|---|---|---|---|
| +2-4 hr | | | | | | | | |
| EphB1 +/+ vs. EphB1 -/- | 3.967 | 2.397 | 1.57 | 1.968 | 6 | 7 | 1.128 | 34 |
| EphB1 +/+ vs. EphB1 +/- | 3.967 | 3.847 | 0.1195 | 1.968 | 6 | 7 | 0.0859 | 34 |
| EphB1 -/- vs. EphB1 +/- | 2.397 | 3.847 | -1.45 | 1.891 | 7 | 7 | 1.085 | 34 |
| +6-8 hr | | | | | | | | |
| EphB1 +/+ vs. EphB1 -/- | 11.84 | 5.601 | 6.242 | 1.968 | 6 | 7 | 4.486 | 34 |
| EphB1 +/+ vs. EphB1 +/- | 11.84 | 5.1 | 6.743 | 1.968 | 6 | 7 | 4.846 | 34 |
| EphB1 -/- vs. EphB1 +/- | 5.601 | 5.1 | 0.5014 | 1.891 | 7 | 7 | 0.3751 | 34 |

Fig. 5H - two-way ANOVA (mech)

| | | | | | |
|---|---|---|---|---|---|
| Table Analyzed | Zymosan - VF - Injured Paw | | | | |
| Two-way ANOVA | Ordinary | | | | |
| Alpha | 0.05 | | | | |
| Source of Variation | % of total variation | P value | P value summary | Significant? | |
| Interaction | 9.9 | 0.0115 | * | Yes | |
| Row Factor | 47.22 | <0.0001 | **** | Yes | |
| Column Factor | 11.43 | 0.0063 | ** | Yes | |
| ANOVA table | SS (Type III) | DF | MS | F (DFn, DFd) | P value |
| Interaction | 17.12 | 2 | 8.56 | F (2, 34) = 5.111 | P=0.0115 |
| Row Factor | 81.65 | 1 | 81.65 | F (1, 34) = 48.76 | P<0.0001 |
| Column Factor | 19.77 | 2 | 9.887 | F (2, 34) = 5.904 | P=0.0063 |
| Residual | 56.94 | 34 | 1.675 | | |
| Difference between row means | | | | | |
| Predicted (LS) mean of Baseline | 6.651 | | | | |
| Predicted (LS) mean of 4-6 hr | 3.786 | | | | |
| Difference between predicted means | 2.865 | | | | |
| SE of difference | 0.4103 | | | | |
| 95% CI of difference | 2.031 to 3.699 | | | | |
| Data summary | | | | | |
| Number of columns (Column Factor) | 3 | | | | |
| Number of rows (Row Factor) | 2 | | | | |
| Number of values | 40 | | | | |

Compare cell means regardless of rows and columns

| | |
|---|---|
| Number of families | 1 |
| Number of comparisons per family | 15 |
| Alpha | 0.05 |

| Tukey's multiple comparisons test | Predicted (LS) Mean diff. | 95.00% CI of diff. | Below threshold? | Summary | Adjusted P Value |
|---|---|---|---|---|---|
| Baseline:+/+ vs. Baseline:E1/E1 | -0.2619 | -2.435 to 1.911 | No | ns | 0.9991 |
| Baseline:+/+ vs. Baseline:E1/+ | -0.4405 | -2.614 to 1.733 | No | ns | 0.9894 |
| Baseline:+/+ vs. 4-6 hr:+/+ | 3.917 | 1.662 to 6.172 | Yes | *** | 0.0001 |
| Baseline:+/+ vs. 4-6 hr:E1/E1 | 0.7738 | -1.399 to 2.947 | No | ns | 0.8879 |
| Baseline:+/+ vs. 4-6 hr:E1/+ | 3.202 | 1.029 to 5.375 | Yes | ** | 0.0012 |
| Baseline:E1/E1 vs. Baseline:E1/+ | -0.1786 | -2.266 to 1.909 | No | ns | 0.9998 |
| Baseline:E1/E1 vs. 4-6 hr:+/+ | 4.179 | 2.006 to 6.352 | Yes | **** | <0.0001 |
| Baseline:E1/E1 vs. 4-6 hr:E1/E1 | 1.036 | -1.052 to 3.124 | No | ns | 0.6682 |
| Baseline:E1/E1 vs. 4-6 hr:E1/+ | 3.464 | 1.376 to 5.552 | Yes | *** | 0.0002 |
| Baseline:E1/+ vs. 4-6 hr:+/+ | 4.357 | 2.184 to 6.530 | Yes | **** | <0.0001 |
| Baseline:E1/+ vs. 4-6 hr:E1/E1 | 1.214 | -0.8735 to 3.302 | No | ns | 0.5066 |
| Baseline:E1/+ vs. 4-6 hr:E1/+ | 3.643 | 1.555 to 5.731 | Yes | *** | 0.0001 |
| 4-6 hr:+/+ vs. 4-6 hr:E1/E1 | -3.143 | -5.316 to -0.9698 | Yes | ** | 0.0015 |
| 4-6 hr:+/+ vs. 4-6 hr:E1/+ | -0.7143 | -2.887 to 1.459 | No | ns | 0.9173 |
| 4-6 hr:E1/E1 vs. 4-6 hr:E1/+ | 2.429 | 0.3408 to 4.516 | Yes | * | 0.0149 |

| Test details | Predicted (LS) Mean 1 | Predicted (LS) Mean 2 | Predicted (LS) Mean diff. | SE of diff. | N1 | N2 | q | DF |
|---|---|---|---|---|---|---|---|---|
| Baseline:+/+ vs. Baseline:E1/E1 | 6.417 | 6.679 | -0.2619 | 0.72 | 6 | 7 | 0.5144 | 34 |
| Baseline:+/+ vs. Baseline:E1/+ | 6.417 | 6.857 | -0.4405 | 0.72 | 6 | 7 | 0.8652 | 34 |
| Baseline:+/+ vs. 4-6 hr:+/+ | 6.417 | 2.5 | 3.917 | 0.7472 | 6 | 6 | 7.413 | 34 |
| Baseline:+/+ vs. 4-6 hr:E1/E1 | 6.417 | 5.643 | 0.7738 | 0.72 | 6 | 7 | 1.52 | 34 |
| Baseline:+/+ vs. 4-6 hr:E1/+ | 6.417 | 3.214 | 3.202 | 0.72 | 6 | 7 | 6.29 | 34 |
| Baseline:E1/E1 vs. Baseline:E1/+ | 6.679 | 6.857 | -0.1786 | 0.6917 | 7 | 7 | 0.3651 | 34 |
| Baseline:E1/E1 vs. 4-6 hr:+/+ | 6.679 | 2.5 | 4.179 | 0.72 | 7 | 6 | 8.208 | 34 |
| Baseline:E1/E1 vs. 4-6 hr:E1/E1 | 6.679 | 5.643 | 1.036 | 0.6917 | 7 | 7 | 2.117 | 34 |
| Baseline:E1/E1 vs. 4-6 hr:E1/+ | 6.679 | 3.214 | 3.464 | 0.6917 | 7 | 7 | 7.083 | 34 |
| Baseline:E1/+ vs. 4-6 hr:+/+ | 6.857 | 2.5 | 4.357 | 0.72 | 7 | 6 | 8.559 | 34 |
| Baseline:E1/+ vs. 4-6 hr:E1/E1 | 6.857 | 5.643 | 1.214 | 0.6917 | 7 | 7 | 2.483 | 34 |
| Baseline:E1/+ vs. 4-6 hr:E1/+ | 6.857 | 3.214 | 3.643 | 0.6917 | 7 | 7 | 7.448 | 34 |
| 4-6 hr:+/+ vs. 4-6 hr:E1/E1 | 2.5 | 5.643 | -3.143 | 0.72 | 6 | 7 | 6.173 | 34 |
| 4-6 hr:+/+ vs. 4-6 hr:E1/+ | 2.5 | 3.214 | -0.7143 | 0.72 | 6 | 7 | 1.403 | 34 |
| 4-6 hr:E1/E1 vs. 4-6 hr:E1/+ | 5.643 | 3.214 | 2.429 | 0.6917 | 7 | 7 | 4.965 | 34 |

Fig. 5H - one-way ANOVA (mechR)

| | | | | | |
|---|---|---|---|---|---|
| Table Analyzed | Zymosan - VF - Injured Paw - Ratio | | | | |
| Data sets analyzed | A-D | | | | |
| Distribution assumption | Normal (Gaussian) | | | | |
| ANOVA summary | | | | | |
| F | 3.605 | | | | |
| P value | 0.0314 | | | | |
| P value summary | * | | | | |
| Significant diff. among means (P < 0.05)? | Yes | | | | |
| R squared | 0.351 | | | | |
| Brown-Forsythe test | | | | | |
| F (DFn, DFd) | 0.7170 (3, 20) | | | | |
| P value | 0.5534 | | | | |
| P value summary | ns | | | | |
| Are SDs significantly different (P < 0.05)? | No | | | | |
| Bartlett's test | | | | | |
| Bartlett's statistic (corrected) | 6.186 | | | | |
| P value | 0.1029 | | | | |
| P value summary | ns | | | | |
| Are SDs significantly different (P < 0.05)? | No | | | | |
| ANOVA table | SS | DF | MS | F (DFn, DFd) | P value |
| Treatment (between columns) | 13.77 | 3 | 4.59 | F (3, 20) = 3.605 | P=0.0314 |
| Residual (within columns) | 25.46 | 20 | 1.273 | | |
| Total | 39.24 | 23 | | | |
| Data summary | | | | | |
| Number of treatments (columns) | 4 | | | | |
| Number of values (total) | 24 | | | | |

Fig. 5H - post-hoc - mecRATIO

| | |
|---|---|
| Number of families | 1 |
| Number of comparisons per family | 3 |
| Alpha | 0.05 |

| Dunnett's multiple comparisons test | Mean diff. | 95.00% CI of diff. | Below threshold? | Summary | Adjusted P Value | A-? | |
|---|---|---|---|---|---|---|---|
| +/+ vs. E1/E1 | 1.927 | 0.3305 to 3.523 | Yes | * | 0.0162 | B | E1/E1 |
| +/+ vs. E1/+ | 0.5352 | -1.061 to 2.131 | No | ns | 0.7326 | C | E1/+ |
| +/+ vs. tFB1/tFB1 | 1.297 | -0.5553 to 3.149 | No | ns | 0.2117 | D | tFB1/tFB1 |

| Test details | Mean 1 | Mean 2 | Mean diff. | SE of diff. | n1 | n2 | q | DF |
|---|---|---|---|---|---|---|---|---|
| +/+ vs. E1/E1 | 3.207 | 1.28 | 1.927 | 0.6278 | 6 | 7 | 3.069 | 20 |
| +/+ vs. E1/+ | 3.207 | 2.671 | 0.5352 | 0.6278 | 6 | 7 | 0.8526 | 20 |
| +/+ vs. tFB1/tFB1 | 3.207 | 1.91 | 1.297 | 0.7284 | 6 | 4 | 1.78 | 20 |

Fig. 5I - two-way ANOVA (mech)

| Table Analyzed | Zymosan - VF - Left Paw | | | | |
|---|---|---|---|---|---|
| Two-way ANOVA | Ordinary | | | | |
| Alpha | 0.05 | | | | |
| Source of Variation | % of total variation | P value | P value summary | Significant? | |
| Interaction | 9.501 | 0.0136 | * | Yes | |
| Row Factor | 35.8 | <0.0001 | **** | Yes | |
| Column Factor | 28.07 | 0.0002 | *** | Yes | |
| ANOVA table | SS (Type III) | DF | MS | F (DFn, DFd) | P value |
| Interaction | 8.867 | 1 | 8.867 | F (1, 18) = 7.479 | P=0.0136 |
| Row Factor | 33.41 | 1 | 33.41 | F (1, 18) = 28.18 | P<0.0001 |
| Column Factor | 26.2 | 1 | 26.2 | F (1, 18) = 22.10 | P=0.0002 |
| Residual | 21.34 | 18 | 1.186 | | |
| Difference between column means | | | | | |
| Predicted (LS) mean of +/+ | 3.708 | | | | |
| Predicted (LS) mean of EphrinB2-lacZ/6YFdV | 5.9 | | | | |
| Difference between predicted means | -2.192 | | | | |
| SE of difference | 0.4662 | | | | |
| 95% CI of difference | -3.171 to -1.212 | | | | |
| Difference between row means | | | | | |
| Predicted (LS) mean of Baseline VF | 6.042 | | | | |
| Predicted (LS) mean of VF (+4-6 hr) | 3.567 | | | | |
| Difference between predicted means | 2.475 | | | | |
| SE of difference | 0.4662 | | | | |
| 95% CI of difference | 1.495 to 3.455 | | | | |
| Interaction CI | | | | | |
| Mean diff, A1 - B1 | -0.9167 | | | | |
| Mean diff, A2 - B2 | -3.467 | | | | |
| (A1 -B1) - (A2 - B2) | 2.55 | | | | |
| 95% CI of difference | 0.5910 to 4.509 | | | | |
| (B1 - A1) - (B2 - A2) | -2.55 | | | | |
| 95% CI of difference | -4.509 to -0.5910 | | | | |
| Data summary | | | | | |
| Number of columns (Column Factor) | 2 | | | | |
| Number of rows (Row Factor) | 2 | | | | |
| Number of values | 22 | | | | |

Compare cell means regardless of rows and columns

| | |
|---|---|
| Number of families | 1 |
| Number of comparisons per family | 6 |
| Alpha | 0.05 |

| Tukey's multiple comparisons test | Predicted (LS) Mean diff. | 95.00% CI of diff. | Below threshold? | Summary | Adjusted P Value |
|---|---|---|---|---|---|
| Baseline:+/+ vs. Baseline:EphrinB2-lacZ/6YI | -0.9167 | -2.780 to 0.9468 | No | ns | 0.521 |
| Baseline:+/+ vs. Zymosan:+/+ | 3.75 | 1.973 to 5.527 | Yes | **** | <0.0001 |
| Baseline:+/+ vs. Zymosan:EphrinB2-lacZ/6Y | 0.2833 | -1.580 to 2.147 | No | ns | 0.9726 |
| Baseline:EphrinB2-lacZ/6YFdV vs. Zymosar | 4.667 | 2.803 to 6.530 | Yes | **** | <0.0001 |
| Baseline:EphrinB2-lacZ/6YFdV vs. Zymosar | 1.2 | -0.7464 to 3.146 | No | ns | 0.3319 |
| Zymosan:+/+ vs. Zymosan:EphrinB2-lacZ/6Y | -3.467 | -5.330 to -1.603 | Yes | *** | 0.0003 |

| Test details | Predicted (LS) Mean 1 | Predicted (LS) Mean 2 | Predicted (LS) Mean diff. | SE of diff. | N1 | N2 | q | DF |
|---|---|---|---|---|---|---|---|---|
| Baseline:+/+ vs. Baseline:EphrinB2-lacZ/6YI | 5.583 | 6.5 | -0.9167 | 0.6593 | 6 | 5 | 1.966 | 18 |
| Baseline:+/+ vs. Zymosan:+/+ | 5.583 | 1.833 | 3.75 | 0.6287 | 6 | 6 | 8.436 | 18 |
| Baseline:+/+ vs. Zymosan:EphrinB2-lacZ/6Y | 5.583 | 5.3 | 0.2833 | 0.6593 | 6 | 5 | 0.6077 | 18 |
| Baseline:EphrinB2-lacZ/6YFdV vs. Zymosar | 6.5 | 1.833 | 4.667 | 0.6593 | 5 | 6 | 10.01 | 18 |
| Baseline:EphrinB2-lacZ/6YFdV vs. Zymosar | 6.5 | 5.3 | 1.2 | 0.6887 | 5 | 5 | 2.464 | 18 |
| Zymosan:+/+ vs. Zymosan:EphrinB2-lacZ/6Y | 1.833 | 5.3 | -3.467 | 0.6593 | 6 | 5 | 7.436 | 18 |

Fig. 5I - t-test - mec

| | |
|---|---|
| Table Analyzed | Zymosan - VF - Left Paw - Ratio |
| Column B | EphrinB2-lacZ/6YFdV |
| vs. | vs. |
| Column A | +/+ |
| Unpaired t test | |
| P value | 0.0039 |
| P value summary | ** |
| Significantly different (P < 0.05)? | Yes |
| One- or two-tailed P value? | Two-tailed |
| t, df | t=3.845, df=9 |
| How large is the effect? | |
| Mean of column A | 3.403 |
| Mean of column B | 1.288 |
| Difference between means (B - A) ± SEM | -2.115 ± 0.5501 |
| 95% confidence interval | -3.360 to -0.8709 |
| R squared (eta squared) | 0.6216 |
| F test to compare variances | |
| F, DFn, Dfd | 4.833, 5, 4 |
| P value | 0.1521 |
| P value summary | ns |
| Significantly different (P < 0.05)? | No |
| Data analyzed | |
| Sample size, column A | 6 |
| Sample size, column B | 5 |

Fig. 5J - two-way ANOVA (therm)

| | | | | | |
|---|---|---|---|---|---|
| Table Analyzed | Thermal - Injured Paw (seconds) | | | | |
| Two-way ANOVA | Ordinary | | | | |
| Alpha | 0.05 | | | | |
| Source of Variation | % of total variation | P value | P value summary | Significant? | |
| Interaction | 6.221 | 0.0176 | * | Yes | |
| Row Factor | 83.25 | <0.0001 | **** | Yes | |
| Column Factor | 4.061 | 0.0538 | ns | No | |
| ANOVA table | SS | DF | MS | F (DFn, DFd) | P value |
| Interaction | 20.89 | 2 | 10.44 | F (2, 12) = 5.768 | P=0.0176 |
| Row Factor | 279.5 | 1 | 279.5 | F (1, 12) = 154.4 | P<0.0001 |
| Column Factor | 13.63 | 2 | 6.817 | F (2, 12) = 3.765 | P=0.0538 |
| Residual | 21.73 | 12 | 1.811 | | |
| Difference between row means | | | | | |
| Mean of Baseline | 11.98 | | | | |
| Mean of 4-6 hr | 4.103 | | | | |
| Difference between means | 7.881 | | | | |
| SE of difference | 0.6343 | | | | |
| 95% CI of difference | 6.499 to 9.263 | | | | |
| Data summary | | | | | |
| Number of columns (Column Factor) | 3 | | | | |
| Number of rows (Row Factor) | 2 | | | | |
| Number of values | 18 | | | | |

Fig. 5J - post-hoc - thermal

Within each row, compare columns (simple effects within rows)

| | |
|---|---|
| Number of families | 2 |
| Number of comparisons per family | 2 |
| Alpha | 0.05 |

| Dunnett's multiple comparisons test | Mean diff. | 95.00% CI of diff. | Below threshold? | Summary | Adjusted P Value |
|---|---|---|---|---|---|
| Baseline | | | | | |
| Vehicle vs. 3511-I | -0.1333 | -2.883 to 2.616 | No | ns | 0.9893 |
| Vehicle vs. BQPB4 | 0.59 | -2.159 to 3.339 | No | ns | 0.8158 |
| 4-6 hr | | | | | |
| Vehicle vs. 3511-I | -2.947 | -5.696 to -0.1972 | Yes | * | 0.0362 |
| Vehicle vs. BQPB4 | -4.683 | -7.433 to -1.934 | Yes | ** | 0.0021 |

| Test details | Mean 1 | Mean 2 | Mean diff. | SE of diff. | N1 | N2 | q | DF |
|---|---|---|---|---|---|---|---|---|
| Baseline | | | | | | | | |
| Vehicle vs. 3511-I | 12.14 | 12.27 | -0.1333 | 1.099 | 3 | 3 | 0.1214 | 12 |
| Vehicle vs. BQPB4 | 12.14 | 11.55 | 0.59 | 1.099 | 3 | 3 | 0.537 | 12 |
| 4-6 hr | | | | | | | | |
| Vehicle vs. 3511-I | 1.56 | 4.507 | -2.947 | 1.099 | 3 | 3 | 2.682 | 12 |
| Vehicle vs. BQPB4 | 1.56 | 6.243 | -4.683 | 1.099 | 3 | 3 | 4.263 | 12 |

Fig. 5J - one-way ANOVA tRatio

| | | | | | |
|---|---|---|---|---|---|
| Table Analyzed | Thermal - Injured Paw Ratio (BL/Zymosan) | | | | |
| Data sets analyzed | A, C, D | | | | |
| Distribution assumption | Normal (Gaussian) | | | | |
| ANOVA summary | | | | | |
| F | 14.11 | | | | |
| P value | 0.0054 | | | | |
| P value summary | ** | | | | |
| Significant diff. among means (P < 0.05)? | Yes | | | | |
| R squared | 0.8247 | | | | |
| Brown-Forsythe test | | | | | |
| F (DFn, DFd) | 0.7626 (2, 6) | | | | |
| P value | 0.5069 | | | | |
| P value summary | ns | | | | |
| Are SDs significantly different (P < 0.05)? | No | | | | |
| Bartlett's test | | | | | |
| Bartlett's statistic (corrected) | | | | | |
| P value | | | | | |
| P value summary | | | | | |
| Are SDs significantly different (P < 0.05)? | | | | | |
| ANOVA table | SS | DF | MS | F (DFn, DFd) | P value |
| Treatment (between columns) | 59.03 | 2 | 29.51 | F (2, 6) = 14.11 | P=0.0054 |
| Residual (within columns) | 12.55 | 6 | 2.092 | | |
| Total | 71.58 | 8 | | | |
| Data summary | | | | | |
| Number of treatments (columns) | 3 | | | | |
| Number of values (total) | 9 | | | | |

Fig. 5J - post-hoc - tRatio

| | |
|---|---|
| Number of families | 1 |
| Number of comparisons per family | 2 |
| Alpha | 0.05 |

| Dunnett's multiple comparisons test | Mean diff. | 95.00% CI of diff. | Below threshold? | Summary | Adjusted P Value | A-? | |
|---|---|---|---|---|---|---|---|
| Vehicle vs. 3511-I | 4.47 | 1.089 to 7.851 | Yes | * | 0.0162 | C | 3511-I |
| Vehicle vs. BQPB4 | 6.047 | 2.666 to 9.427 | Yes | ** | 0.0039 | D | BQPB4 |

| Test details | Mean 1 | Mean 2 | Mean diff. | SE of diff. | n1 | n2 | q | DF |
|---|---|---|---|---|---|---|---|---|
| Vehicle vs. 3511-I | 7.937 | 3.467 | 4.47 | 1.181 | 3 | 3 | 3.785 | 6 |
| Vehicle vs. BQPB4 | 7.937 | 1.89 | 6.047 | 1.181 | 3 | 3 | 5.121 | 6 |

Fig. 5J - two-way ANOVA (mech)

| | | | | | |
|---|---|---|---|---|---|
| Table Analyzed | VF - Injured Paw | | | | |
| Two-way ANOVA | Ordinary | | | | |
| Alpha | 0.05 | | | | |
| Source of Variation | % of total variation | P value | P value summary | Significant? | |
| Interaction | 13.9 | 0.2979 | ns | No | |
| Row Factor | 24.33 | 0.0236 | * | Yes | |
| Column Factor | 14.62 | 0.088 | ns | No | |
| ANOVA table | SS | DF | MS | F (DFn, DFd) | P value |
| Interaction | 16.22 | 4 | 4.056 | F (4, 18) = 1.327 | P=0.2979 |
| Row Factor | 28.39 | 2 | 14.19 | F (2, 18) = 4.645 | P=0.0236 |
| Column Factor | 17.06 | 2 | 8.528 | F (2, 18) = 2.791 | P=0.0880 |
| Residual | 55 | 18 | 3.056 | | |
| Data summary | | | | | |
| Number of columns (Column Factor) | 3 | | | | |
| Number of rows (Row Factor) | 3 | | | | |
| Number of values | 27 | | | | |

Within each column, compare rows (simple effects within columns)

| | |
|---|---|
| Number of families | 3 |
| Number of comparisons per family | 2 |
| Alpha | 0.05 |

| Dunnett's multiple comparisons test | Mean diff. | 95.00% CI of diff. | Below threshold? | Summary | Adjusted P Value |
|---|---|---|---|---|---|
| Vehicle | | | | | |
| Baseline vs. 2-4 hr | 2.833 | -0.5902 to 6.257 | No | ns | 0.1107 |
| Baseline vs. 6-8 hr | 3.833 | 0.4098 to 7.257 | Yes | ** | 0.0278 |
| 3511-I | | | | | |
| Baseline vs. 2-4 hr | 3.667 | 0.2431 to 7.090 | Yes | * | 0.0354 |
| Baseline vs. 6-8 hr | 2.333 | -1.090 to 5.757 | No | ns | 0.2045 |
| BQPB4 | | | | | |
| Baseline vs. 2-4 hr | 0.3333 | -3.090 to 3.757 | No | ns | 0.961 |
| Baseline vs. 6-8 hr | 0 | -3.424 to 3.424 | No | ns | >0.9999 |

| Test details | Mean 1 | Mean 2 | Mean diff. | SE of diff. | N1 | N2 | q | DF |
|---|---|---|---|---|---|---|---|---|
| Vehicle | | | | | | | | |
| Baseline vs. 2-4 hr | 6.333 | 3.5 | 2.833 | 1.427 | 3 | 3 | 1.985 | 18 |
| Baseline vs. 6-8 hr | 6.333 | 2.5 | 3.833 | 1.427 | 3 | 3 | 2.686 | 18 |
| 3511-I | | | | | | | | |
| Baseline vs. 2-4 hr | 7.167 | 3.5 | 3.667 | 1.427 | 3 | 3 | 2.569 | 18 |
| Baseline vs. 6-8 hr | 7.167 | 4.833 | 2.333 | 1.427 | 3 | 3 | 1.635 | 18 |
| BQPB4 | | | | | | | | |
| Baseline vs. 2-4 hr | 6.167 | 5.833 | 0.3333 | 1.427 | 3 | 3 | 0.2335 | 18 |
| Baseline vs. 6-8 hr | 6.167 | 6.167 | 0 | 1.427 | 3 | 3 | 0 | 18 |

Fig. 5J - one-way ANOVA mRatio

| | | | | | |
|---|---|---|---|---|---|
| Table Analyzed | VF - Pain Ratio separate columns (BL/Zymosan) | | | | |
| Data sets analyzed | E, G, H | | | | |
| ANOVA summary | | | | | |
| F | 5.887 | | | | |
| P value | 0.0385 | | | | |
| P value summary | * | | | | |
| Significant diff. among means (P < 0.05)? | Yes | | | | |
| R squared | 0.6624 | | | | |
| Brown-Forsythe test | | | | | |
| F (DFn, DFd) | 2.567 (2, 6) | | | | |
| P value | 0.1565 | | | | |
| P value summary | ns | | | | |
| Are SDs significantly different (P < 0.05)? | No | | | | |
| Bartlett's test | | | | | |
| Bartlett's statistic (corrected) | | | | | |
| P value | | | | | |
| P value summary | | | | | |
| Are SDs significantly different (P < 0.05)? | | | | | |
| ANOVA table | SS | DF | MS | F (DFn, DFd) | P value |
| Treatment (between columns) | 7.277 | 2 | 3.639 | F (2, 6) = 5.887 | P=0.0385 |
| Residual (within columns) | 3.708 | 6 | 0.618 | | |
| Total | 10.99 | 8 | | | |
| Data summary | | | | | |
| Number of treatments (columns) | 3 | | | | |
| Number of values (total) | 9 | | | | |

Fig. 5J - post-hoc - mRatio

| | |
|---|---|
| Number of families | 1 |
| Number of comparisons per family | 2 |
| Alpha | 0.05 |

| Dunnett's multiple comparisons test | Mean Diff. | 95.00% CI of diff. | Below threshold? | Summary | Adjusted P Value | E-? |
|---|---|---|---|---|---|---|
| Vehicle (6-8 hr) vs. 3511-I (6-8 hr) | 1.64 | -0.1978 to 3.478 | No | ns | 0.0746 | G 1-I (6-8 hr) |
| Vehicle (6-8 hr) vs. BQPB4 (6-8 hr) | 2.093 | 0.2556 to 3.931 | Yes | * | 0.0303 | H B4 (6-8 hr) |

| Test details | Mean 1 | Mean 2 | Mean Diff. | SE of diff. | n1 | n2 | q | DF |
|---|---|---|---|---|---|---|---|---|
| Vehicle (6-8 hr) vs. 3511-I (6-8 hr) | 3.11 | 1.47 | 1.64 | 0.6419 | 3 | 3 | 2.555 | 6 |
| Vehicle (6-8 hr) vs. BQPB4 (6-8 hr) | 3.11 | 1.017 | 2.093 | 0.6419 | 3 | 3 | 3.261 | 6 |

Fig. 5K - two-way ANOVA (therm)

| | | | | | |
|---|---|---|---|---|---|
| Table Analyzed | Bar Graph- Thermal - Left Paw (seconds) | | | | |
| Two-way ANOVA | Ordinary | | | | |
| Alpha | 0.05 | | | | |
| Source of Variation | % of total variation | P value | P value summary | Significant? | |
| Interaction | 3.77 | 0.015 | * | Yes | |
| Row Factor | 69.29 | <0.0001 | **** | Yes | |
| Column Factor | 3.286 | 0.0253 | * | Yes | |
| ANOVA table | SS (Type III) | DF | MS | F (DFn, DFd) | P value |
| Interaction | 21.84 | 3 | 7.281 | F (3, 28) = 4.144 | P=0.0150 |
| Row Factor | 401.5 | 1 | 401.5 | F (1, 28) = 228.5 | P<0.0001 |
| Column Factor | 19.04 | 3 | 6.347 | F (3, 28) = 3.613 | P=0.0253 |
| Residual | 49.19 | 28 | 1.757 | | |
| Difference between row means | | | | | |
| Predicted (LS) mean of Baseline | 11.09 | | | | |
| Predicted (LS) mean of 4-6 hr | 4.009 | | | | |
| Difference between predicted means | 7.084 | | | | |
| SE of difference | 0.4686 | | | | |
| 95% CI of difference | 6.124 to 8.044 | | | | |
| Data summary | | | | | |
| Number of columns (Column Factor) | 4 | | | | |
| Number of rows (Row Factor) | 2 | | | | |
| Number of values | 36 | | | | |

Within each row, compare columns (simple effects within rows)

| | |
|---|---|
| Number of families | 2 |
| Number of comparisons per family | 3 |
| Alpha | 0.05 |

| Dunnett's multiple comparisons test | Predicted (LS) Mean diff. | 95.00% CI of diff. | Below threshold? | Summary | Adjusted P Value |
|---|---|---|---|---|---|
| Baseline | | | | | |
| Vehicle vs. BQPB4 (20 mg/kg, PO) | -0.1567 | -2.073 to 1.760 | No | ns | 0.9947 |
| Vehicle vs. BQPB4 (10 mg/kg, PO) | 0.8167 | -1.531 to 3.164 | No | ns | 0.7383 |
| Vehicle vs. BQPB4 (5 mg/kg, PO) | 0.8467 | -1.501 to 3.194 | No | ns | 0.7175 |
| 4-6 hr | | | | | |
| Vehicle vs. BQPB4 (20 mg/kg, PO) | -3.37 | -5.286 to -1.454 | Yes | *** | 0.0004 |
| Vehicle vs. BQPB4 (10 mg/kg, PO) | -2.85 | -5.197 to -0.5028 | Yes | * | 0.0143 |
| Vehicle vs. BQPB4 (5 mg/kg, PO) | -2.177 | -4.524 to 0.1705 | No | ns | 0.0742 |

| Test details | Predicted (LS) Mean 1 | Predicted (LS) Mean 2 | Predicted (LS) Mean diff. | SE of diff. | N1 | N2 | q |
|---|---|---|---|---|---|---|---|
| Baseline | | | | | | | |
| Vehicle vs. BQPB4 (20 mg/kg, PO) | 11.47 | 11.63 | -0.1567 | 0.7652 | 6 | 6 | 0.2047 |
| Vehicle vs. BQPB4 (10 mg/kg, PO) | 11.47 | 10.65 | 0.8167 | 0.9372 | 6 | 3 | 0.8714 |
| Vehicle vs. BQPB4 (5 mg/kg, PO) | 11.47 | 10.62 | 0.8467 | 0.9372 | 6 | 3 | 0.9034 |
| 4-6 hr | | | | | | | |
| Vehicle vs. BQPB4 (20 mg/kg, PO) | 1.91 | 5.28 | -3.37 | 0.7652 | 6 | 6 | 4.404 |
| Vehicle vs. BQPB4 (10 mg/kg, PO) | 1.91 | 4.76 | -2.85 | 0.9372 | 6 | 3 | 3.041 |
| Vehicle vs. BQPB4 (5 mg/kg, PO) | 1.91 | 4.087 | -2.177 | 0.9372 | 6 | 3 | 2.322 |

Fig. 5K - one-way ANOVA tRatio

| | | | | | |
|---|---|---|---|---|---|
| Table Analyzed | Bar Graph- Thermal - Left Paw Ratio (BL/Zymosan) | | | | |
| Data sets analyzed | A-D | | | | |
| Distribution assumption | Normal (Gaussian) | | | | |
| ANOVA summary | | | | | |
| F | 8.614 | | | | |
| P value | 0.0017 | | | | |
| P value summary | ** | | | | |
| Significant diff. among means (P < 0.05)? | Yes | | | | |
| R squared | 0.6486 | | | | |
| Brown-Forsythe test | | | | | |
| F (DFn, DFd) | 1.321 (3, 14) | | | | |
| P value | 0.307 | | | | |
| P value summary | ns | | | | |
| Are SDs significantly different (P < 0.05)? | No | | | | |
| Bartlett's test | | | | | |
| Bartlett's statistic (corrected) | | | | | |
| P value | | | | | |
| P value summary | | | | | |
| Are SDs significantly different (P < 0.05)? | | | | | |
| ANOVA table | SS | DF | MS | F (DFn, DFd) | P value |
| Treatment (between columns) | 55.16 | 3 | 18.39 | F (3, 14) = 8.614 | P=0.0017 |
| Residual (within columns) | 29.88 | 14 | 2.134 | | |
| Total | 85.04 | 17 | | | |
| Data summary | | | | | |
| Number of treatments (columns) | 4 | | | | |
| Number of values (total) | 18 | | | | |

Fig. 5K - post-hoc - tRatio

| | |
|---|---|
| Number of families | 1 |
| Number of comparisons per family | 3 |
| Alpha | 0.05 |

| Dunnett's multiple comparisons test | Mean diff. | 95.00% CI of diff. | Below threshold? | Summary | Adjusted P Value | A-? |
|---|---|---|---|---|---|---|
| Vehicle vs. BQPB4 (20 mg/kg, PO) | 3.93 | 1.689 to 6.171 | Yes | ** | 0.001 | B ng/kg, PO) |
| Vehicle vs. BQPB4 (10 mg/kg, PO) | 3.735 | 0.9908 to 6.479 | Yes | ** | 0.0078 | C ng/kg, PO) |
| Vehicle vs. BQPB4 (5 mg/kg, PO) | 3.042 | 0.2974 to 5.786 | Yes | * | 0.0289 | D ng/kg, PO) |

| Test details | Mean 1 | Mean 2 | Mean diff. | SE of diff. | n1 | n2 | q | DF |
|---|---|---|---|---|---|---|---|---|
| Vehicle vs. BQPB4 (20 mg/kg, PO) | 6.362 | 2.432 | 3.93 | 0.8435 | 6 | 6 | 4.659 | 14 |
| Vehicle vs. BQPB4 (10 mg/kg, PO) | 6.362 | 2.627 | 3.735 | 1.033 | 6 | 3 | 3.615 | 14 |
| Vehicle vs. BQPB4 (5 mg/kg, PO) | 6.362 | 3.32 | 3.042 | 1.033 | 6 | 3 | 2.944 | 14 |

Fig. 5K - t-test - tRatio

| | |
|---|---|
| Table Analyzed | Bar Graph- Thermal - Left Paw Ratio (BL/Zymosan) (BQPB4 pooled) |
| Column B | BQPB4 (20-10-5 mg/kg, PO) |
| vs. | vs. |
| Column A | Vehicle |
| Unpaired t test | |
| P value | <0.0001 |
| P value summary | **** |
| Significantly different (P < 0.05)? | Yes |
| One- or two-tailed P value? | Two-tailed |
| t, df | t=5.217, df=16 |
| How large is the effect? | |
| Mean of column A | 6.362 |
| Mean of column B | 2.703 |
| Difference between means (B - A) ± SEM | -3.659 ± 0.7014 |
| 95% confidence interval | -5.146 to -2.172 |
| R squared (eta squared) | 0.6298 |
| F test to compare variances | |
| F, DFn, Dfd | 2.748, 5, 11 |
| P value | 0.1508 |
| P value summary | ns |
| Significantly different (P < 0.05)? | No |
| Data analyzed | |
| Sample size, column A | 6 |
| Sample size, column B | 12 |

### Table S7: Statistics for Figure 6.

Fig. 6A - t-test Results

| | |
|---|---|
| Table Analyzed | CFA 2+3 - DH Tom Neuron Ratios in WT, E1/+, and E1/E1 Mut |
| Column B | EphB1 +/- |
| vs. | vs. |
| Column A | EphB1 +/+ |
| Unpaired t test | |
| P value | 0.0094 |
| P value summary | ** |
| Significantly different (P < 0.05)? | Yes |
| One- or two-tailed P value? | Two-tailed |
| t, df | t=3.764, df=6 |
| How large is the effect? | |
| Mean of column A | 2.845 |
| Mean of column B | 1.98 |
| Difference between means (B - A) ± SEM | -0.8650 ± 0.2298 |
| 95% confidence interval | -1.427 to -0.3027 |
| R squared (eta squared) | 0.7025 |
| F test to compare variances | |
| F, DFn, Dfd | 11.23, 3, 3 |
| P value | 0.0774 |
| P value summary | ns |
| Significantly different (P < 0.05)? | No |
| Data analyzed | |
| Sample size, column A | 4 |
| Sample size, column B | 4 |

Fig. 6B - two-way ANOVA by sect

| | | | | | |
|---|---|---|---|---|---|
| Table Analyzed | CFA 20, 3, 17 - all sections analyzed | | | | |
| Two-way ANOVA | Ordinary | | | | |
| Alpha | 0.05 | | | | |
| Source of Variation | % of total variation | P value | P value summary | Significant? | |
| Row Factor | 24.51 | 0.2051 | ns | No | |
| Column Factor | 36.18 | <0.0001 | **** | Yes | |
| ANOVA table | SS (Type III) | DF | MS | F (DFn, DFd) | P value |
| Row Factor | 188.6 | 57 | 3.308 | F (57, 99) = 1.207 | P=0.2051 |
| Column Factor | 278.4 | 3 | 92.8 | F (3, 99) = 33.84 | P<0.0001 |
| Residual | 271.5 | 99 | 2.742 | | |
| Data summary | | | | | |
| Number of columns (Column Factor) | 4 | | | | |
| Number of rows (Row Factor) | 58 | | | | |
| Number of values | 160 | | | | |

Fig. 6B - post hoc by section

Compare column means (main column effect)

| | |
|---|---|
| Number of families | 1 |
| Number of comparisons per family | 6 |
| Alpha | 0.05 |

| Tukey's multiple comparisons test | Mean diff. | 95.00% CI of diff. | Below threshold? | Summary | Adjusted P Value |
|---|---|---|---|---|---|
| +/+ vs. +/- | 3.399 | 2.409 to 4.390 | Yes | **** | <0.0001 |
| +/+ vs. -/- | 3.074 | 2.040 to 4.107 | Yes | **** | <0.0001 |
| +/+ vs. +/+ (no CFA) | 3.415 | 2.235 to 4.596 | Yes | **** | <0.0001 |
| +/- vs. -/- | -0.3258 | -1.255 to 0.6036 | No | ns | 0.7963 |
| +/- vs. +/+ (no CFA) | 0.016 | -1.139 to 1.171 | No | ns | >0.9999 |
| -/- vs. +/+ (no CFA) | 0.3418 | -0.8466 to 1.530 | No | ns | 0.8758 |

| Test details | Mean 1 | Mean 2 | Mean diff. | SE of diff. | N1 | N2 | q | DF |
|---|---|---|---|---|---|---|---|---|
| +/+ vs. +/- | 5.117 | 1.718 | 3.399 | 0.379 | 36 | 48 | 12.68 | 99 |
| +/+ vs. -/- | 5.117 | 2.044 | 3.074 | 0.3956 | 36 | 52 | 10.99 | 99 |
| +/+ vs. +/+ (no CFA) | 5.117 | 1.702 | 3.415 | 0.4517 | 36 | 24 | 10.69 | 99 |
| +/- vs. -/- | 1.718 | 2.044 | -0.3258 | 0.3557 | 48 | 52 | 1.296 | 99 |
| +/- vs. +/+ (no CFA) | 1.718 | 1.702 | 0.016 | 0.4418 | 48 | 24 | 0.0512 | 99 |
| -/- vs. +/+ (no CFA) | 2.044 | 1.702 | 0.3418 | 0.4548 | 52 | 24 | 1.063 | 99 |

Fig. 6B - two-way ANOVA by mous

| | | | | | |
|---|---|---|---|---|---|
| Table Analyzed | CFA 20, 3, 17 - grouped by mouse | | | | |
| Two-way ANOVA | Ordinary | | | | |
| Alpha | 0.05 | | | | |
| Source of Variation | % of total variation | P value | P value summary | Significant? | |
| Row Factor | 9.035 | 0.1188 | ns | No | |
| Column Factor | 85.89 | 0.0001 | *** | Yes | |
| ANOVA table | SS (Type III) | DF | MS | F (DFn, DFd) | P value |
| Row Factor | 1.289 | 4 | 0.3224 | F (4, 7) = 2.704 | P=0.1188 |
| Column Factor | 12.26 | 3 | 4.086 | F (3, 7) = 34.28 | P=0.0001 |
| Residual | 0.8344 | 7 | 0.1192 | | |
| Data summary | | | | | |
| Number of columns (Column Factor) | 4 | | | | |
| Number of rows (Row Factor) | 5 | | | | |
| Number of values | 15 | | | | |

Fig. 6B - post hoc by mouse

Compare column means (main column effect)

| | |
|---|---|
| Number of families | 1 |
| Number of comparisons per family | 6 |
| Alpha | 0.05 |

| Tukey's multiple comparisons test | Mean diff. | 95.00% CI of diff. | Below threshold? | Summary | \djusted P Value |
|---|---|---|---|---|---|
| +/+ vs. +/- | 1.791 | 0.9829 to 2.599 | Yes | *** | 0.0007 |
| +/+ vs. -/- | 1.981 | 1.173 to 2.789 | Yes | *** | 0.0004 |
| +/+ vs. +/+ (no CFA) | 2.599 | 1.556 to 3.642 | Yes | *** | 0.0003 |
| +/- vs. -/- | 0.1903 | -0.6178 to 0.9984 | No | ns | 0.8615 |
| +/- vs. +/+ (no CFA) | 0.8079 | -0.2354 to 1.851 | No | ns | 0.1328 |
| -/- vs. +/+ (no CFA) | 0.6176 | -0.4257 to 1.661 | No | ns | 0.2872 |

| Test details | Mean 1 | Mean 2 | Mean diff. | SE of diff. | N1 | N2 | q | DF |
|---|---|---|---|---|---|---|---|---|
| +/+ vs. +/- | 3.388 | 1.597 | 1.791 | 0.2441 | 4 | 4 | 10.37 | 7 |
| +/+ vs. -/- | 3.388 | 1.406 | 1.981 | 0.2441 | 4 | 5 | 11.48 | 7 |
| +/+ vs. +/+ (no CFA) | 3.388 | 0.7889 | 2.599 | 0.3152 | 4 | 2 | 11.66 | 7 |
| +/- vs. -/- | 1.597 | 1.406 | 0.1903 | 0.2441 | 4 | 5 | 1.102 | 7 |
| +/- vs. +/+ (no CFA) | 1.597 | 0.7889 | 0.8079 | 0.3152 | 4 | 2 | 3.625 | 7 |
| -/- vs. +/+ (no CFA) | 1.406 | 0.7889 | 0.6176 | 0.3152 | 5 | 2 | 2.771 | 7 |

Fig. 6C - one-way ANOVA Results

| | | | | | |
|---|---|---|---|---|---|
| Table Analyzed | CFA Test 2 - TOM Neuron Counts in DH in V, A20, and MCD | | | | |
| Data sets analyzed | A-C | | | | |
| Distribution assumption | Normal (Gaussian) | | | | |
| ANOVA summary | | | | | |
| F | 3.39 | | | | |
| P value | 0.0934 | | | | |
| P value summary | ns | | | | |
| Significant diff. among means (P < 0.05)? | No | | | | |
| R squared | 0.492 | | | | |
| Brown-Forsythe test | | | | | |
| F (DFn, DFd) | 0.4735 (2, 7) | | | | |
| P value | 0.6414 | | | | |
| P value summary | ns | | | | |
| Are SDs significantly different (P < 0.05)? | No | | | | |
| Bartlett's test | | | | | |
| Bartlett's statistic (corrected) | | | | | |
| P value | | | | | |
| P value summary | | | | | |
| Are SDs significantly different (P < 0.05)? | | | | | |
| ANOVA table | SS | DF | MS | F (DFn, DFd) | P value |
| Treatment (between columns) | 2.644 | 2 | 1.322 | F (2, 7) = 3.390 | P=0.0934 |
| Residual (within columns) | 2.73 | 7 | 0.39 | | |
| Total | 5.375 | 9 | | | |
| Data summary | | | | | |
| Number of treatments (columns) | 3 | | | | |
| Number of values (total) | 10 | | | | |

Fig. 6C - post-hoc

| | | | | | | | | |
|---|---|---|---|---|---|---|---|---|
| Number of families | 1 | | | | | | | |
| Number of comparisons per family | 1 | | | | | | | |
| Alpha | 0.05 | | | | | | | |
| Bonferroni's multiple comparisons test | Mean diff. | 95.00% CI of diff. | Below threshold? | Summary | Adjusted P Value | | | |
| Vehicle vs. A20 | 1.355 | 0.006875 to 2.703 | Yes | * | 0.0491 | A-B | | |
| Test details | Mean 1 | Mean 2 | Mean diff. | SE of diff. | n1 | n2 | t | DF |
| Vehicle vs. A20 | 2.92 | 1.565 | 1.355 | 0.5701 | 3 | 2 | 2.377 | 7 |

Fig. 6C - t-test

| | |
|---|---|
| Table Analyzed | CFA Test 2 - TOM Neuron Counts in DH in V, A20, and MCD |
| Column B | A20 |
| vs. | vs. |
| Column A | Vehicle |
| Unpaired t test with Welch's correction | |
| P value | 0.0322 |
| P value summary | * |
| Significantly different (P < 0.05)? | Yes |
| One- or two-tailed P value? | Two-tailed |
| Welch-corrected t, df | t=3.791, df=3.000 |
| How large is the effect? | |
| Mean of column A | 2.92 |
| Mean of column B | 1.565 |
| Difference between means (B - A) ± SEM | -1.355 ± 0.3574 |
| 95% confidence interval | -2.493 to -0.2174 |
| R squared (eta squared) | 0.8273 |
| F test to compare variances | |
| F, DFn, Dfd | 3.060, 2, 1 |
| P value | 0.7495 |
| P value summary | ns |
| Significantly different (P < 0.05)? | No |
| Data analyzed | |
| Sample size, column A | 3 |
| Sample size, column B | 2 |

Fig. 6D - one-way ANOVA Results

| | | | | | |
|---|---|---|---|---|---|
| Table Analyzed | CFA 18 - Tom+ neuron ratio | | | | |
| Data sets analyzed | A-C | | | | |
| Distribution assumption | Normal (Gaussian) | | | | |
| ANOVA summary | | | | | |
| F | 30.48 | | | | |
| P value | 0.0016 | | | | |
| P value summary | ** | | | | |
| Significant diff. among means (P < 0.05)? | Yes | | | | |
| R squared | 0.9242 | | | | |
| Brown-Forsythe test | | | | | |
| F (DFn, DFd) | 1.477 (2, 5) | | | | |
| P value | 0.3134 | | | | |
| P value summary | ns | | | | |
| Are SDs significantly different (P < 0.05)? | No | | | | |
| Bartlett's test | | | | | |
| Bartlett's statistic (corrected) | | | | | |
| P value | | | | | |
| P value summary | | | | | |
| Are SDs significantly different (P < 0.05)? | | | | | |
| ANOVA table | SS | DF | MS | F (DFn, DFd) | P value |
| Treatment (between columns) | 0.7425 | 2 | 0.3713 | F (2, 5) = 30.48 | P=0.0016 |
| Residual (within columns) | 0.0609 | 5 | 0.01218 | | |
| Total | 0.8034 | 7 | | | |
| Data summary | | | | | |
| Number of treatments (columns) | 3 | | | | |
| Number of values (total) | 8 | | | | |

Fig. 6D - post-hoc

| | |
|---|---|
| Number of families | 1 |
| Number of comparisons per family | 3 |
| Alpha | 0.05 |

| Tukey's multiple comparisons test | Mean diff. | 95.00% CI of diff. | Below threshold? | Summary | Adjusted P Value | |
|---|---|---|---|---|---|---|
| no CFA vs. +CFA | -0.675 | -0.9860 to -0.3640 | Yes | ** | 0.0021 | A-B |
| no CFA vs. +CFA + A20-I | -0.15 | -0.5091 to 0.2091 | No | ns | 0.4267 | A-C |
| +CFA vs. +CFA + A20-I | 0.525 | 0.2140 to 0.8360 | Yes | ** | 0.0063 | B-C |

| Test details | Mean 1 | Mean 2 | Mean diff. | SE of diff. | n1 | n2 | q | DF |
|---|---|---|---|---|---|---|---|---|
| no CFA vs. +CFA | 1.11 | 1.785 | -0.675 | 0.09558 | 2 | 4 | 9.988 | 5 |
| no CFA vs. +CFA + A20-I | 1.11 | 1.26 | -0.15 | 0.1104 | 2 | 2 | 1.922 | 5 |
| +CFA vs. +CFA + A20-I | 1.785 | 1.26 | 0.525 | 0.09558 | 4 | 2 | 7.768 | 5 |

| Compact letter display | |
|---|---|
| +CFA | A |
| +CFA + A20-I | B |
| no CFA | B |

Fig. S9C - t-test

| | |
|---|---|
| Table Analyzed | CFA 20 Hand-counted vs Computer Counted |
| Column C | E1 Hand-Collected |
| vs. | vs. |
| Column B | WT Hand-Collected |
| Unpaired t test | |
| P value | 0.009 |
| P value summary | ** |
| Significantly different (P < 0.05)? | Yes |
| One- or two-tailed P value? | Two-tailed |
| t, df | t=4.143, df=5 |
| How large is the effect? | |
| Mean of column B | 1.956 |
| Mean of column C | 1.291 |
| Difference between means (C - B) ± SEM | -0.6650 ± 0.1605 |
| 95% confidence interval | -1.078 to -0.2524 |
| R squared (eta squared) | 0.7745 |
| F test to compare variances | |
| F, DFn, Dfd | 2.255, 3, 2 |
| P value | 0.6438 |
| P value summary | ns |
| Significantly different (P < 0.05)? | No |
| Data analyzed | |
| Sample size, column B | 3 |
| Sample size, column C | 4 |

| | |
|---|---|
| Table Analyzed | CFA 20 Hand-counted vs Computer Counted |
| Column E | E1 Computer |
| vs. | vs. |
| Column D | WT Computer |
| Unpaired t test | |
| P value | 0.001 |
| P value summary | *** |
| Significantly different (P < 0.05)? | Yes |
| One- or two-tailed P value? | Two-tailed |
| t, df | t=6.887, df=5 |
| How large is the effect? | |
| Mean of column D | 3.612 |
| Mean of column E | 1.424 |
| Difference between means (E - D) ± SEM | -2.188 ± 0.3177 |
| 95% confidence interval | -3.005 to -1.371 |
| R squared (eta squared) | 0.9046 |
| F test to compare variances | |
| F, DFn, Dfd | 17.61, 3, 2 |
| P value | 0.1084 |
| P value summary | ns |
| Significantly different (P < 0.05)? | No |
| Data analyzed | |
| Sample size, column D | 3 |
| Sample size, column E | 4 |

Fig. S9D - one-way ANOVA

| | |
|---|---|
| Table Analyzed | Zymosan 6 - testing 3511-I and BQPB4 |
| Data sets analyzed | A, B, D |
| Distribution assumption | Normal (Gaussian) |
| ANOVA summary | |
| F | 7.86 |
| P value | 0.0005 |
| P value summary | *** |
| Significant diff. among means (P < 0.05)? | Yes |
| R squared | 0.04917 |
| Brown-Forsythe test | |
| F (DFn, DFd) | [illegible].3668 (2, 304) |
| P value | 0.6932 |
| P value summary | ns |
| Are SDs significantly different (P < 0.05)? | No |
| Bartlett's test | |
| Bartlett's statistic (corrected) | 0.2696 |
| P value | 0.8739 |
| P value summary | ns |
| Are SDs significantly different (P < 0.05)? | No |

| ANOVA table | SS | DF | MS | F (DFn, DFd) | P value |
|---|---|---|---|---|---|
| Treatment (between columns) | 18.87 | 2 | 9.434 | F (2, 304) = 7.860 | P=0.0005 |
| Residual (within columns) | 364.9 | 304 | 1.2 | | |
| Total | 383.8 | 306 | | | |

| | |
|---|---|
| Data summary | |
| Number of treatments (columns) | 3 |
| Number of values (total) | 307 |

Fig. S9D - post-hoc

| | |
|---|---|
| Number of families | 1 |
| Number of comparisons per family | 3 |
| Alpha | 0.05 |

| Tukey's multiple comparisons test | Mean diff. | 95.00% CI of diff. | Below threshold? | Summary | Adjusted P Value | |
|---|---|---|---|---|---|---|
| +/+ vs. 3511-I | 0.4354 | 0.09885 to 0.7719 | Yes | ** | 0.0071 | A-B |
| +/+ vs. BQPB4 | 0.5739 | 0.1906 to 0.9572 | Yes | ** | 0.0014 | A-D |
| 3511-I vs. BQPB4 | 0.1385 | -0.2606 to 0.5377 | No | ns | 0.6926 | B-D |

| Test details | Mean 1 | Mean 2 | Mean diff. | SE of diff. | n1 | n2 | q | DF |
|---|---|---|---|---|---|---|---|---|
| +/+ vs. 3511-I | 2.012 | 1.576 | 0.4354 | 0.1429 | 132 | 106 | 4.309 | 304 |
| +/+ vs. BQPB4 | 2.012 | 1.438 | 0.5739 | 0.1628 | 132 | 69 | 4.987 | 304 |
| 3511-I vs. BQPB4 | 1.576 | 1.438 | 0.1385 | 0.1695 | 106 | 69 | 1.156 | 304 |

| Compact letter display | |
|---|---|
| +/+ | A |
| 3511-I | B |
| BQPB4 | B |

### Table S8: Statistics for Figure 7.

Fig. 7a - t-test

| | |
|---|---|
| Table Analyzed | Data 1 |
| Column B | Data Set-B |
| vs. | vs. |
| Column A | Data Set-A |
| Unpaired t test | |
| P value | 0.0112 |
| P value summary | * |
| Significantly different (P < 0.05)? | Yes |
| One- or two-tailed P value? | Two-tailed |
| t, df | t=3.611, df=6 |
| How large is the effect? | |
| Mean of column A | 100.7 |
| Mean of column B | 49.2 |
| Difference between means (B - A) ± SEM | -51.47 ± 14.25 |
| 95% confidence interval | -86.34 to -16.59 |
| R squared (eta squared) | 0.6849 |
| F test to compare variances | |
| F, DFn, Dfd | 1.181, 2, 4 |
| P value | 0.7904 |
| P value summary | ns |
| Significantly different (P < 0.05)? | No |
| Data analyzed | |
| Sample size, column A | 3 |
| Sample size, column B | 5 |

Fig. 7B - t-test

| | |
|---|---|
| Table Analyzed | Data 1 |
| Column B | Data Set-B |
| vs. | vs. |
| Column A | Data Set-A |
| Unpaired t test | |
| P value | 0.0007 |
| P value summary | *** |
| Significantly different (P < 0.05)? | Yes |
| One- or two-tailed P value? | Two-tailed |
| t, df | t=5.049, df=9 |
| How large is the effect? | |
| Mean of column A | 73.8 |
| Mean of column B | 25.17 |
| Difference between means (B - A) ± SEM | -48.63 ± 9.633 |
| 95% confidence interval | -70.42 to -26.84 |
| R squared (eta squared) | 0.739 |
| F test to compare variances | |
| F, DFn, Dfd | 2.603, 4, 5 |
| P value | 0.3223 |
| P value summary | ns |
| Significantly different (P < 0.05)? | No |
| Data analyzed | |
| Sample size, column A | 5 |
| Sample size, column B | 6 |
